# Regulation of neuronal mRNA splicing and Tau isoform ratio by ATXN3 through deubiquitylation of splicing factors

**DOI:** 10.1101/711424

**Authors:** Andreia Neves-Carvalho, Sara Duarte-Silva, Joana M. Silva, Liliana Meireles-Costa, Daniela Monteiro-Fernandes, Joana S. Correia, Beatriz Rodrigues, Sasja Heetveld, Bruno Almeida, Natalia Savytska, Jorge Diogo Da Silva, Andreia Teixeira-Castro, Ioannis Sotiropoulos, Ana Luísa Carvalho, Peter Heutink, Ka Wan Li, Patrícia Maciel

## Abstract

The ubiquitylation/deubiquitylation balance in cells is maintained by deubiquitylating enzymes, including ATXN3. The precise role of this protein, mutated in spinocerebellar ataxia type 3 (SCA3), remains elusive, as few substrates for its deubiquitylating activity were identified. Here, we characterized the ubiquitome of neuronal cells lacking ATXN3, and found altered polyubiquitylation in a large proportion of proteins involved in RNA metabolism, including splicing factors. Using transcriptomic analysis and reporter minigenes we confirmed that splicing was globally altered in these cells. Among the candidate targets of ATXN3 was SRSF7 (9G8), a key regulator of MAPT (Tau) exon 10 splicing. Loss-of-function of ATXN3 led to reduced SRSF7 levels and a deregulation of MAPT exon 10 splicing, resulting in a decreased 4R/3R-Tau ratio. Similar alterations were found in cellular models of expanded polyQ ATXN3 and SCA3 patient brains, pointing to a relevant role of this mechanism in SCA3, and establishing a previously unsuspected link between two key proteins involved in different neurodegenerative disorders.

## 1. Introduction

Ubiquitin signaling is now recognized as a fundamental molecular mechanism tightly regulating a broad range of intracellular events [1–3]. Ubiquitylation is a highly dynamic biochemical modification in which an ubiquitin (Ub) moiety is attached to a target protein. This process is catalyzed by the sequential actions of an Ub-activating enzyme (E1), Ub-conjugating enzymes (E2) and Ub-protein ligases (E3) that bind Ub to different lysine (K) residues in the substrate, resulting in mono or poly Ub chains (Reviewed in [4]). Ubiquitylated substrates are then recognized by proteins containing ubiquitin binding domains and directed to different fates. For example, K48-linked polyUb usually targets proteins for proteasomal degradation [4], while K63-linked polyUb regulates protein activation, subcellular localization or degradation in the lysosome (autophagy) and is known to be relevant for DNA repair (Reviewed in [5]). Ubiquitylation and proteolysis by the Ubiquitin-proteasome pathway (UPP) are now well-known as important mechanisms in the nervous system as this proteolytic pathway is known to degrade misfolded or short-lived regulatory proteins (Reviewed in [1, 6]). Impairment of the UPP has been connected to several neurodegenerative diseases such as Alzheimer’s (Reviewed in [7]), Parkinson’s (Reviewed in [8]) and Huntington’s (Reviewed in [9]) diseases. Like most post-transcriptional modifications, ubiquitylation is a reversible signal and is counterbalanced by the activity of deubiquitylating (DUB) enzymes, which remove Ub units and play an important role in the modulation of proteins by the proteasome as well as the autophagic pathway [10–12], and in Ub signaling in general. Several pieces of evidence have suggested that Ub may also regulate splicing: (i) Ub and Ub-like proteins have been shown to copurify with splicing complexes [13, 14], (ii) ubiquitylated splicing factors have been identified in proteomic screenings [15, 16], and (iii) several functional domains related to the Ub pathway have been identified in important spliceosome proteins, such as the Jab1/MPN domain of the essential U5 SnRNP component Prp8 [17–20]. The action of DUBs has a major impact on the ubiquitylated proteome (also known as ubiquitome). Ataxin-3 (ATXN3) is a protein with DUB activity known to be involved in Spinocerebellar ataxia type 3 (SCA3)/ Machado Joseph disease (MJD), a neurodegenerative disorder of adult onset caused by the expansion of a polyglutamine (polyQ) tract in this protein. Interestingly, in addition to being involved in global protein quality control responses [21, 22], normal ATXN3 appears to participate in cell adhesion and in the organization of cytoskeleton network [23–26], to be involved in transcriptional regulation and DNA repair [27–29], and to be required for neuronal differentiation [25]. Nevertheless, the relative relevance of these functions and the physiological role of this highly conserved but, apparently, genetically non-essential protein remain to be clarified, and its substrates are also not well characterized, particularly in neurons. Given the known relevance of the UPP in neuronal function and its link to nervous system diseases, we sought to clarify the precise role of ATXN3 in neuronal cells. To identify potential candidates for the DUB activity of this protein, we characterized changes in the ubiquitome of neuronal cells lacking ATXN3 (ATXN3^shRNA^ cells) by mass-spectrometry, and identified for the first time a role for ATXN3 in RNA metabolism. We found a global perturbation of splicing in the absence of this protein, among which a mis-splicing of tau exon 10 which perturbs the 4R/3R tau ratio. Our findings describe a novel mechanism in neuronal cells of partial loss of function of ATXN3 upon expansion of a polyQ tract, that may contribute for SCA3 pathogenesis, and links two key proteins involved in different neurodegenerative disorders.

## 2. Materials and Methods

*Cell lines and primary culturesSH-SY5Y* <u>cell cultures: SH-SY5Y</u> human female neuroblastoma cell line (ATCC number CRL-2266) was cultured in Dulbecco’s modified eagle medium: nutrient mixture (DMEM)/F-12 supplemented with 10% (v/v) fetal bovine serum (FBS) (Biochrom), 2mM glutaMAX (Invitrogen), 100 U/mL penicillin (Invitrogen), 100 μg/mL streptomycin (Invitrogen) and 25 ng/mL puromycin (Sigma Aldrich). Medium was changed every 2 days. Cells were transfected with a shRNA sequence targeting ATXN3 or a scrambled shRNA sequence as described elsewhere *[25]*. Differentiation of stably infected cell lines was induced by 0.1 μM all-trans-retinoic acid (RA) (Sigma Aldrich) in opti-MEM (Invitrogen) supplemented with 0.5% FBS for 7 days. Medium was replaced every 2 days. <u>Primary cultures of cerebellum:</u> cerebellar neuron cultures were prepared from P7 Wistar rats. Briefly, upon dissection, cerebella were submitted to a trypsin-based enzymatic digestion followed by mechanical dissociation. Isolated cells were plated on poly-d-lysine (Sigma Aldrich) and lysine (Sigma Aldrich) pre-coated 6-well plates at a density of 1.25×106 cells/mL using Neurobasal A (Invitrogen) supplemented with 1x B27 (Invitrogen), 0.1 mg/mL Kanamycin (Invitrogen), 1 mM Glutamax I (Invitrogen) and 10 ng/mL bFGF (Invitrogen) for 7 days. Half of the medium was changed every 4 days.

<u>Low-density cortical cultures (Banker cultures).</u> For imaging experiments, low-density cultures were prepared as previously described [30]. Cortices were dissected, treated with 0.06% (wt/vol) trypsin for 15 min at 37°C and dissociated in Ca^2+^ and Mg^2+^-free Hank’s balanced salt solution (HBSS; in mM: 5.36 KCl, 0.44 KH2PO4,137 NaCl, 4.16 NaHCO3, 0.34 Na2HPO4.2H2O, 5 glucose, 1 sodium pyruvate, 10 HEPES and 0.001% phenol red). Cells were then plated in neuronal plating medium onto poly-D-lysine coated coverslips (0.1 mg/ml) at the final density of 3 × 105 cells per 60 mm culture dishes. After 2-4h, coverslips were flipped over an astroglial feeder layer (grown in MEM supplemented with 10% horse serum, 0.6% glucose, and 1% penicillin-streptomycin) and maintained in neuronal culture medium (NBM supplemented with SM1 supplement [1:50 dilution], 0.5 mM glutamine, 0.12 mg/ml gentamycin, 25 μM glutamate). The neuronal cultures were treated with 5 μM cytosine arabinoside at DIV 2-3, to prevent glia overgrowth. The cultures were maintained in humidified incubator at 37°C with 5% CO_2_ and 95% air, and fed once per week with neuronal culture medium without glutamate, by replacing one-third of the medium per dish.

### Neuron transfection

Primary cultures of cortical neurons were transiently transfected using the calcium phosphate transfection protocol adapted from Jiang and colleagues [31] at 10 DIV. Briefly, neurons were treated with 2 mM kynurenic acid in conditioned neuronal culture medium, for 20 min at 37°C, while DNA precipitates were prepared. For DNA precipitate preparation, plasmid DNAs were diluted in Tris-EDTA transfection buffer (in mM: 10 Tris-HCl, 2.5 EDTA, at pH 7.3), followed by the addition of CaCl_2_ solution (2.5 M in 10 mM HEPES) drop-wise to the diluted DNA, to a final concentration of 250 mM CaCl_2_. This mix was added to an equivalent volume of HEPES-buffered saline transfection solution (in mM: 274 NaCl, 10 KCl, 1.4 Na2HPO4, 11 dextrose, 42 HEPES, at pH 7.2), a small fraction at a time (1/8^th^) and the mixture was vortexed gently. The precipitates were then added dropwise to pre-conditioned neurons and the cultures were incubated with the precipitates for 1.5-2h. After incubation, remaining DNA precipitates were dissolved by incubating the neurons with acidified neuron culture medium (in mM: 2 kynurenic acid, ~5 HCl final concentration) for 15-20 min at 37°C. Coverslips were returned to the original astroglial plate. All incubations were performed at 37°C, in a humidified incubator with 5% CO_2_ and 95% air. For imaging experiments, Srsf7 (or the control empty vector pIC111) was cotransfected with GFP-tagged ataxin-3 with 28Q or 84Q, in a total of 5μg of DNA per coverslip. Plasmids were allowed to express for 6 days.

### Immunofluorescence

To label excitatory synapses cortical neurons at DIV16 were fixed with cold 4% PFA / 4% sucrose for 15 minutes, followed by 6 sequential washes with phosphate buffer saline (PBS), permeabilized with 0.25% Triton X-100 during 5min and washed once with PBS. Non-specific staining was blocked with 10% bovine serum albumin (BSA) for 30 min at 37°C. Neurons were incubated overnight at 4°C with chicken polyclonal anti-MAP2 antibody (1:5000; Abcam), rabbit polyclonal anti-GFP antibody (1:500, MBL), mouse monoclonal anti-PSD-95 antibody (1:200; Thermo Scientific) and guinea pig polyclonal anti-VGluT1 antibody (1:5000; Merck Millipore), diluted in 3% BSA. Coverslips were then incubated for 60 min at 37°C with the respective secondary antibodies diluted in 3% BSA: AMCA-conjugated anti-chicken (1:200; Jackson ImmunoResearch), Alexa Fluor^®^ 488 conjugated anti-rabbit (1:500; Invitrogen Molecular Probes); Alexa Fluor^®^ 568 conjugated anti-mouse (1:500; Invitrogen Molecular Probes) and Alexa Fluor® 647 conjugated anti-guinea pig (1:1000; Invitrogen Molecular Probes). Coverslips were mounted with fluorescence mounting media. All solutions were prepared in PBS (in mM: 137 NaCl, 2.7 KCl, 1.8 KH_2_PO_4_, 10 Na_2_HPO_4_•2H_2_O, at pH 7.4).

### Imaging

Fluorescence imaging for puncta analysis was performed using a widefield Zeiss Axio Observer Z1 inverted microscope (Carl Zeiss) equipped with an AxioCam HRm camera and with ZEN blue software (Carl Zeiss). Images for synapse analyses were acquired with a 63x Plan-ApoChromat oil objective (numerical aperture 1.4). For each independent experiment, neurons were cultured and stained simultaneously. Dendrites with similar thickness and appearance were randomly selected using MAP2 and eGFP staining and exposure time was defined to avoid pixel saturation. All experimental conditions within independent preparations were imaged using identical settings.

### Puncta quantification

Images were quantified as previously described [32], using image analysis software FIJI [33], taking advantage of a semi-automatic macro designed for this purpose. The area of interest was randomly selected by using MAP2 and eGFP staining for transfected cells. Dendritic length was measured in the area of interest, using one of the mentioned channels. To quantify the proteins of interest, images were subjected to a user-defined intensity threshold, in order to have defined protein clusters, and user defined background intensity was subtracted in all images. Each protein cluster present in the area of interest was analysed and measurements of intensity, area and number were obtained. Synaptic clusters were selected by overlapping PSD-95 clusters with thresholded VGluT1 clusters. All results were normalized to dendritic length. This analysis was performed blind to the experimental condition.

### Dendritic arborization analysis

Dendritic arborization was studied using the Sholl analysis method. Using Simple Neurite Tracer plug-in of FIJI, dendrites of each transfected neuron labelled with eGFP were identified based on dendritic morphology and on presence of MAP2, and traced in a semi-automatic way. Neurons were then analysed using the concentric Sholl analysis, providing overall analysis of arborization. A minimum of 10 cells per each experimental condition was analysed in a blind manner.

### Pulldown of polyubiquitylated proteins

Cells treated with RA were lysed by sonication on ice in lysis buffer (50 mM Tris-HCl pH 7.5, 0.15 M NaCl, 1mM EDTA, 1% NP-40, 10% Glycerol, complete protease inhibitors (Roche) and 50 μM UB/UBl protease inhibitor PR-619 (LifeSensors). After lysis, 2 mg of total protein extract was incubated with 100 μL of pre-equilibrated Agarose-TUBEs (LifeSensors), overnight at 4°C on a rocking platform. Sedimented beads were washed 3 times with washing buffer (20 mM Tris pH 8.0, 0.15 M NaCl, 0.1% Tween-20) before being eluted with 1x SDS sample buffer (62.5mM Tris-HCl pH 6.8, 10% glycerol, 2% SDS, Bromophenol Blue). Eluted proteins were immediately boiled at 98°C for 15min and ran in a 10% SDS-PAGE gel. After incubation with the FK2 anti-ubiquitin primary antibody (1:2000, Millipore) overnight at 4°C, membranes were incubated with secondary antibody for 1 hour at room temperature (anti-mouse, 1:10.000, Bio-Rad). Antibody binding was detected by chemiluminescence (Clarity kit, Bio-Rad).

### Digestion of proteins from preparative 1D-PAGE gel

The 1D PAGE LC-MS/MS approach was used for protein identification as previously described [34]. Eluted proteins were separated using 1.5 mm and 10% SDS-PAGE gels. The quality of purification was controlled by Coomassie Brilliant Blue g-250 (Sigma) staining before MS analysis. Gel image was acquired the Gel Doc™ EZ system (Bio-rad). After Coomassie staining, all the visible blue-stained protein spots were manually excised from the gel. The gel pieces were destained overnight at room temperature using 50% acetonitrile in 25 mM ammonium bicarbonate buffer, pH 8.5, and then dehydrated with 100% acetonitrile. The shrunken pieces were then reswollen in 50 mM ammonium bicarbonate buffer, dehydrated in 100% acetronitrile and dried in a speedvac® concentrator (Savant) for 30 min. The gel pieces were rehydrated in 60 μL of 20 μg/mL Trypsin (Promega) in 50 mM ammonium bicarbonate solution and incubated for 2h at 55°C. The gel pieces were then incubated with 0.1% trifluoroacetic acid in 50% acetonitrile for 20 min at room temperature in order to extract the remaining peptides from the gel. The tryptic peptides were dried in a speedvac for 2 h.

### Liquid chromatography-tandem mass spectrometry (LC-MS/MS)

After re-dissolution in 17 μL 0.1% acetic acid, samples were separated on a capillary C18 column using a nano LC-ultra 1D plus HPLC system (Eksigent) and analyzed on-line with an electrospray LTQ-Orbitrap Discovery mass spectrometer (Thermo Fisher Scientific). MS/MS spectra were searched against a human database (uniprot_sprot, 2010_01) with the ProteinPilot^™^ software (version 3.0; AB-sciex) using the Paragon^™^ algorithm (version 3.0.0.076) as the search engine. The detected protein threshold (unused protscore (confidence)) in the software was set to 0.10 to achieve 20% confidence and the proteins identified were grouped to minimize redundancy. Peptides with “unused” values < 2 have low confidence and were excluded from analysis. The “unused” value is defined in the handbook of ProteinPilot as a sum of peptide scores from all the non-redundant peptides matched to a protein. Peptides with confidence of ≥ 99% would have a peptide score of 2. Tryptic peptides shared by multiple proteins were assigned to the winner protein.

### RNA extraction and Array hybridization

Total RNA was isolated from ATXN3^shRNA^ and SCR^shRNA^ cells using an miRNeasy mini kit (Qiagen) and quality assessment was achieved using RNA 6000 Nano labchip (Bioanalyzer, Agilent) and by a Nanodrop spectrophotometer (Thermo). Total RNAs RIN values were between 8.7 and 9.3. Affymetrix Human Transcriptome Array 2.0 ST arrays were hybridized according Affymetrix recommendations using the Ambion WT protocol (Life technologies, France) and Affymetrix labelling and hybridization kits. Raw data, transcript data and exon data were controlled with Expression console (Affymetrix).

### Microarray data analysis

Affymetrix Human Transcriptome Array 2.0 ST dataset analysis was performed using the GenoSplice technology (www.genosplice.com). Data was normalized using quantile normalization. Background corrections were made with antigenomic probes and probes were selected according to their %GC, cross-hybridization status and potential overlap with repeat region as previously described [35–37]. Only probes targeting exons and exon-exon junctions annotated from FAST DB® transcripts (release fastdb_2013_2) were selected [38, 39]. Only probes with a DABG P value ≥0.05 in at least half of the arrays were considered for statistical analysis [35–37]. Only genes expressed in at least one compared condition were analyzed. To be considered to be expressed, the DABG P-value had to be ≤0.05 for at least half of the gene probes. We performed an unpaired Student’s t-test to compare gene intensities between ATXN3^shRNA^ and SCR^shRNA^ cells. Genes were considered significantly regulated when fold-change was ≥1.5 and P-value ≤0.05 (unadjusted P-value). Analysis at the splicing level was first performed taking into account only exon probes (‘EXON analysis) in order to potentially detect new alternative events that could be differentially regulated (i.e., without taking into account exon-exon junction probes). Analysis at the splicing level was also performed by taking into account exon-exon junction probes (‘SPLICING PATTERN analysis) using the FAST DB® splicing pattern annotation (i.e., for each gene, all possible splicing patterns were defined and analyzed. All types of alternative events can be analyzed: alternative first exons, alternative terminal exons, cassette exon, mutually exclusive exons, alternative 5’ donor splice site, alternative 3’ acceptor splice sites and intron retention). EXON and SPLICING PATTERN analyses were performed using unpaired Student’s t-test on the splicing-index as previously described [35–37]. Results were considered statistically significant for unadjusted P-values ≤ 0.05 and fold-changes ≥ 1.5 for SPLICING PATTERN analysis and unadjusted P-values ≤ 0.05 and fold-changes ≥ 2.0 for EXON analysis. Gene Ontology (GO), KEGG and REACTOME analyses of differentially regulated genes were performed using DAVID [40].

### Plasmid purification

Hybrid minigene reporter plasmids AdML and α-globin [41, 42] were kindly provided by Prof. Juan Valcárcel (Centre de Regulació Genòmica, Barcelona). Top10 competent cells (Invitrogen) were transformed with 100 ng of plasmid DNA, according to the recommended protocol. Briefly, the cells were incubated with the constructs on ice for 30 min followed by heat shock at 42°C for 1 min. After incubation on ice for 2 min, 500 μL of LB medium was added to the cell vial and incubated at 150 rpm for 60 min at 37°C. Cultures were grown overnight at 37°C in LB/ampicillin plates. The next day, one colony was inoculated in LB/ampicillin (100 mg/mL) at 37°C overnight. Plasmid extraction was carried out using the ZR Plasmid Miniprep^™^ (Zymo Research) according with the manufacturer’s protocol. DNA concentration was determined using Nanodrop (Alfagene) and integrity verified by running 200 ng in an agarose gel.

### Cell transfection

4×10^5^ cells per well were plated in gelatin-coated 6 well plates and incubated for 24 h. Before transfection, the culture medium was changed to DMEM/F-12-AA without antibiotics and supplemented with 5% FBS. Cells were transfected with 200 ng of the reporter plasmids using Lipofectamine^®^ 2000 Transfection Reagent (Invitrogen) according to the manufacturer’s instructions. Briefly, reporter plasmid minigenes and the transfection reagents were appropriately diluted in Opti-MEM medium separately and incubated for 5 min at room temperature. The mixed reagents were then incubated at room temperature for 20 min allowing the formation of transfection complexes. The cells were then incubated for 24 h with the transfection mix.

### Semi-quantitative PCR

PCR amplification of AdML and *α*-globin reporter genes was carried out using Taq DNA Polymerase (Thermo Fisher Scientific) following the manufacturer’s protocol. The cycling conditions were: 95°C for 5 min followed by 24 cycles of denaturing at 95°C for 1 min, annealing at 60°C for 45 sec, extension at 72°C for 1 min and final extension at 72°C for 5 minutes. The PCR product was ran in a 2% agarose gel. Gel analyses and splicing efficiency calculations were performed using Image Lab software (Bio-Rad).

### Sub-cellular fractionation

RA-treated pelleted cells were resuspended in ice RSB buffer (10 mM Tris-HCl pH 7.4, 10 M NaCl) and incubated on ice 3 min. After centrifugation for 5 min, 4000 rpm at 4°C, the pellet was resuspended in RSBG40 buffer (10 mM Tris-HCl pH 7.4, 10 mM NaCl, 3 mM MgCl2, 10% glycerol, 0.5% NP-40, 0,5 mM DTT) and centrifuged for 3 min, 7000 rpm at 4°C. The supernatant was collected as cytoplasmic fraction and stored at −80°C. The nuclear pellet was resuspended in 50 μL B1 buffer (20 mM Tris-HCl pH 7.9, 75 mM NaCl, 5% glycerol, 0.5 mM EDTA, 0.85 mM DTT, 0.125 mM PMSF) and 450 μL B2 buffer (20 mM HEPES pH 7.6, 300 mM NaCl, 0.2 mM EDTA, 1 mM DTT, 7.5 mM MgCl_2_, 1 M Urea, 1% NP-40). Samples were vortexed for 5 sec, incubated on ice for 10 min and centrifuged for 5 min, 15000 rpm at 4°C. The supernatant was collected as nuclear fraction and stored at −80°C.

### Protein synthesis inhibition and proteasome inhibition

RA-treated SH-SY5Y cells were incubated with 5 μM cicloheximide (Merck) during 20, 60 and 120 minutes. For each condition, the half-life (t1/2) of SRSF7 was determined using the N(t) = N0 * (0.5) ^ t/t1/2 equation, where N0 is the initial protein amount (OD normalized to tubulin) and N(t) is the protein amount at t = 60 min. For proteasome inhibition, RA-treated cells were incubated with 5 μM MG132 (Calbiochem) for 24h prior to lysis.

### High-throughput high-content functional imaging

SH-SY5Y cells were seeded at a density of 4×10^3^ cells/well in flat bottom 96-well plates previously coated with Matrigel (BD, Biosciences), and 10 μM all-trans-retinoic acid (Sigma Aldrich) was added the day after plating in DMEM-F12 with 1% FBS, to induce differentiation. After 5 days, cells were washed in DMEM/F-12 and incubated with 50 ng/mL BDNF (Prepotech) in DMEM/F-12 without serum for 3 days. Cells were then labeled for β-III tubulin (1:1000, R&D Systems), scanned at different locations of each well and the quantitative analysis of neurite length were automatically done using the automatic imaging system Thermo Scientific Cellomics^®^ ArrayScan^®^ VTI.

### Immunoprecipitation

RA-treated cells were washed in ice-cold PBS and lysed by sonication on ice in NP-40 buffer. Aliquots were taken from protein extracts as 10% inputs. 1 mg of total protein was pre-cleared for 3h at 4°C by incubation with glutathione-coupled sepharose beads (GE Healthcare) previously equilibrated in Wash buffer 1 (50 mM Tris-HCl pH 7.5, 150 nM NaCl, 1% NP-40, 1x protease inhibitors (Roche)) for 3 times, 10 min at 4°C. Beads were then centrifuged and the supernatant was incubated O/N at 4°C with 50 μL equilibrated beads. After centrifugation, the supernatant was discarded and the beads were washed 2 times with Wash buffer 1 for 10 min at 4°C. Beads were then washed twice with Wash buffer 2 (50 mM Tris-HCl pH 7.5, 500 mM NaCl, 0.1% NP-40) for 10 min at 4°C, and once with Wash buffer 3 (50 mM Tris-HCL pH 7.5, 0.1% NP-40) for 20 min at 4°C. The supernatant was discarded and the bound proteins were eluted with 1x Laemmli buffer, boiled for 5 min at 98°C and run in a 10% SDS-PAGE gel.

### Proximity Ligation Assay (PLA)

SH-SY5Y cells were cultured onto polyD-lysine coated cover slips in 24-well plates using DMEM media as indicated above. PLA protocol was performed as described in [43] and following manufacturer instructions with adaptations. Cells were fixed with 4% PFA, and permeabilized with 0.5% Triton in PBS at room temperature. Blocking was performed at 37°C in a humidity chamber for 1 hour, using Duolink^®^ PLA Blocking Solution (Sigma). Primary antibodies were incubated O/N at 4°C followed by two washing steps of 5 minutes with Wash Buffer A (Sigma) at room temperature. PLUS and MINUS PLA probes were diluted 1:5 in the Duolink Antibody Diluent (Sigma), added to the cover slips and incubated for 1h at 37°C in a humidity chamber. Washing was performed for 5 minutes two times with Washing Buffer A at room temperature followed by incubation with Ligase solution (Sigma) for 30 minutes at 37°C in a humidity chamber. After washing two times during 2 minutes with Washing Buffer A at room temperature, cover slips were incubated with Amplification solution (Sigma) for 100 minutes at 37°C in a humidity chamber protected from light. Then, cover slips were washed two times during 10 minutes with Washing Buffer B (Sigma) at room temperature followed by an additional washing step of 1 minute using 100X diluted Washing Buffer B. Cover slips were mounted on a glass slide with mounting media and DAPI and cells analyzed by confocal microscopy.

### Immunoblotting

RA-treated cells were pelleted and frozen in liquid nitrogen. For cellular extracts 50 μg of total protein isolated in NP-40 buffer (150 mM NaCl, 50 mM Tris-HCl pH 7.6, 0.5% NP-40, protease inhibitors (Roche)) was resolved in 10% SDS-PAGE gels and then transferred to a nitrocellulose membrane. After incubation with the primary antibodies: Tau-5 (1:2000, Abcam), Tau-4R (1:1000, Millipore), Tau-3R (1:2000, Millipore), Tau (E178) (1:1000, Abcam), Histone H3 (1:10.000, Abcam), SFRS7(9G8) (1:1000, kindly provided by Dr. James Stévine), Actin (1:200, DSHB) overnight at 4°C, membranes were incubated with secondary antibodies for 1 hour at room temperature (anti-rabbit or anti-mouse, 1:10.000, Bio-Rad, anti-goat, 1:7500, Pierce). Antibody binding was detected by chemiluminescence(Clarity kit, Bio-Rad).

### qRT-PCR

1 μg of total RNA purified from differentiated cells was reversed transcribed using the One-step SuperScript kit (Bio-Rad). The qRT-PCR reaction was performed in a CFX96 Real-time PCR detection system (Bio-Rad) using the Quantitec SYBR Green kit (Qiagen). Gene expression was normalized to HMBS or B2M levels within each sample. Results are presented as fold change.

### Quantification and statistical analysis

Continuous variables with normal distributions (Shapiro-Wilk test p>0.05) were analyzed with the Student t-test or two-way ANOVA. Non-parametric Mann-Whitney U-test was used when variables were non-continuous or when a continuous variable did not present a normal distribution (Shapiro-Wilk test p<0.05). All statistical analyses were performed using SPSS 22.0 (SPSS Inc., Chicago, IL). A critical value for significance of p<0.05 was used throughout the study. For real-time quantitative PCR data, the same approach was used and results were presented using the ΔΔCt method, as described before [44]. Graphs and statistical analyses of the Low-density cortical cultures were performed using GraphPad Prism 6 software. Outliers were identified using raw experimental data with the Grubbs method. Data are presented as mean ± standard error of the mean (SEM) of three or more experiments, performed in independent preparations, and statistical differences were analysed as indicated in the figure captions.

## 3. Results

### 3.1 Changes in the ubiquitome of cells lacking ATXN3 suggest a link to RNA metabolism

With the goal of identifying substrates of the DUB activity in neuronal cells, we searched for changes in the ubiquitome of neuronally differentiated SH-SY5Y cells lacking ATXN3 (ATXN3^shRNA^ cells–Figure 1a-b, Figure S1a) [25], taking advantage of a methodology that combines capture of the whole spectrum of polyubiquitylated proteins in a cell extract using an enrichment by Tandem Binding Ubiquitin Entities (TUBEs) followed by mass spectrometry analysis (TUBEs-LC-MS/MS) [45]. Figure 1a summarizes the steps followed for the purification and identification of the polyubiquitylated proteins in ATXN3^shRNA^ and scrambled control shRNA (SCR^shRNA^) cells. The integration of the data and the comparison between the proteins identified in ATXN3^shRNA^ versus SCR^shRNA^ cells resulted in a list of proteins with altered polyubiquitylation in cells lacking ATXN3; among these are potential targets of the DUB activity of ATXN3, i.e., putative ATXN3 substrates. In each pulldown experiment, around 1200-1300 proteins were identified. When the results of all the independent experiments were merged, we observed that many of these proteins were only sporadically detected across the different experiments; for the remaining analysis, these proteins were excluded, reducing the list to 615 consistently identified proteins. From these, 193 proteins showed altered levels in the polyubiquitylated fraction in ATXN3^shRNA^ cells when compared to the SCR^shRNA^ controls (p<0.05) (Table S1). The majority of these polyubiquitylated proteins were either not detected (44.04%) or showed decreased levels (24.35%) in ATXN3^shRNA^ cells when compared to the controls. These proteins can be grouped into 10 functional networks, the most representative being related to: (i) Gene expression, DNA replication, recombination and repair (17.7%), (ii) RNA post-transcriptional modification (13.3%), (iii) Molecular transport, RNA trafficking (10.8%), (iv) Cell death and survival (8.9%), and (v) Organ morphology (8.9%) (Figure 1b). The fact that a significant proportion of the proteins with altered polyubiquitylation in ATXN3^shRNA^ cells are RNA metabolism-related proteins (25%), around 8% of them being splicing factors (Table 1), suggested to us that ATXN3 could be playing a role in mRNA splicing, a previously undescribed role for this DUB.

**Figure 1.**
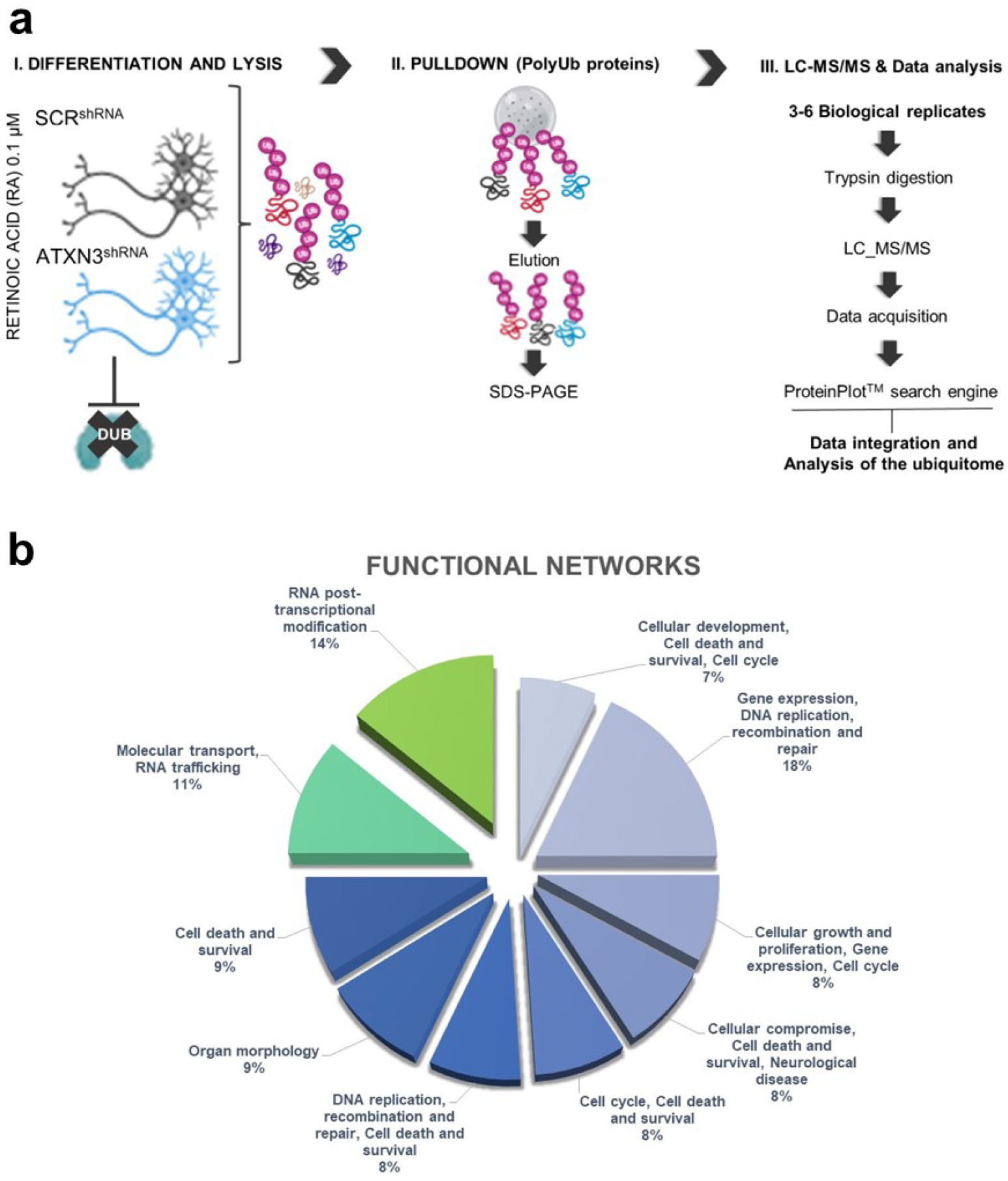
Experimental design – purification of polyubiquitylated proteins using TUBEs. **(a-I)** RA-treated SH-SY5Y cells (with silenced ATXN3 or not) were lysed and protein extracts were incubated with TUBEs. **(a-II)** TUBEs-captured proteins were recovered with Laemmli buffer and analyzed by comassie blue staining of the polyubiquityated proteins purified using TUBEs run in a 1-D SDS-PAGE gel or western-blot with anti-ubiquitin FK2 antibody. **(a-III)** Polyubiquitylated proteins were trypsin-digested and identified by LC-MS/MS. **(b)** After acquisition, data were processed and integrated in functional networks.

**Table 1.** RNA metabolism related proteins with altered ubiquitylation in cells lacking ATXN3.

| RNA METABOLISM PROTEINS WITH ALTERED UBIQUITYLATION IN ATXN3<sup>shRNA</sup> CELLS |  |  |  |
| --- | --- | --- | --- |
| UPREGULATED | p value | DOWNREGULATED | p value |
| RBM10 (Rna-binding protein 10) | 4,87E-19 | DDX41 (DEAD-box protein abstract variant) | 6,22E-06 |
| SNRPB2 (U2 small nuclear ribonucleoprotein B) | 3,26E-11 | DDX56 (ATP-dependent RNA helicase) | 2,38E-05 |
| SFRS2 (Splicing factor arginine/serine-rich 2) | 5,36E-11 | LUC7L (Putative RNA-binding protein Luc-like 1) | 5,00E-04 |
| RBM8A (RNA-binding protein 8A) | 1,56E-06 | SFRS7 (Serine/arginine-rich splicing factor) | 8,00E-04 |
| PRPF8 (Pre-mRNA-processing-splicing factor 8) | 1,00E-03 | DDX10 (Probable ATP-dependent RNA helicase) | 1,00E-02 |
| SFRS5 (Serine/arginine-rich splicing factor 5) | 5,00E-03 | HNRNPA1P10 (Heterogeneous nuclear ribonucleoprotein N) | 2,00E-02 |
| SNRPN (Small nuclear ribonucleoprotein-associated protein N) | 5,00E-03 | FUS (RNA-binding protein) | 3,00E-02 |
| SRSF9 (Serine/arginine-rich splicing factor 9) | 7,00E-03 | RBMX (Heterogeneous nuclear ribonucleoprotein G) | 4,00E-02 |
| RBM12B (RNA-binding protein 12B) | 2,00E-02 | SRRT (Serrate RNA effector molecule) | 4,00E-02 |
| AQR (Intron-binding protein aquarius) | 2,00E-02 |  |  |
| PRPF40A (Pre-mRNA-processing factor 40 homolog A) | 2,00E-02 |  |  |
| HNRNPK (Heterogeneous nuclear ribonucleoprotein K) | 3,00E-02 |  |  |

### 3.2. Cells lacking ATXN3 show global perturbation of splicing

We hypothesized that absence of ATXN3 could lead to a deregulation of the pre-mRNA splicing process. To address this, we used two hybrid minigene reporter plasmids: the AdML minigene, representing constitutive/strong splicing events, and the α-globin minigene, for which the alternative splicing (exon skipping) is indicative of the performance of regulatory splicing factors such as hnRNP and SF proteins [41, 42]. As shown in Figure 2a-c, knockdown of ATXN3 significantly altered the processing of the two splicing reporters, suggesting a general deregulation of splicing in the absence of this protein. Consistent with a global deregulation of splicing in the absence of ATXN3, microarray analysis (Figure 2d) using specific arrays containing additional probes for exon/exon junctions–which have been proved to have a high degree of coherence comparing to RNA-seq – we confirmed that 1993 genes (43%), from the 7450 differentially expressed probes in ATXN3^shRNA^ cells, presented differentially regulated alternative splicing events (Table S2). These genes mainly encode components with function in protein degradation systems, cell adhesion, axon guidance and various signaling pathways (Table S3), the majority of splicing events being decreased when compared to control cells (Table S4). The most affected alternative splicing event types were exon cassette usage (34%), complex splicing (20%) and intron retention (15%) (Figure 2e). To increase our confidence on the microarray data, we validated by qRT-PCR 21 out of 24 splicing events, representing different classes of splicing events (Figure S2a-b). Alternative terminal exon usage, which can be mechanistically linked to alternative splicing, was also seen in 13% of the cases (Figure 2e). Being a key mechanism in the regulation of gene expression, defects in these regulatory processes may affect a multiplicity of cellular functions related to disease. Of notice, the cellular pathways related to synapse function were enriched among the differentially spliced genes: (i) Synapse vesicle, (ii) Synapse, (iii) Postsynapse and (iv) Glutamatergic synapse (Table 2).

**Figure 2.**
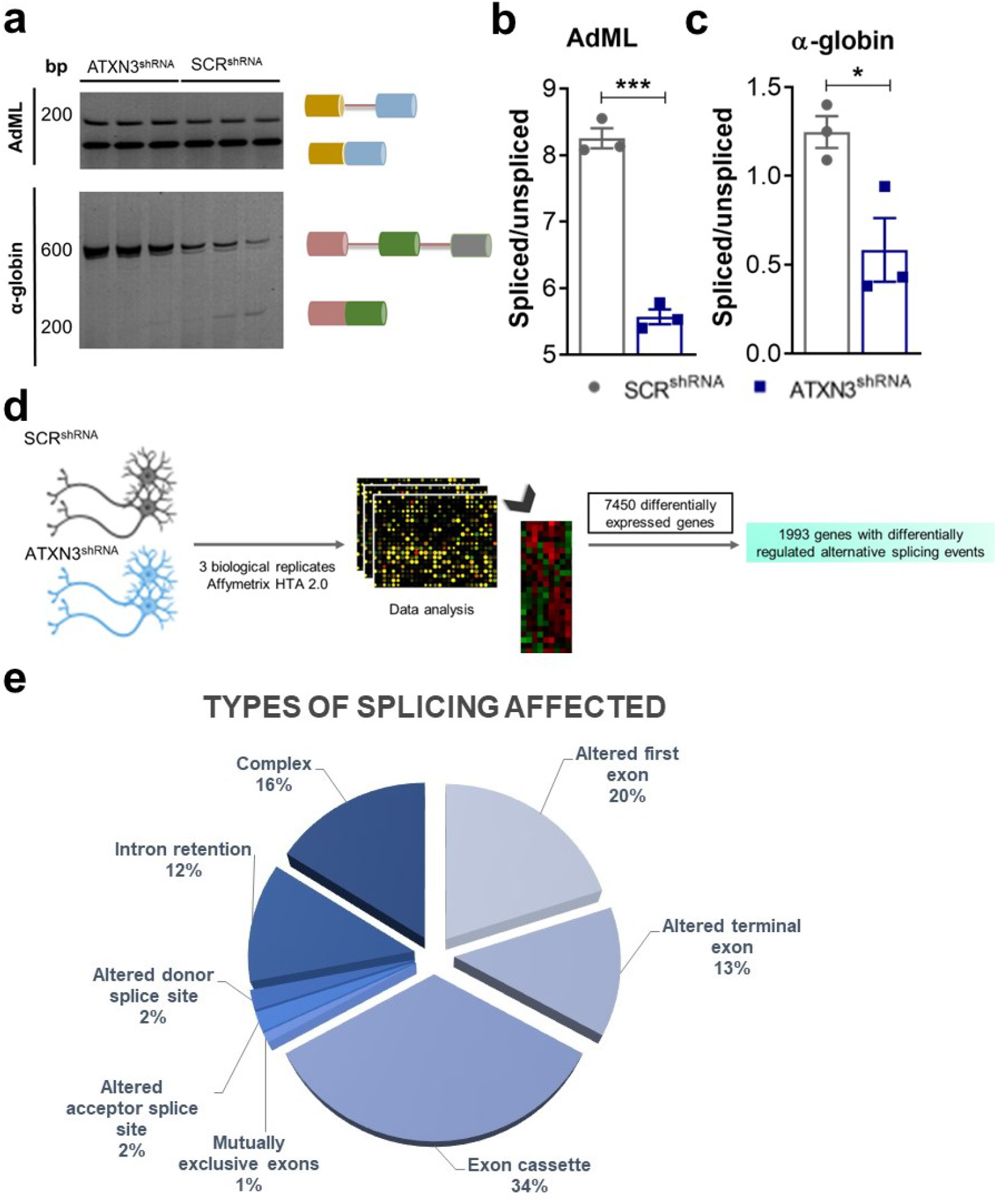
Efficacy of RNA processing assessed by splicing reporter minigenes in ATXN3^shRNA^ cells. **(a-c)** Semi-quantitative analysis of minigene’ alternative splicing showed a decreased efficiency of splicing in ATXN3^shRNA^ cells. Schemes for the splicing products are indicated on **(a)**. Exons are represented as colored cylinders and introns by red lines. **(a, b)** The AdML minigene contains one intron, giving rise to two bands: the upper band corresponds to the unspliced transcript, the lower band to the spliced product. **(a, c)** The α-globin minigene contains two introns and a set of G triplets in intron 2 that promote the recognition of the 5’ splice site leading to skipping of exon 2. **(d) Experimental design** – microarray analysis of alternative splicing using the Affymetrix Human Transcriptome Array 2.0 ST to assess perturbation of global splicing patterns in ATXN3^shRNA^ cells. **(e)** Distribution of the differentially regulated alternative splicing events in ATXN3^shRNA^ cells. n≥3 independent biological replicates in all experiments. p<0,05 was taken as the cut-off. Student’s t-test was used for comparisons. Asterisks represent mean± SEM. *p<0,05; ***p<0,001.

**Table 2.** Gene ontology analysis was performed for PANTHER pathways using the Gene Ontology Consortium database (binomial statistical test with Bonferroni correction for multiple testing: p ≤ 0.05).

| GO category | Fold enrichment | p value |
| --- | --- | --- |
| Synaptic vesicle (GO:0008021) | 2.58 | 3.78E-06 |
| Synapse (GO:0045202)® | 1.82 | 3.65E-14 |
| Postsynapse (GO:0098794) | 1.69 | 3.62E-04 |
| Glutamatergic synapse (GO:0098978) | 1.82 | 7.06E-03 |

### 3.3 ATXN3 modulates SRSF7 degradation by the proteasome

Interestingly, among the splicing factors with differential ubiquitylation was the Arginine/Serine-Rich splicing factor SRSF7 (also known as 9G8), known to be involved in the regulation of alternative splicing of the neurodegeneration-related gene MAPT [46], encoding the Tau protein. As shown in Figure 3a, we detected decreased levels of polyubiquitylated SRSF7 presenting Ub-K48 linkages in cells lacking ATXN3, a type of Ub chain that usually signals proteins for degradation through the UPP.

**Figure 3.**
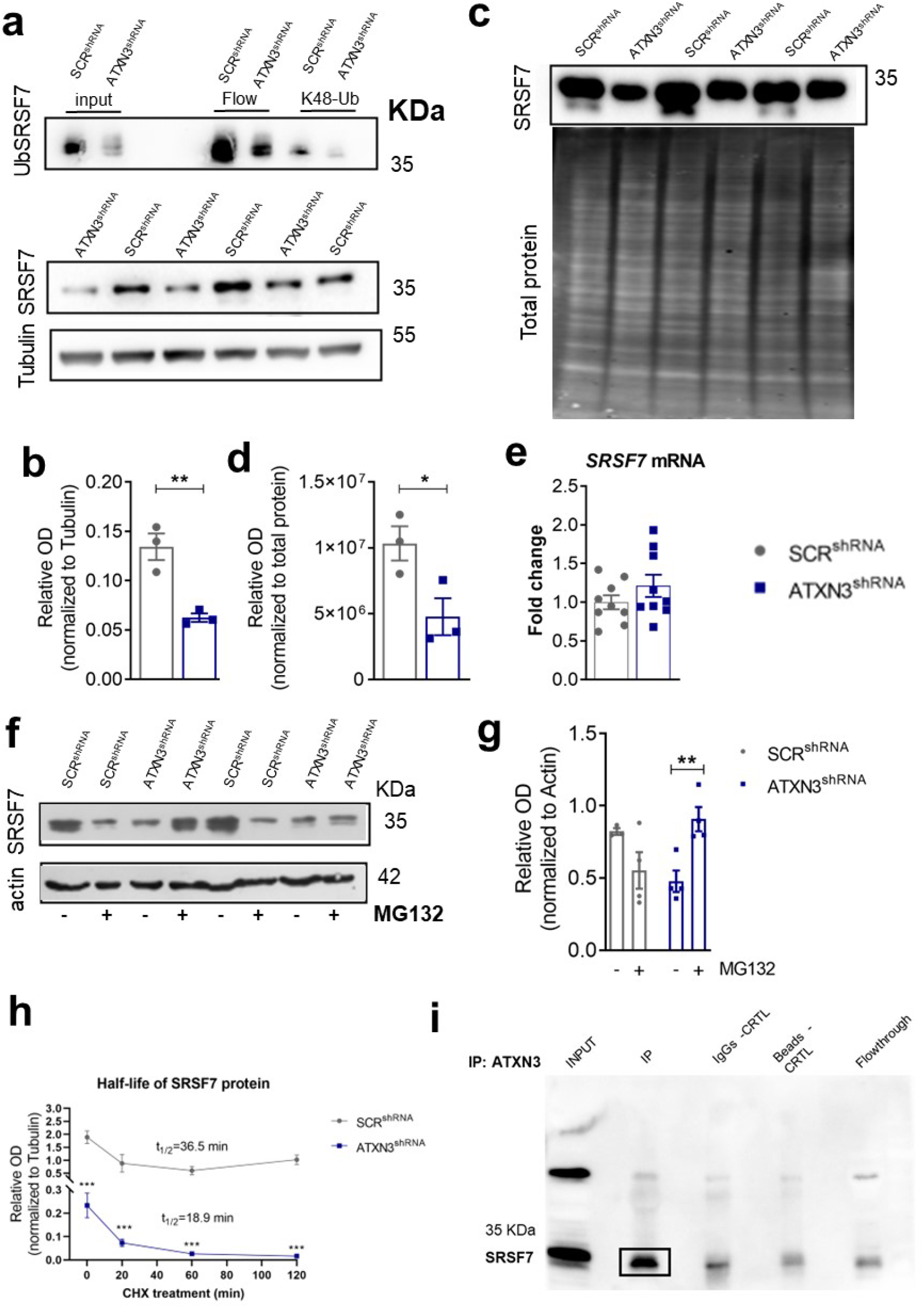
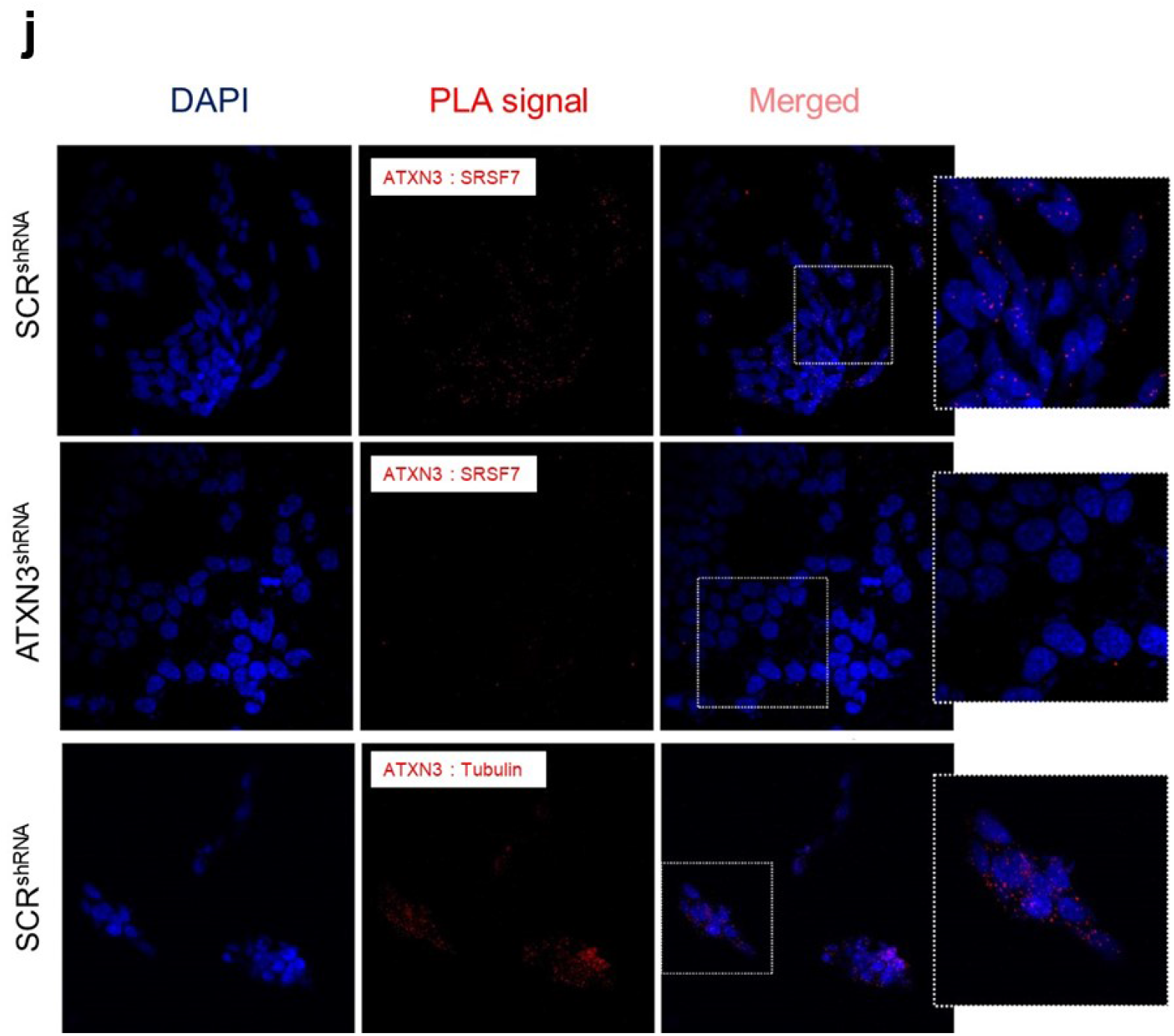
SRSF7 protein levels in ATXN3^shRNA^ and SCR^shRNA^ control cells. **(a-b)** Levels of both total and polyubiquitylated of SRSF7 in ATXN3^shRNA^ cells and the SCR^shRNA^ controls, as assessed by Western-blot analysis after capture of polyubiquitylated proteins with K48-linked polyubiquitin chains. Relative band density for total SRSF7 levels was analyzed and normalized for tubulin. **(c-d)** Nuclear SRSF7 protein levels are decreased in ATXN3^shRNA^ cells. Relative band density for SRSF7 was analyzed and normalized for total protein levels (Azured staining). **(e)** No significant differences were observed in the mRNA levels of SRSF7 between the ATXN3^shRNA^ cells and the SCR^shRNA^ controls, **(f-g)** Protein levels of SRSF7 in ATXN3^shRNA^ cells and SCR^shRNA^ controls upon proteasome inhibition (MG132 treatment). Relative band density for total SRSF7 levels was analyzed and normalized for actin. **(h)** Relative amounts of SRSF7 in ATXN3^shRNA^ cells and SCR^shRNA^ cells at various cycloheximide treatment times. Relative band density for total SRSF7 levels was analyzed and normalized for tubulin. **(i)** In vitro and **(j)** in situ interaction of Human ATXN3 with SRSF7. Scale bar corresponds to 40 μm. n≥3 independent biological replicates in all experiments. Student’s t-test or ANOVA were used for comparisons. Asterisks represent mean±SEM. *p<0.05; **p<0.01, ***p<0.01.

Consistent with the hypothesis that the differential ubiquitylation of SRSF7 can have an impact on its degradation, the steady state levels of total (Figure 3a-b) and nuclear (Figure 3c-d) SRSF7 were decreased in ATXN3^shRNA^ cells as compared with the SCR^shRNA^ control cells. This suggests that SRSF7 is indeed being more degraded in the absence of ATXN3, confirming that ATXN3 affects the expression of SRSF7 at the protein and not at the mRNA level, since both the steady state levels and stability of SRSF7 mRNA were unchanged in the absence of this protein (Figure 3e, Figure S1b, respectively).

Based on the fact that SRSF7 was captured using Agarose-TUBEs that have higher affinity for polyubiquitylated than for monoubiquitylated proteins [45], and considering that ATXN3 is a DUB, we asked if ATXN3 could modify ubiquitylation and regulate degradation of SRSF7 through the UPP. Supporting this possibility, the levels of SRSF7 were significantly increased in ATXN3^shRNA^ cells upon proteasome inhibition with MG132 (Figure 3f-g). Protein stability analysis upon inhibition of protein synthesis by cicloheximide treatment showed a decrease in the stability of SRSF7 in ATXN3^shRNA^ cells (Figure 3h), suggesting that ATXN3 normally acts to inhibit SRSF7 degradation in neurons. In order for ATXN3 to modify the ubiquitylation and stability of SRSF7, the two proteins would need to interact. Co-immunoprecipitation using protein extracts from differentiated neuron-like SH-SY5Y cells confirmed that an interaction between ATXN3 and SRSF7 occurs in these cells (Figure 3i). This interaction was further confirmed in live cells by the Protein Ligation Assay (Figure 3j, Figure S1c). Globally, our data are compatible with the hypothesis of SRSF7 being a substrate of the DUB activity of ATXN3 in neurons.

### 3.4 Tau splicing is deregulated in the absence of ATXN3

Given that one of the splicing factors showing differential ubiquitylation was SRSF7 which modulates tau mRNA splicing [47]–namely the inclusion or not of exon 10 in the mature mRNA (forming the 4R variant of this protein), we designed a very specific qRT-PCR protocol (Figure S3) to analyze mRNA levels of Tau isoforms in ATXN3^shRNA^ cells, in order to assess the functional impact of the reduction of this splicing factor in Tau splicing. We found a significant decrease in the transcript levels of the 4R-Tau isoform (Figure 4a), but no alteration of the 3R-Tau isoform in ATXN3^shRNA^ cells as compared with SCR^shRNA^ control cells (Figure 4b), leading to an altered 4R/3R-Tau ratio (Figure 4c) and decreased total tau levels, both in ATXN3^shRNA^ cells (Figure 4d) as well as in cultured primary cerebellar neurons in which ATXN3 expression was partially silenced by shRNA (Figure 4e-h, Figure S1d). Given that SRSF7, in addition to its activity promoting MAPT exon 10 inclusion also increases the stability of the 4R-encoding mRNA [48], it was interesting to see that absence of ATXN3 led to a decrease in the stability of 4R-Tau mRNA, seen upon treatment with actinomycin to block translation (Figure 4i).

**Figure 4.**
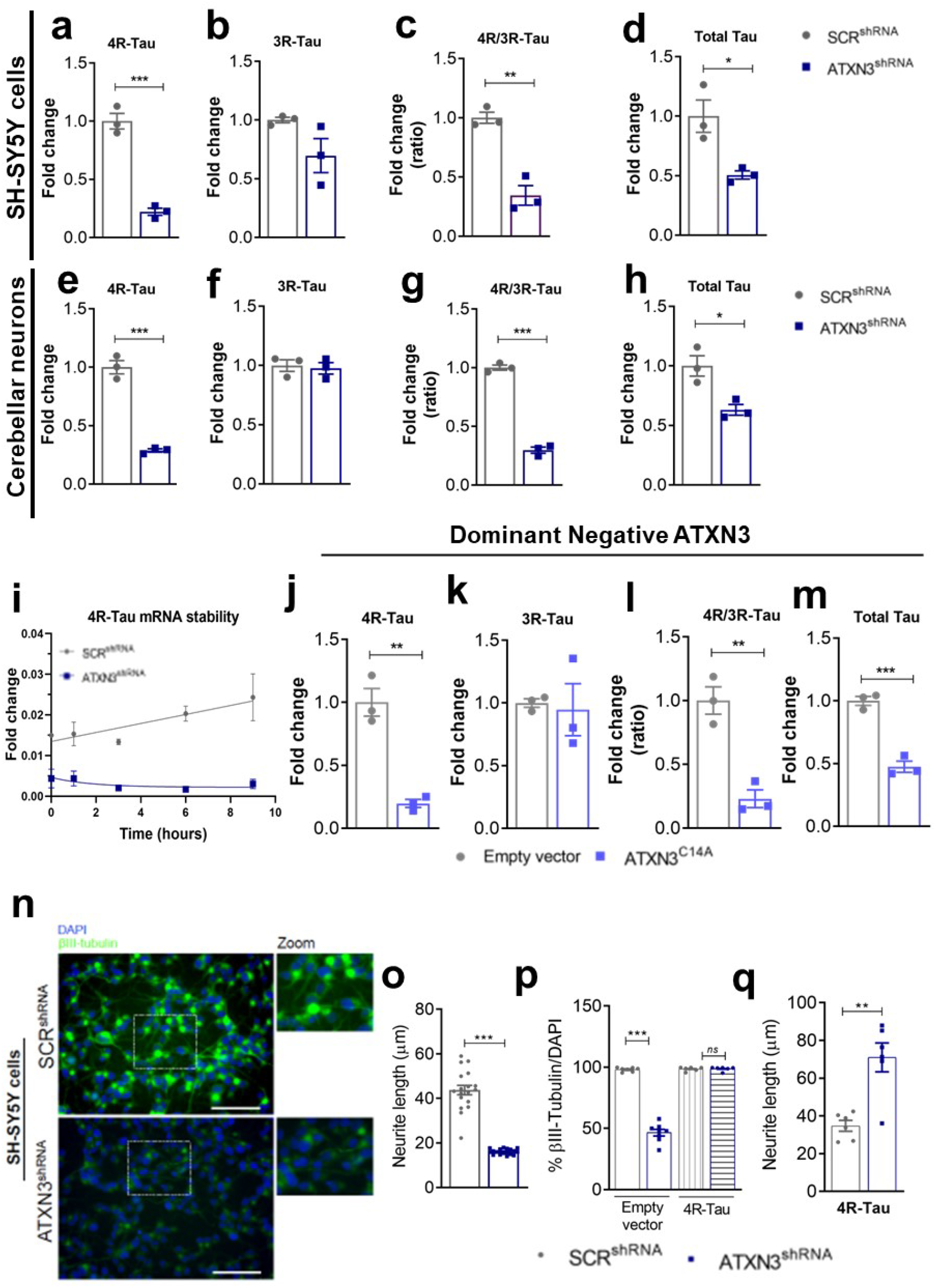
Disruption of 4R/3R-Tau ratio in ATXN3^shRNA^ cells and SCR^shRNA^ controls. **(a)** qRT-PCR analysis of transcription levels of 4R-Tau, **(b)** 3R-Tau isoform, **(c)** 4R/3R-Tau ratio and **(d)** total Tau, in ATXN3^shRNA^ cells and SCR^shRNA^ controls, and in **(e-h)** cerebellar primary neurons. mRNA levels were normalized to the *HMBS* (a-d) or *B2M* (e-h) genes. **(i)** mRNA stability of the 4R tau isoform. mRNA levels were normalized to the *B2M gene.* **(j)** mRNA levels of 4R-Tau isoform, **(k)** 3R-Tau isoform, **(l)** 4R/3R-Tau ratio and **(m)** total Tau in ATXN3_C14A and control cells. mRNA levels were normalized to the *HMBS gene.* The 2-ΔΔCT method was used to calculate the relative fold gene expression. 4R/3R-Tau ratio was obtained by dividing 4R and 3R-Tau mRNA levels. **(n)** Expression of βIII-tubulin in ATXN3^shRNA^ cells and SCR^shRNA^ controls as assessed by immunostaining. **(o)** Average length of the neuritis in ATXN3^shRNA^ cells and SCR^shRNA^ controls. **(p)** Overexpression of 4R-Tau rescued the expression of βIΠ-tubulin and **(q)** the average length of neurites in ATXN3^shRNA^ cells. n≤3 independent biological replicates in all experiments. Student’s t-test was used for comparisons. Asterisks represent mean±SEM. *p<0.05; **p<0.01, ***p<0.01.

Consistently, catalytically inactive (dominant negative) ATXN3 expression also led to alterations in tau expression and in the isoform ratio, similar to those seen in cells with silenced ATXN3: decreased expression of 4R-Tau (Figure 4j), no alterations on the 3R-Tau isoform (Figure 4k), and consequentially a decreased 4R/3R-Tau ratio (Figure 4l) together with decreased total Tau mRNA levels (Figure 4m). In order to test if the reduction of the 4R tau isoform and consequent deregulation of the 4R/3R tau ratio was contributing to the neuronal phenotype observed in ATXN3^shRNA cells^, we overexpressed the 4R-Tau isoform in these cells and evaluated their ability to differentiate into neuronal cells upon retinoic acid treatment. Indeed, we observed a rescue of the expression levels of several neuronal differentiation markers–MAP2, βIII-tubulin and Neurogenin–and a tendency towards a decrease in the expression of Nestin, a marker for proliferative undifferentiated cells (Figure S1e). To explore the functional consequences of this reduction in 4R-Tau levels, we performed a morphometric analysis of these cells; ATXN3^shRNA^ cultures presented a decreased expression of βIII-tubulin and a decreased average length of the neurite as compared with the SCR^shRNA^ controls, which correlated with a more immature stage of neuronal differentiation (Figure 4n-o). Of note, overexpression of 4R-Tau in ATXN3^shRNA^ improved both of these parameters (Figure 4p-q).

### 3.5 Decreased SRSF7 expression and Tau splicing perturbation in a SCA3 cell model and in patient brains

To determine if the presence of an expanded polyglutamine (polyQ) tract within ATXN3 would lead to an alteration in this newly identified physiological function, we analyzed a SH-SY5Y cell line expressing a version of ATXN3 bearing 83 glutamines (ATXN3_83Q). We asked whether the presence of the polyQ tract could also impact MAPT mRNA splicing regulation, altering the 4R/3R-Tau ratio. Interestingly, expression of ATXN3_83Q led to similar changes in the expression of tau isoforms as observed in cells lacking ATXN3 or expressing a catalytic mutant version of this protein: i) decreased expression of 4R-Tau isoform (Figure 5a), ii) no alterations on the 3R-Tau isoform (Figure 5b), iii) a decrease of the 4R/3R-Tau ratio (Figure 5c), and (iv) a decrease in the total levels of Tau mRNA (Figure 5d). Of notice, the overexpression of the wild type ATXN3 (ATXN3_28Q) also caused some degree of perturbation (Figure 5a-d), pointing to the importance of a tight regulation of ATXN3 dosage in neurons. Most interestingly, mildly decreased levels of SRSF7 protein were also present in the brains of SCA3 patients as compared with healthy controls (Figure 5e).

**Figure 5.**
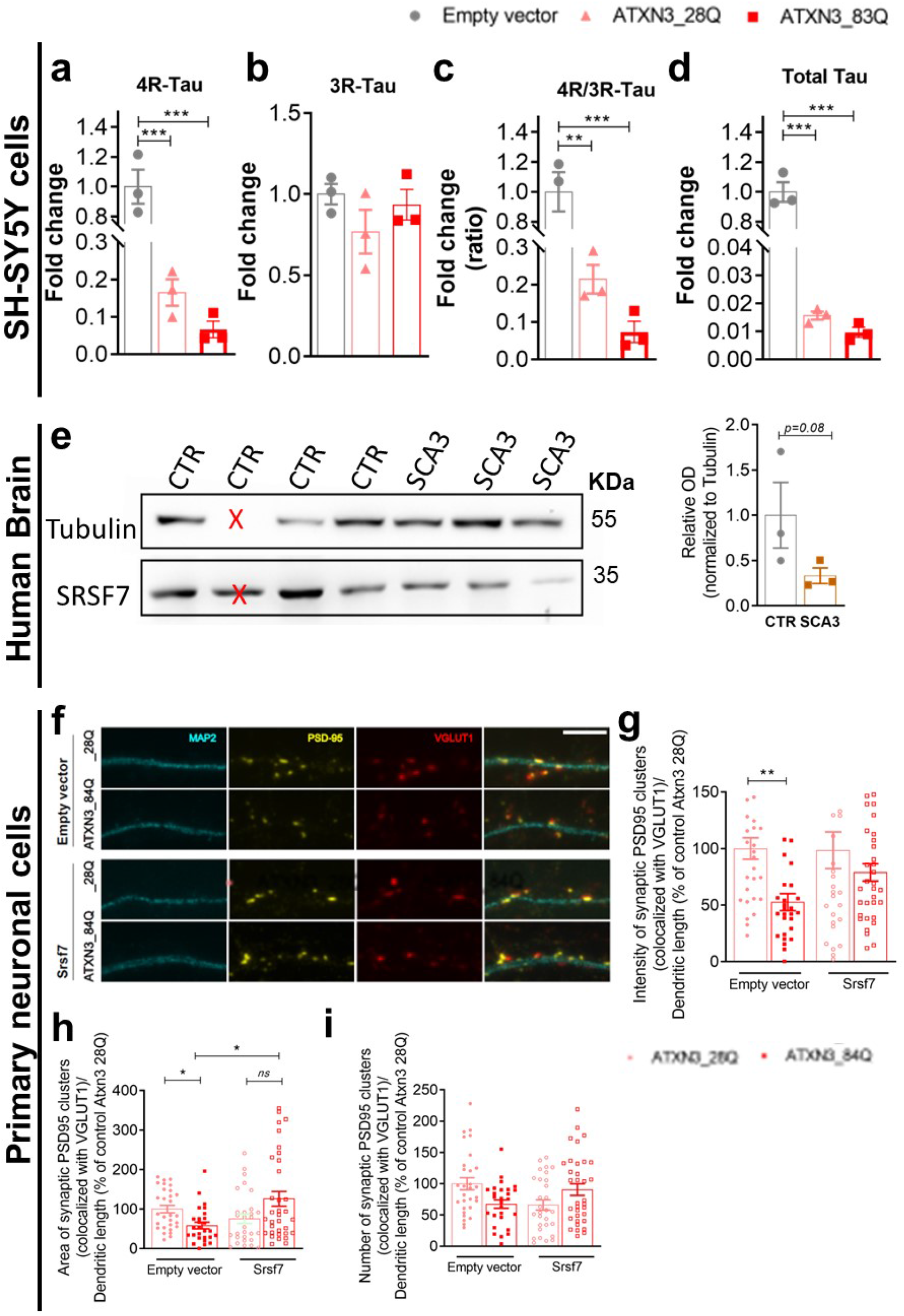
PolyQ expansion in ATXN3 affects Tau splicing. **(a)** mRNA levels of 4R-Tau isoform, **(b)** 3R-Tau isoform, **(c)** 4R/3R-Tau ratio, and **(d)** total Tau in ATXN3_83Q and 28Q-expressing cells. mRNA levels were normalized to the *HMBS gene*. **(e)** SRSF7 protein levels in SCA3 patient’s brains. Relative band density for total SRSF7 levels was analyzed and normalized for Tubulin. Tubulin was not detected in one control sample (labelled with a red cross) and therefore not quantified and included in the final data analysis. **(f)** Representative images of primary cortical neurons at 16 DIV expressing GFP-Ataxin-3 (with 28Q or 84Q) and coexpressing Srsf7 when indicated. Neurons were stained for PSD-95, VGluT1, eGFP and MAP2. Scale bar: 5μm. **(g)** Intensity, **(h)** area and **(i)** number of PSD-95 clusters (co-localizing with VGluT1 clusters) per dendritic length (n=27-34 neurons per experimental condition) and 3 independent experiments. Student’s t-test, ANOVA or Kruskal-Wallis tests were applied when appropriated. *Asterisks represent mean±SEM. *p<0.05; **p<0.01; ***p<0.001.

Knowing that Tau plays a role in synapse formation and maintenance [49], and since synapse related genes were significantly enriched among the differentially spliced genes in ATXN3 silenced cells, we tested whether supplementation of SRSF7 rescued the previously described synaptotoxic effects of expanded ATXN3 expression in primary cortical neurons [50]. For this, we quantified excitatory synaptic clusters by labelling the postsynaptic scaffold protein PSD-95 and the pre-synaptic vesicle protein VGlut1. We saw a reduction in intensity, area, and no differences in number of PSD-95 clusters that co-localize with VGlut1 corresponding to potentially functionally synapses, in cortical neurons expressing mutant ATXN3, a difference that was mitigated by overexpression of SRSF7 (Figure 5f-i), suggesting that reduced levels of this splicing factor are relevant for the negative impact of expanded ATXN3 on excitatory synapses.

Altogether, these results suggest that the functional interaction of ATXN3 with SRSF7 and its impact on molecular mechanisms regulating tau splicing in neurons is of functional relevance in these cells and may be relevant in the pathogenesis of SCA3.

## 4. Discussion

Ubiquitylation is a post-translational modification that controls several aspects of neuronal function by regulating protein abundance. Disruption of this signaling pathway has been described in neurodegenerative diseases. Indeed, since many of these diseases exhibit ubiquitylated protein aggregates, loss of ubiquitin homeostasis may be an important contributor for disease. Considering the importance of DUB enzymes in maintaining the ubiquitylation balance, we focused on the characterization of the ubiquitome of neuronal cells lacking ATXN3 [48]. Using TUBEs, that enable the pulldown of polyubiquitylated proteins without further genetic manipulation or inhibition of the proteasome [48] in combination with LC-MS/MS, we were able to consistently identify 615 proteins per condition, a yield comparable to those described in other studies [51, 52]. Among the proteins identified, approximately one third presented altered levels of polyubiquitylation in ATXN3^shRNA^ cells. Curiously, the majority of these proteins presented decreased levels in the polyubiquitin-enriched fraction signal, suggesting that normally ATXN3 might be preventing their degradation, for instance by editing the substrate’s ubiquitin chains, as we observed before for ITGA5 [24, 25]. Therefore, when ATXN3 is silenced, the polyubiquitin signal is not removed, which may result in an increased degradation of the targeted proteins. The fact that a significant proportion of the proteins with altered polyubiquitylation levels in ATXN3^shRNA^ cells were splicing factors and proteins involved in RNA processing, is suggestive of ATXN3 playing a role in the pre-mRNA splicing process in neuronal cells. This process is particularly important in generating diversity and specificity in the central nervous system, which requires a wide protein repertoire to generate its highly specialized and adaptable neuronal circuits [53]. Given the importance of RNA processing for neuronal function, it comes as no surprise that altered RNA processing is a contributing factor to the pathogenesis of multiple neurodegenerative diseases [54–58]. Disruption of alternative splicing (AS) regulation–resulting from mutations or abnormal expression of RNA-binding proteins–can lead to an imbalance or an inappropriate expression pattern of key protein isoforms, with profound consequences on cellular and organismal physiology. Indeed, the relative concentration of splicing factors and hnRNPs, which may be regulated by the Ub pathway, has been shown to regulate AS [59]. For example, it was shown that the abundance of several splicing factors is differentially affected by proteasome inhibition, and abnormal levels of these regulatory proteins were associated with a different AS pattern [60]. Another study showed that the splicing factor SRSF5 is targeted to proteasomal degradation as the cells undergo terminal differentiation [61]. Consistently, general splicing efficacy was decreased in cells depleted of ATXN3, as shown using artificial reporter minigenes, further supporting the involvement of ATXN3 in the regulation of the splicing process. Additionally, genome wide microarray analysis of endogenous splicing events confirmed that absence of ATXN3 leads to a dramatic alteration in the splicing pattern of a large number of genes in neuronal cells, including genes related to protein degradation, adhesion, axon guidance, synapses and signaling pathways. This is consistent with our previous findings showing that absence of ATXN3 leads to an impairment in neuronal differentiation and adhesion [25], and affects the degradation of several target proteins [23–25]. Significantly, a portion of the predicted splicing regulators of these genes with altered splicing in cells lacking ATXN3 were found to have altered levels of polyubiquitylation in our proteomic analysis, the majority of them being Serine/Arginine (SR)–rich phosphoproteins. Proteins of the SR family are key players in the control of AS, regulating the selection of alternative sites [62, 63]. The fact that SRSF7 (9G8), one of those proteins and a key regulator of MAPT (tau) exon 10 splicing [47, 64], was captured using the Agarose-TUBEs, which have high affinity for polyubiquitin chains [45], suggested that this splicing factor is polyubiquitylated, and thus might be degraded through the UPP. Indeed, while it has been described that the mature human neurons express approximately equal levels of 4R and 3R-Tau isoforms [65, 66], whose expression is developmentally controlled and finely-tuned, we found that loss of function of ATXN3 disrupts this balance in neuronal cells leading to a decreased 4R/3R-Tau ratio, that coincides with (and may be caused by) decreased levels of SRSF7. Recent works have shown imbalanced expression of 3R and 4R-Tau, namely a decreased expression of the 4R isoform as here observed, in the brain of a mouse model of Down syndrome, which shows age-related neurodegeneration, but also in human Pick’s disease, AD and PD patients [67–69], strengthening the idea that a faulty tau exon 10 splicing might be contributing for the pathogenesis of several neurodegenerative diseases. In addition, the decreased levels of the 4R-Tau isoform presented by ATXN3^shRNA^ cells, also correlate with the immature phenotype of these cells, that we previously described [25]. Indeed, Conejero-Godberg and colleagues also assessed the transcriptional profiles related to 3R and 4R-Tau splice variants, revealing that a cluster of genes related to neuronal cell morphology and outgrowth of neurites is upregulated in the presence of 4R isoforms, while genes related to cellular growth and proliferation were downregulated [70]. This is in agreement with previous work from Sennvik’s team establishing a role for 4R-Tau isoforms in promoting neuronal differentiation [71].

Taking into account that loss of function of ATXN3 leads to a deregulation in the amount of SRSF7, at the protein level but not RNA level, we hypothesized that this splicing factor could be a substrate of the DUB activity of ATXN3, which could be modulating its degradation. In line with that, we confirmed that ATXN3 interacts closely with SRSF7 in neuronal cells, suggesting that these proteins are molecular partners in such cells, and establishing a functional link between two key proteins involved in different neurodegenerative diseases. Trying to explore if this interaction could be indicative that ATXN3 is modulating the degradation of SRSF7 through the proteasome, we measured the levels of this protein upon proteasome inhibition; as expected, we saw an accumulation of SRSF7 when the proteasome was inhibited.

Interestingly, we also found a decreased 4R/3R-Tau ratio, as well as decreased protein levels of SRSF7, in cells expressing the mutant ATXN3 protein. This suggests that the mis-splicing of tau may be contributing to SCA3 pathogenesis, as it does to other neurodegenerative diseases. Confirming a relevant role for SRSF7 reduction in the neurotoxic properties of expanded ATXN3, the synaptic loss phenotype of primary neurons overexpressing ATXN3 with 84 glutamines was mitigated by SRSF7 overexpression. Considering that we observed similar phenotype and similar molecular changes using cells lacking ATXN3 and cells expressing either a catalytic inactive mutant form of the protein or the expanded disease-associated form, it is reasonable to think that the DUB activity of ATXN3 is important in this cellular process and that the polyglutamine expansion might cause a partial loss of this normal function of the protein (even in presence of the wild type protein), perhaps through a dominant negative mechanism, thus contributing to SCA3 pathogenesis. Indeed, Winborn and co-workers demonstrated that while overexpression of ATXN3 with a non-pathogenic polyQ tract lead to a reduction in total ubiquitylated protein levels in cells, this reduction was not observed upon overexpression of expanded ATXN3, which may reflect an inefficient deubiquitylation of cellular proteins, pointing to an overall loss of function [72]. Overall, our data suggest that ATXN3 is involved in RNA metabolism and particularly in splicing regulation through the modulation of the ubiquitylation of splicing factors. This regulatory role may be mediated through the DUB activity of ATXN3 by modulating activation, degradation and/or subcellular localization of the splicing factors, or may be an indirect result of the modulation of E3 ligases specific for these targets.

Recently, several pieces of evidence coming from the use of technologies such as exon arrays and RNA-Seq suggested an association between perturbation of AS and several neurodegenerative diseases including Alzheimer’s disease, Huntington’s disease, Pick’s disease, in addition to the more directly splicing-related Spinal Muscular Atrophy and Amyotrophic Lateral Sclerosis [73–75]. Importantly, several ataxia-causing proteins have been described to interact with splicing factors and other proteins related to mRNA metabolism (reviewed in [76, 77]). Also, expression of mutant ATXN3 bearing an expanded polyglutamine tract was previously shown to alter the ability of the subnuclear domains known as Cajal bodies to efficiently participate in small nuclear ribonucleoprotein (snRNP) biogenesis pathway, and to reduce the efficacy of splicing of reporter genes [78, 79]. These findings lead us to propose that ATXN3 plays a role in splicing regulation in neurons, a novel function for this protein, and that a perturbation of this physiological function might contribute for SCA3 pathogenesis.

## Supplementary figure and table captions

**Figure S1. No changes in SRSF7 mRNA stability in ATXN3^shRNA^ cells. (a)** ATXN3 silencing using different shRNA clones as compared to the scrambled sequence and empty vector expression. Tubulin is used as reference. **(b)** No differences were observed in the SRSF7 mRNA stability after cycloheximide (CHX) treatment at different timepoints. mRNA levels were normalized to the *HMBS* gene. **(c)** PLA negative control (no antibodies) and single antibody staining in control cells. **(d)** Knockdown of Ataxin-3 in rat cerebellar primary neurons. Relative band density for Ataxin-3 levels was analyzed and normalized for tubulin protein levels. **(e)** Expression of differentiation markers assessed by qRT-PCR in ATXN3^shRNA^ cells overexpressing 4R tau isoform. mRNA levels were normalized to the *HMBS* gene. The 2-ΔΔCT method was used to calculate the relative fold gene expression. nɤ3 independent biological replicates in all experiments. Student’s t-test was used for comparisons. Asterisks represent mean±SEM. *p<0.05, **p<0.01.

**Figure S2. qRT-PCR validation of the expression microarray data. (a)** Schematic representation of the strategy used to design a set of primers were design to evaluate a set of different splicing events. **(b)** qRT-PCR validation of the microarrays data. mRNA levels were normalized to the *B2M* gene. The 2-ΔΔCT method was used to calculate the relative fold gene expression.

**Figure S3. Experimental design to analyze the expression of Tau isoforms by qRT-PCR. (a)** MAPT and the six isoforms found in adult human brain. Exons 1, 4, 5, 7, 9 and 11-13 are constitutively spliced (blue) and exon 0, which is part of the promoter, and 14 are non-coding (white). Exon 6 and 8 are not transcribed in human brain (orange) and the unusually long exon 4a (grey) is expressed only in the peripheral nervous system. Alternative splicing of exon 2 (green), 3 (purple) and 10 (red) gives rise to the six isoforms. The repeats of tau (R1-R4) are indicated and 4R and 3R tau forward (FW) and reverse (RV) primer locations are shown. **(b)** Screenshot of a qPCR experiment demonstrating the specificity of primers for 4R and 3R tau. cDNA of 4R0N and 3R0N tau constructs were used as template. Primer pairs specific for 4R tau amplify 4R0N cDNA constructs and not 3R0N constructs. Primer pairs specific for 3R tau, using locked nucleic acids, amplify 3R0N cDNA constructs and not 4R0N cDNA constructs.

**Table S1. Differences in polyubiquitylated proteins identified by TUBEs-MS in SH-SY5Y cells lacking ATXN3.** List of identified proteins with altered levels of ubiquitylation in RA-treated ATXN3^shRNA^ cells as compared with the SCR^shRNA^ controls (p<0.05). These proteins were present in at least 3 independent experiments. In red and green are proteins with increased and decreased polyubiquitylated levels, respectively. $ indicates protein splicing factors.

**Table S2. Differentially regulated alternative splicing events in ATXN3^shRNA^ cells.** List of genes with altered splicing in RA-treated ATXN3^shRNA^ cells as compared with the SCR^shRNA^ controls.

**Table S3. KEGG pathway analysis of genes with altered splicing in ATXN3^shRNA^ cells.** The genes identified on the KEGG pathway analysis presented at least one differentially regulated exon/splicing pattern in ATXN3^shRNA^ cells.

**Table S4. Changes in alternative types events in ATXN3^shRNA^ cells.**

## Author Contributions

Conceptualization, Andreia Neves-Carvalho, Sara Duarte-Silva, Andreia Teixeira-Castro, Ioannis Sotiropoulos, Peter Heutink, Ka Wan Li and Patrícia Maciel; Data curation, Andreia Neves-Carvalho, Sara Duarte-Silva, Joana Silva and Patrícia Maciel; Formal analysis, Sara Duarte-Silva and Joana Silva; Funding acquisition, Patrícia Maciel; Investigation, Andreia Neves-Carvalho, Sara Duarte-Silva, Joana Silva, Liliana Meireles-Costa, Daniela Monteiro-Fernandes, Joana Correia, Beatriz Rodrigues, Sasja Heetveld, Bruno Almeida, Natalia Savytska, Jorge Da Silva, Ana Luisa Carvalho, Ka Wan Li and Patrícia Maciel; Methodology, Andreia Neves-Carvalho, Sara Duarte-Silva, Joana Silva and Patrícia Maciel; Project administration, Patrícia Maciel; Resources, Andreia Teixeira-Castro, Ioannis Sotiropoulos, Ka Wan Li and Patrícia Maciel; Software, Patrícia Maciel; Supervision, Peter Heutink and Patrícia Maciel; Validation, Liliana Meireles-Costa and Bruno Almeida; Visualization, Patrícia Maciel; Writing–original draft, Andreia Neves-Carvalho and Sara Duarte-Silva; Writing–review & editing, Patricia Maciel.

## Institutional Review Board Statement

Human post-mortem brain samples were obtained from the Michigan Brain Bank (University of Michigan, Ann Arbor, MI).

## Informed Consent Statement

Informed consent was obtained from all subjects involved in the study.

## Data Availability Statement

Not applicable.

## Acknowledgments

The authors want to thank João Relvas, PhD, for the help with the revisions of the paper. A special thanks to the Translational Neurogenetics team (Maciel lab) for all the fruitful discussions, particularly to Dr. Marta Costa for the kind help with qRT-PCRs data analysis.

## References

1. Hegde, A.N. and S.C. Upadhya, Role of ubiquitin-proteasome-mediated proteolysis in nervous system disease. Biochim Biophys Acta, 2011. 1809(2): p. 128–40.

2. Yi, J.J. and M.D. Ehlers, Emerging roles for ubiquitin and protein degradation in neuronal function. Pharmacol Rev, 2007. 59(1): p. 14–39.

3. Baptista, M.S., C.B. Duarte, and P. Maciel, Role of the ubiquitin-proteasome system in nervous system function and disease: using C. elegans as a dissecting tool. Cell Mol Life Sci, 2012. 69(16): p. 2691–715.

4. Hershko, A. and A. Ciechanover, The ubiquitin system. Annu Rev Biochem, 1998. 67: p. 425–79.

5. Komander, D., The emerging complexity of protein ubiquitination. Biochem Soc Trans, 2009. 37(Pt 5): p. 937–53.

6. Tai, H.C. and E.M. Schuman, Ubiquitin, the proteasome and protein degradation in neuronal function and dysfunction. Nat Rev Neurosci, 2008. 9(11): p. 826–38.

7. de Vrij, F.M., et al., Protein quality control in Alzheimer’s disease by the ubiquitin proteasome system. Prog Neurobiol, 2004. 74(5): p. 249–70.

8. Upadhya, S.C. and A.N. Hegde, Ubiquitin-proteasome pathway components as therapeutic targets for CNS maladies. Curr Pharm Des, 2005. 11(29): p. 3807–28.

9. Rubinsztein, D.C., The roles of intracellular protein-degradation pathways in neurodegeneration. Nature, 2006. 443(7113): p. 780–6.

10. Nijman, S.M., et al., A genomic and functional inventory of deubiquitinating enzymes. Cell, 2005. 123(5): p. 773–86.

11. Komander, D., M.J. Clague, and S. Urbe, Breaking the chains: structure and function of the deubiquitinases. Nat Rev Mol Cell Biol, 2009. 10(8): p. 550–63.

12. Clague, M.J., J.M. Coulson, and S. Urbe, Cellular functions of the DUBs. J Cell Sci, 2012. **125**(Pt 2): p. 277–86.

13. Rappsilber, J., et al., Large-scale proteomic analysis of the human spliceosome. Genome Res, 2002. 12(8): p. 1231–45.

14. Makarov, E.M., et al., Small nuclear ribonucleoprotein remodeling during catalytic activation of the spliceosome. Science, 2002. 298(5601): p. 2205–8.

15. Wagner, S.A., et al., A proteome-wide, quantitative survey of in vivo ubiquitylation sites reveals widespread regulatory roles. Mol Cell Proteomics, 2011. 10(10): p. M111 013284.

16. Peng, J., et al., A proteomics approach to understanding protein ubiquitination. Nat Biotechnol, 2003. 21(8): p. 921–6.

17. Bellare, P., et al., Ubiquitin binding by a variant Jab1/MPN domain in the essential pre-mRNA splicing factor Prp8p. RNA, 2006. 12(2): p. 292–302.

18. Kramer, A., et al., Mammalian splicing factor SF3a120 represents a new member of the SURP family of proteins and is homologous to the essential splicing factor PRP21p of Saccharomyces cerevisiae. RNA, 1995. 1(3): p. 260–72.

19. Lygerou, Z., G. Christophides, and B. Seraphin, A novel genetic screen for snRNP assembly factors in yeast identifies a conserved protein, Sad1p, also required for pre-mRNA splicing. Mol Cell Biol, 1999. 19(3): p. 2008–20.

20. Makarova, O.V., E.M. Makarov, and R. Luhrmann, The 65 and 110 kDa SR-related proteins of the U4/U6.U5 tri-snRNP are essential for the assembly of mature spliceosomes. EMBO J, 2001. 20(10): p. 2553–63.

21. Burnett, B., F. Li, and R.N. Pittman, The polyglutamine neurodegenerative protein ataxin-3 binds polyubiquitylated proteins and has ubiquitin protease activity. Hum Mol Genet, 2003. 12(23): p. 3195–205.

22. Doss-Pepe, E.W., et al., Ataxin-3 interactions with rad23 and valosin-containing protein and its associations with ubiquitin chains and the proteasome are consistent with a role in ubiquitin-mediated proteolysis. Mol Cell Biol, 2003. 23(18): p. 6469–83.

23. Rodrigues, A.J., et al., Absence of ataxin-3 leads to cytoskeletal disorganization and increased cell death. Biochim Biophys Acta, 2010. 1803(10): p. 1154–63.

24. do Carmo Costa, M., et al., Ataxin-3 plays a role in mouse myogenic differentiation through regulation of integrin subunit levels. PLoS One, 2010. 5(7): p. e11728.

25. Neves-Carvalho, A., et al., Dominant negative effect of polyglutamine expansion perturbs normal function of ataxin-3 in neuronal cells. Hum Mol Genet, 2015. 24(1): p. 100–17.

26. Zhong, X. and R.N. Pittman, Ataxin-3 binds VCP/p97 and regulates retrotranslocation of ERAD substrates. Hum Mol Genet, 2006. 15(16): p. 2409–20.

27. Wang, G., et al., Ataxin-3, the MJD1 gene product, interacts with the two human homologs of yeast DNA repair protein RAD23, HHR23A and HHR23B. Hum Mol Genet, 2000. 9(12): p. 1795–803.

28. Chatterjee, A., et al., The role of the mammalian DNA end-processing enzyme polynucleotide kinase 3’-phosphatase in spinocerebellar ataxia type 3 pathogenesis. PLoS Genet, 2015. 11(1): p. e1004749.

29. Gao, R., et al., Inactivation of PNKP by mutant ATXN3 triggers apoptosis by activating the DNA damage-response pathway in SCA3. PLoS Genet, 2015. 11(1): p. e1004834.

30. Roppongi, R.T., K.P. Champagne-Jorgensen, and T.J. Siddiqui, Low-Density Primary Hippocampal Neuron Culture. J Vis Exp, 2017(122).

31. Jiang, M., L. Deng, and G. Chen, High Ca(2+)-phosphate transfection efficiency enables single neuron gene analysis. Gene Ther, 2004. 11(17): p. 1303–11.

32. Silva, M.M., et al., MicroRNA-186-5p controls GluA2 surface expression and synaptic scaling in hippocampal neurons. Proc Natl Acad Sci U S A, 2019. 116(12): p. 5727–5736.

33. Schindelin, J., et al., Fiji: an open-source platform for biological-image analysis. Nat Methods, 2012. 9(7): p. 676–82.

34. Colledge, M., et al., Targeting of PKA to glutamate receptors through a MAGUK-AKAP complex. Neuron, 2000. 27(1): p. 107–19.

35. Furney, S.J., et al., SF3B1 mutations are associated with alternative splicing in uveal melanoma. Cancer Discov, 2013. 3(10): p. 1122–1129.

36. Gandoura, S., et al., Gene-and exon-expression profiling reveals an extensive LPS-induced response in immune cells in patients with cirrhosis. J Hepatol, 2013. 58(5): p. 936–48.

37. Wang, E., et al., Global profiling of alternative splicing events and gene expression regulated by hnRNPH/F. PLoS One, 2012. 7(12): p. e51266.

38. de la Grange, P., et al., FAST DB: a website resource for the study of the expression regulation of human gene products. Nucleic Acids Res, 2005. 33(13): p. 4276–84.

39. de la Grange, P., et al., A new advance in alternative splicing databases: from catalogue to detailed analysis of regulation of expression and function of human alternative splicing variants. BMC Bioinformatics, 2007. 8: p. 180.

40. Huang da, W., B.T. Sherman, and R.A. Lempicki, Systematic and integrative analysis of large gene lists using DAVID bioinformatics resources. Nat Protoc, 2009. 4(1): p. 44–57.

41. Guth, S., et al., Evidence for substrate-specific requirement of the splicing factor U2AF(35) and for its function after polypyrimidine tract recognition by U2AF(65). Mol Cell Biol, 1999. 19(12): p. 8263–71.

42. Pacheco, T.R., et al., In vivo requirement of the small subunit of U2AF for recognition of a weak 3’ splice site. Mol Cell Biol, 2006. 26(21): p. 8183–90.

43. Fredriksson, S., et al., Protein detection using proximity-dependent DNA ligation assays. Nat Biotechnol, 2002. 20(5): p. 473–7.

44. Pfaffl, M.W., A new mathematical model for relative quantification in real-time RT-PCR. Nucleic Acids Res, 2001. 29(9): p. e45.

45. Hjerpe, R., et al., Efficient protection and isolation of ubiquitylated proteins using tandem ubiquitin-binding entities. EMBO Rep, 2009. 10(11): p. 1250–8.

46. Cavaloc, Y., et al., Characterization and cloning of the human splicing factor 9G8: a novel 35 kDa factor of the serine/arginine protein family. EMBO J, 1994. 13(11): p. 2639–49.

47. Gao, L., et al., SR protein 9G8 modulates splicing of tau exon 10 via its proximal downstream intron, a clustering region for frontotemporal dementia mutations. Mol Cell Neurosci, 2007. 34(1): p. 48–58.

48. Qian, W., et al., Splicing factor SC35 promotes tau expression through stabilization of its mRNA. FEBS Lett, 2011. 585(6): p. 875–80.

49. Pooler, A.M., W. Noble, and D.P. Hanger, A role for tau at the synapse in Alzheimer’s disease pathogenesis. Neuropharmacology, 2014. **76 Pt A**: p. 1–8.

50. Matos, C.A., et al., Ataxin-3 phosphorylation decreases neuronal defects in spinocerebellar ataxia type 3 models. J Cell Biol, 2016. 212(4): p. 465–80.

51. Lopitz-Otsoa, F., et al., Integrative analysis of the ubiquitin proteome isolated using Tandem Ubiquitin Binding Entities (TUBEs). J Proteomics, 2012. 75(10): p. 2998–3014.

52. Li, K.W., et al., Identifying true protein complex constituents in interaction proteomics: the example of the DMXL2 protein complex. Proteomics, 2012. 12(15-16): p. 2428–32.

53. Ule, J. and R.B. Darnell, RNA binding proteins and the regulation of neuronal synaptic plasticity. Curr Opin Neurobiol, 2006. 16(1): p. 102–10.

54. Anderson, P. and P. Ivanov, tRNA fragments in human health and disease. FEBS Lett, 2014. 588(23): p. 4297–304.

55. Belzil, V.V., T.F. Gendron, and L. Petrucelli, RNA-mediated toxicity in neurodegenerative disease. Mol Cell Neurosci, 2013. 56: p. 406–19.

56. Bentmann, E., C. Haass, and D. Dormann, Stress granules in neurodegeneration--lessons learnt from TAR DNA binding protein of 43 kDa and fused in sarcoma. FEBS J, 2013. 280(18): p. 4348–70.

57. Ling, S.C., M. Polymenidou, and D.W. Cleveland, Converging mechanisms in ALS and FTD: disrupted RNA and protein homeostasis. Neuron, 2013. 79(3): p. 416–38.

58. Halliday, G., et al., Mechanisms of disease in frontotemporal lobar degeneration: gain of function versus loss of function effects. Acta Neuropathol, 2012. 124(3): p. 373–82.

59. Busch, A. and K.J. Hertel, Evolution of SR protein and hnRNP splicing regulatory factors. Wiley Interdiscip Rev RNA, 2012. 3(1): p. 1–12.

60. Moulton, V.R., A.R. Gillooly, and G.C. Tsokos, Ubiquitination regulates expression of the serine/arginine-rich splicing factor 1 (SRSF1) in normal and systemic lupus erythematosus (SLE) T cells. J Biol Chem, 2014. 289(7): p. 4126–34.

61. Breig, O. and F. Baklouti, Proteasome-mediated proteolysis of SRSF5 splicing factor intriguingly co-occurs with SRSF5 mRNA upregulation during late erythroid differentiation. PLoS One, 2013. 8(3): p. e59137.

62. Graveley, B.R., Sorting out the complexity of SR protein functions. RNA, 2000. 6(9): p. 1197–211.

63. Fu, X.D., The superfamily of arginine/serine-rich splicing factors. RNA, 1995. 1(7): p. 663–80.

64. Ding, S., et al., Regulation of alternative splicing of tau exon 10 by 9G8 and Dyrk1A. Neurobiol Aging, 2012. 33(7): p. 1389–99.

65. Goedert, M., et al., Multiple isoforms of human microtubule-associated protein tau: sequences and localization in neurofibrillary tangles of Alzheimer’s disease. Neuron, 1989. 3(4): p. 519–26.

66. Goedert, M. and R. Jakes, Expression of separate isoforms of human tau protein: correlation with the tau pattern in brain and effects on tubulin polymerization. EMBO J, 1990. 9(13): p. 4225–30.

67. Yin, X., et al., Dyrk1A overexpression leads to increase of 3R-tau expression and cognitive deficits in Ts65Dn Down syndrome mice. Sci Rep, 2017. 7(1): p. 619.

68. Magdalinou, N.K., et al., Identification of candidate cerebrospinal fluid biomarkers in parkinsonism using quantitative proteomics. Parkinsonism Relat Disord, 2017. 37: p. 65–71.

69. Luk, C., et al., Development and assessment of sensitive immuno-PCR assays for the quantification of cerebrospinal fluid three-and four-repeat tau isoforms in tauopathies. J Neurochem, 2012. 123(3): p. 396–405.

70. Chen, S., et al., MAPT isoforms: differential transcriptional profiles related to 3R and 4R splice variants. J Alzheimers Dis, 2010. 22(4): p. 1313–29.

71. Sennvik, K., et al., Tau-4R suppresses proliferation and promotes neuronal differentiation in the hippocampus of tau knockin/knockout mice. FASEB J, 2007. 21(9): p. 2149–61.

72. Winborn, B.J., et al., The deubiquitinating enzyme ataxin-3, a polyglutamine disease protein, edits Lys63 linkages in mixed linkage ubiquitin chains. J Biol Chem, 2008. 283(39): p. 26436–43.

73. Singh, N.N. and R.N. Singh, Alternative splicing in spinal muscular atrophy underscores the role of an intron definition model. RNA Biol, 2011. 8(4): p. 600–6.

74. Orozco, D. and D. Edbauer, FUS-mediated alternative splicing in the nervous system: consequences for ALS and FTLD. J Mol Med (Berl), 2013. 91(12): p. 1343–54.

75. Mills, J.D. and M. Janitz, Alternative splicing of mRNA in the molecular pathology of neurodegenerative diseases. Neurobiol Aging, 2012. 33(5): p. 1012 e11–24.

76. Ranum, L.P. and T.A. Cooper, RNA-mediated neuromuscular disorders. Annu Rev Neurosci, 2006. 29: p. 259–77.

77. McLoughlin, H.S., L.R. Moore, and H.L. Paulson, Pathogenesis of SCA3 and implications for other polyglutamine diseases. Neurobiol Dis, 2020. 134: p. 104635.

78. Sun, J., et al., Differential effects of polyglutamine proteins on nuclear organization and artificial reporter splicing. J Neurosci Res, 2007. 85(11): p. 2306–17.

79. Orr, H.T., SCA1-phosphorylation, a regulator of Ataxin-1 function and pathogenesis. Prog Neurobiol, 2012. 99(3): p. 179–85.

