## Supplementary figures and images for "Regulation of neuronal mRNA splicing and Tau isoform ratio by ATXN3 through deubiquitylation of splicing factors"

### Figure S1

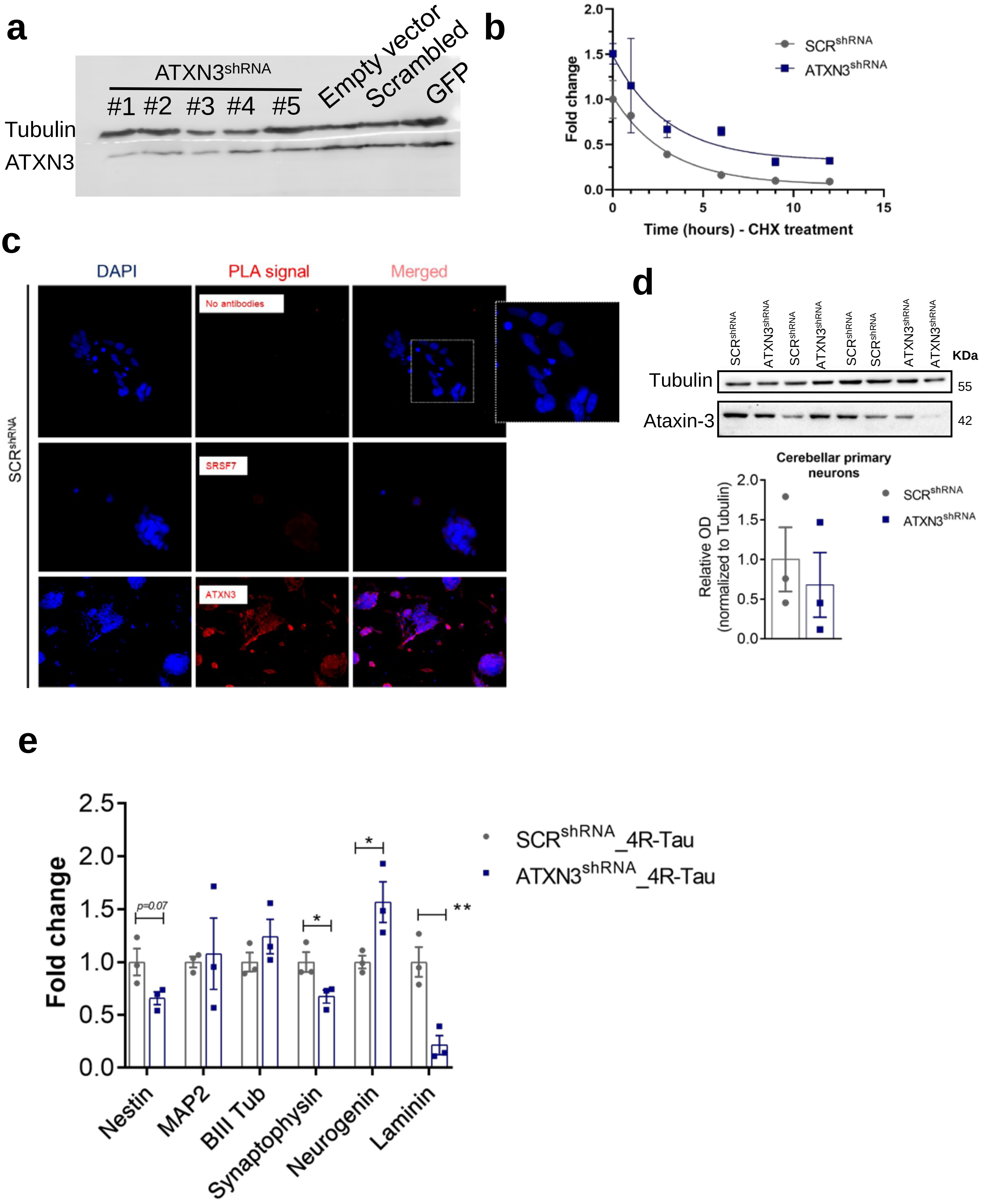

### Figure S3

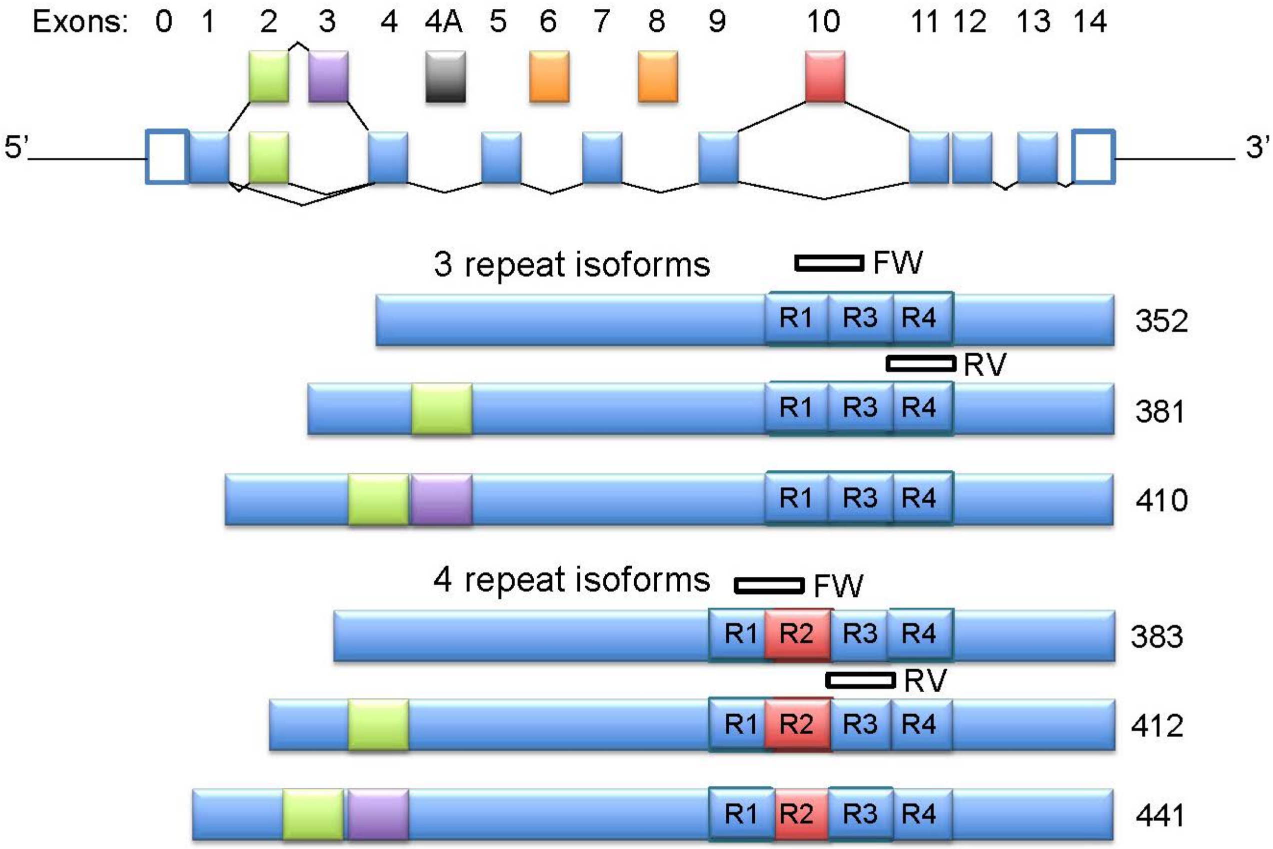

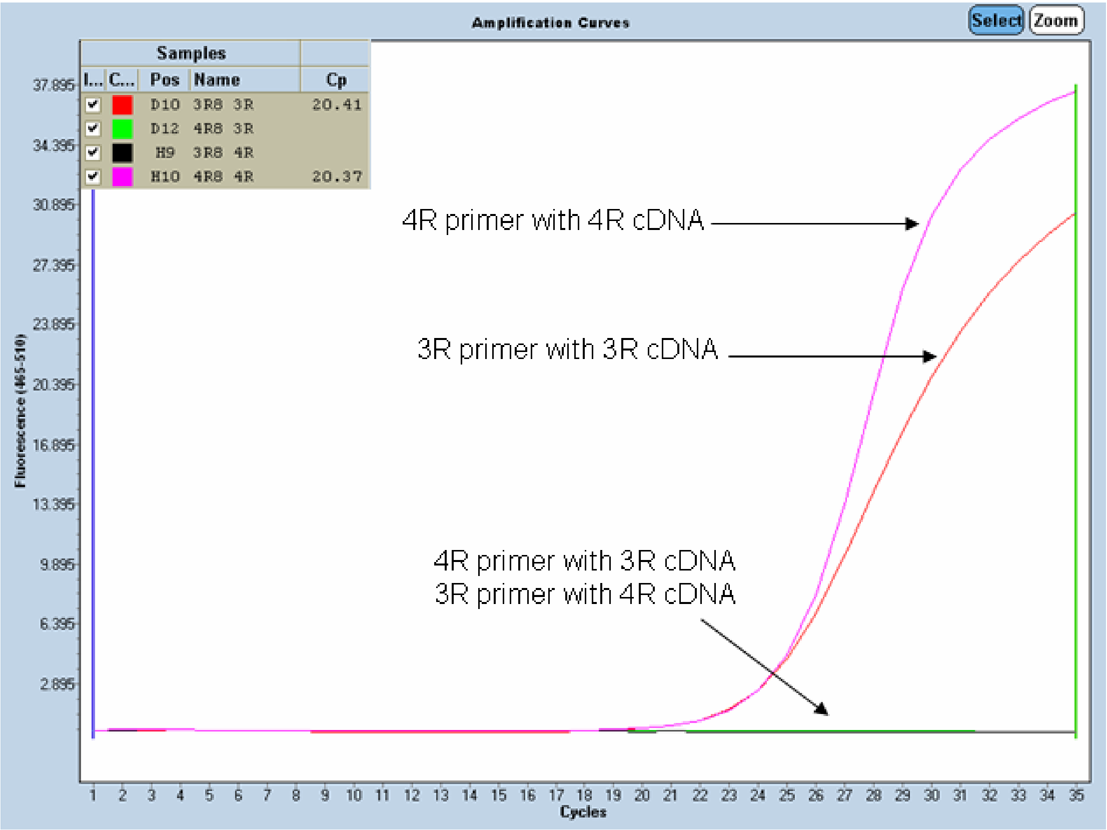
