## Supplementary material for "Regulation of neuronal mRNA splicing and Tau isoform ratio by ATXN3 through deubiquitylation of splicing factors": Figure S2

**a**

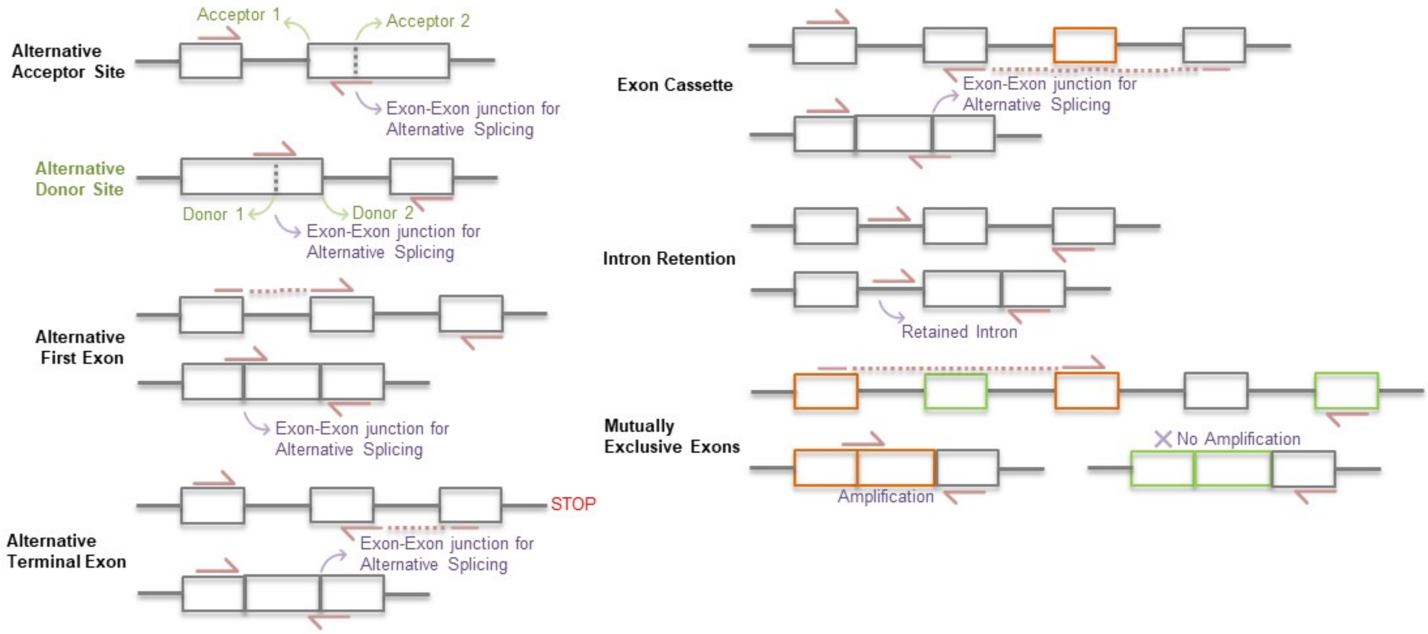


**b**

| Gene | Type | Direction | Exon(s) | Primer_Forward | Primer_Reverse | qRT-PCR Confirmation (Fold change relative to control) |
|---|---|---|---|---|---|---|
| **NEO1** | Alt. Acceptor Site | Down | e16-e17 | CTCGTGTTCCTGAAGTGCCT | CTTCAGAAGTGTCTGTGTGAGG | 0.002 |
| **EXOC7** | Alt. Acceptor Site | Down | e7-e8 | CGACTCCCTGATACAGGATGC | CTCAGGCTTGGTCTGCTTGA | 0.245 |
| **DOCK9** | Alt. Acceptor Site | Up | e42-43 | ATGTGTGCGGCTCTGTGTTA | AGGAGAAGCTGGTGTGCTTA | 1.405 |
| **ADD1** | Alt. Donor Site | Down | e11-e12 | AGGAAGGGCAGAATGGAAGC | TCCATCCTCTTTAGTCCACAAGC | 0.076 |
| **INPP4A** | Alt. Donor Site | Up | e17-e18 | CTGAACGTGGACAAGAGCCT | GGAGGGGGACTGCAATCTTT | 0.059 |
| **PPIE** | Alt. Donor Site | Up | e4-e5 | GCAGCTATCGACAACATGAATGA | CTCCTGGGTCTCTGCTTTGG | 1.967 |
| **EWSR1** | Alt. First Exon | Down | e1-8 | GACGGAACCATTCCAAACAGC | GGTGTAAGCACTGTAGCCCT | N/A |
| **TMEM181** | Alt. First Exon | Down | e1 | TACAGGCTGGCGCCCAT | CCGCACATTTCCCTGCACAT | 0.133 |
| **GPR18** | Alt. First Exon | Up | e1-e2 | GCAAAAGTCCACAAGCTCGATA | GTGTGGCTGTTTCAGTGCAG | 1.465 |
| **TMPO** | Alt. Terminal Exon | Up | e5 | CTTCAAAAGGCGGACCTCTG | GTCTGCGGCAACTAGCACTAA | 2.839 |
| **LRRFIP1** | Alt. Terminal Exon | Up | e21 | ATGCTCGAGGAAATCCGACA | GCGGAGCTCTCTTTGGAGTT | N/A |
| **RABGAP1L** | Alt. Terminal Exon | Down | e32/e33-37 | TGCTGCATATGCCAGAGGAA | TCTTGGAACAGTTCACCTGC | 0.779 |
| **PAPD4** | Complex | Down | e3 | GAATGCAGACTTGTCTAGAGCTG | GCCGTTTACCGTCAAGAGGA | 0.280 |
| **EEF1D** | Complex | Down | e1, e5 | CAGCCTGAGACCCAACAGAAA | CTCTGGCTCTCGCAATGTCA | 0.057 |
| **TPM1** | Complex | Up | e2-e3 | GCCGCCAAGGCTGAAG | CGGACTTGGCCTTCTGAGAG | Only detected in ATXN3$^{shRNA}$ cells |
| **VPS13C** | Exon Cassette | Down | e6,e7 | GGACATCAAGCCTGGACGTAA | GCAATCCAGTTTGGGCGTTT | Only detected in SCRshRNA cells |
| **EML4** | Exon Cassette | Up | e4 | AATTCGAGCATCACCTTCTCCC | TTCTCCTTCTTGGTTGATGATGAC | 1.126 |
| **SH3GLB1** | Exon Cassette | Down | e6,e7 | GCCAGGCTGAGAAGACAGAA | TGATGGGCATGTGTACTGCT | 0.728 |
| **MAPK8IP3** | Intron Retention | Up | e5 | CAGCTCCAGCTACCAGTGTC | TCCCCACTTCTTTGCCCATT | 0.020 |
| **SUPT7L** | Intron Retention | Up | e2 | CCTTTCGGGCAAGGATCTCA | AGACGTCGGTTGTGCTGAAT | 2.679 |
| **SRSF1** | Intron Retention | Down | e4 | ATCTCATGAGGGAGAAACTGCC | TGTTGCTTCTGCTACGGCTT | 0.293 |
| **MEAF6** | Mutually Exclusive Exons | Down | e7-e8 | TCTGTGCAGGGAGTGAAACC | GTGTCTTCTAATAGTCAGCTCGTG | 0.308 |
| **DLG1** | Mutually Exclusive Exons | Up | e23 | CAAGGACCAGAGTGAGCAGG | CAGTGTTTGCCCTTTTCTGC | 2.294 |
| **FYN** | Mutually Exclusive Exons | Down | e11 | CCTTGAGGAAGCGCAGATCA | AAACTTCCTTTGTTCATATACTCGG | 0.188 |
