## Supplementary material for "Regulation of neuronal mRNA splicing and Tau isoform ratio by ATXN3 through deubiquitylation of splicing factors": Table S1

| **Name** | **ATXN3$^{shRNA}$** | **p value** |
|---|---|---|
| **Network 1 – Gene expression, DNA replication and repair** | | |
| PGRMC1 (Membrane-associated progesterone receptor) | | 1.12E-14 |
| RANBP2 (E3 SUMO-protein ligase) | | 6.16E-06 |
| NASP (Nuclear autoantigenic sperm protein) | | 6.21E-06 |
| PURB (Transcriptional activator protein Pur-beta) | | 6.22E-06 |
| EHMT2 (Uncharacterized protein) | | 6.22E-06 |
| CAP1 (Adenylyl cyclase-associated protein 1) | | 5.54E-05 |
| TNPO1 (Transportin-1) | | 9.80E-05 |
| DR1 (Protein Dr1) | | 3.00E-04 |
| TRIM28 (Transcription intermediary factor 1-beta) | | 2.00E-03 |
| EIF6 (Eukaryotic translation initiation factor 6) | | 2.00E-03 |
| PLEC (Plectin) | | 4.00E-03 |
| SNRPN (Small nuclear ribonucleoprotein-associated protein N) | | 5.00E-03 |
| SFRS9 (Serine/arginine-rich splicing factor 9) | \$ | 7.00E-03 |
| HDAC2 (Histone deacetylase 2) | | 9.00E-03 |
| POLR2E (DNA-directed RNA polymerases I, II, and III subunit) | | 1.00E-02 |
| HSP90B1 (Endoplasmin) | | 1.00E-02 |
| SMARCB1 (Integraseinteractor 1b protein) | | 2.00E-02 |
| UBC (Ubiquitin C splice variant) | | 2.00E-02 |
| SKIV2L2 (Superkillerviralicidic activity 2-like 2) | \$ | 3.00E-02 |
| FUS (RNA-binding protein) | \$ | 3.00E-02 |
| HNRNPK (Heterogeneous nuclear ribonucleoprotein K) | \$ | 3.00E-02 |
| FLOT1 (Flotillin-1) | | 4.00E-02 |
| CBX5 (Uncharacterized protein) | | 4.00E-02 |
| GNB2L1 (Guanine nucleotide-binding protein subunit beta-2) | | 4.00E-02 |
| SMARCC2 (SWI/SNF complex subunit SMARCC2) | | 4.00E-02 |
| TPM3 (Isoform 2 of Tropomyosin alpha-3 chain) | | 4.00E-02 |
| C14orf166 (UPF0568 protein) | | 4.00E-02 |
| ATRX (Transcriptional regulator) | | 4.00E-02 |
| GAPDH (Glyceraldehyde-3-phosphate dehydrogenase) | | 5.00E-02 |
| **Newtwork 2 – RNA pot-transcriptional modification** | | |
| RPS10 (RPS10-NUDT3 protein) | | 6.22E-06 |
| C1QBP (Complement 1Q subcomponent-binding protein) | | 8.00E-04 |
| BPTF (Nucleosome-remodeling factor subunit) | | 1.00E-03 |
| PRPF8 (Pre-mRNA-processing-splicing factor 8) | \$ | 1.00E-03 |
| SFRS5 (Serine/arginine-rich splicing factor 5) | \$ | 5.00E-03 |
| EMD (Emerin) | | 9.00E-03 |
| HMGB3 (Uncharacterized protein) | | 1.00E-02 |
| NUP205 (Nuclear pore complex protein) | | 1.00E-02 |
| PRKCA (Protein kinase C alpha type) | | 2.00E-02 |
| HNRNPA1P10 (Heterogeneous nuclear ribonucleoprotein A1) | | 2.00E-02 |
| LRPPRC (Leucine-rich PPR motif-containing protein) | | 2.00E-02 |
| AQR (Intron-binding protein aquarius) | | 2.00E-02 |
| CAPZA1 (F-actin-capping protein subunit alpha-1) | | 3.00E-02 |

| | | |
|---|---|---|
| EEF1B2 (Elongation factor 1-beta) | | 3.00E-02 |
| RPL9 (60S ribosomal protein L9) | | 3.00E-02 |
| RPL18 (Uncharacterized protein) | | 4.00E-02 |
| SRRT (Serrate RNA effector molecule) | | 4.00E-02 |
| RBMX (Heterogeneous nuclear ribonucleoprotein G) | $ | 4.00E-02 |
| CNTN1 (Contactin-1) | | 4.00E-02 |
| THOC2 (THO complex subunit) | | 6.00E-02 |
| IARS (Uncharacterized protein) | | 8.00E-02 |
| **Network 3 – Moleculr transport, RNA trafficking** | | |
| DBN1 (Uncharacterized protein) | | 6.22E-06 |
| UQCRC2 (Cytochrome b-c1 complex subunit 2) | | 1.38E-05 |
| CANX (Calnexin) | | 2.00E-04 |
| SPTBN1 (Spectrin beta chain) | | 1.00E-03 |
| CALR (Calreticulin) | | 2.00E-03 |
| KIF5C (Kinesin heavy chain isoform 5C) | | 2.00E-03 |
| PSMA4 (Proteasome subunit alpha type-4) | | 2.00E-03 |
| RAE1 (Uncharacterized protein) | | 2.00E-03 |
| CAND1 (Cullin-associated NEDD8-dissociated protein 1) | | 3.00E-03 |
| YWHAG (14-3-3 protein gamma) | | 5.00E-03 |
| PDIA3 (Protein disulfide-isomerase A3) | | 1.00E-02 |
| NUP107 (Nuclear pore complex protein) | | 1.00E-02 |
| NUP160 (Nuclear pore complex protein) | | 1.00E-02 |
| PTPLAD1 (Butyrate-induced transcript 1) | | 3.00E-02 |
| PSMC5 (26S protease regulatory subunit 8) | | 4.00E-02 |
| CTPS (CTP synthase 1) | | 4.00E-02 |
| **Network 4 – Cell death and survival** | | |
| PSME2 (Uncharacterized protein) | | 1.78E-16 |
| GPI (Glucose-6-phosphate isomerase) | | 1.54E-05 |
| ASNS (Asparagine synthetase) | | 2.00E-04 |
| ESYT1 (Uncharacterized protein) | | 2.00E-04 |
| ATAD3A (ATPase family AAA domain-containing protein 3A) | | 8.00E-04 |
| ANXA5 (Annexin A5) | | 1.00E-03 |
| HSPH1 (Heat-shock protein 105 kDa) | | 2.00E-03 |
| RCC1 (Regulator of chromosome condensation) | | 2.00E-03 |
| HSPD1 (60 kDa heat shock protein) | | 2.00E-03 |
| PRDX1 (Uncharacterized protein) | | 2.00E-02 |
| VARS (Valyl-tRNAsynthetase) | | 4.00E-02 |
| EIF4A1 (Eukaryotic initiation factor 4A-I) | | 4.00E-02 |
| RARS (Isoform Monomeric of Arginyl-tRNAsynthetase) | | 5.00E-02 |
| RPL21 (60S ribosomal protein L21) | | 5.00E-02 |
| **Network 5 –Organ morphology** | | |
| TBL2 (Uncharacterized protein) | | 7.59E-14 |
| PROSC (Prolinesynthetase co-transcribed) | | 1.19E-13 |
| GSR (Glutathione reductase) | | 6.22E-06 |
| BCLAF1 (Bcl-2-associated transcription factor 1) | | 4.71E-05 |
| PFKM (6-phosphofructokinase) | | 3.00E-03 |
| PDIA6 (Protein disulfide-isomerase A6) | | 4.00E-03 |
| NAP1L1 (Nucleosome assembly protein 1-like 1) | | 7.00E-03 |

| | | |
|---|---|---|
| CKB (Creatine kinase B-type) | | 1.00E-02 |
| AHCY (Adenosylhomocysteinase) | | 3.00E-02 |
| PAICS (Multifunctional protein ADE2) | | 3.00E-02 |
| UBA1 (Ubiquitin-like modifier-activating enzyme 1) | | 3.00E-02 |
| RFC4 (Uncharacterized protein) | | 3.00E-02 |
| MDH2 (Malate dehydrogenase) | | 4.00E-02 |
| MYH10 (Uncharacterized protein) | | 4.00E-02 |
| **Network 6 - DNA replication, recombination and repair, cell death and Survival** | | |
| ATXN10 (Ataxin-10) | | 1.78E-16 |
| SNRPB2 (U2 small nuclear ribonucleoprotein B) | | 3.26E-11 |
| SFRS2 (Splicing factor arginine/serine-rich 2) | $ | 5.36E-11 |
| RBM8A (RNA-binding protein 8A) | | 1.56E-06 |
| MDC1 (Uncharacterized protein) | | 6.22E-06 |
| SFRS7 (Serine/arginine-rich splicing factor 7) | $ | 8.00E-04 |
| NHP2 (Uncharacterized protein) | | 3.00E-03 |
| TFAP2B (Isoform 2 of Transcription factor AP-2-beta) | | 2.00E-02 |
| MAP1B (Microtubule-associated protein 1B) | | 2.00E-02 |
| CRKL (Crk-like protein) | | 2.00E-02 |
| CAD (Uncharacterized protein) | | 4.00E-02 |
| VIM (Vimentin) | | 4.00E-02 |
| **Network 7 - Cell cycle, Cell death and survival** | | |
| CS (Citrate synthase) | | 1.78E-16 |
| UBXN1 (UBX domain-containing protein 1) | | 1.78E-16 |
| TUBA4A (Tubulin alpha-4 chain) | | 3.00E-03 |
| SAFB (Uncharacterized protein) | | 4.00E-03 |
| ACLY (Uncharacterized protein) | | 8.00E-03 |
| BRD1 (Bromodomain-containing protein 1) | | 1.00E-02 |
| SON (Isoform C of Protein SON) | | 1.00E-02 |
| PHB2 (Prohibitin-2) | | 2.00E-02 |
| PRPF40A (Pre-mRNA-processing factor 40 homolog A) | $ | 2.00E-02 |
| PHB (Prohibitin) | | 3.00E-02 |
| HSP90AB1 (Heat shock protein HSP 90-beta) | | 3.00E-02 |
| WDR3 (WD repeat-containing protein 3) | | 4.00E-02 |
| **Network 8 - Cellular compromise, Cell death and survival** | | |
| MYL6 (Myosin light polypeptide 6) | | 7.60E-14 |
| ENO2 (Enolase) | | 6.16E-06 |
| ERLIN2 (Uncharacterized protein) | | 2.48E-05 |
| RPL23A (60S ribosomal protein) | | 5.63E-05 |
| CLIC1 (Chloride intracellular channel protein 1) | | 1.00E-03 |
| ATAD2B (ATPase family AAA domain-containing protein 2B) | | 1.00E-03 |
| ATP5B (ATP synthase subunit beta) | | 7.00E-03 |
| CDK5 (Cyclin-dependent kinase 5) | | 9.00E-03 |
| SPTAN1 (Spectrin alpha chain) | | 2.00E-02 |
| SPIN1 (Spindlin-1) | | 2.00E-02 |
| GNAO1 (Guanine nucleotide-binding protein G(o) subunit alpha) | | 4.00E-02 |
| **Network 9 - Cellular growth and proliferation, Gene expression, Cell cycle** | | |
| GNAL (Guanine nucleotide-binding protein G(olf) subunit alpha) | | 2.05E-10 |
| EIF3L (Eukaryotic translation initiation factor 3) | | 6.22E-06 |

| | | |
|---|---|---|
| EIF3B (Uncharacterized protein) | | 6.22E-06 |
| TMEM33 (Transmembrane protein 33) | | 6.22E-06 |
| RAB5C (Uncharacterized protein) | | 9.71E-05 |
| SIX6 (Homeobox protein) | | 9.80E-05 |
| RPA1 (Replication protein A 70 kDa) | | 2.00E-02 |
| COPA (Coatomer subunit alpha) | | 2.00E-02 |
| MAB21L1 (Protein mab-21-like 1) | | 2.00E-02 |
| RAB1B (Ras-related protein) | | 2.00E-02 |
| RBM12B (RNA-binding protein 12B) | | 2.00E-02 |
| **Network 10 . Cellular development, Cell death and survival, Cell cycle** | | |
| RBM10 (RNA-binding protein 10) | | 4.78E-19 |
| CCT8 (T-complex protein 1 subunit theta) | | 5.72E-08 |
| TCP1 (T-complex protein 1 subunit alpha) | | 2.77E-06 |
| PSMA1 (Proteasome subunit alpha type-1) | | 6.22E-06 |
| MAP1S (BPY2 interacting protein 1) | | 5.43E-05 |
| GLTSCR2 (Uncharacterized protein) | | 9.80E-05 |
| CCT5 (T-complex protein 1 subunit epsilon) | | 2.16E-04 |
| GDI1 (Rab GDP dissociation inhibitor alpha) | | 9.16E-04 |
| TUBB2C (Tubulin beta-2C chain) | | 4.76E-02 |
| CCT4 (T-complex protein 1 subunit delta) | | 4.78E-02 |
| **Other** | | |
| CUTA (Isoform A of Protein CutA) | | 7.59E-14 |
| GNAI3 (Guanine nucleotide-binding protein G(k) subunit alpha) | | 7.59E-14 |
| TRAP1 (Uncharacterized protein) | | 6.58E-13 |
| ABT1 (Activator of basal transcription 1) | | 2.34E-12 |
| PSMA8 (Proteasome subunit alpha type-7-like) | | 7.71E-12 |
| POTEE (POTE ankyrin domain family member E) | | 7.49E-11 |
| C9orf114 (Uncharacterized protein) | | 1.56E-06 |
| SEPT7 (Uncharacterized protein) | | 6.21E-06 |
| HMBOX1 (Uncharacterized protein) | | 6.21E-06 |
| GLOD4 (CGI-150 protein) | | 6.22E-06 |
| DDX41 (DEAD-box protein abstrakt variant) | | 6.22E-06 |
| DDX56 (ATP-dependent RNA helicase) | | 2.38E-05 |
| PSMA8 (Uncharacterized protein) | | 2.48E-05 |
| TOMM22 (Mitochondrial import receptor subunit) | | 2.54E-05 |
| HIST2H2BD (Histone H2B type 2-D) | | 9.17E-05 |
| BLVRA (Biliverdinreductase A) | | 2.00E-04 |
| ACTL6B (Actin-like protein 6B) | | 3.00E-04 |
| LUC7L (Putative RNA-binding protein Luc7-like 1) | | 5.00E-04 |
| CHMP5 (Charged multivesicular body protein 5) | | 6.00E-04 |
| ISOC1 (Isochorismatase domain-containing protein 1) | | 7.00E-04 |
| RPL24 (60S ribosomal protein L24) | $ | 2.00E-03 |
| ABCF1 (ATP-binding cassette sub-family F member 1) | | 3.00E-03 |
| BMS1 (Ribosome biogenesis protein) | | 4.00E-03 |
| PDS5B (Sister chromatid cohesion protein) | | 4.00E-03 |
| PSMD1 (26S proteasome non-ATPase regulatory subunit 1) | | 5.00E-03 |
| DIMT1 (Probable dimethyladenosinetransferase) | | 6.00E-03 |
| WDR75 (WD repeat-containing protein 75) | | 7.00E-03 |

| | |
|---|---|
| DDX10 (Probable ATP-dependent RNA helicase) | 1.00E-02 |
| DPM1 (Uncharacterized protein) | 1.00E-02 |
| SLC25A11 (Mitochondrial 2-oxoglutarate/malate carrier) | 1.00E-02 |
| FN3K (Fructosamine-3-kinase) | 2.00E-02 |
| ZNF828 (ZNF828 Zinc finger protein 828) | 2.00E-02 |
| DDX49 (Probable ATP-dependent RNA helicase) | 2.00E-02 |
| EIF5B (Eukaryotic translation initiation factor 5B) | 2.00E-02 |
| RTL1 (Retrotransposon-like protein 1) | 3.00E-02 |
| RANP1 (Uncharacterized protein) | 3.00E-02 |
| DHX8 (Uncharacterized protein) | 3.00E-02 |
| MYEF2 (Myelin expression factor 2) | 4.00E-02 |
| HMGA1 (High mobility group protein HMG-I/HMG-Y) | 4.00E-02 |
| PELP1 (Proline-, glutamic acid-, leucine-rich protein 1) | 4.00E-02 |
