## Supplementary material for "Regulation of neuronal mRNA splicing and Tau isoform ratio by ATXN3 through deubiquitylation of splicing factors": Table S2

| Alternative Event Type | Number of Regulated Events |
|---|---|
| **Alter. First Exon** | 742 |
| **Alter. Terminal Exon** | 494 |
| **Exon Cassette** | 1248 |
| **Mutually Exclusive Exons** | 37 |
| **Alter. Acceptor Splice Site** | 73 |
| **Alter. Donor Splice Site** | 72 |
| **Intron Retention** | 438 |
| **Complex** | 580 |

### List of the 3,684 Differentially Regulated Alternative Splicing Events (from 1,993 Distinct Genes)

| Gene Symbol | Alternative Event Type | Involved exon(s) | Regulation Splicing Index | Average Splicing-Index Fold-Change | EnsEMBL ID |
|---|---|---|---|---|---|
| PRSS3 | Alter. First Exon | e1,e3/e2 | up (e1,e3) | 25,11 | ENSG00000 010438 |
| FOXO3 | Alter. First Exon | e3/e5 | down (e5) | 20,18 | ENSG00000 118689 |
| PTPRS | Exon Cassette | e15 | down | 17,65 | ENSG00000 105426 |
| IFI30 // PIK3R2 | Alter. First Exon | e1-19/e20 | down (e1-19) | 14,96 | ENSG00000 105647 // ENSG00000 216490 |
| NPFFR2 | Alter. First Exon | e1/e3-4 | up (e1) | 14,72 | ENSG00000 056291 |
| NPFFR2 | Alter. First Exon | e1/e3 | up (e1) | 14,72 | ENSG00000 056291 |
| FAM163A | ter. Terminal Exo | e7/e8,e9 | down (e8,e9) | 14,59 | ENSG00000 143340 |
| PRSS3 | Alter. First Exon | e1/e2 | up (e1) | 14,41 | ENSG00000 010438 |
| FOXO3 | Alter. First Exon | e2/e5 | up (e2) | 14,11 | ENSG00000 118689 |
| TUBB3 | Alter. First Exon | e4-8/e5-6,e9 | up (e4-8) | 14,05 | ENSG00000 198211 |
| // NME1-NME2 // | Complex | e8 | down | 13,82 | ENSG00000 011052 // ENSG00000 239672 // ENSG00000 243678 |

| | | | | | |
|---|---|---|---|---|---|
| F6 // HEXA // PA | Exon Cassette | e53 | down | 13,77 | ENSG00000137817 // ENSG00000140488 // ENSG00000213614 |
| OLFM3 | Exon Cassette | e6 | up | 13,23 | ENSG00000118733 |
| IFI30 // PIK3R2 | Alter. First Exon | e1-18/e19 | down (e1-18) | 13,13 | ENSG00000105647 // ENSG00000216490 |
| LINC01021 | Complex | e2/e5,e6-7 | down (e5,e6-7) | 13,04 | ENSG00000250337 |
| LINC01021 | Exon Cassette | e5-6 | down | 12,45 | ENSG00000250337 |
| GPC3 | Alter. First Exon | e1/e4 | down (e1) | 12,44 | ENSG00000147257 |
| VPS13C | Exon Cassette | e6,e7 | down | 12,31 | ENSG00000129003 |
| DRAM2 | Complex | e1,e2 | up | 12,17 | ENSG00000156171 |
| DRAM2 | Exon Cassette | e2 | up | 12,01 | ENSG00000156171 |
| TPD52L2 | Exon Cassette | e6,e7 | down | 11,59 | ENSG00000101150 |
| TUBB3 | Complex | e9-11 | up | 11,53 | ENSG00000198211 |
| NPNT | Exon Cassette | e6 | down | 11,3 | ENSG00000168743 |
| EWSR1 | Alter. First Exon | e1-8/e9 | down (e1-8) | 11,13 | ENSG00000182944 |
| TPD52L2 | Exon Cassette | e7 | down | 10,73 | ENSG00000101150 |
| F2 // INS // INS-I | ter. Terminal Exo | 6/e11,e12-1 | down (e11,e12-13) | 10,64 | ENSG00000129965 // ENSG00000167244 // ENSG00000254647 |
| STMN2 | Exon Cassette | e7 | up | 10,17 | ENSG00000104435 |
| TUBB3 | Complex | e9-10 | up | 10,15 | ENSG00000198211 |
| EML4 | Exon Cassette | e4 | up | 9,66 | ENSG00000143924 |
| EPB41L2 | Exon Cassette | e21 | down | 9,63 | ENSG00000079819 |
| MAPK8IP3 | Intron Retention | e5 | up | 9,59 | ENSG00000138834 |
| SH3GLB1 | Exon Cassette | e6,e7 | down | 9,48 | ENSG00000097033 |

| | | | | | |
|---|---|---|---|---|---|
| DMKN | Exon Cassette | e11-12 | down | 9,46 | ENSG00000 161249 |
| ZNF415 | Exon Cassette | e7 | up | 9,39 | ENSG00000 170954 |
| CNTNAP2 | Alter. First Exon | -11,e13-17/e | down (e1-11,e13-17 | 9,38 | ENSG00000 174469 |
| PAPD4 | Complex | e3/e4 | down (e3) | 9,26 | ENSG00000 164329 |
| PYGL | Exon Cassette | e2 | up | 9,18 | ENSG00000 100504 |
| MEAF6 | Exon Cassette | e7 | down | 9,15 | ENSG00000 163875 |
| SORBS2 | Complex | e19-22 | up | 9,13 | ENSG00000 154556 |
| CNTNAP2 | Alter. First Exon | e12-17/e18 | down (e12-17) | 8,91 | ENSG00000 174469 |
| EEF1D | Complex | e1,e5 | down | 8,74 | ENSG00000 104529 |
| --- | Alter. First Exon | e2-8/e9 | up (e2-8) | 8,73 | --- |
| EEF1D | Exon Cassette | e3,e5 | down | 8,6 | ENSG00000 104529 |
| SLC47A1 | Alter. First Exon | e1/e2 | down (e2) | 8,55 | ENSG00000 142494 |
| DYNC1I2 | Exon Cassette | e7 | down | 8,47 | ENSG00000 077380 |
| ZNF20 // ZNF625 | Alter. First Exon | e1-3,e5/e6 | down (e1-3,e5) | 8,44 | ENSG00000 132010 // ENSG00000 257591 |
| DCBLD1 | Alter. First Exon | e1/e2 | down (e1) | 8,42 | ENSG00000 164465 |
| SGIP1 | Exon Cassette | e7 | up | 8,34 | ENSG00000 118473 |
| PLEKHA6 | Exon Cassette | e10,e11-12 | up | 8,31 | ENSG00000 143850 |
| SLCO3A1 | Complex | e16,e17 | up | 8,28 | ENSG00000 176463 |
| SORBS2 | Complex | e20-22 | up | 8,27 | ENSG00000 154556 |
| PLEKHA6 | Exon Cassette | e11,e12 | up | 8,24 | ENSG00000 143850 |
| ZNF454 | Alter. First Exon | e1/e2 | up (e1) | 8,16 | ENSG00000 178187 |
| NEO1 | lter. Acceptor Sit | e17 | down | 7,98 | ENSG00000 067141 |
| LMO3 | Alter. First Exon | e1/e3 | up (e3) | 7,88 | ENSG00000 048540 |
| SORBS2 | Exon Cassette | e19,e20 | up | 7,86 | ENSG00000 154556 |
| PLEKHA6 | Exon Cassette | e10,e11 | up | 7,8 | ENSG00000 143850 |

| | | | | | |
|---|---|---|---|---|---|
| EPB41L2 | Complex | e21 | up | 7,69 | ENSG00000079819 |
| TPD52L2 | Exon Cassette | e6-8 | down | 7,62 | ENSG00000101150 |
| PLEKHA6 | Exon Cassette | e11-12 | up | 7,6 | ENSG00000143850 |
| SLCO3A1 | ter. Terminal Exo | e16-17/e18 | up (e16-17) | 7,53 | ENSG00000176463 |
| MEAF6 | Exon Cassette | e8 | up | 7,44 | ENSG00000163875 |
| DYX1C1 // DYX1 | ter. Terminal Exo | 0/e15,e16-2 | up (e15,e16-22) | 7,39 | ENSG00000256061 // ENSG00000260916 // ENSG00000261771 |
| PTPRF | Exon Cassette | e16 | down | 7,31 | ENSG00000142949 |
| ZNF20 // ZNF625 | Alter. First Exon | e1-5/e6 | down (e1-5) | 7,17 | ENSG00000132010 // ENSG00000257591 |
| PDE4B | Complex | e1/e3-4 | up (e3-4) | 7,15 | ENSG00000184588 |
| CD99 | ter. Terminal Exo | e2/e3-11 | down (e3-11) | 7,14 | ENSG00000002586 |
| HAND2-AS1 | Complex | e6 | up | 7,13 | ENSG00000237125 |
| KCNMA1 | ter. Terminal Exo | e29-30,e32 | ,e25,e29-30,e32-34 | 7 | ENSG00000156113 |
| ETV4 | Exon Cassette | e3 | down | 7 | ENSG00000175832 |
| KCNH2 | Complex | 6,e7-9,e15-1 | down | 6,97 | ENSG00000055118 |
| ADD1 | Exon Cassette | e18 | down | 6,87 | ENSG00000087274 |
| KCNMA1 | ter. Terminal Exo | e28-30,e32- | ,e25,e28-30,e32-34 | 6,81 | ENSG00000156113 |
| GRIK2 | Exon Cassette | e19 | up | 6,77 | ENSG00000164418 |
| CD99 | ter. Terminal Exo | 2/e3-8,e10-1 | down (e3-8,e10-11) | 6,71 | ENSG00000002586 |
| FGFR2 | Complex | e15/e16 | down (e15) | 6,62 | ENSG00000066468 |
| CD99 | ter. Terminal Exo | 2/e3,e4-8,e1 | down (e3,e4-8,e10) | 6,59 | ENSG00000002586 |
| CLSTN2 | Complex | e2,e3 | down | 6,58 | ENSG00000158258 |
| RIIAD1 | Alter. First Exon | e1-5/e6 | up (e1-5) | 6,53 | ENSG00000178796 |
| SPAG9 | Alter. First Exon | e1-4/e5 | up (e1-4) | 6,5 | ENSG00000008294 |

| | | | | | |
|---|---|---|---|---|---|
| F2 // INS // INS-I( | Alter. First Exon | e1-2/e4 | up (e1-2) | 6,48 | ENSG00000129965 // ENSG00000167244 // ENSG00000254647 |
| DLG1 | Exon Cassette | e26 | down | 6,4 | ENSG00000075711 |
| ZNF423 | Alter. First Exon | e3-4/e6 | up (e3-4) | 6,39 | ENSG00000102935 |
| FAT1 | Exon Cassette | e28-29 | down | 6,37 | ENSG00000083857 |
| TMEM181 | Alter. First Exon | e1/e2-4 | down (e1) | 6,34 | ENSG00000146433 |
| PLEKHA6 | Exon Cassette | e10-11 | up | 6,31 | ENSG00000143850 |
| PAX6 | Intron Retention | e10 | down | 6,3 | ENSG00000007372 |
| PAX6 | Alter. Donor Site | e10 | down | 6,28 | ENSG00000007372 |
| --- | Complex | e2,e4,e6 | down | 6,28 | --- |
| MDM2 | Alter. First Exon | e1-4/e3 | up (e3) | 6,25 | ENSG00000135679 |
| PDZRN3 | Alter. First Exon | e1-3/e7 | up (e1-3) | 6,25 | ENSG00000121440 |
| GPR19 | Alter. First Exon | e1-2/e5 | up (e1-2) | 6,22 | ENSG00000183150 |
| ULBP3 | Complex | e5-6 | down | 6,21 | ENSG00000131019 |
| KCNH2 | Complex | e6-16 | down | 6,2 | ENSG00000055118 |
| ARSG | Alter. First Exon | e1/e2 | down (e1) | 6,15 | ENSG00000141337 |
| PDE4B | Alter. First Exon | e1-4/e5 | up (e1-4) | 6,11 | ENSG00000184588 |
| REST | Complex | e10-11 | down | 6,11 | ENSG00000084093 |
| LRRN1 | Alter. First Exon | e1/e2,e3 | up (e1) | 6,09 | ENSG00000175928 |
| TPD52L1 | Alter. First Exon | e1/e3 | down (e1) | 6,09 | ENSG00000111907 |
| SNAP91 | Exon Cassette | e4-6 | up | 6,09 | ENSG00000065609 |
| IQCJ-SCHIP1 // : | Exon Cassette | e8 | up | 6,05 | ENSG00000151967 // ENSG00000214216 // ENSG00000250588 |
| ELMO1 | Complex | e19/e22 | up (e19) | 6,02 | ENSG00000155849 |

| | | | | | |
|---|---|---|---|---|---|
| MACF1 | Exon Cassette | e66 | up | 5,99 | ENSG00000127603 |
| WLS | Exon Cassette | e12 | down | 5,99 | ENSG00000116729 |
| L3MBTL3 | Exon Cassette | e5 | down | 5,99 | ENSG00000198945 |
| CNR1 | Complex | e1/e2,e4 | up (e2,e4) | 5,98 | ENSG00000118432 |
| CNR1 | Exon Cassette | e2,e4 | up | 5,98 | ENSG00000118432 |
| SLCO3A1 | Alter. First Exon | e2-4/e5 | up (e2-4) | 5,95 | ENSG00000176463 |
| RT222 // SMARC | Exon Cassette | e18 | up | 5,92 | ENSG00000073584 // ENSG00000213424 |
| DKK3 | Alter. First Exon | e2/e4 | up (e2) | 5,91 | ENSG00000050165 |
| WDR35 | Exon Cassette | e11 | down | 5,9 | ENSG00000118965 |
| PRKCH | Alter. First Exon | e1-11/e12 | up (e1-11) | 5,88 | ENSG00000027075 |
| KIF2A | Exon Cassette | e19 | down | 5,88 | ENSG00000068796 |
| PRKCH | Alter. First Exon | 3,e5-11/e12 | up (e3,e5-11) | 5,87 | ENSG00000027075 |
| TMEM181 | Alter. First Exon | e1/e2,e3 | down (e1) | 5,87 | ENSG00000146433 |
| DPYSL3 | Complex | e2-11 | down | 5,8 | ENSG00000113657 |
| ZNF667 | Alter. First Exon | e1-2/e3 | up (e1-2) | 5,77 | ENSG00000198046 |
| PRKCH | Alter. First Exon | 2,e4-11/e12 | up (e2,e4-11) | 5,74 | ENSG00000027075 |
| TPD52L2 | Exon Cassette | e7-8 | down | 5,74 | ENSG00000101150 |
| LINGO2 | Complex | e2-6/e3-5 | up (e3-5) | 5,74 | ENSG00000174482 |
| PRRG1 | Complex | e1,e2 | down | 5,72 | ENSG00000130962 |
| PHF21A | ualy Exclusive Ex | e15/e16 | up (e16) | 5,71 | ENSG00000135365 |
| HLA-F | ter. Terminal Exo | e6/e8,e9 | down (e8,e9) | 5,7 | ENSG00000204642 |
| BCLAF1 | Exon Cassette | e7 | up | 5,68 | ENSG00000029363 |
| GPC3 | Exon Cassette | e2 | up | 5,68 | ENSG00000147257 |
| SCAMP5 | ualy Exclusive Ex | e2/e3 | up (e2) | 5,63 | ENSG00000198794 |
| SYT5 | Exon Cassette | e3 | down | 5,63 | ENSG00000129990 |

| | | | | | |
|---|---|---|---|---|---|
| GDAP1L1 | Alter. First Exon | e1/e2 | up (e2) | 5,59 | ENSG00000124194 |
| UNC5A | Exon Cassette | e3,e4 | up | 5,59 | ENSG00000113763 |
| KLHL13 | Alter. First Exon | e3-4/e6 | up (e3-4) | 5,58 | ENSG00000003096 |
| MADD | Exon Cassette | e27 | down | 5,49 | ENSG00000110514 |
| F2 // INS // INS-I | ter. Terminal Exo | e6/e11-13 | down (e11-13) | 5,49 | ENSG00000129965 // ENSG00000167244 // ENSG00000254647 |
| ATCAY | Complex | e4-6 | up | 5,47 | ENSG00000167654 |
| MAPK8 | ualy Exclusive Ex | e7/e8 | down (e7) | 5,44 | ENSG00000107643 |
| DMKN | Exon Cassette | e10,e12 | down | 5,42 | ENSG00000161249 |
| C7 | Alter. First Exon | e1/e2 | up (e1) | 5,42 | ENSG00000112936 |
| MEAF6 | ualy Exclusive Ex | e7/e8 | down (e7) | 5,41 | ENSG00000163875 |
| CADM2 | Complex | e9,e11 | down | 5,4 | ENSG00000175161 |
| PLAT | Complex | e7-8 | up | 5,33 | ENSG00000104368 |
| EEF1D | Exon Cassette | e4,e5 | down | 5,26 | ENSG00000104529 |
| NOTCH2 | Exon Cassette | e3 | up | 5,23 | ENSG00000134250 |
| ANK3 | Complex | e43-44 | down | 5,22 | ENSG00000151150 |
| DMKN | Exon Cassette | e5 | down | 5,22 | ENSG00000161249 |
| PDPN | Exon Cassette | e4 | down | 5,2 | ENSG00000162493 |
| CDH1 | Alter. First Exon | e1-2/e3 | down (e1-2) | 5,2 | ENSG00000039068 |
| CDH1 | Alter. First Exon | e1-2/e4 | down (e1-2) | 5,2 | ENSG00000039068 |
| PARP2 | Alter. Donor Site | e2 | up | 5,19 | ENSG00000129484 |
| SCAMP1 | Alter. First Exon | e1-7/e8 | down (e1-7) | 5,18 | ENSG00000085365 |
| SCAMP1 | Alter. First Exon | e1-5,e7/e8 | down (e1-5,e7) | 5,18 | ENSG00000085365 |
| SCAMP1 | Alter. First Exon | e1-3,e6-7/e8 | down (e1-3,e6-7) | 5,18 | ENSG00000085365 |
| CD55 | Exon Cassette | e9,e11 | down | 5,15 | ENSG00000196352 |

| | | | | | |
|---|---|---|---|---|---|
| STMN4 | Exon Cassette | e4 | up | 5,15 | ENSG00000 015592 |
| GPR160 | Exon Cassette | e3 | down | 5,14 | ENSG00000 173890 |
| NCOA7 | lter. Acceptor Sit | e12 | up | 5,14 | ENSG00000 111912 |
| ICAM2 | Complex | e1,e2 | up | 5,13 | ENSG00000 108622 |
| DDX17 | Intron Retention | e12,e13 | up | 5,12 | ENSG00000 100201 |
| LPHN2 | Exon Cassette | e26 | down | 5,11 | ENSG00000 117114 |
| TCF12 | Exon Cassette | e17 | down | 5,09 | ENSG00000 140262 |
| CLDN1 | Complex | e1/e2,e3-4 | up (e2,e3-4) | 5,08 | ENSG00000 163347 |
| ESRRG | Complex | e6,e13/e10 | up (e10) | 5,07 | ENSG00000 196482 |
| ESRRG | Exon Cassette | e10 | up | 5,07 | ENSG00000 196482 |
| PPFIA1 | Exon Cassette | e18 | down | 5,07 | ENSG00000 131626 |
| APP | Exon Cassette | e10-11 | up | 5,04 | ENSG00000 142192 |
| MAGED1 | Alter. First Exon | e1/e2 | down (e1) | 5,02 | ENSG00000 179222 |
| FHL1 | Exon Cassette | e11 | down | 5,02 | ENSG00000 022267 |
| INPP5D | Alter. First Exon | e1-5/e6 | down (e1-5) | 5,01 | ENSG00000 168918 |
| MYT1L | Alter. First Exon | e1-5/e6 | down (e1-5) | 5 | ENSG00000 186487 |
| CADM2 | Exon Cassette | e10 | up | 4,99 | ENSG00000 175161 |
| SELENBP1 | Intron Retention | e9 | down | 4,98 | ENSG00000 143416 |
| ZNF423 | Alter. First Exon | e2,e4/e6 | up (e2,e4) | 4,98 | ENSG00000 102935 |
| ICAM2 | Alter. Donor Site | e1 | up | 4,98 | ENSG00000 108622 |
| ICAM2 | Complex | e1/e2 | up (e1) | 4,98 | ENSG00000 108622 |
| EPB41L3 | Complex | e22/e23,e25 | down (e23,e25) | 4,96 | ENSG00000 082397 |
| EPB41L2 | Exon Cassette | e19 | down | 4,96 | ENSG00000 079819 |
| MDM2 | Alter. First Exon | e2/e3 | down (e2) | 4,94 | ENSG00000 135679 |

| | | | | | |
|---|---|---|---|---|---|
| DYX1C1 // DYX1 | ter. Terminal Exo | 9,e15-22/e1 | up (e9,e15-22) | 4,94 | ENSG00000 256061 // ENSG00000 260916 // ENSG00000 261771 |
| CD47 | Exon Cassette | e9,e10 | down | 4,93 | ENSG00000 196776 |
| F2 // INS // INS-IC | ter. Terminal Exo | e12/e13 | down (e13) | 4,9 | ENSG00000 129965 // ENSG00000 167244 // ENSG00000 254647 |
| DKK3 | Alter. First Exon | e3/e4 | up (e3) | 4,9 | ENSG00000 050165 |
| CLASP1 | ualy Exclusive Ex | e22/e23 | up (e22) | 4,88 | ENSG00000 074054 |
| DKK3 | Alter. First Exon | e1/e4 | up (e1) | 4,86 | ENSG00000 050165 |
| NF1 | Exon Cassette | e32 | up | 4,86 | ENSG00000 196712 |
| RNF144A | Alter. First Exon | e1/e2 | up (e1) | 4,83 | ENSG00000 151692 |
| PPEF1 | Alter. First Exon | e1-3/e4 | up (e1-3) | 4,83 | ENSG00000 086717 |
| MYT1L | ter. Terminal Exo | e24/e25-28 | down (e25-28) | 4,82 | ENSG00000 186487 |
| SYNRG | Exon Cassette | e8 | up | 4,81 | ENSG00000 006114 |
| SH2D3C | Complex | e3,e9/e4-7 | up (e4-7) | 4,79 | ENSG00000 095370 |
| SMPDL3A | Alter. First Exon | e1/e3 | down (e1) | 4,78 | ENSG00000 172594 |
| RPL3 | lter. Acceptor Sit | e6 | up | 4,77 | ENSG00000 100316 |
| SH3GLB1 | Exon Cassette | e6 | down | 4,73 | ENSG00000 097033 |
| KN2B-AS1 // MT | Exon Cassette | e28,e30-31 | down | 4,72 | ENSG00000 099810 // ENSG00000 240498 |
| GALC | Alter. First Exon | e2/e3 | up (e2) | 4,7 | ENSG00000 054983 |
| CD55 | Exon Cassette | e9,e10 | down | 4,69 | ENSG00000 196352 |
| TUBB3 | Exon Cassette | e7 | up | 4,69 | ENSG00000 198211 |
| REST | Exon Cassette | e10 | down | 4,69 | ENSG00000 084093 |
| ZIC2 | Complex | e2,e3-4 | down | 4,68 | ENSG00000 043355 |

| | | | | | |
|---|---|---|---|---|---|
| ZNF160 | lter. Acceptor Sit | e3 | up | 4,64 | ENSG00000170949 |
| FN1 | Exon Cassette | e41 | down | 4,63 | ENSG00000115414 |
| FOXP2 | ter. Terminal Exo | 7/e21,e22-2 | down (e21,e22-25) | 4,62 | ENSG00000128573 |
| SORBS1 | Exon Cassette | e11,e12 | up | 4,6 | ENSG00000095637 |
| ZNF423 | Alter. First Exon | e1,e4/e6 | up (e1,e4) | 4,58 | ENSG00000102935 |
| PCSK5 | ter. Terminal Exo | /e7-21,e23-4 | own (e7-21,e23-39 | 4,57 | ENSG00000099139 |
| MLH1 | Exon Cassette | e3 | down | 4,56 | ENSG00000076242 |
| EPB41L2 | Exon Cassette | e19,e20 | down | 4,56 | ENSG00000079819 |
| SLC2A14 | Alter. First Exon | e2/e3-14 | down (e3-14) | 4,54 | ENSG00000173262 |
| REST | ter. Terminal Exo | -8,e11/e14- | up (e7-8,e11) | 4,54 | ENSG00000084093 |
| REST | ter. Terminal Exo | 8-11/e14-15 | up (e8-11) | 4,54 | ENSG00000084093 |
| REST | ter. Terminal Exo | -8,e11/e14- | up (e6-8,e11) | 4,54 | ENSG00000084093 |
| REST | ter. Terminal Exo | 8,e11/e13,e | up (e7-8,e11) | 4,54 | ENSG00000084093 |
| REST | ter. Terminal Exo | 8-11/e13,e1 | up (e8-11) | 4,54 | ENSG00000084093 |
| REST | ter. Terminal Exo | 8,e11/e13,e | up (e6-8,e11) | 4,54 | ENSG00000084093 |
| GIT2 | Exon Cassette | e17 | up | 4,53 | ENSG00000139436 |
| MYT1L | Alter. First Exon | e1-2,e4-5/e6 | down (e1-2,e4-5) | 4,53 | ENSG00000186487 |
| MCC | Alter. First Exon | e1-3/e5 | up (e1-3) | 4,52 | ENSG00000171444 |
| --- | Exon Cassette | e3,e4 | down | 4,52 | --- |
| GALC | Alter. First Exon | e1/e3 | up (e1) | 4,51 | ENSG00000054983 |
| TJAP1 | Exon Cassette | e10 | down | 4,51 | ENSG00000137221 |
| EYA1 | Intron Retention | e5,e6 | up | 4,49 | ENSG00000104313 |
| ADD3 | Exon Cassette | e16 | down | 4,47 | ENSG00000148700 |
| TMPO | Exon Cassette | e6,e7-8 | up | 4,45 | ENSG00000120802 |
| PRSS16 | Exon Cassette | e3-8 | down | 4,45 | ENSG00000112812 |
| CD47 | ter. Terminal Exo | 8/e9,e10-12 | down (e9,e10-12) | 4,44 | ENSG00000196776 |

| | | | | | |
|---|---|---|---|---|---|
| ANK3 | Exon Cassette | e44 | down | 4,39 | ENSG00000151150 |
| NCAM1 | Complex | e18/e19,e20 | up (e19,e20) | 4,39 | ENSG00000149294 |
| ANK2 | Complex | e42,e43 | down | 4,37 | ENSG00000145362 |
| NEDD4L | Alter. First Exon | e2-7/e9 | down (e2-7) | 4,34 | ENSG00000049759 |
| PDZRN3 | Alter. First Exon | e4/e7 | down (e7) | 4,34 | ENSG00000121440 |
| LINGO2 | Exon Cassette | e3-5 | up | 4,34 | ENSG00000174482 |
| YWHAQ | Complex | e1-2 | down | 4,31 | ENSG00000134308 |
| CASP8 | ter. Terminal Exo | 3,e16-18/e1 | down (e13,e16-18) | 4,29 | ENSG00000064012 |
| UNC5C | Exon Cassette | e9 | up | 4,29 | ENSG00000182168 |
| XIST | Complex | e9,e10 | up | 4,29 | ENSG00000229807 |
| TPM1 | Complex | e2,e3 | up | 4,27 | ENSG00000140416 |
| NEDD4L | Alter. First Exon | e1,e7/e9 | down (e1,e7) | 4,27 | ENSG00000049759 |
| NEDD4L | Alter. First Exon | e6-7/e9 | down (e6-7) | 4,27 | ENSG00000049759 |
| CAST | Alter. First Exon | e2/e3 | down (e2) | 4,27 | ENSG00000153113 |
| PPP2R3A | Exon Cassette | e9 | up | 4,26 | ENSG00000073711 |
| MAP7 | Alter. First Exon | e3/e4 | down (e3) | 4,23 | ENSG00000135525 |
| CD99 | Complex | e2/e3 | up (e2) | 4,23 | ENSG00000002586 |
| NEDD4L | Alter. First Exon | e6-7,e10/e9 | down (e6-7,e10) | 4,22 | ENSG00000049759 |
| YWHAQ | Complex | e1,e2 | down | 4,22 | ENSG00000134308 |
| ATL2 | Complex | e14,e15 | down | 4,22 | ENSG00000119787 |
| SORBS1 | Exon Cassette | e12 | up | 4,21 | ENSG00000095637 |
| SLCO3A1 | Alter. First Exon | e2,e4/e5 | up (e2,e4) | 4,21 | ENSG00000176463 |
| TGLF9P // PAR | Exon Cassette | e6 | down | 4,2 | ENSG00000227345 |
| W // SNORD116- | Intron Retention | e38,e39 | up | 4,18 | ENSG00000207137 |
| AEBP1 | Intron Retention | e12 | up | 4,17 | ENSG00000106624 |
| IGDCC4 | Alter. First Exon | e7-18/e19 | up (e7-18) | 4,15 | ENSG00000103742 |

| | | | | | |
|---|---|---|---|---|---|
| MCOLN3 | Exon Cassette | e5 | down | 4,14 | ENSG00000 055732 |
| CDH23 | Alter. First Exon | e49-66/e67 | down (e49-66) | 4,13 | ENSG00000 107736 |
| CDH23 | Alter. First Exon | e50-66/e67 | down (e50-66) | 4,13 | ENSG00000 107736 |
| SYNRG | Exon Cassette | e7-8 | up | 4,13 | ENSG00000 006114 |
| CCNJL | Exon Cassette | e10 | up | 4,13 | ENSG00000 135083 |
| ST7 // ST7-OT3 | Alter. First Exon | e1/e3 | up (e1) | 4,12 | ENSG00000 004866 |
| ERC1 | Exon Cassette | e19 | down | 4,1 | ENSG00000 082805 |
| SPINT2 | Exon Cassette | e2 | up | 4,1 | ENSG00000 167642 |
| GUCY1B3 | Exon Cassette | e2 | up | 4,09 | ENSG00000 061918 |
| FAM168A | Exon Cassette | e4 | down | 4,08 | ENSG00000 054965 |
| TMPO | ter. Terminal Exo | e5/e9-10 | up (e5) | 4,06 | ENSG00000 120802 |
| ZNF611 | Exon Cassette | e9 | up | 4,06 | ENSG00000 213020 |
| LRRFIP1 | ter. Terminal Exo | 21/e23,e24-2 | up (e21) | 4,06 | ENSG00000 124831 |
| DCUN1D4 | Exon Cassette | e3 | up | 4,06 | ENSG00000 109184 |
| SORBS1 | Exon Cassette | e17,e20 | up | 4,05 | ENSG00000 095637 |
| NCAM1 | Complex | 16/e17,e18-2 | up (e17,e18-20) | 4,05 | ENSG00000 149294 |
| NCAM1 | Exon Cassette | e17,e18-20 | up | 4,05 | ENSG00000 149294 |
| TMPO | ter. Terminal Exo | e5/e9-11 | up (e5) | 4,05 | ENSG00000 120802 |
| NRXN1 | Alter. First Exon | 1-3,e5,e8/e | up (e1-3,e5,e8) | 4,05 | ENSG00000 179915 |
| RTN4 | Exon Cassette | e8 | down | 4,04 | ENSG00000 115310 |
| BIN1 | Exon Cassette | e8 | down | 4,04 | ENSG00000 136717 |
| SLC6A2 | Exon Cassette | e6 | up | 4,03 | ENSG00000 103546 |
| DLG1 | ualy Exclusive Ex | e23/e24 | up (e23) | 4,03 | ENSG00000 075711 |
| FUS | Intron Retention | e7 | up | 4,02 | ENSG00000 089280 |
| FLT1 | ter. Terminal Exo | e18,e21-24, | 6-16,e18,e21-24,e2 | 4,01 | ENSG00000 102755 |
| GPR56 | Complex | e6/e7 | up (e7) | 4,01 | ENSG00000 205336 |

| | | | | | |
|---|---|---|---|---|---|
| SLC25A12 | Exon Cassette | e5 | down | 4,01 | ENSG00000 115840 |
| ENPP5 | Exon Cassette | e2 | down | 4,01 | ENSG00000 112796 |
| EPB41L3 | Complex | e22/e24,e25 | down (e24,e25) | 4 | ENSG00000 082397 |
| CLVS1 | Alter. First Exon | e1-5/e6 | down (e1-5) | 4 | ENSG00000 177182 |
| F2 // INS // INS-IG | Alter. First Exon | 1-2,e4-5/e1 | up (e1-2,e4-5) | 3,98 | ENSG00000 129965 // ENSG00000 167244 // ENSG00000 254647 |
| UNC13D | Intron Retention | e11 | up | 3,98 | ENSG00000 092929 |
| ASCC1 | Alter. Donor Site | e4 | down | 3,97 | ENSG00000 138303 |
| ERC1 | Exon Cassette | e6 | up | 3,97 | ENSG00000 082805 |
| GABRB3 | Exon Cassette | e9,e10-11 | up | 3,97 | ENSG00000 166206 |
| HPCAL1 | Alter. First Exon | e2/e3,e5 | up (e3,e5) | 3,97 | ENSG00000 115756 |
| HPCAL1 | Alter. First Exon | e3-5/e4 | up (e3-5) | 3,97 | ENSG00000 115756 |
| HPCAL1 | Alter. First Exon | e3-5/e6 | up (e3-5) | 3,97 | ENSG00000 115756 |
| CLASP1 | Exon Cassette | e22 | up | 3,97 | ENSG00000 074054 |
| TMEM63B | Exon Cassette | e6 | up | 3,97 | ENSG00000 137216 |
| NPNT | Complex | e15 | up | 3,95 | ENSG00000 168743 |
| SLC47A1 | Intron Retention | e18 | up | 3,94 | ENSG00000 142494 |
| ZNF28 | Alter. First Exon | e1-3/e4 | down (e1-3) | 3,94 | ENSG00000 198538 |
| / ARPC4-TTLL3 | ter. Terminal Exo | e12,e14-16, | down (e5-6) | 3,93 | ENSG00000 214021 // ENSG00000 241553 // ENSG00000 250151 |
| HNRNPD | Exon Cassette | e9 | up | 3,92 | ENSG00000 138668 |
| SCARB1 | Exon Cassette | e16 | up | 3,91 | ENSG00000 073060 |
| PDE4D | Exon Cassette | e12-15 | up | 3,91 | ENSG00000 113448 |

| | | | | | |
|---|---|---|---|---|---|
| MRPS28 // TPD5: | Exon Cassette | e9 | down | 3,91 | ENSG00000 076554 // ENSG00000 147586 |
| CLASP2 | Complex | e9 | up | 3,9 | ENSG00000 163539 |
| FAM129A | ter. Terminal Exo | 2/e3-6,e8-1( | up (e3-6,e8-16) | 3,89 | ENSG00000 135842 |
| AKAP13 | Exon Cassette | e14 | up | 3,89 | ENSG00000 170776 |
| VGLL4 | Alter. First Exon | e1-3/e4 | up (e1-3) | 3,89 | ENSG00000 144560 |
| EPB41L3 | Exon Cassette | e23,e25 | down | 3,88 | ENSG00000 082397 |
| HLDB2 // PLCXD | Alter. First Exon | e1-3/e4 | down (e1-3) | 3,88 | ENSG00000 144824 // ENSG00000 240891 |
| LRCH3 | Exon Cassette | e21 | down | 3,88 | ENSG00000 186001 |
| FAM129A | ter. Terminal Exo | e2/e3,e8-12 | up (e3,e8-12) | 3,87 | ENSG00000 135842 |
| CD55 | Alter. Donor Site | e7 | down | 3,86 | ENSG00000 196352 |
| CD55 | Intron Retention | e7 | down | 3,86 | ENSG00000 196352 |
| MFAP2 | Intron Retention | e7 | up | 3,86 | ENSG00000 117122 |
| SLC4A4 | Alter. First Exon | e1/e2 | down (e1) | 3,86 | ENSG00000 080493 |
| HMBOX1 | Intron Retention | e11 | up | 3,86 | ENSG00000 147421 |
| EPB41 | Exon Cassette | e15,e16 | down | 3,84 | ENSG00000 159023 |
| NID2 | Exon Cassette | e13 | down | 3,84 | ENSG00000 087303 |
| AQP3 | Intron Retention | e3 | down | 3,84 | ENSG00000 165272 |
| TMCO1 | Complex | e4,e5 | down | 3,81 | ENSG00000 143183 |
| CARM1 | Exon Cassette | e15 | down | 3,81 | ENSG00000 142453 |
| DMKN | Exon Cassette | e12 | down | 3,81 | ENSG00000 161249 |
| DLG1 | Exon Cassette | e24 | down | 3,81 | ENSG00000 075711 |
| BCLAF1 | Exon Cassette | e11 | up | 3,81 | ENSG00000 029363 |
| KIF26B | Complex | e4,e6-14 | down | 3,79 | ENSG00000 162849 |

| | | | | | |
|---|---|---|---|---|---|
| NIP1 // RAPGEF | Alter. First Exon | 19-24/e25-2 | up (e19-24) | 3,79 | ENSG00000 158987 // ENSG00000 217128 |
| FLNB | Exon Cassette | e27 | up | 3,78 | ENSG00000 136068 |
| SDF4 | Complex | e4,e6-7 | down | 3,77 | ENSG00000 078808 |
| SLAIN1 | Exon Cassette | e5 | down | 3,76 | ENSG00000 139737 |
| AJUBA // HAUS4 | ter. Terminal Exc | -9/e16,e17- | up (e1-9) | 3,76 | ENSG00000 092036 // ENSG00000 129474 |
| DDHD1 | Exon Cassette | e13 | up | 3,76 | ENSG00000 100523 |
| MAATS1 | Exon Cassette | e2 | down | 3,75 | ENSG00000 183833 |
| ELN | Exon Cassette | e32 | up | 3,75 | ENSG00000 049540 |
| FAM129A | ter. Terminal Exc | /e3,e4,e10- | up (e3,e4,e10-13) | 3,74 | ENSG00000 135842 |
| CD9 | Complex | e2-4 | up | 3,74 | ENSG00000 010278 |
| MPZL1 | Exon Cassette | e5 | up | 3,71 | ENSG00000 197965 |
| GTSF1 | Alter. First Exon | e1/e3 | up (e1) | 3,71 | ENSG00000 170627 |
| GTSF1 | Alter. First Exon | e1/e2 | up (e1) | 3,71 | ENSG00000 170627 |
| CLASP1 | Exon Cassette | e25 | down | 3,71 | ENSG00000 074054 |
| PRPF40A | Exon Cassette | e9 | up | 3,71 | ENSG00000 196504 |
| E1 // NFS1 // RB | Intron Retention | e33 | up | 3,71 | ENSG00000 214078 // ENSG00000 244005 // ENSG00000 244462 |
| NIP1 // RAPGEF | Alter. First Exon | ,e20-24/e25 | up (e1-2,e20-24) | 3,7 | ENSG00000 158987 // ENSG00000 217128 |
| NIN | Exon Cassette | e19 | up | 3,68 | ENSG00000 100503 |
| FOXP2 | Exon Cassette | e18,e19 | down | 3,68 | ENSG00000 128573 |
| ETV4 | Alter. First Exon | e1,e3-4/e5 | up (e1,e3-4) | 3,67 | ENSG00000 175832 |
| SGSM1 | Exon Cassette | e13 | up | 3,67 | ENSG00000 167037 |

| | | | | | |
|---|---|---|---|---|---|
| MGST1 | Alter. First Exon | e1/e2 | down (e2) | 3,66 | ENSG00000008394 |
| PRMT1 | Alter. Donor Site | e9 | up | 3,66 | ENSG00000126457 |
| VRK2 | Complex | e8,e9 | down | 3,66 | ENSG00000028116 |
| SCLT1 | Exon Cassette | e8-10,e12 | down | 3,66 | ENSG00000151466 |
| --- | Alter. Donor Site | e1 | down | 3,66 | --- |
| FAM132B | Alter. First Exon | e1-2/e3 | down (e1-2) | 3,65 | ENSG00000178752 |
| MRPS28 // TPD5 | Exon Cassette | e8,e9 | down | 3,65 | ENSG00000076554 // ENSG00000147586 |
| S100A13 | Alter. First Exon | e1-3/e9 | down (e1-3) | 3,64 | ENSG00000189171 |
| FLT1 | Alter. First Exon | e26-27,e29 | (e19-24,e26-27,e2 | 3,64 | ENSG00000102755 |
| NOL12 // TRIOBP | Exon Cassette | e15 | up | 3,64 | ENSG00000100106 // ENSG00000256872 |
| AHRR // PDCD6 | Alter. First Exon | e8-12/e13 | down (e8-12) | 3,64 | ENSG00000063438 // ENSG00000249915 |
| SGIP1 | Exon Cassette | e16-17 | up | 3,63 | ENSG00000118473 |
| CDH23 | Alter. First Exon | 11,e13-27/e | up (e1-11,e13-27) | 3,63 | ENSG00000107736 |
| APOPT1 // KLC1 | Complex | e18,e21 | up | 3,63 | ENSG00000126214 // ENSG00000256053 |
| 2AP // IGHD // IG | Alter. First Exon | e35-38/e39 | up (e35-38) | 3,63 | ENSG00000211896 // ENSG00000211898 // ENSG00000213140 |
| B3GALNT1 | Exon Cassette | e5,e6-7 | up | 3,63 | ENSG00000169255 |
| MDM2 | Alter. First Exon | e1,e4/e2 | down (e2) | 3,62 | ENSG00000135679 |
| DLG4 | Alter. First Exon | e1-3,e5/e6 | up (e1-3,e5) | 3,62 | ENSG00000132535 |
| ARHGAP21 | Alter. First Exon | e9/e10 | down (e9) | 3,61 | ENSG00000107863 |
| SYNRG | Exon Cassette | e7,e8 | up | 3,61 | ENSG00000006114 |

| | | | | | |
|---|---|---|---|---|---|
| SORBS1 | Complex | e11,e12 | up | 3,6 | ENSG00000 095637 |
| // LY75 // LY75- | ter. Terminal Exo | e13/e14-35 | down (e14-35) | 3,6 | ENSG00000 054219 // ENSG00000 241399 // ENSG00000 248672 |
| PRAME | Alter. First Exon | e1/e2 | up (e1) | 3,6 | ENSG00000 185686 |
| TPM1 | Exon Cassette | e3 | up | 3,59 | ENSG00000 140416 |
| NRXN1 | Alter. First Exon | 1-2,e5,e8/e | up (e1-2,e5,e8) | 3,59 | ENSG00000 179915 |
| NR2F1-AS1 | Exon Cassette | e7 | up | 3,59 | ENSG00000 237187 |
| TPD52L1 | ter. Terminal Exo | e7/e8-9 | down (e8-9) | 3,59 | ENSG00000 111907 |
| SLC12A6 | Alter. First Exon | e2-3/e4 | down (e2-3) | 3,58 | ENSG00000 140199 |
| SLC12A6 | Alter. First Exon | e3/e4 | down (e3) | 3,58 | ENSG00000 140199 |
| PTPRK | Exon Cassette | e2 | down | 3,58 | ENSG00000 152894 |
| SORBS1 | lter. Acceptor Sit | e12 | up | 3,57 | ENSG00000 095637 |
| FOXP2 | ter. Terminal Exo | 7/e18,e19-2 | down (e18,e19-24) | 3,57 | ENSG00000 128573 |
| MAP2 | Exon Cassette | e16 | up | 3,56 | ENSG00000 078018 |
| SORBS2 | Complex | e6-8/e9 | down (e6-8) | 3,56 | ENSG00000 154556 |
| SH3PXD2A | Alter. First Exon | e5-6,e9,e12 | wn (e1-3,e5-6,e9,e1 | 3,55 | ENSG00000 107957 |
| MYO18A // TIAF1 | Exon Cassette | e48 | up | 3,55 | ENSG00000 196535 // ENSG00000 221995 |
| ZNF160 | ter. Terminal Exo | e4/e5-6,e8 | down (e5-6,e8) | 3,55 | ENSG00000 170949 |
| ARHGAP21 | Exon Cassette | e9 | down | 3,54 | ENSG00000 107863 |
| PIEZO1 | Complex | e22-37 | down | 3,53 | ENSG00000 103335 |
| ASIC4 | Intron Retention | e4 | up | 3,53 | ENSG00000 072182 |
| SLC12A6 | Alter. First Exon | e1-3/e4 | down (e1-3) | 3,52 | ENSG00000 140199 |
| ABAT | Exon Cassette | e5 | down | 3,52 | ENSG00000 183044 |
| CLASP2 | Alter. First Exon | e1-7/e9 | up (e1-7) | 3,52 | ENSG00000 163539 |

| | | | | | |
|---|---|---|---|---|---|
| LIMCH1 | Exon Cassette | e13,e14-19 | up | 3,52 | ENSG00000064042 |
| KCNMA1 | ter. Terminal Exc | e29-30,e32 | -23,e25,e29-30,e32 | 3,51 | ENSG00000156113 |
| ZNF542 | Complex | e1-2 | up | 3,51 | ENSG00000240225 |
| SRSF11 | Exon Cassette | e7 | up | 3,5 | ENSG00000116754 |
| IGDCC4 | Alter. First Exon | e1-18/e19 | up (e1-18) | 3,5 | ENSG00000103742 |
| GPR56 | Complex | e3,e5 | up | 3,5 | ENSG00000205336 |
| SLC47A1 | Alter. First Exon | e2-8/e9 | up (e2-8) | 3,5 | ENSG00000142494 |
| SLC47A1 | Complex | e8-9/e10 | up (e8-9) | 3,5 | ENSG00000142494 |
| DMKN | Exon Cassette | e9,e11-12 | down | 3,5 | ENSG00000161249 |
| CNTN4 | Alter. First Exon | e1-3,e6-8/e9 | up (e1-3,e6-8) | 3,5 | ENSG00000144619 |
| PRSS16 | Exon Cassette | e4-8 | down | 3,49 | ENSG00000112812 |
| PAXIP1-AS2 | ter. Terminal Exc | e3/e4-8 | down (e4-8) | 3,49 | ENSG00000214106 |
| OXR1 | Exon Cassette | e4,e5-11 | up | 3,49 | ENSG00000164830 |
| GRIN1 | ter. Terminal Exc | e19/e21 | up (e21) | 3,49 | ENSG00000176884 |
| ZNF610 | Exon Cassette | e6 | down | 3,48 | ENSG00000167554 |
| FAM13A | Alter. First Exon | 2-4,e6-9/e1( | up (e2-4,e6-9) | 3,48 | ENSG00000138640 |
| KCNMA1 | ter. Terminal Exc | 5,e29-30,e3 | 10-23,e25,e29-30,e | 3,46 | ENSG00000156113 |
| MRPL42 | Exon Cassette | e3 | up | 3,45 | ENSG00000198015 |
| SLC12A6 | Alter. First Exon | e1,e3/e4 | down (e1,e3) | 3,45 | ENSG00000140199 |
| ATCAY | Complex | e4,e6 | up | 3,45 | ENSG00000167654 |
| ANXA2 | Alter. First Exon | e1-2/e4 | down (e1-2) | 3,44 | ENSG00000182718 |
| FOXP2 | ter. Terminal Exc | e17/e19-25 | down (e19-25) | 3,43 | ENSG00000128573 |
| NFIB | Alter. First Exon | e1,e3/e4 | down (e1,e3) | 3,43 | ENSG00000147862 |
| PLEKHA6 | ıaly Exclusive Ex | e10/e13 | up (e10) | 3,42 | ENSG00000143850 |
| KCNMA1 | ter. Terminal Exc | 3,e25,e29-3 | 5,e10-23,e25,e29-3 | 3,42 | ENSG00000156113 |
| HPSE | Exon Cassette | e9 | down | 3,42 | ENSG00000173083 |

| | | | | | |
|---|---|---|---|---|---|
| EYA1 | lter. Acceptor Sit | e6 | up | 3,42 | ENSG00000 104313 |
| KCNMA1 | ter. Terminal Exc | 3,e25,e29-3 | e5,e10-23,e25,e29-3 | 3,4 | ENSG00000 156113 |
| KCNMA1 | ter. Terminal Exc | e29-30,e32 | -23,e25,e29-30,e32 | 3,4 | ENSG00000 156113 |
| RIC3 | Exon Cassette | e3,e4-5 | down | 3,4 | ENSG00000 166405 |
| DST | Alter. First Exon | e1-3/e5 | up (e1-3) | 3,4 | ENSG00000 151914 |
| TCF4 | Alter. First Exon | e11-12,e15, | 2,e7,e9,e11-12,e15, | 3,39 | ENSG00000 196628 |
| TNIK | Exon Cassette | e15 | down | 3,39 | ENSG00000 154310 |
| EYA1 | Complex | e6 | up | 3,39 | ENSG00000 104313 |
| KIF1B | ter. Terminal Exc | 26/e27-52,e5 | up (e26) | 3,38 | ENSG00000 054523 |
| ZNF611 | Exon Cassette | e8,e9 | up | 3,38 | ENSG00000 213020 |
| PFN2 | Complex | e3 | down | 3,38 | ENSG00000 070087 |
| FOXP2 | ter. Terminal Exc | 7/e20,e21-2 | down (e20,e21-25) | 3,38 | ENSG00000 128573 |
| PLEC | Alter. First Exon | e4/e6 | down (e4) | 3,38 | ENSG00000 178209 |
| KCNMA1 | ter. Terminal Exc | -26,e29-30, | 0-23,e25-26,e29-30 | 3,37 | ENSG00000 156113 |
| ABAT | Alter. First Exon | e1/e2 | up (e1) | 3,37 | ENSG00000 183044 |
| KCNH7 | ter. Terminal Exc | 9/e10,e11-1 | up (e10,e11-19) | 3,37 | ENSG00000 184611 |
| TFAP2A | Alter. First Exon | e1/e3 | up (e3) | 3,37 | ENSG00000 137203 |
| KCNH2 | Alter. First Exon | 1-5,e7-16/e | down (e1-5,e7-16) | 3,37 | ENSG00000 055118 |
| KCNH2 | Alter. First Exon | 3-5,e7-16/e | down (e3-5,e7-16) | 3,37 | ENSG00000 055118 |
| --- | Exon Cassette | e22 | down | 3,37 | --- |
| KIF1B | ter. Terminal Exc | 26/e27,e28-5 | up (e26) | 3,36 | ENSG00000 054523 |
| KIF1B | ter. Terminal Exc | 26/e27,e28-5 | up (e26) | 3,35 | ENSG00000 054523 |
| LPHN2 | Exon Cassette | e25,e26 | down | 3,35 | ENSG00000 117114 |
| RGMA | Alter. First Exon | e1/e3-6 | down (e3-6) | 3,35 | ENSG00000 182175 |
| TCF4 | Alter. First Exon | e16-20/e21 | up (e16-20) | 3,35 | ENSG00000 196628 |
| TCF4 | Alter. First Exon | e17-20/e21 | up (e17-20) | 3,35 | ENSG00000 196628 |

| | | | | | |
|---|---|---|---|---|---|
| ZNF542 | Alter. First Exon | e1/e2 | up (e2) | 3,35 | ENSG00000240225 |
| PPP1R16B | ter. Terminal Exo | e2/e3-12 | down (e3-12) | 3,35 | ENSG00000101445 |
| ZFP64 | Alter. First Exon | e1-5/e7 | down (e1-5) | 3,35 | ENSG00000020256 |
| FYN | Exon Cassette | e12 | up | 3,35 | ENSG00000010810 |
| TCF4 | Alter. First Exon | 11-12,e15,e | -7,e9,e11-12,e15,e | 3,34 | ENSG00000196628 |
| TCF4 | Alter. First Exon | -12,e15,e1 | 5-9,e11-12,e15,e1 | 3,34 | ENSG00000196628 |
| ELL2 | Complex | e1-8 | up | 3,34 | ENSG00000118985 |
| TPD52L1 | ter. Terminal Exo | e7/e9 | down (e9) | 3,34 | ENSG00000111907 |
| TCF4 | Alter. First Exon | 11-12,e15,e | -7,e9,e11-12,e15,e | 3,32 | ENSG00000196628 |
| TCF4 | Alter. First Exon | e18-20/e21 | up (e18-20) | 3,32 | ENSG00000196628 |
| DMKN | Exon Cassette | e9,e12 | down | 3,32 | ENSG00000161249 |
| CDCA7 | Exon Cassette | e3 | down | 3,32 | ENSG00000144354 |
| WDR35 | Exon Cassette | e11,e12 | down | 3,32 | ENSG00000118965 |
| PPP1R16B | ter. Terminal Exo | 2/e3-7,e9-1 | down (e3-7,e9-12) | 3,32 | ENSG00000101445 |
| NEO1 | Exon Cassette | e21 | down | 3,31 | ENSG00000067141 |
| SLC47A1 | Complex | ,e8-11,e14- | down | 3,31 | ENSG00000142494 |
| TCF4 | Alter. First Exon | -12,e15,e1 | 8-9,e11-12,e15,e1 | 3,31 | ENSG00000196628 |
| FAM221A | Exon Cassette | e6 | up | 3,31 | ENSG00000188732 |
| FLT1 | Alter. First Exon | 24,e26-27,e | 19,e21-24,e26-27,e | 3,3 | ENSG00000102755 |
| FAM135A | Exon Cassette | e11 | up | 3,3 | ENSG00000082269 |
| DHRS2 | Alter. First Exon | e1-5/e6 | down (e1-5) | 3,29 | ENSG00000100867 |
| TCF4 | Alter. First Exon | e15,e19-20 | up (e13,e15,e19-20 | 3,29 | ENSG00000196628 |
| PCSK5 | ter. Terminal Exo | 7,e8-10,e1 | wn (e7,e8-10,e12-2 | 3,29 | ENSG00000099139 |
| RPS6KA6 | Exon Cassette | e2,e3 | up | 3,29 | ENSG00000072133 |
| EPB41 | Exon Cassette | e16 | down | 3,28 | ENSG00000159023 |
| SHC1 | Alter. First Exon | e2/e3 | up (e2) | 3,28 | ENSG00000160691 |

| | | | | | |
|---|---|---|---|---|---|
| DBI | Complex | e1/e3 | up (e3) | 3,28 | ENSG00000 155368 |
| DBI | Complex | e1,e3 | up | 3,28 | ENSG00000 155368 |
| DBI | Exon Cassette | e3 | up | 3,28 | ENSG00000 155368 |
| HAUS3 // POLN | ter. Terminal Exc | e22/e23-32 | down (e23-32) | 3,28 | ENSG00000 130997 // ENSG00000 214367 |
| SMPDL3A | Exon Cassette | e2 | up | 3,28 | ENSG00000 172594 |
| MAPT | Exon Cassette | e3,e4 | up | 3,27 | ENSG00000 186868 |
| TCF4 | Alter. First Exon | 2,e15,e19-2 | (e10-12,e15,e19-2 | 3,27 | ENSG00000 196628 |
| --- | Exon Cassette | e2-5 | up | 3,27 | --- |
| GATA2 | Intron Retention | e6 | up | 3,27 | ENSG00000 179348 |
| CCND1 | Complex | e1-3 | down | 3,26 | ENSG00000 110092 |
| LRRC49 | Exon Cassette | e7 | up | 3,26 | ENSG00000 137821 |
| ACADVL | ter. Terminal Exc | 1/e12-13,e2 | down (e12-13,e22) | 3,26 | ENSG00000 072778 |
| SLC47A1 | Exon Cassette | e8 | up | 3,26 | ENSG00000 142494 |
| FAM13A | Alter. First Exon | e1-9/e10 | up (e1-9) | 3,26 | ENSG00000 138640 |
| RAC1 | Exon Cassette | e5 | up | 3,26 | ENSG00000 136238 |
| RIC3 | Complex | e3-5,e7 | down | 3,25 | ENSG00000 166405 |
| MAPT | Exon Cassette | e3 | up | 3,25 | ENSG00000 186868 |
| EPB41L3 | Exon Cassette | e24,e25 | down | 3,24 | ENSG00000 082397 |
| SUPT7L | Intron Retention | e2 | up | 3,24 | ENSG00000 119760 |
| PMS2CL | ter. Terminal Exc | e4/e5-15 | down (e5-15) | 3,23 | ENSG00000 187953 |
| WLS | Exon Cassette | e2 | up | 3,22 | ENSG00000 116729 |
| REST | ter. Terminal Exc | -8,e11/e14- | up (e5-8,e11) | 3,22 | ENSG00000 084093 |
| REST | ter. Terminal Exc | 8,e11/e13,e | up (e5-8,e11) | 3,22 | ENSG00000 084093 |
| MID1 | Alter. First Exon | e1,e3/e5 | up (e1,e3) | 3,22 | ENSG00000 101871 |
| PLEKHA6 | Exon Cassette | e13 | down | 3,21 | ENSG00000 143850 |

| | | | | | |
|---|---|---|---|---|---|
| NCAM1 | Complex | l6/e17,e18-1 | up (e17,e18-19) | 3,21 | ENSG00000 149294 |
| NCAM1 | Exon Cassette | e17,e18-19 | up | 3,21 | ENSG00000 149294 |
| CD9 | Complex | e2/e3-6 | up (e3-6) | 3,21 | ENSG00000 010278 |
| LTA4H | Complex | e3-13,e15-18 | down | 3,21 | ENSG00000 111144 |
| DHRS2 | lter. Acceptor Sit | e6 | up | 3,21 | ENSG00000 100867 |
| SLCO3A1 | Alter. First Exon | e1,e4/e5 | up (e1,e4) | 3,21 | ENSG00000 176463 |
| KIAA0430 | Intron Retention | e3 | up | 3,21 | ENSG00000 166783 |
| TCF4 | Alter. First Exon | l-12,e15,e19 | 4-9,e11-12,e15,e19 | 3,21 | ENSG00000 196628 |
| HAUS3 // POLN | ter. Terminal Exo | 22/e23,e24-3 | down (e23,e24-32) | 3,21 | ENSG00000 130997 // ENSG00000 214367 |
| TCF4 | Alter. First Exon | -12,e15,e19 | ,e9,e11-12,e15,e19 | 3,2 | ENSG00000 196628 |
| TJAP1 | Complex | e10,e11 | down | 3,2 | ENSG00000 137221 |
| GPR56 | Exon Cassette | e5 | up | 3,19 | ENSG00000 205336 |
| MAST4 | Exon Cassette | e29 | down | 3,19 | ENSG00000 069020 |
| KIAA1324L | Alter. First Exon | e1/e2 | up (e1) | 3,19 | ENSG00000 164659 |
| ZMIZ1 | Exon Cassette | e12 | up | 3,18 | ENSG00000 108175 |
| SLC2A14 | Alter. First Exon | e2/e3,e8-14 | down (e3,e8-14) | 3,18 | ENSG00000 173262 |
| KCNH7 | ter. Terminal Exo | 9/e10,e11-1 | up (e10,e11-12) | 3,18 | ENSG00000 184611 |
| GBA // GBAP1 | Alter. First Exon | e1/e15 | up (e1) | 3,17 | ENSG00000 160766 // ENSG00000 177628 |
| F6 // HEXA // PA | Intron Retention | e35,e36 | up | 3,17 | ENSG00000 137817 // ENSG00000 140488 // ENSG00000 213614 |
| DLL1 | ter. Terminal Exo | e4/e5-11 | up (e5-11) | 3,17 | ENSG00000 198719 |
| DDHD1 | ter. Terminal Exo | e12/e14 | up (e12) | 3,16 | ENSG00000 100523 |
| VGLL4 | Alter. First Exon | e1-2/e4 | up (e1-2) | 3,16 | ENSG00000 144560 |

| | | | | | |
|---|---|---|---|---|---|
| DALRD3 | Complex | e2,e3 | down | 3,16 | ENSG00000178149 |
| TFAP2A | Exon Cassette | e8 | up | 3,16 | ENSG00000137203 |
| RIC3 | Complex | e3,e4-5,e7 | down | 3,15 | ENSG00000166405 |
| TVP23A | lter. Acceptor Sit | e4 | up | 3,14 | ENSG00000166676 |
| IL17RC | Complex | e4-5 | up | 3,14 | ENSG00000163702 |
| LPHN2 | Alter. First Exon | e4-6/e5 | down (e4-6) | 3,13 | ENSG00000117114 |
| F6 // HEXA // PA | Exon Cassette | e21 | down | 3,13 | ENSG00000137817 // ENSG00000140488 // ENSG00000213614 |
| RAPGEF4 | Alter. First Exon | e1/e3 | down (e1) | 3,13 | ENSG00000091428 |
| RAPGEF4 | Alter. First Exon | e1/e2 | down (e1) | 3,13 | ENSG00000091428 |
| HLA-DPB1 | ter. Terminal Exo | e2-4/e5-8 | down (e2-4) | 3,13 | ENSG00000223865 |
| KCNT2 | Complex | e20,e22 | down | 3,12 | ENSG00000162687 |
| NUDT7 | Exon Cassette | e5 | up | 3,12 | ENSG00000140876 |
| SLC4A4 | Alter. First Exon | e1-5/e4 | down (e1-5) | 3,12 | ENSG00000080493 |
| DDX41 | Intron Retention | e5 | up | 3,12 | ENSG00000183258 |
| PPHLN1 | Exon Cassette | e8 | up | 3,1 | ENSG00000134283 |
| KIF1B | Alter. First Exon | -16,e18-25,e | 11,e14-16,e18-25,e | 3,09 | ENSG00000054523 |
| TCF7L2 | Alter. First Exon | e1-3/e8 | down (e1-3) | 3,09 | ENSG00000148737 |
| TCF7L2 | Alter. First Exon | e1-3/e9 | down (e1-3) | 3,09 | ENSG00000148737 |
| IYMAI // PLAGL1 | Complex | e1,e4 | up | 3,09 | ENSG00000118495 |
| FLT1 | ter. Terminal Exo | e4-11,e13,e | own (e4-11,e13,e15 | 3,08 | ENSG00000102755 |
| NOL12 // TRIOBF | Alter. First Exon | e8-18/e20 | down (e8-18) | 3,08 | ENSG00000100106 // ENSG00000256872 |
| ATP11C | Exon Cassette | e30 | down | 3,08 | ENSG00000101974 |
| HLA-DMA | Complex | e1-2/e2 | up (e2) | 3,07 | ENSG00000204257 |

| | | | | | |
|---|---|---|---|---|---|
| NOL8 | Exon Cassette | e4 | up | 3,07 | ENSG00000198000 |
| SLC47A1 | Exon Cassette | e7,e8-15 | down | 3,06 | ENSG00000142494 |
| DMKN | Exon Cassette | e10 | down | 3,06 | ENSG00000161249 |
| CSTF2 | Exon Cassette | e9 | down | 3,06 | ENSG00000101811 |
| SLC47A1 | Exon Cassette | e12,e13 | up | 3,05 | ENSG00000142494 |
| ABI3BP | Exon Cassette | e32 | down | 3,05 | ENSG00000154175 |
| DLL1 | ter. Terminal Exo | e4/e5-10 | up (e5-10) | 3,05 | ENSG00000198719 |
| PDE4B | Alter. First Exon | e5-9/e10 | down (e5-9) | 3,04 | ENSG00000184588 |
| C11orf80 // RCE1 | Exon Cassette | e2,e3 | up | 3,04 | ENSG00000173653 // ENSG00000173715 |
| PFKM | Intron Retention | e15 | up | 3,04 | ENSG00000152556 |
| ANK2 | Alter. First Exon | e1,e4/e2 | up (e1,e4) | 3,04 | ENSG00000145362 |
| ITGA5 | Exon Cassette | e3 | up | 3,03 | ENSG00000161638 |
| TAOK3 | Alter. First Exon | -8,e10-15/e | up (e2-8,e10-15) | 3,03 | ENSG00000135090 |
| ZNF83 | Exon Cassette | e13 | up | 3,03 | ENSG00000167766 |
| REST | ter. Terminal Exo | 3,e11-12/e14 | up (e5-8,e11-12) | 3,03 | ENSG00000084093 |
| REST | ter. Terminal Exo | ,e11-12/e13 | up (e5-8,e11-12) | 3,03 | ENSG00000084093 |
| SCLT1 | Exon Cassette | 0,e12-17,e1 | down | 3,03 | ENSG00000151466 |
| STX1A | Complex | e8,e9 | up | 3,03 | ENSG00000106089 |
| ZNF438 | Exon Cassette | e9 | up | 3,02 | ENSG00000183621 |
| / FXYD6 // FXYD | Exon Cassette | e4 | up | 3,02 | ENSG00000137726 // ENSG00000137731 // ENSG00000255245 |
| PIH1D1 | lter. Acceptor Sit | e2 | up | 3,02 | ENSG00000104872 |
| STK32C | Alter. First Exon | e4/e5 | up (e5) | 3,01 | ENSG00000165752 |
| CCNJL | Exon Cassette | e7 | up | 3,01 | ENSG00000135083 |

| | | | | | |
|---|---|---|---|---|---|
| MID1 | Alter. First Exon | e1,e3/e2 | up (e1,e3) | 3,01 | ENSG00000101871 |
| PROX1 | Complex | e1 | up | 3 | ENSG00000117707 |
| FAM89B | Complex | e2 | down | 3 | ENSG00000176973 |
| CEP68 | Alter. First Exon | e1/e3 | up (e1) | 3 | ENSG00000011523 |
| MLH1 | Exon Cassette | e11-12 | down | 3 | ENSG00000076242 |
| JARID2 | Alter. First Exon | e1-4/e5 | up (e1-4) | 3 | ENSG00000008083 |
| XRCC3 | Exon Cassette | e2 | up | 2,99 | ENSG00000126215 |
| EXOC7 | Exon Cassette | e7 | down | 2,99 | ENSG00000182473 |
| LAMA1 | Complex | e42-56 | up | 2,99 | ENSG00000101680 |
| // PRR5 // PRR5 | Alter. First Exon | e12/e13 | down (e12) | 2,99 | ENSG00000186654 // ENSG00000241484 // ENSG00000248405 |
| JARID2 | Alter. First Exon | e2-4/e5 | up (e2-4) | 2,99 | ENSG00000008083 |
| JARID2 | Alter. First Exon | e3-4/e5 | up (e3-4) | 2,99 | ENSG00000008083 |
| CAV1 | Alter. Donor Site | e1 | down | 2,99 | ENSG00000105974 |
| TAOK3 | Alter. First Exon | -8,e10-15/e | up (e1-8,e10-15) | 2,98 | ENSG00000135090 |
| PLCD4 | Complex | e6,e7-10 | up | 2,98 | ENSG00000115556 |
| SOS1 | Exon Cassette | e24 | up | 2,98 | ENSG00000115904 |
| FAM13A | Alter. First Exon | 4,e6-9,e11/ | up (e2-4,e6-9,e11) | 2,98 | ENSG00000138640 |
| TRIP6 | Exon Cassette | e2 | down | 2,98 | ENSG00000087077 |
| PCSK5 | ter. Terminal Exo | e6/e7-22 | down (e7-22) | 2,98 | ENSG00000099139 |
| SLC37A4 | Alter. First Exon | e1-2/e3 | down (e1-2) | 2,97 | --- |
| FXYD5 | Alter. First Exon | e1-5/e6 | up (e1-5) | 2,97 | ENSG00000089327 |
| ATF2 | Exon Cassette | e14 | up | 2,97 | ENSG00000115966 |
| CNTN4 | Alter. First Exon | e1-3,e6/e4-5 | up (e1-3,e6) | 2,97 | ENSG00000144619 |
| SLC1A3 | Complex | e2,e4 | up | 2,97 | ENSG00000079215 |

| | | | | | |
|---|---|---|---|---|---|
| MYO10 | Complex | e1,e3/e4 | up (e1,e3) | 2,97 | ENSG00000145555 |
| ETV1 | Exon Cassette | e5 | up | 2,97 | ENSG00000006468 |
| SPAG6 | Exon Cassette | e3,e4-5 | up | 2,96 | ENSG00000077327 |
| DUFB8 // SEC31 | Exon Cassette | e23 | down | 2,96 | ENSG00000075826 // ENSG00000166136 |
| REST | Complex | e5,e8,e11 | up | 2,96 | ENSG00000084093 |
| DDX41 | Intron Retention | e6 | up | 2,96 | ENSG00000183258 |
| KRBOX4 | Complex | e6/e7 | up (e6) | 2,96 | ENSG00000147121 |
| NSUN4 | Alter. First Exon | e1/e2 | down (e1) | 2,95 | ENSG00000117481 |
| RTN3 | Exon Cassette | e4 | down | 2,95 | ENSG00000133318 |
| RTN1 | Exon Cassette | e5 | up | 2,95 | ENSG00000139970 |
| HSP90AA1 | Alter. First Exon | e1-2/e4 | down (e1-2) | 2,95 | ENSG00000080824 |
| SULT1C4 | ter. Terminal Exo | e3/e4-7 | down (e4-7) | 2,95 | ENSG00000198075 |
| E1 // NFS1 // RB | Intron Retention | e32 | up | 2,95 | ENSG00000214078 // ENSG00000244005 // ENSG00000244462 |
| CD59 | Alter. First Exon | e1-3/e6 | down (e1-3) | 2,94 | ENSG00000085063 |
| CD59 | Alter. First Exon | e1,e3/e5 | down (e1,e3) | 2,94 | ENSG00000085063 |
| FLT1 | ter. Terminal Exo | 4-11,e13,e1 | wn (e4-11,e13,e15-1 | 2,94 | ENSG00000102755 |
| RNF212 | Exon Cassette | e8-10 | up | 2,94 | ENSG00000178222 |
| S5 // EEF1E1 // T | Alter. First Exon | -7,e9,e11/e | down (e6-7,e9,e11) | 2,94 | ENSG00000124802 // ENSG00000188428 // ENSG00000239264 |
| NSMF | Complex | e4/e5 | up (e5) | 2,94 | ENSG00000165802 |
| MADD | Exon Cassette | e22 | down | 2,93 | ENSG00000110514 |
| ANXA2 | Exon Cassette | e2 | down | 2,93 | ENSG00000182718 |

| | | | | | |
|---|---|---|---|---|---|
| HADHA | ter. Terminal Exo | e10/e11-20 | up (e10) | 2,93 | ENSG00000084754 |
| VGLL4 | Alter. First Exon | e1/e4 | up (e1) | 2,93 | ENSG00000144560 |
| UBXN1 | Intron Retention | e6 | up | 2,92 | ENSG00000162191 |
| MPP3 | Intron Retention | e18 | up | 2,92 | ENSG00000161647 |
| RIMS1 | Exon Cassette | e32,e33-34 | down | 2,92 | ENSG00000079841 |
| MAP7 | Alter. First Exon | e2-3/e4 | down (e2-3) | 2,92 | ENSG00000135525 |
| PITRM1 | Intron Retention | e26 | down | 2,91 | ENSG00000107959 |
| GLI2 | Complex | e14,e15 | down | 2,91 | ENSG00000074047 |
| P4HA2 | Alter. First Exon | e1/e2 | down (e1) | 2,91 | ENSG00000072682 |
| CTNNBIP1 | Alter. First Exon | e1-2,e4/e5 | up (e1-2,e4) | 2,9 | ENSG00000178585 |
| ASTN1 | lter. Acceptor Sit | e7 | up | 2,9 | ENSG00000152092 |
| --- | Exon Cassette | e2 | up | 2,9 | --- |
| CD9 | Complex | e5-6 | down | 2,9 | ENSG00000010278 |
| SLC47A1 | Exon Cassette | e7,e8-16 | down | 2,9 | ENSG00000142494 |
| C17orf104 | ter. Terminal Exo | 0/e11,e12-1 | up (e11,e12-16) | 2,9 | ENSG00000180336 |
| ZNF83 | Exon Cassette | e10,e13 | up | 2,9 | ENSG00000167766 |
| SLC25A17 | Exon Cassette | e7 | up | 2,9 | ENSG00000100372 |
| PNPLA8 | Exon Cassette | e3-4 | up | 2,9 | ENSG00000135241 |
| VGLL4 | Alter. First Exon | e4/e5 | down (e4) | 2,89 | ENSG00000144560 |
| HAND2-AS1 | Exon Cassette | e3 | down | 2,89 | ENSG00000237125 |
| BTN2A1 | Alter. First Exon | e1-7,e10/e9 | up (e1-7,e10) | 2,89 | ENSG00000112763 |
| NLGN3 | Exon Cassette | e3 | up | 2,89 | ENSG00000196338 |
| --- | Complex | e5-7 | down | 2,89 | --- |
| ALDH1L2 | lter. Acceptor Sit | e13 | down | 2,88 | ENSG00000136010 |
| 2AP // IGHD // IG | Alter. First Exon | -16/e24-25,e | down (e15-16) | 2,88 | ENSG00000211896 // ENSG00000211898 // ENSG00000213140 |

| | | | | | |
|---|---|---|---|---|---|
| 2AP // IGHD // IG | Alter. First Exon | -16,e101/e2 | down (e15-16,e101) | 2,88 | ENSG00000211896 // ENSG00000211898 // ENSG00000213140 |
| 2AP // IGHD // IG | Alter. First Exon | 5-16/e17,e1 | down (e15-16) | 2,88 | ENSG00000211896 // ENSG00000211898 // ENSG00000213140 |
| 2AP // IGHD // IG | Alter. First Exon | 5-16,e101/e | down (e15-16,e101) | 2,88 | ENSG00000211896 // ENSG00000211898 // ENSG00000213140 |
| 2AP // IGHD // IG | Alter. First Exon | )-40,e101/e | down (e39-40,e101) | 2,88 | ENSG00000211896 // ENSG00000211898 // ENSG00000213140 |
| NRXN1 | Alter. First Exon | 8-19,e21,e2 | e1-3,e5,e8-19,e21,e | 2,88 | ENSG00000179915 |
| ADD1 | Alter. Donor Site | e12 | down | 2,88 | ENSG00000087274 |
| TMEM63A | ter. Terminal Exo | 21-27/e22-2 | down (e22-26) | 2,87 | ENSG00000196187 |
| CADM1 | Alter. First Exon | e10/e12-13 | down (e10) | 2,87 | ENSG00000182985 |
| 2AP // IGHD // IG | Alter. First Exon | e39-40/e103 | down (e39-40) | 2,87 | ENSG00000211896 // ENSG00000211898 // ENSG00000213140 |
| ZNF160 | ter. Terminal Exo | e4/e5,e6-7 | down (e5,e6-7) | 2,87 | ENSG00000170949 |
| ARHGEF7 | Alter. First Exon | e1/e3 | down (e1) | 2,86 | ENSG00000102606 |
| VRK2 | Complex | e12,e13 | down | 2,86 | ENSG00000028116 |
| BTN2A1 | Alter. First Exon | e1-7/e9 | up (e1-7) | 2,86 | ENSG00000112763 |
| FDFT1 | Alter. First Exon | e1/e2 | up (e1) | 2,86 | ENSG00000079459 |
| KLHL13 | Alter. First Exon | e1-2/e6 | up (e1-2) | 2,86 | ENSG00000003096 |
| ACTN2 | Exon Cassette | e9,e10 | down | 2,85 | ENSG00000077522 |

| | | | | | |
|---|---|---|---|---|---|
| 2AP // IGHD // IG | Alter. First Exon | 16/e26,e27, | down (e15-16) | 2,85 | ENSG00000211896 // ENSG00000211898 // ENSG00000213140 |
| 2AP // IGHD // IG | Alter. First Exon | 16,e101/e2 | down (e15-16,e101) | 2,85 | ENSG00000211896 // ENSG00000211898 // ENSG00000213140 |
| DNMT3A | Alter. First Exon | e1/e2 | up (e1) | 2,85 | ENSG00000119772 |
| KRD18A // FAM9 | ter. Terminal Exo | 16,e17-18,e | n (e16,e17-18,e20- | 2,85 | ENSG00000180071 |
| LPAR1 | Alter. First Exon | e2/e6 | up (e2) | 2,85 | ENSG00000198121 |
| SPATS2 | Exon Cassette | e13 | up | 2,84 | ENSG00000123352 |
| 2AP // IGHD // IG | Alter. First Exon | 7,e103/e39- | up (e17,e103) | 2,84 | ENSG00000211896 // ENSG00000211898 // ENSG00000213140 |
| 2AP // IGHD // IG | Alter. First Exon | 25/e39-40,e | up (e24-25) | 2,84 | ENSG00000211896 // ENSG00000211898 // ENSG00000213140 |
| 2AP // IGHD // IG | Alter. First Exon | 7/e39-40,e1 | up (e17) | 2,84 | ENSG00000211896 // ENSG00000211898 // ENSG00000213140 |
| 2AP // IGHD // IG | Alter. First Exon | e17/e39-40 | up (e17) | 2,84 | ENSG00000211896 // ENSG00000211898 // ENSG00000213140 |
| 2AP // IGHD // IG | Alter. First Exon | 25/e39,e40, | up (e24-25) | 2,84 | ENSG00000211896 // ENSG00000211898 // ENSG00000213140 |

| | | | | | |
|---|---|---|---|---|---|
| 2AP // IGHD // IG | Alter. First Exon | -25,e101/e3! | up (e24-25,e101) | 2,84 | ENSG00000 211896 // ENSG00000 211898 // ENSG00000 213140 |
| 2AP // IGHD // IG | Alter. First Exon | 7/e39,e40,e1 | up (e17) | 2,84 | ENSG00000 211896 // ENSG00000 211898 // ENSG00000 213140 |
| 2AP // IGHD // IG | Alter. First Exon | 7,e101/e39- | up (e17,e101) | 2,84 | ENSG00000 211896 // ENSG00000 211898 // ENSG00000 213140 |
| SLC13A3 | Exon Cassette | e13 | down | 2,84 | ENSG00000 158296 |
| DCY10P1 // NFY | Complex | e31,e34-35 | up | 2,84 | ENSG00000 001167 // ENSG00000 161912 |
| KIF1B | Alter. First Exon | 2,e14-25,e27 | e2,e4-12,e14-25,e27 | 2,83 | ENSG00000 054523 |
| TPM1 | Alter. First Exon | e1,e3/e5 | up (e1,e3) | 2,83 | ENSG00000 140416 |
| ETV4 | Alter. First Exon | e2-4/e5 | up (e2-4) | 2,83 | ENSG00000 175832 |
| NRXN1 | Alter. First Exon | 3-19,e21,e2 | 1-2,e5,e8-19,e21,e | 2,83 | ENSG00000 179915 |
| SDHAP1 | Exon Cassette | e2 | down | 2,83 | ENSG00000 185485 |
| NPNT | Complex | e14,e16 | down | 2,83 | ENSG00000 168743 |
| STXBP5 | Exon Cassette | e23,e24-25 | up | 2,83 | ENSG00000 164506 |
| STXBP5 | Exon Cassette | e23,e24 | up | 2,83 | ENSG00000 164506 |
| FUS | Intron Retention | e6 | up | 2,82 | ENSG00000 089280 |
| PCBP3 | Exon Cassette | e13 | down | 2,82 | ENSG00000 183570 |
| C5orf54 | Exon Cassette | e2 | up | 2,82 | ENSG00000 221886 |
| PRKACB | Alter. First Exon | e1/e4,e5-7 | up (e1) | 2,81 | ENSG00000 142875 |
| CD9 | ter. Terminal Exo | e4/e5,e6-7 | up (e5,e6-7) | 2,81 | ENSG00000 010278 |

| | | | | | |
|---|---|---|---|---|---|
| 2AP // IGHD // IG | Alter. First Exon | -16/e72-73,e | down (e15-16) | 2,81 | ENSG00000211896 // ENSG00000211898 // ENSG00000213140 |
| 2AP // IGHD // IG | Alter. First Exon | 15-16/e72-7 | down (e15-16) | 2,81 | ENSG00000211896 // ENSG00000211898 // ENSG00000213140 |
| 2AP // IGHD // IG | Alter. First Exon | -27/e39-40,e | up (e26-27) | 2,81 | ENSG00000211896 // ENSG00000211898 // ENSG00000213140 |
| 2AP // IGHD // IG | Alter. First Exon | 16/e72,e73,e | down (e15-16) | 2,81 | ENSG00000211896 // ENSG00000211898 // ENSG00000213140 |
| 2AP // IGHD // IG | Alter. First Exon | -16,e100/e7 | down (e15-16,e100) | 2,81 | ENSG00000211896 // ENSG00000211898 // ENSG00000213140 |
| 2AP // IGHD // IG | Alter. First Exon | 5-16/e72,e7 | down (e15-16) | 2,81 | ENSG00000211896 // ENSG00000211898 // ENSG00000213140 |
| 2AP // IGHD // IG | Alter. First Exon | -16,e101/e7 | down (e15-16,e101) | 2,81 | ENSG00000211896 // ENSG00000211898 // ENSG00000213140 |
| 2AP // IGHD // IG | Alter. First Exon | 16/e72,e73,e | down (e15-16) | 2,81 | ENSG00000211896 // ENSG00000211898 // ENSG00000213140 |

| | | | | | |
|---|---|---|---|---|---|
| 2AP // IGHD // IG | Alter. First Exon | 27,e101/e3 | up (e26-27,e101) | 2,81 | ENSG00000211896 // ENSG00000211898 // ENSG00000213140 |
| SLC12A6 | Alter. First Exon | e3/e4,e6 | down (e3) | 2,81 | ENSG00000140199 |
| GTF2IRD2 | ter. Terminal Ex | /e10,e11-16 | up (e8-9) | 2,81 | ENSG00000196275 |
| RGS10 | Alter. First Exon | e2/e3 | down (e2) | 2,8 | ENSG00000148908 |
| TMEM218 | Complex | e1/e2-3 | up (e2-3) | 2,8 | ENSG00000150433 |
| ELAC1 // SMAD4 | Exon Cassette | e10,e11 | up | 2,8 | ENSG00000141642 // ENSG00000141646 |
| CBLN2 | Alter. First Exon | e1-3/e4 | up (e1-3) | 2,8 | ENSG00000141668 |
| CASP8 | ter. Terminal Ex | 13-14/e16-1 | down (e16-18) | 2,8 | ENSG00000064012 |
| TMEM144 | Intron Retention | e1,e2 | down | 2,8 | ENSG00000164124 |
| CNOT8 | Complex | e1-2,e6 | up | 2,8 | ENSG00000155508 |
| ICA1 | Exon Cassette | e13 | up | 2,8 | ENSG00000003147 |
| PDE4B | Alter. First Exon | e1-9/e10 | down (e1-9) | 2,79 | ENSG00000184588 |
| MKL2 | Intron Retention | e12 | up | 2,79 | ENSG00000186260 |
| COLEC12 | Alter. First Exon | e1/e2 | down (e1) | 2,79 | ENSG00000158270 |
| CBLN2 | Alter. First Exon | e2-3/e4 | up (e2-3) | 2,79 | ENSG00000141668 |
| ZFAS1 | Exon Cassette | e3-4 | up | 2,79 | ENSG00000177410 |
| ANK2 | Exon Cassette | e40 | down | 2,79 | ENSG00000145362 |
| MAPK10 | Alter. First Exon | e3/e5 | down (e3) | 2,79 | ENSG00000109339 |
| 2AP // IGHD // IG | Alter. First Exon | 5-16/e64,e6 | down (e15-16) | 2,78 | ENSG00000211896 // ENSG00000211898 // ENSG00000213140 |

| | | | | | |
|---|---|---|---|---|---|
| 2AP // IGHD // IG | Alter. First Exon | 16,e100/e6 | down (e15-16,e100) | 2,78 | ENSG00000 211896 // ENSG00000 211898 // ENSG00000 213140 |
| ZNF257 | Exon Cassette | e2 | up | 2,78 | ENSG00000 197134 |
| LRRFIP1 | Alter. First Exon | e3-20,e23/e | up (e1,e3-20,e23) | 2,78 | ENSG00000 124831 |
| CD109 | Complex | 17/e18,e19-3 | down (e18,e19-33) | 2,78 | ENSG00000 156535 |
| 2AP // IGHD // IG | Alter. First Exon | 40/e72-73,e | down (e39-40) | 2,77 | ENSG00000 211896 // ENSG00000 211898 // ENSG00000 213140 |
| 2AP // IGHD // IG | Alter. First Exon | 39-40/e72-7 | down (e39-40) | 2,77 | ENSG00000 211896 // ENSG00000 211898 // ENSG00000 213140 |
| 2AP // IGHD // IG | Alter. First Exon | 40,e101/e7 | down (e39-40,e101) | 2,77 | ENSG00000 211896 // ENSG00000 211898 // ENSG00000 213140 |
| 2AP // IGHD // IG | Alter. First Exon | 39-40/e72,e7 | down (e39-40) | 2,77 | ENSG00000 211896 // ENSG00000 211898 // ENSG00000 213140 |
| 2AP // IGHD // IG | Alter. First Exon | 40/e72,e73,e | down (e39-40) | 2,77 | ENSG00000 211896 // ENSG00000 211898 // ENSG00000 213140 |
| SS18 | Exon Cassette | e5 | up | 2,77 | ENSG00000 141380 |
| DSC2 | Exon Cassette | e18 | down | 2,77 | ENSG00000 134755 |
| SH3YL1 | Alter. First Exon | e1/e2 | up (e2) | 2,77 | ENSG00000 035115 |
| DCY10P1 // NFY | Complex | e31,e33-36 | up | 2,77 | ENSG00000 001167 // ENSG00000 161912 |

| | | | | | |
|---|---|---|---|---|---|
| MACF1 | Exon Cassette | e101 | up | 2,76 | ENSG00000 127603 |
| WNK1 | Exon Cassette | e30 | down | 2,76 | ENSG00000 060237 |
| DLEU1 | Exon Cassette | e30 | down | 2,76 | ENSG00000 176124 |
| 2AP // IGHD // IG | Alter. First Exon | 101/e68-69, | down (e39-40,e101) | 2,76 | ENSG00000 211896 // ENSG00000 211898 // ENSG00000 213140 |
| SEMA6D | Exon Cassette | e21 | up | 2,76 | ENSG00000 137872 |
| HPSE | Exon Cassette | e9-10 | down | 2,76 | ENSG00000 173083 |
| LZTS1 | Complex | e1-2/e3 | down (e1-2) | 2,76 | ENSG00000 061337 |
| ANGEL2 | Complex | e1/e4 | down (e4) | 2,75 | ENSG00000 174606 |
| 2AP // IGHD // IG | Alter. First Exon | )-40,e101/e | down (e39-40,e101) | 2,75 | ENSG00000 211896 // ENSG00000 211898 // ENSG00000 213140 |
| TNPO1 | Alter. First Exon | e1/e2 | down (e1) | 2,75 | ENSG00000 083312 |

| | | | | | |
|---|---|---|---|---|---|
| )HGA11 // PCDH | Alter. Donor Site | e20 | up | 2,75 | ENSG00000 081853 // ENSG00000 204956 // ENSG00000 240184 // ENSG00000 240764 // ENSG00000 242419 // ENSG00000 253159 // ENSG00000 253305 // ENSG00000 253485 // ENSG00000 253537 // ENSG00000 253731 // ENSG00000 253767 // ENSG00000 253846 // ENSG00000 253873 // ENSG00000 253910 // ENSG00000 253953 // ENSG00000 254122 // |
| ZNF454 | Alter. Donor Site | e1 | down | 2,75 | ENSG00000 178187 |
| LEF1 | Exon Cassette | e12 | up | 2,74 | ENSG00000 138795 |
| HARS2 | Intron Retention | e6 | up | 2,74 | ENSG00000 112855 |
| CD109 | ter. Terminal Exo | 7,e34/e18-3 | down (e18-33) | 2,74 | ENSG00000 156535 |
| CD109 | Complex | e17/e18-33 | down (e18-33) | 2,74 | ENSG00000 156535 |
| RELA | lter. Acceptor Sit | e5 | up | 2,73 | ENSG00000 173039 |
| RIC8B | Exon Cassette | e3 | up | 2,73 | ENSG00000 111785 |
| )XO1 // LINC005 | Alter. First Exon | e2/e3 | down (e2) | 2,73 | ENSG00000 150907 // ENSG00000 215483 |
| )XO1 // LINC005 | Alter. First Exon | e1/e2 | down (e2) | 2,73 | ENSG00000 150907 // ENSG00000 215483 |

| | | | | | |
|---|---|---|---|---|---|
| ARHGAP44 | Exon Cassette | e17-19 | down | 2,73 | ENSG00000006740 |
| SULT1C4 | Exon Cassette | e3 | up | 2,73 | ENSG00000198075 |
| CPXM1 | Complex | e11 | up | 2,73 | ENSG00000088882 |
| EXOG | Complex | e3,e5-6 | down | 2,73 | ENSG00000157036 |
| DCY10P1 // NFY | Complex | e31,e32-36 | up | 2,73 | ENSG00000001167 // ENSG00000161912 |
| XRRA1 | Exon Cassette | e13 | up | 2,72 | ENSG00000166435 |
| BCAS3 | Exon Cassette | e18 | down | 2,72 | ENSG00000141376 |
| NR2C2 | Intron Retention | e15 | up | 2,72 | ENSG00000177463 |
| CEP44 | Alter. Donor Site | e1 | up | 2,72 | ENSG00000164118 |
| CDK5 | Intron Retention | e7 | up | 2,72 | ENSG00000164885 |
| MAPK8 | Complex | e8-9 | up | 2,71 | ENSG00000107643 |
| TMEM218 | Complex | e1-2 | up | 2,71 | ENSG00000150433 |
| GPR180 | Exon Cassette | e4 | up | 2,71 | ENSG00000152749 |
| VRK2 | Exon Cassette | e12 | down | 2,71 | ENSG00000028116 |
| FAM13A | Alter. First Exon | e1-9,e11/e1( | up (e1-9,e11) | 2,71 | ENSG00000138640 |
| HYMAI // PLAGL1 | Complex | e2,e4 | up | 2,71 | ENSG00000118495 |
| XAGE1D | Complex | e1,e2 | up | 2,71 | ENSG00000204376 |
| Sep-06 | Complex | e14,e15 | down | 2,71 | ENSG00000125354 |
| BTAF1 | Alter. Donor Site | e9 | up | 2,7 | ENSG00000095564 |
| EXOC7 | lter. Acceptor Sit | e8 | up | 2,7 | ENSG00000182473 |
| ARL17B | ter. Terminal Exo | e5/e6,e7 | up (e5) | 2,7 | ENSG00000228696 |
| 266-1 // MYT1 // | lter. Terminal Exo | e20/e21-25 | down (e21-25) | 2,7 | ENSG00000149656 // ENSG00000196132 // ENSG00000203880 |
| MLH1 | Exon Cassette | e12 | down | 2,7 | ENSG00000076242 |

| | | | | | |
|---|---|---|---|---|---|
| SLC4A4 | Alter. First Exon | e2-5/e4 | down (e2-5) | 2,7 | ENSG00000080493 |
| CD109 | ter. Terminal Exo | 17-34/e18-3 | down (e18-33) | 2,7 | ENSG00000156535 |
| SGCE | Exon Cassette | e9 | up | 2,7 | ENSG00000127990 |
| RMCX5-GPRASP | Complex | e2/e3 | up (e3) | 2,7 | ENSG00000125962 // ENSG00000158301 |
| RMCX5-GPRASP | Exon Cassette | e3 | up | 2,7 | ENSG00000125962 // ENSG00000158301 |
| SRSF11 | Alter. First Exon | e1-3/e5 | down (e1-3) | 2,69 | ENSG00000116754 |
| RIC8B | Exon Cassette | e14,e15 | up | 2,69 | ENSG00000111785 |
| ATCAY | Alter. First Exon | 'e3,e4,e6-15 | wn (e3,e4,e6-15,e1 | 2,69 | ENSG00000167654 |
| LSR | Exon Cassette | e3 | up | 2,69 | ENSG00000105699 |
| CPLX2 | Complex | e3-6 | up | 2,69 | ENSG00000145920 |
| ESRRG | Exon Cassette | e10,e11 | up | 2,68 | ENSG00000196482 |
| TACC2 | Exon Cassette | e30 | up | 2,68 | ENSG00000138162 |
| ANK3 | Complex | e1,e5/e2 | up (e2) | 2,68 | ENSG00000151150 |
| TIMM9 | Exon Cassette | e3 | down | 2,68 | ENSG00000100575 |
| MAP4K4 | ter. Terminal Exo | e17/e19-33 | up (e19-33) | 2,68 | ENSG00000071054 |
| DALRD3 | lter. Acceptor Sit | e3 | down | 2,68 | ENSG00000178149 |
| PCBP4 | Intron Retention | e6,e7 | up | 2,68 | ENSG00000090097 |
| SREK1 | Exon Cassette | e6 | up | 2,68 | ENSG00000153914 |
| MYO10 | Complex | e1-4/e3 | up (e3) | 2,68 | ENSG00000145555 |
| KRD18A // FAM9 | ter. Terminal Exo | /e16-18,e20 | own (e16-18,e20-22 | 2,68 | ENSG00000180071 |
| TCF7L2 | Exon Cassette | e18 | up | 2,67 | ENSG00000148737 |
| SLAIN1 | Alter. First Exon | e2-3/e4,e5 | up (e2-3) | 2,67 | ENSG00000139737 |

| | | | | | |
|---|---|---|---|---|---|
| 1S1 // IL4I1 // NU | Alter. First Exon | e21/e22 | up (e22) | 2,67 | ENSG00000 104951 // ENSG00000 204673 // ENSG00000 213024 |
| COLEC11 | Exon Cassette | e8 | down | 2,67 | ENSG00000 118004 |
| PLCD4 | Complex | e6,e10 | up | 2,67 | ENSG00000 115556 |
| HLA-DMA | ter. Terminal Exo | e1-3/e2,e4-6 | up (e2,e4-6) | 2,67 | ENSG00000 204257 |
| ZNF655 | ter. Terminal Exo | e4-5,e7/e10 | up (e4-5,e7) | 2,67 | ENSG00000 197343 |
| PNPLA8 | Complex | e5 | down | 2,67 | ENSG00000 135241 |
| EFHC1 | Intron Retention | e6 | up | 2,66 | ENSG00000 096093 |
| RIMS1 | Exon Cassette | e31,e32-34 | down | 2,66 | ENSG00000 079841 |
| PCSK5 | ter. Terminal Exo | e6/e7-15 | down (e7-15) | 2,66 | ENSG00000 099139 |
| KRBOX4 | Complex | e5/e6 | up (e6) | 2,66 | ENSG00000 147121 |
| RABGAP1L | ter. Terminal Exo | 30/e32,e33-3 | down (e32,e33-37) | 2,65 | ENSG00000 152061 |
| DTX3 | Intron Retention | e1,e2 | up | 2,65 | ENSG00000 178498 |
| ZC3H18 | Exon Cassette | e4 | up | 2,65 | ENSG00000 158545 |
| CADPS | Alter. First Exon | 16-18,e20-2 | e8-14,e16-18,e20-2 | 2,65 | ENSG00000 163618 |
| ZDHHC14 | lter. Acceptor Sit | e14 | down | 2,65 | ENSG00000 175048 |
| DST | Exon Cassette | e103 | up | 2,65 | ENSG00000 151914 |
| ANKRD19P | ter. Terminal Exo | -12/e13,e14 | up (e11-12) | 2,65 | ENSG00000 187984 |
| MOV10 | Alter. First Exon | e1/e2 | down (e1) | 2,64 | ENSG00000 155363 |
| PTMS | lter. Acceptor Sit | e5 | up | 2,64 | ENSG00000 159335 |
| MAP4K4 | ter. Terminal Exo | /e19-23,e25 | up (e19-23,e25-33) | 2,64 | ENSG00000 071054 |
| FLT4 | lter. Acceptor Sit | e4 | up | 2,64 | ENSG00000 037280 |
| STEAP1 | ter. Terminal Exo | e4/e5 | down (e5) | 2,64 | ENSG00000 164647 |
| DCTN3 | Intron Retention | e3 | up | 2,64 | ENSG00000 137100 |
| FUBP1 | Intron Retention | e20 | up | 2,63 | ENSG00000 162613 |

| | | | | | |
|---|---|---|---|---|---|
| FGD4 | Alter. First Exon | e2/e3 | down (e2) | 2,63 | ENSG00000 139132 |
| TBX3 | Exon Cassette | e3 | up | 2,63 | ENSG00000 135111 |
| CRIP2 | Exon Cassette | e5,e8 | down | 2,63 | ENSG00000 182809 |
| SLC47A1 | Exon Cassette | e4 | up | 2,63 | ENSG00000 142494 |
| HIPK2 | lter. Acceptor Sit | e8 | up | 2,63 | ENSG00000 064393 |
| ELAVL2 | Alter. First Exon | e1,e4/e2 | down (e1,e4) | 2,63 | ENSG00000 107105 |
| NDUFV1 | Intron Retention | e5 | up | 2,62 | ENSG00000 167792 |
| TMEM218 | Complex | e1-3 | up | 2,62 | ENSG00000 150433 |
| KIF21A | Exon Cassette | e12 | up | 2,62 | ENSG00000 139116 |
| FLT1 | Alter. First Exon | -27,e29-35/ | own (e24-27,e29-35 | 2,62 | ENSG00000 102755 |
| DCLK1 | ter. Terminal Exo | 8/e9-18,e2( | up (e8) | 2,62 | ENSG00000 133083 |
| ABAT | ter. Terminal Exo | e4/e5-18 | down (e5-18) | 2,62 | ENSG00000 183044 |
| NFATC1 | Exon Cassette | e3 | up | 2,62 | ENSG00000 131196 |
| GDAP1L1 | Exon Cassette | e5-6 | down | 2,62 | ENSG00000 124194 |
| VCAN | Exon Cassette | e7 | down | 2,62 | ENSG00000 038427 |
| FGFR1 | Complex | e6,e8 | down | 2,62 | ENSG00000 077782 |
| LPAR1 | Alter. First Exon | e3/e6 | up (e3) | 2,62 | ENSG00000 198121 |
| KRBOX4 | Complex | e5-6/e7 | up (e5-6) | 2,62 | ENSG00000 147121 |
| AP1S2 | Complex | e4/e5 | up (e5) | 2,62 | ENSG00000 182287 |
| AP1S2 | Complex | e4,e8/e5 | up (e5) | 2,62 | ENSG00000 182287 |
| PLEKHA6 | Exon Cassette | e12,e13 | down | 2,61 | ENSG00000 143850 |
| TUBB3 | Alter. First Exon | 3-8/e5-6,e9 | up (e3-8) | 2,61 | ENSG00000 198211 |
| C16orf93 | lter. Acceptor Sit | e5 | up | 2,61 | ENSG00000 196118 |
| ELP5 | ter. Terminal Exo | e6/e7-9 | up (e6) | 2,61 | ENSG00000 170291 |
| HMG20B | Intron Retention | e3,e4 | up | 2,61 | ENSG00000 064961 |
| AAK1 | Complex | e14,e15-22 | down | 2,61 | ENSG00000 115977 |

| | | | | | |
|---|---|---|---|---|---|
| UMPS | Exon Cassette | e2 | up | 2,61 | ENSG00000114491 |
| EIF4A2 | ter. Terminal Exc | e10/e11 | up (e10) | 2,61 | ENSG00000156976 |
| STXBP5 | Exon Cassette | e23 | up | 2,61 | ENSG00000164506 |
| TRIM14 | ter. Terminal Exc | e9-10/e11 | up (e9-10) | 2,61 | ENSG00000106785 |
| ZBTB40 | Intron Retention | e5 | up | 2,6 | ENSG00000184677 |
| PHYHIPL | Exon Cassette | e3 | up | 2,6 | ENSG00000165443 |
| CADM1 | Exon Cassette | e10 | down | 2,6 | ENSG00000182985 |
| FGFR1OP2 | Exon Cassette | e5 | up | 2,6 | ENSG00000111790 |
| PS31P5 // THSD | Complex | e17-18 | up | 2,6 | ENSG00000243406 |
| C15orf27 | Alter. First Exon | 1-6,e8-11/e1 | up (e1-6,e8-11) | 2,6 | ENSG00000169758 |
| EXOC7 | Exon Cassette | e8 | up | 2,6 | ENSG00000182473 |
| COLEC11 | Exon Cassette | e6,e8 | down | 2,6 | ENSG00000118004 |
| LRRFIP2 | Exon Cassette | e20 | up | 2,6 | ENSG00000093167 |
| CADPS | Alter. First Exon | ,e23-24,e26 | -18,e21,e23-24,e26 | 2,6 | ENSG00000163618 |
| CADPS | Alter. First Exon | e21,e23-24, | e16-19,e21,e23-24, | 2,6 | ENSG00000163618 |
| PRDM8 | Alter. First Exon | e2-7/e8 | up (e2-7) | 2,6 | ENSG00000152784 |
| ZNF454 | Complex | e1,e2 | down | 2,6 | ENSG00000178187 |
| ZNF655 | ter. Terminal Exc | 4-5,e7/e9-1( | up (e4-5,e7) | 2,6 | ENSG00000197343 |
| SMARCD3 | Intron Retention | e11 | up | 2,6 | ENSG00000082014 |
| KLF10 | Alter. First Exon | e1/e2 | up (e1) | 2,6 | ENSG00000155090 |
| SHISA4 | Complex | e1 | up | 2,59 | ENSG00000198892 |
| C15orf27 | Alter. First Exon | e1-11/e12 | up (e1-11) | 2,59 | ENSG00000169758 |
| PTPRS | Exon Cassette | 13-14,e16-1 | up | 2,59 | ENSG00000105426 |
| ZNF260 | Complex | e2-3 | up | 2,59 | ENSG00000254004 |
| CASP8 | Complex | e4-5/e7 | down (e4-5) | 2,59 | ENSG00000064012 |
| RTKN | Alter. First Exon | e1/e2 | up (e1) | 2,59 | ENSG00000114993 |

| | | | | | |
|---|---|---|---|---|---|
| RASGEF1B | Exon Cassette | e3 | up | 2,59 | ENSG00000 138670 |
| KN2B-AS1 // MT | ualy Exclusive Ex | e19-23/e25 | down (e19-23) | 2,59 | ENSG00000 099810 // ENSG00000 240498 |
| ANGPTL2 | Complex | e2-5 | up | 2,59 | ENSG00000 136859 |
| NFASC | Exon Cassette | e26-27 | up | 2,58 | ENSG00000 163531 |
| NRP1 | ter. Terminal Exo | e10/e11,e13 | down (e11,e13) | 2,58 | ENSG00000 099250 |
| C11orf30 | Exon Cassette | e5 | up | 2,58 | ENSG00000 158636 |
| BTBD11 | Alter. First Exon | e1-3/e4 | down (e1-3) | 2,58 | ENSG00000 151136 |
| ELP5 | ter. Terminal Exo | e6/e7-10 | up (e6) | 2,58 | ENSG00000 170291 |
| SLC13A3 | Alter. First Exon | e1-2/e4 | down (e1-2) | 2,58 | ENSG00000 158296 |
| FOXP2 | Exon Cassette | e19 | down | 2,58 | ENSG00000 128573 |
| GTF2IRD2 | ter. Terminal Exo | e7-8,e10-16, | up (e6) | 2,58 | ENSG00000 196275 |
| FGFR1 | Exon Cassette | e6 | down | 2,58 | ENSG00000 077782 |
| DOCK7 | Exon Cassette | e24 | down | 2,57 | ENSG00000 116641 |
| S100A13 | Complex | e4,e5-9 | down | 2,57 | ENSG00000 189171 |
| SYNRG | Alter. Donor Site | e8 | up | 2,57 | ENSG00000 006114 |
| YPEL1 | Complex | e2,e3 | up | 2,57 | ENSG00000 100027 |
| USP45 | Exon Cassette | e14 | down | 2,57 | ENSG00000 123552 |
| TTF1 | Exon Cassette | e2 | down | 2,57 | ENSG00000 125482 |
| DTX3 | lter. Acceptor Sit | e2 | up | 2,56 | ENSG00000 178498 |
| APA1 // RALGA | Complex | e20/e21 | up (e20) | 2,56 | ENSG00000 174373 // ENSG00000 229419 |
| NUDT7 | Exon Cassette | e4,e5 | up | 2,56 | ENSG00000 140876 |
| CADPS | Alter. First Exon | e21,e23-24, | e16-18,e21,e23-24, | 2,56 | ENSG00000 163618 |
| MAGI1 | Exon Cassette | e28 | down | 2,56 | ENSG00000 151276 |
| FAM221A | Exon Cassette | e6,e7 | up | 2,56 | ENSG00000 188732 |

| | | | | | |
|---|---|---|---|---|---|
| ST7 // ST7-OT3 | Exon Cassette | e4 | up | 2,56 | ENSG00000 004866 |
| RBM8A | Intron Retention | e2-3 | up | 2,55 | ENSG00000 131795 |
| TMEM218 | Complex | e1,e3 | up | 2,55 | ENSG00000 150433 |
| OSBPL8 | Complex | e7,e8 | down | 2,55 | ENSG00000 091039 |
| SEMA6D | Exon Cassette | e20,e21 | up | 2,55 | ENSG00000 137872 |
| 1S1 // IL4I1 // NU | Intron Retention | e2-3 | up | 2,55 | ENSG00000 104951 // ENSG00000 204673 // ENSG00000 213024 |
| TRAF3IP2-AS1 | ter. Terminal Exc | e4/e6-8 | up (e4) | 2,55 | ENSG00000 231889 |
| NUDT10 | Alter. First Exon | e1/e2 | up (e1) | 2,55 | ENSG00000 122824 |
| RPS6KC1 | Intron Retention | e14 | up | 2,54 | ENSG00000 136643 |
| CTNNBIP1 | Alter. First Exon | e1-4/e5 | up (e1-4) | 2,54 | ENSG00000 178585 |
| OSBPL8 | Exon Cassette | e7 | down | 2,54 | ENSG00000 091039 |
| NDRG4 | lter. Acceptor Sit | e8 | up | 2,54 | ENSG00000 103034 |
| --- | Exon Cassette | e5,e6-8 | up | 2,54 | --- |
| NOSIP | Complex | e7 | up | 2,54 | ENSG00000 142546 |
| PLCD4 | ter. Terminal Exc | 1/e12,e13-1 | up (e11) | 2,54 | ENSG00000 115556 |
| PPP2R3A | Exon Cassette | e2 | down | 2,54 | ENSG00000 073711 |
| QARS | lter. Acceptor Sit | e11 | up | 2,54 | ENSG00000 172053 |
| REST | ter. Terminal Exc | -8,e12/e14- | up (e5-8,e12) | 2,54 | ENSG00000 084093 |
| REST | ter. Terminal Exc | 8,e12/e13,e | up (e5-8,e12) | 2,54 | ENSG00000 084093 |
| CAMK2D | Exon Cassette | e15,e16 | down | 2,54 | ENSG00000 145349 |
| FGFR1 | Exon Cassette | e5,e6 | down | 2,54 | ENSG00000 077782 |
| ESRRG | Complex | e9,e10-11 | up | 2,53 | ENSG00000 196482 |
| ETS1 | Exon Cassette | e5-8 | down | 2,53 | ENSG00000 134954 |
| WARS | Exon Cassette | e3 | up | 2,53 | ENSG00000 140105 |

| | | | | | |
|---|---|---|---|---|---|
| TPM1 | Alter. First Exon | e1-3/e5 | up (e1-3) | 2,53 | ENSG00000140416 |
| ATP5G1 | Alter. First Exon | e1/e2 | down (e1) | 2,53 | ENSG00000159199 |
| ARL17B | ter. Terminal Exo | e5/e7,e8-9 | up (e5) | 2,53 | ENSG00000228696 |
| NRG1 | Complex | e5/e6,e10 | up (e6,e10) | 2,53 | ENSG00000157168 |
| GRIN1 | ter. Terminal Exo | e19/e20-21 | up (e20-21) | 2,53 | ENSG00000176884 |
| SFR1 | Intron Retention | e1,e2 | up | 2,52 | ENSG00000156384 |
| GABARAPL1 | Exon Cassette | e2,e3 | up | 2,52 | ENSG00000139112 |
| TYK2 | Complex | e1-7 | up | 2,52 | ENSG00000105397 |
| HPSE | Complex | e2-4,e6-9 | up | 2,52 | ENSG00000173083 |
| NR2F1-AS1 | Exon Cassette | e7,e10 | up | 2,52 | ENSG00000237187 |
| PTPRK | Complex | e1-4/e3 | up (e3) | 2,52 | ENSG00000152894 |
| --- | Exon Cassette | e9 | up | 2,52 | --- |
| SEC11A | Exon Cassette | e2 | down | 2,51 | ENSG00000140612 |
| MGRN1 | Exon Cassette | e12 | up | 2,51 | ENSG00000102858 |
| ABAT | ter. Terminal Exo | e4/e6-18 | down (e6-18) | 2,51 | ENSG00000183044 |
| CEP68 | Complex | e3,e4 | up | 2,51 | ENSG00000011523 |
| DENND6B | Intron Retention | e7 | up | 2,51 | ENSG00000205593 |
| REST | ter. Terminal Exo | -9,e12/e14- | up (e5-9,e12) | 2,51 | ENSG00000084093 |
| REST | ter. Terminal Exo | 9,e12/e13,e | up (e5-9,e12) | 2,51 | ENSG00000084093 |
| FAM13A | Alter. First Exon | e6-9,e11-12 | p (e2-4,e6-9,e11-12 | 2,51 | ENSG00000138640 |
| NONO | Exon Cassette | e3 | up | 2,51 | ENSG00000147140 |
| TCF7L2 | Alter. First Exon | e2-3/e5 | down (e2-3) | 2,5 | ENSG00000148737 |
| PLEKHA5 | Complex | e1,e2-3,e7 | up | 2,5 | ENSG00000052126 |
| IKBIP | Complex | e1-2/e4 | up (e1-2) | 2,5 | ENSG00000166130 |
| EIF5A | ter. Terminal Exo | e6/e7-10 | up (e6) | 2,5 | ENSG00000132507 |
| ATP5A1 | Alter. Donor Site | e3 | up | 2,5 | ENSG00000152234 |

| | | | | | |
|---|---|---|---|---|---|
| ZNF135 | Exon Cassette | e2-3 | up | 2,5 | ENSG00000176293 |
| ZNF266 | Intron Retention | e7 | up | 2,5 | ENSG00000174652 |
| RTN4 | Exon Cassette | e7-8 | down | 2,5 | ENSG00000115310 |
| KLHL5 | Alter. First Exon | e1/e2 | up (e2) | 2,5 | ENSG00000109790 |
| MATR3 // SNHG4 | Alter. First Exon | e1-4/e9 | up (e1-4) | 2,5 | ENSG00000015479 |
| RWDD2A | Intron Retention | e2 | up | 2,5 | ENSG00000013392 |
| PNPLA8 | Exon Cassette | e3 | up | 2,5 | ENSG00000135241 |
| PRKACB | Alter. First Exon | e1/e4,e7 | up (e1) | 2,49 | ENSG00000142875 |
| DCLK1 | ter. Terminal Exo | e8/e14-20 | up (e8) | 2,49 | ENSG00000133083 |
| NK1G1 // KIAA01 | ter. Terminal Exo | e10,e12-17,( | up (e5) | 2,49 | ENSG00000166803 // ENSG00000169118 |
| MAP2K4 | Exon Cassette | e2 | up | 2,49 | ENSG00000065559 |
| GPR155 | Exon Cassette | e2 | up | 2,49 | ENSG00000163328 |
| // PRR5 // PRR5 | Alter. First Exon | e4-7/e13 | down (e4-7) | 2,49 | ENSG00000186654 // ENSG00000241484 // ENSG00000248405 |
| ANK2 | Exon Cassette | e32 | down | 2,49 | ENSG00000145362 |
| GTF2IRD2 | ter. Terminal Exo | 6/e7-8,e10-1 | up (e6) | 2,49 | ENSG00000196275 |
| KRBOX4 | Complex | e5,e6 | up | 2,49 | ENSG00000147121 |
| ZNF195 | Exon Cassette | e6 | up | 2,48 | ENSG00000005801 |
| GREB1 | Complex | e3/e4 | down (e4) | 2,48 | ENSG00000196208 |
| FN1 | Exon Cassette | e43 | up | 2,48 | ENSG00000115414 |
| MATR3 // SNHG4 | ter. Terminal Exo | e4/e9-22 | up (e4) | 2,48 | ENSG00000015479 |
| PPP1R18 | Alter. First Exon | e1/e2 | up (e1) | 2,48 | ENSG00000146112 |
| PLEKHA6 | Exon Cassette | e11,e13 | down | 2,47 | ENSG00000143850 |
| ACTA2 | Exon Cassette | e3 | up | 2,47 | ENSG00000107796 |

| | | | | | |
|---|---|---|---|---|---|
| TBCK | Exon Cassette | e7 | up | 2,47 | ENSG00000145348 |
| MATR3 // SNHG4 | ter. Terminal Exo | e4/e10-22 | up (e4) | 2,47 | ENSG00000015479 |
| J2 // RASA4 // U | Exon Cassette | e3 | up | 2,47 | ENSG00000105808 // ENSG00000267368 // ENSG00000267645 |
| FAM167A | Alter. First Exon | e1/e3,e4 | up (e3,e4) | 2,47 | ENSG00000154319 |
| SRSF11 | Alter. First Exon | e1-3/e5-7 | down (e1-3) | 2,46 | ENSG00000116754 |
| RGS4 | Complex | e5-6 | down | 2,46 | ENSG00000117152 |
| MKI67 | Exon Cassette | e7 | down | 2,46 | ENSG00000148773 |
| ERBB3 | Complex | e1,e4-17 | up | 2,46 | ENSG00000065361 |
| ZFAND6 | Alter. First Exon | e2,e5/e4 | up (e2,e5) | 2,46 | ENSG00000086666 |
| ZNF423 | Alter. First Exon | e1,e4/e3 | up (e3) | 2,46 | ENSG00000102935 |
| NCBP2 | Intron Retention | e3 | up | 2,46 | ENSG00000114503 |
| KIAA0226 | Exon Cassette | e14 | up | 2,46 | ENSG00000145016 |
| REST | Alter. First Exon | e1/e2-3 | down (e1) | 2,46 | ENSG00000084093 |
| REST | Alter. First Exon | e1/e2 | down (e1) | 2,46 | ENSG00000084093 |
| REST | Alter. First Exon | e1/e4 | down (e1) | 2,46 | ENSG00000084093 |
| TRAF3IP2-AS1 | ter. Terminal Exo | e4/e8 | up (e4) | 2,46 | ENSG00000231889 |
| TRAF3IP2-AS1 | ter. Terminal Exo | e2-3,e5/e6-8 | up (e2-3,e5) | 2,46 | ENSG00000231889 |
| ZC4H2 | Exon Cassette | e3 | up | 2,46 | ENSG00000126970 |
| OSBPL8 | Exon Cassette | e2,e3-4 | up | 2,45 | ENSG00000091039 |
| DCLK1 | ter. Terminal Exo | e8/e9-20 | up (e8) | 2,45 | ENSG00000133083 |
| F6 // HEXA // PA | Exon Cassette | e51 | down | 2,45 | ENSG00000137817 // ENSG00000140488 // ENSG00000213614 |
| ORC6 | ter. Terminal Exo | e4/e5-7 | up (e4) | 2,45 | ENSG00000091651 |

| | | | | | |
|---|---|---|---|---|---|
| ACADVL | ter. Terminal Exo | e11/e12-22 | down (e12-22) | 2,45 | ENSG00000 072778 |
| NARF | Intron Retention | e12 | up | 2,45 | ENSG00000 141562 |
| SNRNP70 | Exon Cassette | e8 | up | 2,45 | ENSG00000 104852 |
| RTKN | Alter. First Exon | e1/e2-3 | up (e1) | 2,45 | ENSG00000 114993 |
| ARFIP1 | Exon Cassette | e4 | up | 2,45 | ENSG00000 164144 |
| TACC1 | ter. Terminal Exo | e8,e10,e12- | up (e8,e10,e12-21) | 2,45 | ENSG00000 147526 |
| EBNA1BP2 | Intron Retention | e3 | up | 2,44 | ENSG00000 117395 |
| P2RX7 | Exon Cassette | e6 | up | 2,44 | ENSG00000 089041 |
| PDK2 | Alter. First Exon | e1,e4/e3 | up (e1,e4) | 2,44 | ENSG00000 005882 |
| DMKN | Exon Cassette | e9,e10,e12 | down | 2,44 | ENSG00000 161249 |
| ZNF329 | Exon Cassette | e5 | up | 2,44 | ENSG00000 181894 |
| CNTNAP5 | Complex | e18,e19-24 | down | 2,44 | ENSG00000 155052 |
| 266-1 // MYT1 // I | Exon Cassette | e9 | up | 2,44 | ENSG00000 149656 // ENSG00000 196132 // ENSG00000 203880 |
| HAUS3 // POLN | ter. Terminal Exo | 7-8/e9,e10-2 | down (e7-8) | 2,44 | ENSG00000 130997 // ENSG00000 214367 |
| HLA-DMB | Exon Cassette | e4 | down | 2,44 | ENSG00000 242574 |
| NFASC | Exon Cassette | e30,e31-32 | up | 2,43 | ENSG00000 163531 |
| CD59 | Alter. First Exon | e1-3/e5 | down (e1-3) | 2,43 | ENSG00000 085063 |
| LGR5 | Complex | e11,e12 | up | 2,43 | ENSG00000 139292 |
| C14orf2 | Complex | e4 | up | 2,43 | ENSG00000 156411 |
| C14orf2 | Exon Cassette | e4 | up | 2,43 | ENSG00000 156411 |
| MFGE8 | Exon Cassette | e4 | up | 2,43 | ENSG00000 140545 |
| GPR56 | Complex | e6-16 | up | 2,43 | ENSG00000 205336 |
| ETV4 | Alter. First Exon | e3-4/e5 | up (e3-4) | 2,43 | ENSG00000 175832 |

| | | | | | |
|---|---|---|---|---|---|
| --- | Exon Cassette | e5,e6,e11 | up | 2,43 | --- |
| ZNF638 | Complex | e2/e3 | up (e3) | 2,43 | ENSG00000075292 |
| FBN2 | Exon Cassette | e9 | up | 2,43 | ENSG00000138829 |
| DPYSL2 | Complex | e1,e4-12 | down | 2,43 | ENSG00000092964 |
| FZD3 | Exon Cassette | e2 | up | 2,43 | ENSG00000104290 |
| NRG1 | Exon Cassette | e6 | up | 2,43 | ENSG00000157168 |
| STMN2 | Alter. First Exon | e1/e2 | up (e1) | 2,43 | ENSG00000104435 |
| EPB41 | Exon Cassette | e15,e16-17 | down | 2,42 | ENSG00000159023 |
| KCNT2 | Exon Cassette | e20 | down | 2,42 | ENSG00000162687 |
| FAM13C | ter. Terminal Exo | e12/e13,e14 | up (e13,e14) | 2,42 | ENSG00000148541 |
| MARK3 | Exon Cassette | e16-17 | up | 2,42 | ENSG00000075413 |
| CTSH | Exon Cassette | e3 | down | 2,42 | ENSG00000103811 |
| SLC44A2 | Complex | e23 | down | 2,42 | ENSG00000129353 |
| KIAA0930 | Alter. First Exon | e4/e6 | up (e4) | 2,42 | ENSG00000100364 |
| DDX46 | Exon Cassette | e13 | up | 2,42 | ENSG00000145833 |
| MATR3 // SNHG4 | ter. Terminal Exo | 4/e6-7,e9-2 | up (e4) | 2,42 | ENSG00000015479 |
| NSUN5 | Intron Retention | e2 | up | 2,42 | ENSG00000130305 |
| FGFR1 | Exon Cassette | e6,e7 | down | 2,42 | ENSG00000077782 |
| GARNL3 | Exon Cassette | e24,e25 | up | 2,42 | ENSG00000136895 |
| KRBOX4 | Exon Cassette | e6 | up | 2,42 | ENSG00000147121 |
| LPHN2 | Alter. First Exon | e1-3,e6/e5 | down (e1-3,e6) | 2,41 | ENSG00000117114 |
| USF1 | Alter. First Exon | e1/e2 | up (e1) | 2,41 | ENSG00000158773 |
| BBIP1 | Exon Cassette | e4 | up | 2,41 | ENSG00000214413 |
| XRRA1 | Exon Cassette | e11,e12 | down | 2,41 | ENSG00000166435 |
| CCDC90B | Complex | e1-3 | down | 2,41 | ENSG00000137500 |
| CACNB3 | Exon Cassette | e6 | down | 2,41 | ENSG00000167535 |

| | | | | | |
|---|---|---|---|---|---|
| 2AP // IGHD // IG | Alter. First Exon | )-40,e103/e | down (e39-40,e103) | 2,41 | ENSG00000 211896 // ENSG00000 211898 // ENSG00000 213140 |
| GREB1L | Exon Cassette | e12 | up | 2,41 | ENSG00000 141449 |
| ZNF260 | Exon Cassette | e4 | up | 2,41 | ENSG00000 254004 |
| VRK2 | Exon Cassette | e21 | down | 2,41 | ENSG00000 028116 |
| ZNF638 | lter. Acceptor Sit | e3 | up | 2,41 | ENSG00000 075292 |
| WDR6 | Intron Retention | e4 | up | 2,41 | ENSG00000 178252 |
| REST | ualy Exclusive Ex | e5-8/e14 | up (e5-8) | 2,41 | ENSG00000 084093 |
| REST | ualy Exclusive Ex | e5-8/e13 | up (e5-8) | 2,41 | ENSG00000 084093 |
| HLA-DMB | Exon Cassette | e5 | up | 2,41 | ENSG00000 242574 |
| 1 // FAM188B // | Complex | e27/e27,e31 | down (e27,e31) | 2,41 | ENSG00000 106125 // ENSG00000 240583 // ENSG00000 241644 |
| 1 // FAM188B // | Complex | e27,e31/e27 | down (e27,e31) | 2,41 | ENSG00000 106125 // ENSG00000 240583 // ENSG00000 241644 |
| GTPBP10 | Exon Cassette | e3 | up | 2,41 | ENSG00000 105793 |
| PNPLA8 | Complex | e2/e3-5 | up (e3-5) | 2,41 | ENSG00000 135241 |
| NAV2 | Exon Cassette | e25 | up | 2,4 | ENSG00000 166833 |
| DOCK9 | lter. Acceptor Sit | e43 | up | 2,4 | ENSG00000 088387 |
| EML5 | Exon Cassette | e12 | up | 2,4 | ENSG00000 165521 |
| CHRNB4 | Alter. Donor Site | e12 | up | 2,4 | ENSG00000 117971 |
| FAM83G | ter. Terminal Exo | e5-6/e7 | down (e7) | 2,4 | ENSG00000 188522 |
| AHRR // PDCD6 | Intron Retention | e4 | up | 2,4 | ENSG00000 063438 // ENSG00000 249915 |

| | | | | | |
|---|---|---|---|---|---|
| CDH23 | lter. Acceptor Sit | e8 | down | 2,39 | ENSG00000107736 |
| PAX6 | Alter. Donor Site | e3 | down | 2,39 | ENSG00000007372 |
| SULT1A1 | Alter. First Exon | e1-4/e6 | down (e1-4) | 2,39 | ENSG00000196502 |
| MED24 | Complex | e8 | up | 2,39 | ENSG00000008838 |
| MED24 | Exon Cassette | e8 | up | 2,39 | ENSG00000008838 |
| INPP4A | Alter. Donor Site | e17 | up | 2,39 | ENSG00000040933 |
| KIDINS220 | Exon Cassette | e28,e29 | up | 2,39 | ENSG00000134313 |
| ITSN1 | Exon Cassette | e38 | up | 2,39 | ENSG00000205726 |
| CAMK2D | Exon Cassette | e16 | down | 2,39 | ENSG00000145349 |
| MAGI2-AS3 | Exon Cassette | e9 | up | 2,39 | ENSG00000234456 |
| TPM3 | Intron Retention | e11 | up | 2,38 | ENSG00000143549 |
| FGFR1OP2 | ter. Terminal Exo | e5/e6-7 | up (e5) | 2,38 | ENSG00000111790 |
| RIC8B | Exon Cassette | e11,e12-15 | up | 2,38 | ENSG00000111785 |
| SNHG16 | Complex | e1,e2 | up | 2,38 | ENSG00000163597 |
| SRSF1 | Intron Retention | e4 | down | 2,38 | ENSG00000136450 |
| GTPBP3 | Complex | e10 | down | 2,38 | ENSG00000130299 |
| ZNF83 | Exon Cassette | e11,e13 | up | 2,38 | ENSG00000167766 |
| EXOC1 | Exon Cassette | e11 | down | 2,38 | ENSG00000090989 |
| REST | Complex | e5,e8/e12 | up (e5,e8) | 2,38 | ENSG00000084093 |
| REST | Exon Cassette | e7-8 | up | 2,38 | ENSG00000084093 |
| REST | Exon Cassette | e8 | up | 2,38 | ENSG00000084093 |
| REST | Exon Cassette | e6,e8 | up | 2,38 | ENSG00000084093 |
| RARS2 | Alter. First Exon | e1/e2 | up (e1) | 2,38 | ENSG00000146282 |
| PPFIA1 | Complex | e24,e25-26 | down | 2,37 | ENSG00000131626 |
| ACADVL | Intron Retention | e12 | up | 2,37 | ENSG00000072778 |
| GPATCH8 | Exon Cassette | e5 | up | 2,37 | ENSG00000186566 |

| | | | | | |
|---|---|---|---|---|---|
| OAZ1 | Intron Retention | e2 | up | 2,37 | ENSG00000104904 |
| NDUFAF7 | Intron Retention | e5,e6 | up | 2,37 | ENSG00000003509 |
| CD200 | Exon Cassette | e3 | down | 2,37 | ENSG00000091972 |
| TRAF3IP2-AS1 | ter. Terminal Exo | e2-3,e5/e8 | up (e2-3,e5) | 2,37 | ENSG00000231889 |
| MVB12B | ter. Terminal Exo | e8/e9,e10-12 | up (e8) | 2,37 | ENSG00000196814 |
| AQP3 | Intron Retention | e4 | down | 2,37 | ENSG00000165272 |
| STAG2 | Exon Cassette | e4 | up | 2,37 | ENSG00000101972 |
| SRSF4 | Intron Retention | e6,e7 | up | 2,36 | ENSG00000116350 |
| TNR | Alter. First Exon | e1-2/e3 | down (e1-2) | 2,36 | ENSG00000116147 |
| DCLK1 | ter. Terminal Exo | e9-16,e18,e | up (e8) | 2,36 | ENSG00000133083 |
| APBA2 | Exon Cassette | e11 | down | 2,36 | ENSG00000034053 |
| NCOR1 | Exon Cassette | e25 | up | 2,36 | ENSG00000141027 |
| EXOC7 | Complex | e7-8 | down | 2,36 | ENSG00000182473 |
| RBBP8 | Alter. First Exon | e2/e3 | down (e2) | 2,36 | ENSG00000101773 |
| ATP9B | Exon Cassette | e36 | down | 2,36 | ENSG00000166377 |
| OX17 // POPDC | ualy Exclusive Ex | e4-5/e8-9 | up (e4-5) | 2,36 | ENSG00000121577 // ENSG00000138495 |
| OX17 // POPDC | Exon Cassette | e4,e5 | up | 2,36 | ENSG00000121577 // ENSG00000138495 |
| MYB | Exon Cassette | e9,e10 | up | 2,36 | ENSG00000118513 |
| TSEN15 | Exon Cassette | e5 | up | 2,35 | ENSG00000198860 |
| SRSF4 | Exon Cassette | e3,e4-5 | up | 2,35 | ENSG00000116350 |
| PEX11A | Exon Cassette | e3 | up | 2,35 | ENSG00000166821 |
| KANSL1 | Exon Cassette | e11 | up | 2,35 | ENSG00000120071 |
| ZNF83 | Exon Cassette | e10,e11,e13 | up | 2,35 | ENSG00000167766 |
| ZNF550 | Exon Cassette | e5 | up | 2,35 | ENSG00000251369 |

| | | | | | |
|---|---|---|---|---|---|
| POLR1B | Exon Cassette | e2 | up | 2,35 | ENSG00000125630 |
| CNTNAP5 | ter. Terminal Exc | e18-24/e25 | down (e18-24) | 2,35 | ENSG00000155052 |
| SYNGR1 // TAB1 | Intron Retention | e3 | down | 2,35 | ENSG00000100321 // ENSG00000100324 |
| OPRM1 | Complex | e1,e2-3 | up | 2,35 | ENSG00000112038 |
| C7orf43 | lter. Acceptor Sit | e9 | up | 2,35 | ENSG00000146826 |
| PLEC | Alter. First Exon | e4/e7 | down (e4) | 2,35 | ENSG00000178209 |
| SRSF11 | Alter. First Exon | e1-3/e5,e6-7 | down (e1-3) | 2,34 | ENSG00000116754 |
| TCF7L2 | Alter. First Exon | 1-3,e5,e10/e | down (e1-3,e5,e10) | 2,34 | ENSG00000148737 |
| TCF7L2 | Alter. First Exon | 1-3,e5,e10/e | down (e1-3,e5,e10) | 2,34 | ENSG00000148737 |
| TCF7L2 | Alter. First Exon | e1-3,e5-6/e8 | down (e1-3,e5-6) | 2,34 | ENSG00000148737 |
| TCF7L2 | Alter. First Exon | e1-3,e5-6/e9 | down (e1-3,e5-6) | 2,34 | ENSG00000148737 |
| RELA | Intron Retention | e4,e5 | up | 2,34 | ENSG00000173039 |
| NUMA1 | Exon Cassette | e21 | up | 2,34 | ENSG00000137497 |
| ZDHHC7 | Exon Cassette | e4 | up | 2,34 | ENSG00000153786 |
| C00338 // SEC1 | ter. Terminal Exc | 4/e8-9,e11-2 | down (e8-9,e11-25) | 2,34 | ENSG00000129657 // ENSG00000234912 |
| SERPINE2 | Alter. First Exon | e3-4/e5 | down (e3-4) | 2,34 | ENSG00000135919 |
| TOP2B | Alter. First Exon | e1/e2 | down (e1) | 2,34 | ENSG00000077097 |
| ADH5 | ter. Terminal Exc | e6/e7-9 | up (e6) | 2,34 | ENSG00000197894 |
| MATR3 // SNHG4 | Alter. First Exon | e1-4,e6-7/e9 | up (e1-4,e6-7) | 2,34 | ENSG00000015479 |
| FAM126A | Exon Cassette | e2 | up | 2,34 | ENSG00000122591 |
| MRRF | Complex | e5,e6-7 | up | 2,34 | ENSG00000148187 |
| ODF2 | Exon Cassette | e5 | up | 2,34 | ENSG00000136811 |
| DNALI1 | ter. Terminal Exc | e3/e4-6 | down (e4-6) | 2,33 | ENSG00000163879 |
| C1orf85 | Complex | e1-2 | up | 2,33 | ENSG00000198715 |

| | | | | | |
|---|---|---|---|---|---|
| OPN3 | Alter. First Exon | e1-2/e3,e4 | up (e1-2) | 2,33 | ENSG00000 054277 |
| VEZT | Exon Cassette | e18 | up | 2,33 | ENSG00000 028203 |
| DCLK1 | ter. Terminal Exo | e8/e9-14 | up (e8) | 2,33 | ENSG00000 133083 |
| TJP1 | Exon Cassette | e24 | up | 2,33 | ENSG00000 104067 |
| 1S1 // IL4I1 // NU | Alter. First Exon | e19/e21 | up (e19) | 2,33 | ENSG00000 104951 // ENSG00000 204673 // ENSG00000 213024 |
| EPB41L1 | Complex | e2,e10/e3-7 | up (e3-7) | 2,33 | ENSG00000 088367 |
| VGLL4 | Alter. First Exon | e4-6/e7 | down (e4-6) | 2,33 | ENSG00000 144560 |
| WDR46 | Complex | e1/e2,e3 | down (e2,e3) | 2,33 | ENSG00000 227057 |
| CASP2 | Intron Retention | e10 | up | 2,33 | ENSG00000 106144 |
| FAM120C | ter. Terminal Exo | 3,e4-12,e15 | up (e2) | 2,33 | ENSG00000 184083 |
| MXI1 | Exon Cassette | e8 | up | 2,32 | ENSG00000 119950 |
| ORC6 | ter. Terminal Exo | e4/e5-6 | up (e4) | 2,32 | ENSG00000 091651 |
| --- | Exon Cassette | e5-8,e10-11 | up | 2,32 | --- |
| HNRNPL | Exon Cassette | e8 | up | 2,32 | ENSG00000 104824 |
| WDR6 | Complex | e1/e4 | up (e4) | 2,32 | ENSG00000 178252 |
| HAUS3 // POLN | Complex | 18/e19,e20-2 | up (e19,e20-21) | 2,32 | ENSG00000 130997 // ENSG00000 214367 |
| HAUS3 // POLN | Exon Cassette | e19-21 | up | 2,32 | ENSG00000 130997 // ENSG00000 214367 |
| MFSD8 | Exon Cassette | e11 | up | 2,32 | ENSG00000 164073 |
| WDR46 | Complex | e2 | down | 2,32 | ENSG00000 227057 |
| WDR46 | Intron Retention | e5 | up | 2,32 | ENSG00000 227057 |
| VPS52 | Intron Retention | e13 | up | 2,32 | ENSG00000 223501 |
| BBS9 | Exon Cassette | e17 | down | 2,32 | ENSG00000 122507 |

| | | | | | |
|---|---|---|---|---|---|
| OXR1 | Alter. First Exon | e1,e4-11/e3 | up (e1,e4-11) | 2,32 | ENSG00000164830 |
| POMGNT1 | Intron Retention | e12 | up | 2,31 | ENSG00000085998 |
| C1orf85 | Exon Cassette | e2 | up | 2,31 | ENSG00000198715 |
| LDB1 | ter. Terminal Exo | e11/e12 | up (e11) | 2,31 | ENSG00000198728 |
| VPS13C | ter. Terminal Exo | e83/e84-87 | down (e83) | 2,31 | ENSG00000129003 |
| TXNDC11 | Complex | e1/e1-2 | up (e1-2) | 2,31 | ENSG00000153066 |
| MAP4K4 | ter. Terminal Exo | /e18-23,e25 | up (e18-23,e25-33) | 2,31 | ENSG00000071054 |
| PDIA5 | Intron Retention | e16 | down | 2,31 | ENSG00000065485 |
| OX17 // POPDC | ualy Exclusive Ex | e5/e7-9 | up (e5) | 2,31 | ENSG00000121577 // ENSG00000138495 |
| FGF1 | Exon Cassette | e8 | down | 2,31 | ENSG00000113578 |
| IICAL1 // ZBTB2 | Intron Retention | e24 | up | 2,31 | ENSG00000112365 // ENSG00000135596 |
| 1 // FAM188B // | Complex | e27,e31 | down | 2,31 | ENSG00000106125 // ENSG00000240583 // ENSG00000241644 |
| MID1 | Alter. First Exon | e13/e14,e15 | down (e14,e15) | 2,31 | ENSG00000101871 |
| TCF7L2 | Alter. First Exon | e2-3,e5-6/e8 | down (e2-3,e5-6) | 2,3 | ENSG00000148737 |
| TCF7L2 | Alter. First Exon | e2-3,e5-6/e9 | down (e2-3,e5-6) | 2,3 | ENSG00000148737 |
| TCF7L2 | Exon Cassette | e20 | down | 2,3 | ENSG00000148737 |
| DBI | Complex | e1,e2 | up | 2,3 | ENSG00000155368 |
| DBI | Complex | e1-2 | up | 2,3 | ENSG00000155368 |
| KIDINS220 | Exon Cassette | e28 | up | 2,3 | ENSG00000134313 |
| HAND2-AS1 | Exon Cassette | e3,e4 | down | 2,3 | ENSG00000237125 |
| HPSE | Exon Cassette | e5 | down | 2,3 | ENSG00000173083 |
| ORC3 | Exon Cassette | e15 | up | 2,3 | ENSG00000135336 |

| | | | | | |
|---|---|---|---|---|---|
| FYN | ualy Exclusive Ex | e11/e12 | down (e11) | 2,3 | ENSG00000010810 |
| MAP7 | Exon Cassette | e3 | down | 2,3 | ENSG00000135525 |
| BNIP3L | Alter. First Exon | e2/e3 | up (e2) | 2,3 | ENSG00000104765 |
| NLGN3 | Alter. First Exon | e1-3/e4 | up (e1-3) | 2,3 | ENSG00000196338 |
| FAM120C | ter. Terminal Exo | e2/e3-17 | up (e2) | 2,3 | ENSG00000184083 |
| RAP1GAP | Complex | e1-5/e2,e4 | up (e2,e4) | 2,29 | ENSG00000076864 |
| NRP1 | ter. Terminal Exo | 0/e11,e13-1 | down (e11,e13-14) | 2,29 | ENSG00000099250 |
| RIC3 | Exon Cassette | e3,e4 | down | 2,29 | ENSG00000166405 |
| CD9 | Alter. First Exon | e1/e2 | down (e2) | 2,29 | ENSG00000010278 |
| ATF7IP | Alter. First Exon | e1/e2 | up (e1) | 2,29 | ENSG00000171681 |
| HNRNPC | Complex | e3,e4 | up | 2,29 | ENSG00000092199 |
| GPR56 | Complex | e6,e7,e9-16 | up | 2,29 | ENSG00000205336 |
| MAPT | Alter. First Exon | e1-3/e4 | up (e1-3) | 2,29 | ENSG00000186868 |
| NOTCH3 | Exon Cassette | e16 | up | 2,29 | ENSG00000074181 |
| RAB3A | Alter. First Exon | e1/e2 | up (e1) | 2,29 | ENSG00000105649 |
| ZNF772 | Exon Cassette | e4 | up | 2,29 | ENSG00000197128 |
| CAD11 // NPHP | Intron Retention | e44 | up | 2,29 | ENSG00000113971 // ENSG00000240303 |
| HPSE | Complex | e2,e3-4,e7-9 | up | 2,29 | ENSG00000173083 |
| OMMD5 // ZNF25 | Alter. First Exon | 1-5,e7-10/e1 | down (e1-5,e7-10) | 2,29 | ENSG00000170619 // ENSG00000196150 |
| PPIE | Alter. Donor Site | e4 | up | 2,28 | ENSG00000084072 |
| OLFM3 | Alter. First Exon | e1-4/e5 | up (e1-4) | 2,28 | ENSG00000118733 |
| IGSF9 | Exon Cassette | e12 | up | 2,28 | ENSG00000085552 |
| FGD4 | ter. Terminal Exo | e21/e22-23 | up (e21) | 2,28 | ENSG00000139132 |
| DBI | Complex | e1/e2 | up (e2) | 2,28 | ENSG00000155368 |

| | | | | | |
|---|---|---|---|---|---|
| DBI | Exon Cassette | e2 | up | 2,28 | ENSG00000155368 |
| GTPBP10 | Exon Cassette | e3,e4 | up | 2,28 | ENSG00000105793 |
| TMEM2 | Alter. First Exon | e20-22/e23 | down (e20-22) | 2,28 | ENSG00000135048 |
| TCF7L2 | Exon Cassette | e6 | down | 2,27 | ENSG00000148737 |
| TMTC4 | ter. Terminal Exo | 5/e16,e17-1 | up (e15) | 2,27 | ENSG00000125247 |
| LRRC49 | Exon Cassette | e7,e9 | up | 2,27 | ENSG00000137821 |
| ELMOD3 | Exon Cassette | e2,e3 | up | 2,27 | ENSG00000115459 |
| PRKRA | Exon Cassette | e5 | up | 2,27 | ENSG00000180228 |
| BID | Complex | e4-7 | up | 2,27 | ENSG00000015475 |
| HPSE | Complex | e2-9 | up | 2,27 | ENSG00000173083 |
| MATR3 // SNHG4 | Alter. First Exon | e1-2,e4/e9 | up (e1-2,e4) | 2,27 | ENSG00000015479 |
| DUSP22 | Intron Retention | e8 | up | 2,27 | ENSG00000112679 |
| ENY2 | Intron Retention | e1 | up | 2,27 | ENSG00000120533 |
| FAM213B | Intron Retention | e6 | up | 2,26 | ENSG00000157870 |
| LRRC27 | Exon Cassette | e16 | up | 2,26 | ENSG00000148814 |
| SYT9 | Exon Cassette | e4 | down | 2,26 | ENSG00000170743 |
| SDHD | ualy Exclusive Ex | e4/e5 | up (e5) | 2,26 | ENSG00000204370 |
| SDHD | Exon Cassette | e5 | up | 2,26 | ENSG00000204370 |
| DLEU1 | Complex | e30 | down | 2,26 | ENSG00000176124 |
| NBR2 | ter. Terminal Exo | e7/e9 | up (e7) | 2,26 | ENSG00000198496 |
| TMUB2 | Exon Cassette | e3 | up | 2,26 | ENSG00000168591 |
| ZNF780B | Exon Cassette | e3 | up | 2,26 | ENSG00000128000 |
| RAB28 | Exon Cassette | e8 | up | 2,26 | ENSG00000157869 |
| MYB | Complex | e8/e10,e12 | up (e10,e12) | 2,26 | ENSG00000118513 |
| ST7 // ST7-OT3 | ter. Terminal Exo | e25/e26 | up (e25) | 2,26 | ENSG00000004866 |
| ZC4H2 | Alter. First Exon | e1/e2 | up (e1) | 2,26 | ENSG00000126970 |

| | | | | | |
|---|---|---|---|---|---|
| CCDC30 | ualy Exclusive Ex | 9,e10-11,e1 | p (e9,e10-11,e13-14 | 2,25 | ENSG00000 186409 |
| PRKACB | Alter. First Exon | e1/e3,e7 | up (e1) | 2,25 | ENSG00000 142875 |
| PRKACB | Alter. First Exon | e1/e3 | up (e1) | 2,25 | ENSG00000 142875 |
| CACNB2 | ualy Exclusive Ex | e11/e12 | up (e11) | 2,25 | ENSG00000 165995 |
| UBXN1 | Intron Retention | e7 | up | 2,25 | ENSG00000 162191 |
| TMPO | ter. Terminal Exc | e4/e5,e9-10 | up (e4) | 2,25 | ENSG00000 120802 |
| DSC3 | Exon Cassette | e16 | down | 2,25 | ENSG00000 134762 |
| COL3A1 | Alter. First Exon | 1-36,e47/e4 | down (e1-36,e47) | 2,25 | ENSG00000 168542 |
| PCBP3 | ter. Terminal Exc | e11/e12-14 | up (e12-14) | 2,25 | ENSG00000 183570 |
| HLA-C | Complex | e2-3 | up | 2,25 | ENSG00000 204525 |
| KN2B-AS1 // MT | Exon Cassette | e20,e21,e23 | down | 2,25 | ENSG00000 099810 // ENSG00000 240498 |
| TOR2A | Intron Retention | e3 | down | 2,25 | ENSG00000 160404 |
| XRRA1 | Exon Cassette | e10-12 | down | 2,24 | ENSG00000 166435 |
| CCDC91 | Exon Cassette | e7 | up | 2,24 | ENSG00000 123106 |
| EIF4B | Exon Cassette | e7 | up | 2,24 | ENSG00000 063046 |
| TMPO | ter. Terminal Exc | e4/e5,e9-11 | up (e4) | 2,24 | ENSG00000 120802 |
| SRSF5 | Intron Retention | e6-7 | up | 2,24 | ENSG00000 100650 |
| --- | Alter. Donor Site | e2 | up | 2,24 | --- |
| 8 // EIF4A1 // SE | Intron Retention | e22-23 | up | 2,24 | ENSG00000 129226 // ENSG00000 161956 // ENSG00000 161960 |
| TNFRSF11A | Exon Cassette | e9 | up | 2,24 | ENSG00000 141655 |
| TNFRSF11A | Exon Cassette | e9-10 | up | 2,24 | ENSG00000 141655 |
| SNRNP70 | Complex | e8,e9 | up | 2,24 | ENSG00000 104852 |
| CSNK1E | ter. Terminal Exc | e8/e9 | up (e8) | 2,24 | ENSG00000 213923 |

| | | | | | |
|---|---|---|---|---|---|
| DNAL4 // SUN2 | Complex | e12,e13 | down | 2,24 | ENSG00000 100242 // ENSG00000 100246 |
| ZBTB38 | Alter. First Exon | 1-5,e7,e9/e1 | up (e1-5,e7,e9) | 2,24 | ENSG00000 177311 |
| CLASP2 | ter. Terminal Exo | ,e23-24,e27 | up (e20-21) | 2,24 | ENSG00000 163539 |
| PPIP5K2 | Exon Cassette | e28 | up | 2,24 | ENSG00000 145725 |
| WDR55 | Intron Retention | e5 | up | 2,24 | ENSG00000 120314 |
| MYB | Complex | 8/e10,e11-1 | up (e10,e11-12) | 2,24 | ENSG00000 118513 |
| SELENBP1 | lter. Acceptor Sit | e2 | down | 2,23 | ENSG00000 143416 |
| TMEM218 | Complex | e1 | down | 2,23 | ENSG00000 150433 |
| P2RX7 | Exon Cassette | e5 | down | 2,23 | ENSG00000 089041 |
| MYO9A | Exon Cassette | e27 | up | 2,23 | ENSG00000 066933 |
| C16orf45 | Complex | e1,e3 | down | 2,23 | ENSG00000 166780 |
| ZNF559 // ZNF55 | Intron Retention | e7 | up | 2,23 | ENSG00000 188321 // ENSG00000 188629 |
| ZNF559 // ZNF55 | Exon Cassette | e3 | up | 2,23 | ENSG00000 188321 // ENSG00000 188629 |
| SRSF6 | Exon Cassette | e3 | up | 2,23 | ENSG00000 124193 |
| IQGAP2 | Exon Cassette | e20 | down | 2,23 | ENSG00000 145703 |
| ULBP2 | ter. Terminal Exo | e3/e4,e5 | down (e4,e5) | 2,23 | ENSG00000 131015 |
| DDC | Exon Cassette | e4 | up | 2,23 | ENSG00000 132437 |
| BAG1 | Complex | e6-7 | down | 2,23 | ENSG00000 107262 |
| ATP2B3 | ter. Terminal Exo | e20/e21,e23 | down (e21,e23) | 2,23 | ENSG00000 067842 |
| TCF7L2 | Alter. First Exon | e1-3,e5/e8 | down (e1-3,e5) | 2,22 | ENSG00000 148737 |
| TCF7L2 | Alter. First Exon | e1-3,e5/e9 | down (e1-3,e5) | 2,22 | ENSG00000 148737 |
| TIAL1 | Intron Retention | e12 | up | 2,22 | ENSG00000 151923 |
| CAPRIN2 | Exon Cassette | e5 | up | 2,22 | ENSG00000 110888 |

| | | | | | |
|---|---|---|---|---|---|
| DCTN2 | Complex | e2/e4 | up (e4) | 2,22 | ENSG00000 175203 |
| FLT1 | Alter. First Exon | -27,e29-35/ | own (e25-27,e29-35 | 2,22 | ENSG00000 102755 |
| APA1 // RALGA | ter. Terminal Exo | e43/e46 | down (e46) | 2,22 | ENSG00000 174373 // ENSG00000 229419 |
| COL3A1 | Alter. First Exon | e1-47/e48 | down (e1-47) | 2,22 | ENSG00000 168542 |
| KIF1A | Complex | 2,e3-38,e40 | wn (e2,e3-38,e40-4 | 2,22 | ENSG00000 130294 |
| TCEA2 | Intron Retention | e11 | up | 2,22 | ENSG00000 171703 |
| BID | Exon Cassette | e4,e5-6 | up | 2,22 | ENSG00000 015475 |
| KIAA0226 | Intron Retention | e12 | up | 2,22 | ENSG00000 145016 |
| SLC1A3 | Complex | e4-9,e11 | up | 2,22 | ENSG00000 079215 |
| FGF1 | Complex | e8-9 | down | 2,22 | ENSG00000 113578 |
| ATXN1 | Exon Cassette | e3 | up | 2,22 | ENSG00000 124788 |
| KIF13A | Exon Cassette | e27 | up | 2,22 | ENSG00000 137177 |
| ZNF138 | Exon Cassette | e3,e4,e7 | up | 2,22 | ENSG00000 197008 |
| MRRF | Exon Cassette | e6,e7 | up | 2,22 | ENSG00000 148187 |
| ATP2B3 | Exon Cassette | e21 | down | 2,22 | ENSG00000 067842 |
| DMAP1 | Intron Retention | e5,e6-7 | up | 2,21 | ENSG00000 178028 |
| DCP2 // CYB5R | ter. Terminal Exo | e6/e7-10 | down (e7-10) | 2,21 | ENSG00000 157211 // ENSG00000 215883 |
| CADM1 | Exon Cassette | e10,e11 | down | 2,21 | ENSG00000 182985 |
| VPS39 | Complex | e3/e4 | up (e3) | 2,21 | ENSG00000 166887 |
| GPR56 | Complex | e6,e7-16 | up | 2,21 | ENSG00000 205336 |
| MAP4K4 | Exon Cassette | e17 | down | 2,21 | ENSG00000 071054 |
| EPB41L1 | Complex | 2,e10/e3,e | up (e3,e5) | 2,21 | ENSG00000 088367 |
| PPIP5K2 | Exon Cassette | e28,e29-30 | up | 2,21 | ENSG00000 145725 |
| PPIA | Exon Cassette | e2 | up | 2,21 | ENSG00000 196262 |

| | | | | | |
|---|---|---|---|---|---|
| LRRCC1 | Alter. Donor Site | e1 | up | 2,21 | ENSG00000133739 |
| RUNX1T1 | Exon Cassette | e15 | up | 2,21 | ENSG00000079102 |
| BGT1 // RALGD | Intron Retention | e27 | up | 2,21 | ENSG00000148288 // ENSG00000160271 |
| EXTL2 | Exon Cassette | e3 | up | 2,2 | ENSG00000162694 |
| CCT3 | Exon Cassette | e2 | up | 2,2 | ENSG00000163468 |
| DTX3 | Intron Retention | e2 | up | 2,2 | ENSG00000178498 |
| DPP8 | Exon Cassette | e19 | up | 2,2 | ENSG00000074603 |
| LUC7L3 | Intron Retention | e12-13 | up | 2,2 | ENSG00000108848 |
| AMZ2P1 | Intron Retention | e2 | up | 2,2 | ENSG00000214174 |
| --- | Exon Cassette | e5,e6,e9-11 | up | 2,2 | --- |
| SLC8A2 | Exon Cassette | e2 | down | 2,2 | ENSG00000118160 |
| MTA3 | ter. Terminal Exo | 7/e18,e19-2 | up (e17) | 2,2 | ENSG00000057935 |
| SNRPB | Exon Cassette | e3 | up | 2,2 | ENSG00000125835 |
| BCL2L1 | Exon Cassette | e4 | up | 2,2 | ENSG00000171552 |
| BTD | Alter. First Exon | e1/e2 | down (e1) | 2,2 | ENSG00000169814 |
| FAM13A | Alter. First Exon | -9,e11-12/e | up (e1-9,e11-12) | 2,2 | ENSG00000138640 |
| RIMS1 | Alter. First Exon | 25,e28,e30 | e10-22,e25,e28,e3 | 2,2 | ENSG00000079841 |
| RNF146 | Exon Cassette | e4,e6 | up | 2,2 | ENSG00000118518 |
| HLA-DMB | ter. Terminal Exo | e3/e5,e6 | down (e3) | 2,2 | ENSG00000242574 |
| GRIN1 | Exon Cassette | e20 | down | 2,2 | ENSG00000176884 |
| PTER | Exon Cassette | e2 | down | 2,19 | ENSG00000165983 |
| C11orf73 | ter. Terminal Exo | e6-7/e8 | up (e6-7) | 2,19 | ENSG00000149196 |
| XRRA1 | Exon Cassette | e11 | down | 2,19 | ENSG00000166435 |
| IKBIP | Exon Cassette | e2 | up | 2,19 | ENSG00000166130 |
| GPHN | Exon Cassette | e7 | up | 2,19 | ENSG00000171723 |

| | | | | | |
|---|---|---|---|---|---|
| LRP2 | Alter. First Exon | e1-50/e51 | up (e1-50) | 2,19 | ENSG00000 081479 |
| KLHL5 | Exon Cassette | e2 | up | 2,19 | ENSG00000 109790 |
| TRAF3IP2-AS1 | ter. Terminal Exo | e4/e6-7 | up (e4) | 2,19 | ENSG00000 231889 |
| IYMAI // PLAGL1 | Exon Cassette | e4 | up | 2,19 | ENSG00000 118495 |
| ANXA1 | Complex | e1/e2-3 | down (e2-3) | 2,19 | ENSG00000 135046 |
| BGT1 // RALGD | Alter. Donor Site | e27 | up | 2,19 | ENSG00000 148288 // ENSG00000 160271 |
| PRKACB | Alter. First Exon | e1/e4 | up (e1) | 2,18 | ENSG00000 142875 |
| FNBP1L | Intron Retention | e16 | up | 2,18 | ENSG00000 137942 |
| PF10 // NOTCH2 | Complex | 102,e103-10 | down | 2,18 | ENSG00000 163386 // ENSG00000 213240 |
| ZEB1 | Alter. First Exon | e1-2/e3 | up (e1-2) | 2,18 | ENSG00000 148516 |
| TCF7L2 | Alter. First Exon | e2-3,e5/e8 | down (e2-3,e5) | 2,18 | ENSG00000 148737 |
| TCF7L2 | Alter. First Exon | e2-3,e5/e9 | down (e2-3,e5) | 2,18 | ENSG00000 148737 |
| TMEM80 | Intron Retention | e5-6 | up | 2,18 | ENSG00000 177042 |
| ATG16L2 | Complex | e4,e5-6 | down | 2,18 | ENSG00000 168010 |
| MFGE8 | Exon Cassette | e3,e4 | up | 2,18 | ENSG00000 140545 |
| --- | Exon Cassette | e5,e9-11 | up | 2,18 | --- |
| PRNP | Complex | e1-2 | down | 2,18 | ENSG00000 171867 |
| TMEM189-UBE2V | Complex | e11,e15 | up | 2,18 | ENSG00000 124208 // ENSG00000 240849 // ENSG00000 244687 |
| TMEM189-UBE2V | ualy Exclusive Ex | e11/e13 | up (e11) | 2,18 | ENSG00000 124208 // ENSG00000 240849 // ENSG00000 244687 |

| | | | | | |
|---|---|---|---|---|---|
| TMEM189-UBE2V | Exon Cassette | e11 | up | 2,18 | ENSG00000124208 // ENSG00000240849 // ENSG00000244687 |
| FAIM | ualy Exclusive Ex | e3/e4 | up (e3) | 2,18 | ENSG00000158234 |
| CLASP2 | ter. Terminal Exo | e23-24,e27-2 | up (e21) | 2,18 | ENSG00000163539 |
| HPSE | Exon Cassette | e6 | up | 2,18 | ENSG00000173083 |
| TRAF3IP2-AS1 | ter. Terminal Exo | e2-3,e5/e6-7 | up (e2-3,e5) | 2,18 | ENSG00000231889 |
| USP11 | Intron Retention | e19 | up | 2,18 | ENSG00000102226 |
| HAUS7 // TREX2 | ter. Terminal Exo | e8/e9-12 | up (e8) | 2,18 | ENSG00000183479 |
| OSBPL9 | Exon Cassette | e16 | up | 2,17 | ENSG00000117859 |
| SMG7 | Exon Cassette | e19 | down | 2,17 | ENSG00000116698 |
| AKT3 | Complex | e14 | down | 2,17 | ENSG00000117020 |
| SEPHS1 | ter. Terminal Exo | e6/e7,e10 | up (e6) | 2,17 | ENSG00000086475 |
| TMEM218 | Exon Cassette | e2 | up | 2,17 | ENSG00000150433 |
| OSBPL8 | Alter. First Exon | e6-8/e9 | down (e6-8) | 2,17 | ENSG00000091039 |
| ATXN3 | Exon Cassette | e2,e4 | down | 2,17 | ENSG00000066427 |
| ATXN3 | Exon Cassette | e2 | down | 2,17 | ENSG00000066427 |
| ZFAND6 | Exon Cassette | e6 | down | 2,17 | ENSG00000086666 |
| PSMC3IP | Alter. Donor Site | e3 | up | 2,17 | ENSG00000131470 |
| PSMC3IP | Alter. Donor Site | e7 | up | 2,17 | ENSG00000131470 |
| PSMC3IP | Intron Retention | e7 | up | 2,17 | ENSG00000131470 |
| ZNF85 | Alter. First Exon | e1-3/e4,e6 | down (e1-3) | 2,17 | --- |
| KHK | Exon Cassette | e2-3 | up | 2,17 | ENSG00000138030 |
| PDE10A | Complex | e5/e6 | up (e5) | 2,17 | ENSG00000112541 |
| ETV1 | Exon Cassette | e15 | up | 2,17 | ENSG00000006468 |
| NRG1 | Complex | e5/e6-7,e10 | up (e6-7,e10) | 2,17 | ENSG00000157168 |

| | | | | | |
|---|---|---|---|---|---|
| SH3GLB2 | Intron Retention | e8 | up | 2,17 | ENSG00000148341 |
| DDAH1 | Alter. First Exon | e4/e5,e7 | up (e4) | 2,16 | ENSG00000153904 |
| TMED5 | Exon Cassette | e3 | down | 2,16 | ENSG00000117500 |
| RSBN1 | Exon Cassette | e3 | up | 2,16 | ENSG00000081019 |
| BCCIP | ter. Terminal Exo | e6/e8 | up (e6) | 2,16 | ENSG00000107949 |
| TPCN2 | Exon Cassette | e17 | up | 2,16 | ENSG00000162341 |
| FLT1 | Alter. First Exon | e15-16,e18,( | 11,e13,e15-16,e18,( | 2,16 | ENSG00000102755 |
| OXA1L | Intron Retention | e5 | up | 2,16 | ENSG00000155463 |
| EMC4 | Complex | e3/e5 | up (e5) | 2,16 | ENSG00000128463 |
| NEO1 | lter. Acceptor Sit | e8 | up | 2,16 | ENSG00000067141 |
| ZSCAN29 | Alter. First Exon | e1/e2 | down (e1) | 2,16 | ENSG00000140265 |
| LLGL1 | Intron Retention | e17 | up | 2,16 | ENSG00000131899 |
| ANKRD13B | Complex | e15-17 | down | 2,16 | ENSG00000198720 |
| PHF12 | Complex | e3,e4 | up | 2,16 | ENSG00000109118 |
| ABCA5 | Exon Cassette | e38 | up | 2,16 | ENSG00000154265 |
| --- | Exon Cassette | e5-11 | up | 2,16 | --- |
| --- | Exon Cassette | e5,e6-11 | up | 2,16 | --- |
| HPCAL1 | Alter. First Exon | e1/e2 | up (e1) | 2,16 | ENSG00000115756 |
| HPCAL1 | Alter. First Exon | e1/e4 | up (e1) | 2,16 | ENSG00000115756 |
| HPCAL1 | Alter. First Exon | e1/e6 | up (e1) | 2,16 | ENSG00000115756 |
| SNHG17 | Complex | e6-8 | up | 2,16 | ENSG00000196756 |
| ERAP1 | ter. Terminal Exo | e20/e22 | up (e20) | 2,16 | ENSG00000164307 |
| PMPCB | Intron Retention | e12 | up | 2,16 | ENSG00000105819 |
| SAG4 // MAGEA2 | Complex | e2,e3 | up | 2,16 | ENSG00000183305 // ENSG00000242599 |
| ACOT7 | Exon Cassette | e12 | up | 2,15 | ENSG00000097021 |
| SEPHS1 | ter. Terminal Exo | e6/e7-8,e10 | up (e6) | 2,15 | ENSG00000086475 |

| | | | | | |
|---|---|---|---|---|---|
| ZWINT | Intron Retention | e2 | up | 2,15 | ENSG00000 122952 |
| DGKZ | Alter. First Exon | e3-29/e30 | up (e3-29) | 2,15 | ENSG00000 149091 |
| SLC3A2 | Exon Cassette | e9 | down | 2,15 | ENSG00000 168003 |
| ARHGAP32 | Intron Retention | e22 | up | 2,15 | ENSG00000 134909 |
| SYNE2 | Exon Cassette | e118 | down | 2,15 | ENSG00000 054654 |
| ATXN3 | Exon Cassette | e2,e3 | down | 2,15 | ENSG00000 066427 |
| ATXN3 | Exon Cassette | e2,e3-4 | down | 2,15 | ENSG00000 066427 |
| RPAIN | Intron Retention | e4-5 | up | 2,15 | ENSG00000 129197 |
| NELFCD | Intron Retention | e13 | up | 2,15 | ENSG00000 101158 |
| RBM5 // RBM6 | Intron Retention | e30 | up | 2,15 | ENSG00000 003756 // ENSG00000 004534 |
| PCBP4 | Exon Cassette | e3 | up | 2,15 | ENSG00000 090097 |
| OX17 // POPDC | ualy Exclusive E: | e5/e7,e9 | up (e5) | 2,15 | ENSG00000 121577 // ENSG00000 138495 |
| STAG1 | Exon Cassette | e10 | up | 2,15 | ENSG00000 118007 |
| TRIM2 | ter. Terminal Exo | 7/e8-10,e12 | up (e6-7) | 2,15 | ENSG00000 109654 |
| TRIM2 | ter. Terminal Exo | 7/e8-10,e12 | up (e6-7) | 2,15 | ENSG00000 109654 |
| HINT1 | Exon Cassette | e4,e5 | up | 2,15 | ENSG00000 169567 |
| KLHL7 | ter. Terminal Exo | e6/e8-14 | up (e6) | 2,15 | ENSG00000 122550 |
| POLR2J4 | ter. Terminal Exo | /e9-10,e12- | down (e9-10,e12-15 | 2,15 | ENSG00000 214783 |
| KIF13B | Exon Cassette | e40 | up | 2,15 | ENSG00000 197892 |
| PHACTR4 | Exon Cassette | e14 | up | 2,14 | ENSG00000 204138 |
| YIPF1 | Exon Cassette | e12 | down | 2,14 | ENSG00000 058799 |
| DPH5 | Exon Cassette | e3 | up | 2,14 | ENSG00000 117543 |
| TCF7L2 | Alter. First Exon | e1-3/e5 | down (e1-3) | 2,14 | ENSG00000 148737 |
| DNAJC24 | Complex | e4,e5 | up | 2,14 | ENSG00000 170946 |

| | | | | | |
|---|---|---|---|---|---|
| RIC3 | Exon Cassette | e3,e4-5,e7 | down | 2,14 | ENSG00000166405 |
| RAB6A | ualy Exclusive Ex | e5/e6 | up (e5) | 2,14 | ENSG00000175582 |
| OSBPL8 | Alter. First Exon | e1,e5/e6 | up (e1,e5) | 2,14 | ENSG00000091039 |
| C12orf76 | Complex | e8,e9 | up | 2,14 | ENSG00000174456 |
| MCF2L | Complex | e27-31,e34 | down | 2,14 | ENSG00000126217 |
| P1 // NEDD8 // N | Alter. Donor Site | e5 | up | 2,14 | ENSG00000100926 // ENSG00000129559 // ENSG00000196497 // ENSG00000213920 // ENSG00000254505 // ENSG00000255526 |
| TXNDC11 | Complex | e1-2 | up | 2,14 | ENSG00000153066 |
| LEPREL4 | Alter. First Exon | e1/e2 | down (e1) | 2,14 | ENSG00000141696 |
| --- | Exon Cassette | e6,e7-11 | up | 2,14 | --- |
| M228A // FAM22 | Complex | e4/e5 | up (e4) | 2,14 | ENSG00000186453 // ENSG00000219626 |
| MYD88 | Exon Cassette | e3 | down | 2,14 | ENSG00000172936 |
| PHF1 | Intron Retention | e6 | up | 2,14 | ENSG00000112511 |
| IAA1984 // RABL | Alter. First Exon | e3-4/e5 | down (e3-4) | 2,14 | ENSG00000196642 // ENSG00000213213 |
| FAM3A | Exon Cassette | e7 | down | 2,14 | ENSG00000071889 |
| NRP1 | ter. Terminal Exo | e10/e11,e14 | down (e11,e14) | 2,13 | ENSG00000099250 |
| MAP1LC3B | Exon Cassette | e4 | up | 2,13 | ENSG00000140941 |
| MBTD1 | Exon Cassette | e8 | up | 2,13 | ENSG00000011258 |
| Sep-02 | Alter. First Exon | e1-7/e9 | up (e1-7) | 2,13 | ENSG00000168385 |

| | | | | | |
|---|---|---|---|---|---|
| // PRR5 // PRR5 | Alter. First Exon | e4-10/e13 | down (e4-10) | 2,13 | ENSG00000186654 // ENSG00000241484 // ENSG00000248405 |
| REST | ter. Terminal Exo | 5-12/e14-15 | up (e5-12) | 2,13 | ENSG00000084093 |
| REST | ter. Terminal Exo | 5-12/e13,e1 | up (e5-12) | 2,13 | ENSG00000084093 |
| TMEM128 | Intron Retention | e4 | up | 2,13 | ENSG00000132406 |
| HPSE | Complex | e2-5,e7-9 | up | 2,13 | ENSG00000173083 |
| PPIP5K2 | Exon Cassette | e28,e29 | up | 2,13 | ENSG00000145725 |
| IYMAI // PLAGL1 | Complex | e7/e9,e10 | up (e9,e10) | 2,13 | ENSG00000118495 |
| AKAP9 | Exon Cassette | e20 | down | 2,13 | ENSG00000127914 |
| PMS2P4 | ter. Terminal Exo | e5-6/e8 | up (e5-6) | 2,13 | ENSG00000067601 |
| RUNX1T1 | Exon Cassette | e7 | up | 2,13 | ENSG00000079102 |
| ESRRG | Exon Cassette | e12 | up | 2,12 | ENSG00000196482 |
| SLC47A1 | Intron Retention | e10 | down | 2,12 | ENSG00000142494 |
| ZNF254 | Alter. First Exon | 1-2,e4/e9-10 | up (e1-2,e4) | 2,12 | ENSG00000213096 |
| ZNF254 | Alter. First Exon | e1-2,e4/e9 | up (e1-2,e4) | 2,12 | ENSG00000213096 |
| ZNF266 | Intron Retention | e8-9 | up | 2,12 | ENSG00000174652 |
| ITPRIPL1 | Exon Cassette | e2 | up | 2,12 | ENSG00000198885 |
| CASP8 | Exon Cassette | e9 | up | 2,12 | ENSG00000064012 |
| FN1 | Exon Cassette | e41-42 | down | 2,12 | ENSG00000115414 |
| NAPB | ter. Terminal Exo | e10/e11 | up (e10) | 2,12 | ENSG00000125814 |
| GGT7 | Exon Cassette | e3 | down | 2,12 | ENSG00000131067 |
| PREX1 | Intron Retention | e35 | up | 2,12 | ENSG00000124126 |
| MLH1 | Exon Cassette | e17-18 | up | 2,12 | ENSG00000076242 |
| ATP11B | Exon Cassette | e29,e30 | down | 2,12 | ENSG00000058063 |
| OPA1 | Exon Cassette | e7 | up | 2,12 | ENSG00000198836 |

| | | | | | |
|---|---|---|---|---|---|
| TBL1XR1 | Alter. First Exon | e2/e3 | up (e2) | 2,12 | ENSG00000177565 |
| PAM | Exon Cassette | e14 | down | 2,12 | ENSG00000145730 |
| RGL2 | Intron Retention | e6 | up | 2,12 | ENSG00000237441 |
| KLHL7 | ter. Terminal Exo | e7/e8-14 | up (e7) | 2,12 | ENSG00000122550 |
| ZNF138 | Exon Cassette | e2,e3 | up | 2,12 | ENSG00000197008 |
| P1 // TRIM73 // T | Intron Retention | e9 | up | 2,12 | ENSG00000155428 // ENSG00000223705 |
| KN2B-AS1 // MT | Exon Cassette | 8,e20-21,e2 | down | 2,12 | ENSG00000099810 // ENSG00000240498 |
| ARHGEF7 | Exon Cassette | e3-4 | up | 2,11 | ENSG00000102606 |
| ZNRF1 | Exon Cassette | e3 | up | 2,11 | ENSG00000186187 |
| ZNF254 | Alter. First Exon | e1-2/e9-10 | up (e1-2) | 2,11 | ENSG00000213096 |
| ELMOD3 | Exon Cassette | e2,e3-4 | up | 2,11 | ENSG00000115459 |
| NELFCD | Intron Retention | e5 | up | 2,11 | ENSG00000101158 |
| PCED1A | Intron Retention | e4 | up | 2,11 | ENSG00000132635 |
| DGCR8 | Intron Retention | e2 | up | 2,11 | ENSG00000128191 |
| ATXN1 | Complex | e3,e7-9 | down | 2,11 | ENSG00000124788 |
| ETV1 | Alter. First Exon | e4-8/e10 | up (e4-8) | 2,11 | ENSG00000006468 |
| FAM167A | Alter. First Exon | e1/e2,e4 | up (e2,e4) | 2,11 | ENSG00000154319 |
| MSANTD3-TMEF | ter. Terminal Exo | e5-6/e8-16 | up (e5-6) | 2,11 | ENSG00000066697 // ENSG00000241697 // ENSG00000251349 |
| CSTF2 | Exon Cassette | e9,e10 | down | 2,11 | ENSG00000101811 |
| EPB41 | ter. Terminal Exo | /e15-18,e20 | up (e14) | 2,1 | ENSG00000159023 |
| S100PBP | Exon Cassette | e7 | up | 2,1 | ENSG00000116497 |

| | | | | | |
|---|---|---|---|---|---|
| DHX9 // NPL | Exon Cassette | e24 | up | 2,1 | ENSG00000 135829 // ENSG00000 135838 |
| TSEN15 | Exon Cassette | e4,e5 | up | 2,1 | ENSG00000 198860 |
| TCF7L2 | Exon Cassette | e19,e20 | down | 2,1 | ENSG00000 148737 |
| WNK1 | Exon Cassette | e14 | down | 2,1 | ENSG00000 060237 |
| IKBIP | Complex | e1-2 | up | 2,1 | ENSG00000 166130 |
| CRIP2 | Exon Cassette | e5 | down | 2,1 | ENSG00000 182809 |
| MYEF2 | Intron Retention | e10 | up | 2,1 | ENSG00000 104177 |
| CDIPT | Intron Retention | e3 | up | 2,1 | ENSG00000 103502 |
| CCDC40 | Alter. First Exon | e1-7/e8,e9 | up (e1-7) | 2,1 | ENSG00000 141519 |
| ZNF559 // ZNF5 | lter. Acceptor Sit | e8 | down | 2,1 | ENSG00000 188321 // ENSG00000 188629 |
| ZNF254 | Alter. First Exon | e1-2/e9 | up (e1-2) | 2,1 | ENSG00000 213096 |
| ZNF260 | Exon Cassette | e3 | up | 2,1 | ENSG00000 254004 |
| MAP4K4 | Exon Cassette | e17,e18 | down | 2,1 | ENSG00000 071054 |
| C6orf70 | Exon Cassette | e16 | down | 2,1 | ENSG00000 130023 |
| HLA-DOB // TAP | Complex | e8-12 | down | 2,1 | ENSG00000 204267 // ENSG00000 241106 |
| HLA-DMA | Intron Retention | e2,e3 | down | 2,1 | ENSG00000 204257 |
| NRG1 | Exon Cassette | e6,e7 | up | 2,1 | ENSG00000 157168 |
| OXR1 | Alter. First Exon | 1-11,e13/e | up (e1-11,e13) | 2,1 | ENSG00000 164830 |
| SCML1 | Intron Retention | e2-3 | up | 2,1 | ENSG00000 047634 |
| ZNF711 | Exon Cassette | e4 | down | 2,1 | ENSG00000 147180 |
| MPP1 | Exon Cassette | e2 | down | 2,1 | ENSG00000 130830 |
| WASH3P | Alter. First Exon | e1-2/e3 | up (e1-2) | 2,1 | ENSG00000 185596 |
| --- | Alter. First Exon | e1-2/e3 | down (e1-2) | 2,09 | --- |

| | | | | | |
|---|---|---|---|---|---|
| MRPL55 | Alter. Donor Site | e1 | up | 2,09 | ENSG00000162910 |
| VDAC2 | Exon Cassette | e7 | up | 2,09 | ENSG00000165637 |
| HDAC7 | Intron Retention | e25,e26 | up | 2,09 | ENSG00000061273 |
| 1S1 // IL4I1 // NU | Intron Retention | e2,e3 | up | 2,09 | ENSG00000104951 // ENSG00000204673 // ENSG00000213024 |
| ZNF83 | Alter. First Exon | e1-3/e8 | up (e1-3) | 2,09 | ENSG00000167766 |
| PRV1 // PCBP1-A | Alter. First Exon | e1/e2 | down (e1) | 2,09 | ENSG00000179818 // ENSG00000244617 |
| RABL2B | Intron Retention | e2 | up | 2,09 | ENSG00000079974 |
| GUF1 | Exon Cassette | e2 | up | 2,09 | ENSG00000151806 |
| REST | ter. Terminal Exo | 5-11/e14-15 | up (e5-11) | 2,09 | ENSG00000084093 |
| REST | ter. Terminal Exo | 5-11/e13,e1 | up (e5-11) | 2,09 | ENSG00000084093 |
| HAUS3 // POLN | ter. Terminal Exo | 8/e9-21,e23 | down (e7-8) | 2,09 | ENSG00000130997 // ENSG00000214367 |
| STEAP2 | Alter. First Exon | e2/e4 | down (e2) | 2,09 | ENSG00000157214 |
| RMI1 | Complex | e1-2/e3 | up (e1-2) | 2,09 | ENSG00000178966 |
| ALG13 | Exon Cassette | e5 | up | 2,09 | ENSG00000101901 |
| ZYG11B | Exon Cassette | e11 | up | 2,08 | ENSG00000162378 |
| IRF6 | Exon Cassette | e2,e3 | down | 2,08 | ENSG00000117595 |
| CAMK2G | Exon Cassette | e22 | up | 2,08 | ENSG00000148660 |
| RAD52 | ter. Terminal Exo | e5,e6-9,e11 | down (e2) | 2,08 | ENSG00000002016 |
| PARP11 | Exon Cassette | e2 | up | 2,08 | ENSG00000111224 |
| TBX2 | Complex | e1,e2-6 | down | 2,08 | ENSG00000121068 |
| ZNF90 | Exon Cassette | e2 | down | 2,08 | ENSG00000213988 |
| DOCK10 | Exon Cassette | e48 | up | 2,08 | ENSG00000135905 |

| | | | | | |
|---|---|---|---|---|---|
| GOLGA4 | Exon Cassette | e28 | down | 2,08 | ENSG00000144674 |
| IP6K2 | ter. Terminal Exo | 9-10/e12-1 | up (e9-10) | 2,08 | ENSG00000068745 |
| FGFR3 | Complex | e7,e8 | down | 2,08 | ENSG00000068078 |
| AFF1 | Alter. First Exon | e1-3/e4 | up (e1-3) | 2,08 | ENSG00000172493 |
| RASA1 | Exon Cassette | e14 | up | 2,08 | ENSG00000145715 |
| PPAP2A | Complex | e1-2/e3 | down (e1-2) | 2,08 | ENSG00000067113 |
| RNF146 | Exon Cassette | e3,e6 | up | 2,08 | ENSG00000118518 |
| HMBOX1 | Complex | e1,e2 | up | 2,08 | ENSG00000147421 |
| RPL23AP53 | Exon Cassette | e2 | up | 2,08 | ENSG00000223508 |
| BICD2 | Intron Retention | e7 | down | 2,08 | ENSG00000185963 |
| OSBPL9 | Complex | e15/e16 | up (e16) | 2,07 | ENSG00000117859 |
| CAMSAP2 | Exon Cassette | e5 | down | 2,07 | ENSG00000118200 |
| SEPHS1 | Alter. Donor Site | e9 | up | 2,07 | ENSG00000086475 |
| SEPHS1 | Exon Cassette | e9 | up | 2,07 | ENSG00000086475 |
| ARFIP2 | Intron Retention | e3,e4 | up | 2,07 | ENSG00000132254 |
| CS | Exon Cassette | e2 | up | 2,07 | ENSG00000062485 |
| RSRC2 | Alter. Donor Site | e4 | up | 2,07 | ENSG00000111011 |
| RIMBP2 | Exon Cassette | e2 | up | 2,07 | ENSG00000060709 |
| ANKRD10 | Alter. First Exon | e1-3/e4 | down (e1-3) | 2,07 | ENSG00000088448 |
| CCNB1IP1 | Exon Cassette | e6 | up | 2,07 | ENSG00000100814 |
| 2AP // IGHD // IG | Alter. First Exon | e70/e71 | up (e70) | 2,07 | ENSG00000211896 // ENSG00000211898 // ENSG00000213140 |
| SLC12A6 | Exon Cassette | e6 | up | 2,07 | ENSG00000140199 |
| TNRC6A | Exon Cassette | e15 | down | 2,07 | ENSG00000090905 |

| | | | | | |
|---|---|---|---|---|---|
| DNM2 // QTRT1 | Intron Retention | e7 | up | 2,07 | ENSG00000 079805 // ENSG00000 213339 |
| GTPBP3 | Complex | e4-9/e10 | up (e4-9) | 2,07 | ENSG00000 130299 |
| STX10 | Intron Retention | e7 | up | 2,07 | ENSG00000 104915 |
| MLH1 | Exon Cassette | e8 | up | 2,07 | ENSG00000 076242 |
| SLC1A3 | Complex | e4-11 | up | 2,07 | ENSG00000 079215 |
| CPLX2 | Alter. First Exon | e1-2,e4-6/e3 | up (e1-2,e4-6) | 2,07 | ENSG00000 145920 |
| --- | Exon Cassette | e8-9 | up | 2,07 | --- |
| MID1 | Exon Cassette | e3 | up | 2,07 | ENSG00000 101871 |
| Sep-06 | ter. Terminal Exo | e12-13/e17 | up (e12-13) | 2,07 | ENSG00000 125354 |
| CCDC30 | ualy Exclusive Ex | e10,e11,e13 | up (e10,e11,e13-15 | 2,06 | ENSG00000 186409 |
| TEAD1 | Complex | e7 | up | 2,06 | ENSG00000 187079 |
| TEAD1 | Exon Cassette | e7 | up | 2,06 | ENSG00000 187079 |
| TMEM218 | Exon Cassette | e2,e3 | up | 2,06 | ENSG00000 150433 |
| FOXM1 | Exon Cassette | e9 | up | 2,06 | ENSG00000 111206 |
| HDAC7 | Exon Cassette | e26 | up | 2,06 | ENSG00000 061273 |
| SAMD4A | Exon Cassette | e3,e4-6 | up | 2,06 | ENSG00000 020577 |
| 2AP // IGHD // IG | ter. Terminal Exo | 01/e99,e13 | up (e99,e130-133) | 2,06 | ENSG00000 211896 // ENSG00000 211898 // ENSG00000 213140 |
| 2AP // IGHD // IG | ter. Terminal Exo | 99,e130-13 | o (e54,e99,e130-13 | 2,06 | ENSG00000 211896 // ENSG00000 211898 // ENSG00000 213140 |
| TTBK2 | Exon Cassette | e10,e11-12 | up | 2,06 | ENSG00000 128881 |
| NFATC3 | Alter. First Exon | e3/e4 | up (e3) | 2,06 | ENSG00000 072736 |
| NUDT7 | Exon Cassette | e4 | up | 2,06 | ENSG00000 140876 |

| | | | | | |
|---|---|---|---|---|---|
| RPS15A | Complex | e1 | down | 2,06 | ENSG00000 134419 |
| SLC26A6 | Intron Retention | e4 | down | 2,06 | ENSG00000 225697 |
| OCLN | Complex | e4/e6,e8-9 | up (e4) | 2,06 | ENSG00000 197822 |
| CCHCR1 | Complex | e1,e3 | up | 2,06 | ENSG00000 204536 |
| SNK2B // LY6G5 | Intron Retention | e5 | up | 2,06 | ENSG00000 204435 // ENSG00000 240053 |
| CHCHD7 | Exon Cassette | e2 | up | 2,06 | ENSG00000 170791 |
| DAB2IP | Alter. First Exon | e4/e5,e6-7 | up (e5,e6-7) | 2,06 | ENSG00000 136848 |
| SRRM1 | Exon Cassette | e10 | up | 2,05 | ENSG00000 133226 |
| VPS45 | Intron Retention | e7 | up | 2,05 | ENSG00000 136631 |
| HAX1 | Intron Retention | e2 | up | 2,05 | ENSG00000 143575 |
| DUFB8 // SEC31 | Exon Cassette | e32 | down | 2,05 | ENSG00000 075826 // ENSG00000 166136 |
| DNAJC24 | Exon Cassette | e4 | up | 2,05 | ENSG00000 170946 |
| GABARAPL1 | Complex | e1,e2-3 | up | 2,05 | ENSG00000 139112 |
| LIMA1 | Alter. First Exon | e1-3/e4 | up (e1-3) | 2,05 | ENSG00000 050405 |
| PTGR2 // ZNF41( | Alter. Donor Site | e1 | down | 2,05 | ENSG00000 119725 // ENSG00000 140043 |
| PTGR2 // ZNF41( | Complex | e1 | down | 2,05 | ENSG00000 119725 // ENSG00000 140043 |
| P1 // NEDD8 // N | ter. Terminal Exc | 23/e24,e25-! | down (e24,e25-54) | 2,05 | ENSG00000 100926 // ENSG00000 129559 // ENSG00000 196497 // ENSG00000 213920 // ENSG00000 254505 // ENSG00000 255526 |

| | | | | | |
|---|---|---|---|---|---|
| DOC2A | Alter. Donor Site | e11 | up | 2,05 | ENSG00000 149927 |
| MGAT5B | Exon Cassette | e12 | up | 2,05 | ENSG00000 167889 |
| --- | Exon Cassette | e6,e11 | up | 2,05 | --- |
| IOL12 // TRIOBF | Alter. First Exon | e9-14,e16-1 | wn (e1-7,e9-14,e16- | 2,05 | ENSG00000 100106 // ENSG00000 256872 |
| RNF146 | Exon Cassette | e5,e6 | up | 2,05 | ENSG00000 118518 |
| ZNF655 | ter. Terminal Exc | e4-7/e10 | up (e4-7) | 2,05 | ENSG00000 197343 |
| AP1S2 | ter. Terminal Exc | e4,e8/e6 | down (e4,e8) | 2,05 | ENSG00000 182287 |
| DEPDC1 | Exon Cassette | e9 | down | 2,04 | ENSG00000 024526 |
| MRPL55 | Complex | e1-3 | up | 2,04 | ENSG00000 162910 |
| FBXO18 | Alter. First Exon | e4-5/e6 | up (e4-5) | 2,04 | ENSG00000 134452 |
| INPP5F | ter. Terminal Exc | /e11-16,e18 | own (e11-16,e18-22 | 2,04 | ENSG00000 198825 |
| ERP29 | Exon Cassette | e2 | up | 2,04 | ENSG00000 089248 |
| TAOK3 | Alter. First Exon | e1/e2 | down (e1) | 2,04 | ENSG00000 135090 |
| DLEU1 | Exon Cassette | e14 | down | 2,04 | ENSG00000 176124 |
| ARHGEF7 | Exon Cassette | e3 | up | 2,04 | ENSG00000 102606 |
| MYO5C | Exon Cassette | e5 | down | 2,04 | ENSG00000 128833 |
| COG4 | Intron Retention | e7,e8 | up | 2,04 | ENSG00000 103051 |
| SS18 | Exon Cassette | e3,e5 | up | 2,04 | ENSG00000 141380 |
| SLC9A8 | lter. Acceptor Sit | e7 | down | 2,04 | ENSG00000 197818 |
| USP25 | Exon Cassette | e20 | up | 2,04 | ENSG00000 155313 |
| --- | Exon Cassette | e2 | up | 2,04 | --- |
| CLASP2 | ter. Terminal Exc | ,e23-24,e27 | up (e20-21) | 2,04 | ENSG00000 163539 |
| LSAMP | ter. Terminal Exc | e2/e3-8 | down (e3-8) | 2,04 | ENSG00000 185565 |
| :OX17 // POPDC | Exon Cassette | e8-9 | down | 2,04 | ENSG00000 121577 // ENSG00000 138495 |
| TRIM2 | Alter. First Exon | e3/e4-5 | down (e4-5) | 2,04 | ENSG00000 109654 |

| | | | | | |
|---|---|---|---|---|---|
| RIMS1 | Alter. First Exon | 22,e25,e28,( | (e8,e10-22,e25,e28 | 2,04 | ENSG00000079841 |
| RIMS1 | Alter. First Exon | ,e25,e28,e3 | vn (e9-22,e25,e28,e | 2,04 | ENSG00000079841 |
| UPP1 | lter. Acceptor Sit | e3 | up | 2,04 | ENSG00000183696 |
| CAMK2B | Exon Cassette | e15 | up | 2,04 | ENSG00000058404 |
| RRAGB | Exon Cassette | e4 | down | 2,04 | ENSG00000083750 |
| RCC1 // SNHG3 | Exon Cassette | e5 | down | 2,03 | ENSG00000180198 // ENSG00000242125 |
| NFASC | Exon Cassette | e30,e31-33 | up | 2,03 | ENSG00000163531 |
| STMN1 | Intron Retention | e5 | up | 2,03 | ENSG00000117632 |
| CYP2R1 | Alter. First Exon | e2/e3 | down (e2) | 2,03 | ENSG00000186104 |
| CTSC | Alter. Donor Site | e5 | up | 2,03 | ENSG00000109861 |
| SLC37A4 | Alter. First Exon | e1/e3 | down (e1) | 2,03 | --- |
| RPH3A | Alter. First Exon | e2-10/e11 | down (e2-10) | 2,03 | ENSG00000089169 |
| C14orf93 | Complex | e1/e2-3 | up (e2-3) | 2,03 | ENSG00000100802 |
| 5orf57 // MRPL42 | Alter. Donor Site | e1 | up | 2,03 | ENSG00000128891 |
| ERBB2 | Alter. First Exon | e1-5,e8/e9 | down (e1-5,e8) | 2,03 | ENSG00000141736 |
| ZNF271 | Complex | e1,e2 | up | 2,03 | ENSG00000257267 |
| ZNF615 | Exon Cassette | e2 | up | 2,03 | ENSG00000197619 |
| PREPL | Exon Cassette | e2 | up | 2,03 | ENSG00000138078 |
| FAM136A | Complex | e2 | up | 2,03 | ENSG00000035141 |
| FKBP7 | Complex | e3-4 | down | 2,03 | ENSG00000079150 |
| BID | Exon Cassette | e4-5 | up | 2,03 | ENSG00000015475 |
| CD200 | Exon Cassette | e2-3 | down | 2,03 | ENSG00000091972 |
| SLC30A5 | lter. Acceptor Sit | e4 | down | 2,03 | ENSG00000145740 |
| CLK4 | Intron Retention | e7 | up | 2,03 | ENSG00000113240 |

| | | | | | |
|---|---|---|---|---|---|
| GAL3ST4 // GPC | Complex | e12,e13 | down | 2,03 | ENSG00000 197093 // ENSG00000 213420 |
| IL7 | Exon Cassette | e6 | up | 2,03 | ENSG00000 104432 |
| TRMT1L | Exon Cassette | e11 | up | 2,02 | ENSG00000 121486 |
| NRP1 | ter. Terminal Exo | e10/e11-12 | down (e11-12) | 2,02 | ENSG00000 099250 |
| SLC12A6 | Alter. First Exon | e3/e5,e6 | down (e3) | 2,02 | ENSG00000 140199 |
| SYNRG | Exon Cassette | e23 | up | 2,02 | ENSG00000 006114 |
| NSF // NSFP1 | Complex | e3-4 | up | 2,02 | ENSG00000 073969 // ENSG00000 260075 |
| ZNF583 | Alter. First Exon | e1-3/e4 | up (e1-3) | 2,02 | ENSG00000 198440 |
| ZNF571 | Exon Cassette | e2 | up | 2,02 | ENSG00000 180479 |
| ELMOD3 | Exon Cassette | e2 | up | 2,02 | ENSG00000 115459 |
| CDCA7 | Exon Cassette | e2,e3 | down | 2,02 | ENSG00000 144354 |
| PITX2 | Exon Cassette | e5 | up | 2,02 | ENSG00000 164093 |
| AHRR // PDCD6 | Exon Cassette | e16 | down | 2,02 | ENSG00000 063438 // ENSG00000 249915 |
| WDR55 | Intron Retention | e2 | up | 2,02 | ENSG00000 120314 |
| RIMS1 | Alter. First Exon | 0-22,e25,e3 | n (e7,e10-22,e25,e | 2,02 | ENSG00000 079841 |
| C7orf55-LUC7L2 | Alter. First Exon | e2,e4/e8 | down (e2,e4) | 2,02 | ENSG00000 146963 // ENSG00000 164898 // ENSG00000 269955 |
| RIMS2 | Exon Cassette | e24,e25-29 | down | 2,02 | ENSG00000 176406 |
| DNM1 | ualy Exclusive Ex | e10/e11 | down (e10) | 2,02 | ENSG00000 106976 |
| KIAA0020 | ter. Terminal Exo | e1-2/e3-19 | down (e3-19) | 2,02 | ENSG00000 080608 |
| TRO | Alter. Donor Site | e1 | up | 2,02 | ENSG00000 067445 |
| GDI1 | Intron Retention | e5 | up | 2,02 | ENSG00000 203879 |

| | | | | | |
|---|---|---|---|---|---|
| MID1 | Exon Cassette | e14,e15 | down | 2,02 | ENSG00000 101871 |
| PPIE | Intron Retention | e4 | up | 2,01 | ENSG00000 084072 |
| GAP11 // FAM25 | Intron Retention | e8 | down | 2,01 | ENSG00000 151303 // ENSG00000 188100 |
| NT5C2 | Exon Cassette | e12 | up | 2,01 | ENSG00000 076685 |
| ATG16L2 | Exon Cassette | e4-6 | down | 2,01 | ENSG00000 168010 |
| VPS11 | Complex | e1,e2 | up | 2,01 | --- |
| RNH1 | Complex | e3-4 | up | 2,01 | ENSG00000 023191 |
| UBXN1 | Intron Retention | e1 | up | 2,01 | ENSG00000 162191 |
| TMEM218 | Exon Cassette | e3 | up | 2,01 | ENSG00000 150433 |
| ROGDI | Intron Retention | e4 | up | 2,01 | ENSG00000 067836 |
| ZNF821 | Exon Cassette | e3 | up | 2,01 | ENSG00000 102984 |
| C17orf76-AS1 | Intron Retention | e3 | up | 2,01 | ENSG00000 175061 |
| ZNF606 | ter. Terminal Exo | e5/e6,e7-9 | up (e5) | 2,01 | ENSG00000 166704 |
| LSM14B | Exon Cassette | e6 | up | 2,01 | ENSG00000 149657 |
| EPHA3 | ter. Terminal Exo | e7-8/e9-19 | up (e9-19) | 2,01 | ENSG00000 044524 |
| EPHA3 | ter. Terminal Exo | e7/e9-19 | up (e9-19) | 2,01 | ENSG00000 044524 |
| EPHA3 | ter. Terminal Exo | 7-8/e9,e10-1 | up (e9,e10-19) | 2,01 | ENSG00000 044524 |
| HNRNPDL | Exon Cassette | e8 | up | 2,01 | ENSG00000 152795 |
| ZSCAN26 | Intron Retention | e3 | up | 2,01 | --- |
| CAMK2B | Exon Cassette | e15-16 | up | 2,01 | ENSG00000 058404 |
| LGALS8 | Exon Cassette | e11 | up | 2 | ENSG00000 116977 |
| HYI | Intron Retention | e3,e4 | up | 2 | ENSG00000 178922 |
| NT5C2 | Intron Retention | e14 | up | 2 | ENSG00000 076685 |
| SIPA1 | ter. Terminal Exo | 11,e13-19/e | down (e9-11,e13-19 | 2 | ENSG00000 213445 |
| ARFIP2 | lter. Acceptor Sit | e4 | up | 2 | ENSG00000 132254 |
| SLC37A4 | Exon Cassette | e8 | down | 2 | --- |

| | | | | | |
|---|---|---|---|---|---|
| PML | ter. Terminal Exo | e8/e9,e10 | up (e8) | 2 | ENSG00000140464 |
| IREB2 | Intron Retention | e8 | up | 2 | ENSG00000136381 |
| C16orf59 | Intron Retention | e2 | up | 2 | ENSG00000162062 |
| TVP23C // TVP2 | ter. Terminal Exo | e6/e10-11 | up (e6) | 2 | ENSG00000175106 // ENSG00000239704 // ENSG00000259024 |
| RPL28 | ter. Terminal Exo | e3/e5-7 | up (e3) | 2 | ENSG00000108107 |
| PLEKHH2 | Alter. First Exon | e1-9/e10 | down (e1-9) | 2 | ENSG00000152527 |
| GNAS | Alter. First Exon | e8/e9 | down (e8) | 2 | ENSG00000087460 |
| CCDC14 | Intron Retention | e3 | up | 2 | ENSG00000175455 |
| RNF146 | Exon Cassette | e6 | up | 2 | ENSG00000118518 |
| HLA-DOB // TAP2 | Alter. First Exon | e1-11/e15 | up (e1-11) | 2 | ENSG00000204267 // ENSG00000241106 |
| FBXL6 | Intron Retention | e3 | up | 2 | ENSG00000182325 |
| CCDC180 | Alter. First Exon | 12,e14-15/e | up (e1-12,e14-15) | 2 | ENSG00000197816 |
| KIAA0020 | ter. Terminal Exo | 1-2/e3,e4-1 | down (e3,e4-19) | 2 | ENSG00000080608 |
| LPAR1 | Alter. First Exon | e2/e6,e7-8 | up (e2) | 2 | ENSG00000198121 |
| NFASC | ter. Terminal Exo | e22,e23-30 | vn (e22,e23-30,e34- | 1,99 | ENSG00000163531 |
| MOB2 | Alter. First Exon | e1/e2 | up (e1) | 1,99 | ENSG00000182208 |
| FBXO3 | ter. Terminal Exo | e11/e12 | up (e11) | 1,99 | ENSG00000110429 |
| NUMA1 | Alter. First Exon | e1-3,e5/e7 | up (e1-3,e5) | 1,99 | ENSG00000137497 |
| RASA3 | Exon Cassette | e4,e5 | up | 1,99 | ENSG00000185989 |
| HNRNPC | Exon Cassette | e4 | up | 1,99 | ENSG00000092199 |
| RORA | Complex | e1/e2 | down (e2) | 1,99 | ENSG00000069667 |
| C1D26 // ZNF28 | Alter. First Exon | e1-6/e8 | down (e1-6) | 1,99 | ENSG00000214946 // ENSG00000255104 |

| | | | | | |
|---|---|---|---|---|---|
| SYT4 | Complex | e2-4 | down | 1,99 | ENSG00000132872 |
| RPL28 | ter. Terminal Exo | e3/e5-6 | up (e3) | 1,99 | ENSG00000108107 |
| ZNF606 | ter. Terminal Exo | e5/e6-8 | up (e5) | 1,99 | ENSG00000166704 |
| BRE | Exon Cassette | e2 | up | 1,99 | ENSG00000158019 |
| FKBP7 | Exon Cassette | e3 | down | 1,99 | ENSG00000079150 |
| AAR2 | Complex | e1 | down | 1,99 | ENSG00000131043 |
| TBC1D20 | Intron Retention | e4 | up | 1,99 | ENSG00000125875 |
| MYD88 | Exon Cassette | e2-3 | down | 1,99 | ENSG00000172936 |
| REST | ualy Exclusive Ex | e5/e14 | up (e5) | 1,99 | ENSG00000084093 |
| REST | ualy Exclusive Ex | e5/e13 | up (e5) | 1,99 | ENSG00000084093 |
| HAND2-AS1 | Complex | e4,e5 | up | 1,99 | ENSG00000237125 |
| CCDC180 | Alter. First Exon | e1-12/e17 | up (e1-12) | 1,99 | ENSG00000197816 |
| GDI1 | Intron Retention | e6 | up | 1,99 | ENSG00000203879 |
| NFASC | ter. Terminal Exo | e21,e22-30 | vn (e21,e22-30,e34- | 1,98 | ENSG00000163531 |
| DUFB8 // SEC31 | Complex | e32,e33 | down | 1,98 | ENSG00000075826 // ENSG00000166136 |
| CADM1 | Alter. First Exon | e1,e3-9/e10 | up (e1,e3-9) | 1,98 | ENSG00000182985 |
| ATXN3 | Exon Cassette | e3 | down | 1,98 | ENSG00000066427 |
| ATXN3 | Exon Cassette | e3,e4 | down | 1,98 | ENSG00000066427 |
| MED24 | Exon Cassette | e7 | down | 1,98 | ENSG00000008838 |
| C17orf58 | Alter. Donor Site | e2 | down | 1,98 | ENSG00000186665 |
| EPOR // RGL3 | Complex | e5-7 | up | 1,98 | ENSG00000187266 // ENSG00000205517 |
| OSBPL6 | Exon Cassette | e12,e13 | down | 1,98 | ENSG00000079156 |
| NABP1 | Exon Cassette | e5 | down | 1,98 | ENSG00000173559 |

| | | | | | |
|---|---|---|---|---|---|
| CLK1 // PPIL3 | Intron Retention | e11,e12 | up | 1,98 | ENSG00000013441 // ENSG00000240344 |
| PLCB1 | Alter. First Exon | e1/e2,e4,e7 | down (e2,e4,e7) | 1,98 | ENSG00000182621 |
| C1QTNF6 | Intron Retention | e7 | down | 1,98 | ENSG00000133466 |
| RBM5 // RBM6 | Intron Retention | e29 | up | 1,98 | ENSG00000003756 // ENSG00000004534 |
| CLASP2 | ter. Terminal Exo | e23-24,e27-3 | up (e21) | 1,98 | ENSG00000163539 |
| DLG1 | Exon Cassette | e9 | up | 1,98 | ENSG00000075711 |
| NUP54 | Exon Cassette | e2,e3 | up | 1,98 | ENSG00000138750 |
| MAP3K1 | Intron Retention | e18 | up | 1,98 | ENSG00000095015 |
| PHF15 | ter. Terminal Exo | e11/e12-13 | up (e12-13) | 1,98 | ENSG00000043143 |
| NADK2 | Exon Cassette | e11 | up | 1,98 | ENSG00000152620 |
| MCC | ter. Terminal Exo | e19/e20-22 | up (e19) | 1,98 | ENSG00000171444 |
| APBB3 | Intron Retention | e1 | up | 1,98 | ENSG00000113108 |
| ZNF138 | Exon Cassette | e3-4 | up | 1,98 | ENSG00000197008 |
| ZNF655 | ter. Terminal Exo | e4-7/e9-10 | up (e4-7) | 1,98 | ENSG00000197343 |
| MTUS1 | ter. Terminal Exo | 8-9,e11,e13 | wn (e8-9,e11,e13-2 | 1,98 | ENSG00000129422 |
| USP11 | Alter. Donor Site | e1 | up | 1,98 | ENSG00000102226 |
| DHX9 // NPL | Alter. First Exon | -12,e14-15/e | up (e9-12,e14-15) | 1,97 | ENSG00000135829 // ENSG00000135838 |
| BSDC1 | Intron Retention | e11 | up | 1,97 | ENSG00000160058 |
| ZEB1 | Alter. First Exon | e2/e3 | up (e2) | 1,97 | ENSG00000148516 |
| SLC15A4 | Exon Cassette | e3 | up | 1,97 | ENSG00000139370 |
| BAZ1A | Exon Cassette | e13 | up | 1,97 | ENSG00000198604 |
| PML | ter. Terminal Exo | e8/e9-10 | up (e8) | 1,97 | ENSG00000140464 |
| C17orf85 | Complex | e7,e8 | up | 1,97 | ENSG00000074356 |

| | | | | | |
|---|---|---|---|---|---|
| C18orf54 | Complex | e3 | down | 1,97 | ENSG00000166845 |
| GTPBP3 | Alter. First Exon | e1/e3,e4 | down (e3,e4) | 1,97 | ENSG00000130299 |
| PVR | Alter. Donor Site | e6 | up | 1,97 | ENSG00000073008 |
| RAPGEF4 | Complex | e19-21 | up | 1,97 | ENSG00000091428 |
| DNAJC27 | Exon Cassette | e5 | up | 1,97 | ENSG00000115137 |
| PLCB1 | Alter. First Exon | e2,e4/e6 | down (e2,e4) | 1,97 | ENSG00000182621 |
| --- | Complex | e1-2 | up | 1,97 | --- |
| LSM14B | ter. Terminal Ex( | 6/e7,e8-9,e1 | up (e6) | 1,97 | ENSG00000149657 |
| TMEM230 | Complex | e1 | down | 1,97 | ENSG00000089063 |
| --- | Exon Cassette | e2,e3 | up | 1,97 | --- |
| MAPKAPK3 | Alter. First Exon | e1-3/e5 | down (e1-3) | 1,97 | ENSG00000114738 |
| RNF44 | Intron Retention | e11 | up | 1,97 | ENSG00000146083 |
| NCOA7 | Alter. First Exon | 3,e5-14/e1 | down (e3,e5-14) | 1,97 | ENSG00000111912 |
| OPRM1 | Exon Cassette | e8-9 | down | 1,97 | ENSG00000112038 |
| CUL7 | Intron Retention | e10,e11 | up | 1,97 | ENSG00000044090 |
| HYMAI // PLAGL1 | Exon Cassette | e9 | up | 1,97 | ENSG00000118495 |
| FOXP2 | Exon Cassette | e18-20 | down | 1,97 | ENSG00000128573 |
| TRIQK | Exon Cassette | e6 | up | 1,97 | ENSG00000205133 |
| IAA1984 // RABL | Alter. First Exon | e1-4/e5 | down (e1-4) | 1,97 | ENSG00000196642 // ENSG00000213213 |
| DCTN3 | Intron Retention | e4 | up | 1,97 | ENSG00000137100 |
| DDI2 // RSC1A1 | ter. Terminal Ex( | e5/e7,e8-11 | up (e5) | 1,96 | ENSG00000197312 // ENSG00000215695 |
| PIP5K1A | Exon Cassette | e4 | up | 1,96 | ENSG00000143398 |
| BBIP1 | ter. Terminal Ex( | e4/e5-6 | up (e4) | 1,96 | ENSG00000214413 |
| PRDM11 | ter. Terminal Ex( | e9/e10 | up (e9) | 1,96 | ENSG00000019485 |
| DGKZ | Alter. First Exon | e4-29/e30 | up (e4-29) | 1,96 | ENSG00000149091 |

| | | | | | |
|---|---|---|---|---|---|
| UVRAG | Exon Cassette | e15 | up | 1,96 | ENSG00000198382 |
| ARID2 | Exon Cassette | e23 | up | 1,96 | ENSG00000189079 |
| PPP2R5C | ualy Exclusive Ex | e3/e5 | down (e3) | 1,96 | ENSG00000078304 |
| PML | ter. Terminal Exo | e7/e9-10 | down (e9-10) | 1,96 | ENSG00000140464 |
| PSMA4 | Intron Retention | e3 | up | 1,96 | ENSG00000041357 |
| NFAT5 | Exon Cassette | e4,e5 | up | 1,96 | ENSG00000102908 |
| CYTH1 | Exon Cassette | e16 | up | 1,96 | ENSG00000108669 |
| --- | Exon Cassette | e6-8,e10-11 | up | 1,96 | --- |
| PRMT1 | Alter. First Exon | e1-3/e4 | up (e1-3) | 1,96 | ENSG00000126457 |
| DPP10 | Alter. First Exon | e1-6/e4 | down (e1-6) | 1,96 | ENSG00000175497 |
| MLH1 | Complex | e1-3 | down | 1,96 | ENSG00000076242 |
| MCC | ter. Terminal Exo | e19/e20-23 | up (e19) | 1,96 | ENSG00000171444 |
| MCC | ter. Terminal Exo | 9/e20,e21-2 | up (e19) | 1,96 | ENSG00000171444 |
| BTN2A3P | lter. Acceptor Sit | e8 | up | 1,96 | ENSG00000124549 |
| HMGA1 | Complex | e4-5 | down | 1,96 | ENSG00000137309 |
| NCOA7 | Alter. First Exon | ,e5,e7-14/e | down (e3,e5,e7-14) | 1,96 | ENSG00000111912 |
| WDR46 | Complex | e1,e2 | up | 1,96 | ENSG00000227057 |
| VPS28 | Intron Retention | e5 | up | 1,96 | --- |
| GKAP1 | Exon Cassette | e7 | up | 1,96 | ENSG00000165113 |
| USP21 | Intron Retention | e11 | up | 1,95 | ENSG00000143258 |
| 1A // CDK11B // | Exon Cassette | e15 | down | 1,95 | ENSG00000008128 // ENSG00000078369 // ENSG00000248333 |
| NES | Complex | e1-4 | down | 1,95 | ENSG00000132688 |
| STT3A | Exon Cassette | e2 | up | 1,95 | ENSG00000134910 |
| H19 | Complex | e5/e6 | down (e5) | 1,95 | ENSG00000130600 |

| | | | | | |
|---|---|---|---|---|---|
| / FXYD6 // FXYD | Exon Cassette | e3-4 | up | 1,95 | ENSG00000 137726 // ENSG00000 137731 // ENSG00000 255245 |
| DYRK4 | Exon Cassette | e13 | down | 1,95 | ENSG00000 010219 |
| RIC8B | Exon Cassette | e11 | up | 1,95 | ENSG00000 111785 |
| RIC8B | Exon Cassette | e12,e13-15 | up | 1,95 | ENSG00000 111785 |
| PML | ter. Terminal Exo | e7-8/e9,e10 | up (e7-8) | 1,95 | ENSG00000 140464 |
| UBE2I | Exon Cassette | e3 | up | 1,95 | ENSG00000 103275 |
| --- | ter. Terminal Exo | 3/e14,e15-2 | up (e14,e15-28) | 1,95 | --- |
| BOLA3-AS1 | Complex | e1-2 | down | 1,95 | ENSG00000 225439 |
| BCL2L1 | Complex | e3,e4 | up | 1,95 | ENSG00000 171552 |
| SNHG17 | Intron Retention | e7-8 | up | 1,95 | ENSG00000 196756 |
| UBE2G2 | Alter. First Exon | e1,e5/e6 | down (e1,e5) | 1,95 | ENSG00000 184787 |
| DAC10 // MAPK1 | ter. Terminal Exo | 3/e14,e15-3 | up (e13) | 1,95 | ENSG00000 100429 // ENSG00000 188130 |
| WDR6 | lter. Acceptor Sit | e4 | up | 1,95 | ENSG00000 178252 |
| SLC9B2 | ter. Terminal Exo | e13/e14 | up (e13) | 1,95 | ENSG00000 164038 |
| OPRM1 | Complex | e1,e3 | up | 1,95 | ENSG00000 112038 |
| NAA38 | Intron Retention | e3 | up | 1,95 | ENSG00000 128534 |
| PDGFA | Alter. First Exon | e1-2/e3 | down (e1-2) | 1,95 | ENSG00000 197461 |
| RIMS2 | Exon Cassette | 24,e25-27,e2 | down | 1,95 | ENSG00000 176406 |
| EIF3E | Alter. First Exon | e1/e2 | down (e1) | 1,95 | ENSG00000 104408 |
| SEMA4D | Alter. First Exon | 1-2,e4-15/e2 | up (e1-2,e4-15) | 1,95 | ENSG00000 187764 |
| SEMA4D | Alter. First Exon | e1-15/e20 | up (e1-15) | 1,95 | ENSG00000 187764 |
| FBXW2 | Alter. Donor Site | e2 | up | 1,95 | ENSG00000 119402 |
| HS2ST1 | Alter. First Exon | e2-7/e10 | down (e2-7) | 1,94 | ENSG00000 153936 |

| | | | | | |
|---|---|---|---|---|---|
| TPAF1 // EFCAB | Exon Cassette | e7 | down | 1,94 | ENSG00000123472 // ENSG00000159658 |
| CD59 | Alter. First Exon | e1-3,e6/e7 | down (e1-3,e6) | 1,94 | ENSG00000085063 |
| AHNAK | ter. Terminal Exo | e5-7/e8-9 | up (e5-7) | 1,94 | ENSG00000124942 |
| CWC15 | Intron Retention | e3 | up | 1,94 | --- |
| EXOSC8 | lter. Acceptor Sit | e6 | down | 1,94 | ENSG00000120699 |
| SRSF5 | lter. Acceptor Sit | e7 | up | 1,94 | ENSG00000100650 |
| PSMA4 | Intron Retention | e1 | up | 1,94 | ENSG00000041357 |
| ACD | Intron Retention | e5 | up | 1,94 | ENSG00000102977 |
| CCDC40 | ter. Terminal Exo | e10/e11,e13 | up (e11,e13) | 1,94 | ENSG00000141519 |
| LTBP4 | Alter. First Exon | e1-4/e5 | down (e1-4) | 1,94 | ENSG00000090006 |
| GIPC1 | Exon Cassette | e2-3 | up | 1,94 | ENSG00000123159 |
| MEIS1 | Intron Retention | e13 | up | 1,94 | ENSG00000143995 |
| GGT7 | Alter. First Exon | e1-3/e4 | down (e1-3) | 1,94 | ENSG00000131067 |
| TMEM189-UBE2V | Alter. First Exon | e3,e5-7,e15/ | up (e1,e3,e5-7,e15) | 1,94 | ENSG00000124208 // ENSG00000240849 // ENSG00000244687 |
| RBX1 // XPNPEP | Exon Cassette | e20 | up | 1,94 | ENSG00000100387 // ENSG00000196236 |
| ETFDH | Exon Cassette | e3 | up | 1,94 | ENSG00000171503 |
| ADAT2 | Alter. First Exon | e1/e2 | down (e1) | 1,94 | ENSG00000189007 |
| EIF3E | ter. Terminal Exo | e6/e7-14 | up (e6) | 1,94 | ENSG00000104408 |
| SH3GLB2 | Intron Retention | e7,e8 | up | 1,94 | ENSG00000148341 |
| PPIE | Exon Cassette | e11 | up | 1,93 | ENSG00000084072 |
| TRIM46 | Exon Cassette | e7 | down | 1,93 | ENSG00000163462 |
| CTNNBIP1 | ter. Terminal Exo | e3/e4-7 | down (e4-7) | 1,93 | ENSG00000178585 |

| | | | | | |
|---|---|---|---|---|---|
| TCTN3 | Intron Retention | e1 | down | 1,93 | ENSG00000 119977 |
| XRRA1 | Exon Cassette | e10-11 | down | 1,93 | ENSG00000 166435 |
| ZBTB1 | Exon Cassette | e3 | up | 1,93 | ENSG00000 126804 |
| AREL1 | Exon Cassette | e2 | up | 1,93 | ENSG00000 119682 |
| C16orf93 | ter. Terminal Exo | e3/e4-9 | down (e4-9) | 1,93 | ENSG00000 196118 |
| SLC44A2 | Exon Cassette | e23 | down | 1,93 | ENSG00000 129353 |
| PKP4 | ter. Terminal Exo | /e18-20,e22 | up (e17) | 1,93 | ENSG00000 144283 |
| CLK1 // PPIL3 | lter. Acceptor Sit | e12 | up | 1,93 | ENSG00000 013441 // ENSG00000 240344 |
| GDAP1L1 | Exon Cassette | e3 | down | 1,93 | ENSG00000 124194 |
| IP6K2 | ter. Terminal Exo | e9-11/e12-15 | up (e9-11) | 1,93 | ENSG00000 068745 |
| CCDC14 | Intron Retention | e4 | up | 1,93 | ENSG00000 175455 |
| FAM114A1 | Complex | e2-6 | up | 1,93 | ENSG00000 197712 |
| TRAF3IP2 | Exon Cassette | e2 | down | 1,93 | ENSG00000 056972 |
| NUDT1 | Alter. First Exon | e1-4/e2-3 | up (e2-3) | 1,93 | ENSG00000 106268 |
| HNRNPA2B1 | Intron Retention | e11 | up | 1,93 | ENSG00000 122566 |
| AASS | Exon Cassette | e2-11 | up | 1,93 | ENSG00000 008311 |
| ZNF767 | Exon Cassette | e3 | up | 1,93 | ENSG00000 133624 |
| NRG1 | Complex | e5/e6,e7,e10 | up (e6,e7,e10) | 1,93 | ENSG00000 157168 |
| --- | Exon Cassette | e5 | down | 1,92 | --- |
| CDC42 | Exon Cassette | e5 | up | 1,92 | ENSG00000 070831 |
| TCEANC2 | Exon Cassette | e3 | down | 1,92 | ENSG00000 116205 |
| RHGAP19 // SLI | ter. Terminal Exo | 21,e22-27,e | (e21,e22-27,e29-5 | 1,92 | ENSG00000 187122 // ENSG00000 213390 |
| CADM1 | ualy Exclusive Ex | e9/e10,e11 | up (e9) | 1,92 | ENSG00000 182985 |
| TMEM218 | Complex | e1/e2 | down (e1) | 1,92 | ENSG00000 150433 |

| | | | | | |
|---|---|---|---|---|---|
| BICD1 | Complex | e7,e8 | up | 1,92 | ENSG00000151746 |
| BICD1 | Exon Cassette | e8 | up | 1,92 | ENSG00000151746 |
| LETMD1 | Exon Cassette | e4-7 | up | 1,92 | ENSG00000050426 |
| 2AP // IGHD // IG | Alter. First Exon | e18-19/e103 | down (e18-19) | 1,92 | ENSG00000211896 // ENSG00000211898 // ENSG00000213140 |
| ERBB2 | Alter. First Exon | e1-4,e8/e9 | down (e1-4,e8) | 1,92 | ENSG00000141736 |
| DDX52 | Intron Retention | e14 | up | 1,92 | ENSG00000141141 |
| ZNF320 | ter. Terminal Exo | e10/e11,e12 | down (e10) | 1,92 | ENSG00000182986 |
| CHMP2A | Complex | e1 | down | 1,92 | ENSG00000130724 |
| ZFAND2B | ter. Terminal Exo | e8/e9-10 | up (e8) | 1,92 | ENSG00000158552 |
| TTN | Exon Cassette | e138-143 | down | 1,92 | ENSG00000155657 |
| PANX2 | Exon Cassette | e2 | up | 1,92 | ENSG00000073150 |
| NKTR | Alter. First Exon | e1-7/e8 | down (e1-7) | 1,92 | ENSG00000114857 |
| ATP2B2 | Complex | e14-19 | down | 1,92 | ENSG00000157087 |
| STAG1 | ter. Terminal Exo | 0/e11,e12-3 | up (e10) | 1,92 | ENSG00000118007 |
| STAG1 | ter. Terminal Exo | 11,e12-33,e | up (e10) | 1,92 | ENSG00000118007 |
| TRIM41 | Intron Retention | e3 | up | 1,92 | ENSG00000146063 |
| HLA-DMB | ualy Exclusive Ex | e4/e5 | down (e4) | 1,92 | ENSG00000242574 |
| TRIQK | Complex | e6/e7 | up (e6) | 1,92 | ENSG00000205133 |
| ZNF706 | Exon Cassette | e4 | up | 1,92 | ENSG00000120963 |
| SEMA4D | Alter. First Exon | 1-17,e19/e2 | up (e1-17,e19) | 1,92 | ENSG00000187764 |
| ATP11C | Exon Cassette | e31 | up | 1,92 | ENSG00000101974 |
| NFASC | ter. Terminal Exo | e21,e22-25 | vn (e21,e22-25,e28- | 1,91 | ENSG00000163531 |
| CDCP2 // CYB5R | ter. Terminal Exo | 6/e7,e11-1 | up (e6) | 1,91 | ENSG00000157211 // ENSG00000215883 |

| | | | | | |
|---|---|---|---|---|---|
| CDCP2 // CYB5R | Exon Cassette | e6 | up | 1,91 | ENSG00000157211 // ENSG00000215883 |
| ITGB3BP | Exon Cassette | e9 | down | 1,91 | ENSG00000142856 |
| FAS | Exon Cassette | e7 | up | 1,91 | ENSG00000026103 |
| DGKZ | Alter. First Exon | e2-29/e30 | up (e2-29) | 1,91 | ENSG00000149091 |
| NUMA1 | Alter. First Exon | e1-3,e6/e7 | up (e1-3,e6) | 1,91 | ENSG00000137497 |
| // CHURC1-FNTI | Exon Cassette | e12 | up | 1,91 | ENSG00000125954 // ENSG00000257365 // ENSG00000258289 |
| HYPK // SERF2 | Complex | e5 | up | 1,91 | ENSG00000140264 // ENSG00000242028 |
| ZNF254 | Alter. First Exon | 1-2,e4,e11/e | up (e1-2,e4,e11) | 1,91 | ENSG00000213096 |
| BCL2L11 | Complex | e2/e9-11,e1 | up (e9-11,e13) | 1,91 | ENSG00000153094 |
| BCL2L11 | Exon Cassette | e9-11 | up | 1,91 | ENSG00000153094 |
| CCDC150 | Alter. First Exon | e14-17,e20 | 5,e8-12,e14-17,e2 | 1,91 | ENSG00000144395 |
| CASP8 | Complex | e16-17 | down | 1,91 | ENSG00000064012 |
| TTN | Exon Cassette | 149,e151-15 | up | 1,91 | ENSG00000155657 |
| RAB5A | Exon Cassette | e3 | up | 1,91 | ENSG00000144566 |
| LPP | Alter. First Exon | e1,e5-8/e9 | up (e1,e5-8) | 1,91 | ENSG00000145012 |
| DPH3 | Exon Cassette | e2 | up | 1,91 | ENSG00000154813 |
| SHROOM3 | ter. Terminal Exo | 11,e13-15/e | down (e8-11,e13-15 | 1,91 | ENSG00000138771 |
| GPBP1 | Exon Cassette | e8 | up | 1,91 | ENSG00000062194 |
| ECHDC1 | Exon Cassette | e5 | up | 1,91 | ENSG00000093144 |
| RGL2 | Intron Retention | e14 | up | 1,91 | ENSG00000237441 |
| RABL5 | Intron Retention | e4 | up | 1,91 | ENSG00000128581 |
| CDK16 | Intron Retention | e13,e14 | up | 1,91 | ENSG00000102225 |

| | | | | | |
|---|---|---|---|---|---|
| CUL4B | Alter. First Exon | e1-2/e3 | up (e1-2) | 1,91 | ENSG00000 158290 |
| EPB41 | ter. Terminal Exo | 4/e18,e21-2 | up (e14) | 1,9 | ENSG00000 159023 |
| EPB41 | ter. Terminal Exo | 4/e15,e17-1 | up (e14) | 1,9 | ENSG00000 159023 |
| NFASC | ter. Terminal Exo | 20/e22-29,e3 | own (e22-29,e34-36 | 1,9 | ENSG00000 163531 |
| FAM129A | Exon Cassette | e9 | down | 1,9 | ENSG00000 135842 |
| SEPHS1 | Exon Cassette | e8,e9 | up | 1,9 | ENSG00000 086475 |
| PTMS | Alter. First Exon | e1/e2 | up (e1) | 1,9 | ENSG00000 159335 |
| MYL6 | ter. Terminal Exo | e7-8/e9 | up (e7-8) | 1,9 | ENSG00000 092841 |
| TBC1D30 | ter. Terminal Exo | e14/e15 | up (e14) | 1,9 | ENSG00000 111490 |
| TNFRSF1A | Alter. First Exon | ,e3-6,e8-11/ | up (e1,e3-6,e8-11) | 1,9 | ENSG00000 067182 |
| APOPT1 // KLC1 | Exon Cassette | e19 | down | 1,9 | ENSG00000 126214 // ENSG00000 256053 |
| 2AP // IGHD // IG | ter. Terminal Exo | ,e130-135/e | up (e98,e130-135) | 1,9 | ENSG00000 211896 // ENSG00000 211898 // ENSG00000 213140 |
| 2AP // IGHD // IG | ter. Terminal Exo | 01/e98,e13 | up (e98,e130-135) | 1,9 | ENSG00000 211896 // ENSG00000 211898 // ENSG00000 213140 |
| ZFP90 | Alter. First Exon | e3-6/e9 | down (e3-6) | 1,9 | ENSG00000 184939 |
| LRRC37B | ter. Terminal Exo | 2/e13,e15-1 | down (e13,e15-18) | 1,9 | ENSG00000 185158 |
| ACSF2 | Alter. First Exon | 1-3,e5-11/e1 | down (e1-3,e5-11) | 1,9 | ENSG00000 167107 |
| RDM1 | Exon Cassette | e5 | down | 1,9 | ENSG00000 187456 |
| ZNF271 | Exon Cassette | e2 | up | 1,9 | ENSG00000 257267 |
| RBM39 | Exon Cassette | e3 | up | 1,9 | ENSG00000 131051 |
| CTNNB1 | Complex | e19 | down | 1,9 | ENSG00000 168036 |
| EOMES | Complex | e4-7 | down | 1,9 | ENSG00000 163508 |

| | | | | | |
|---|---|---|---|---|---|
| FAM193A | Alter. First Exon | e3-7/e8 | down (e3-7) | 1,9 | ENSG00000125386 |
| DNAJB14 | ter. Terminal Exo | e2-5/e7-12 | up (e2-5) | 1,9 | ENSG00000164031 |
| MANBA | Exon Cassette | e6 | up | 1,9 | ENSG00000109323 |
| MAPK14 | ualy Exclusive Ex | e9/e10 | up (e9) | 1,9 | ENSG00000112062 |
| SMARCD3 | Intron Retention | e10 | up | 1,9 | ENSG00000082014 |
| ZBTB33 | Exon Cassette | e2 | up | 1,9 | ENSG00000177485 |
| DFFB | lter. Acceptor Sit | e4 | up | 1,89 | ENSG00000169598 |
| DFFB | Complex | e4 | up | 1,89 | ENSG00000169598 |
| DFFB | Complex | e2/e4 | up (e4) | 1,89 | ENSG00000169598 |
| PSMB4 | Intron Retention | e4 | up | 1,89 | ENSG00000159377 |
| USP21 | Intron Retention | e10 | up | 1,89 | ENSG00000143258 |
| NFASC | ter. Terminal Exo | e21,e22-29 | vn (e21,e22-29,e34- | 1,89 | ENSG00000163531 |
| SLC16A1 | Complex | e1,e2 | down | 1,89 | ENSG00000155380 |
| ANGEL2 | Alter. First Exon | e1/e2,e4 | up (e1) | 1,89 | ENSG00000174606 |
| RNH1 | Alter. First Exon | e1/e3 | down (e1) | 1,89 | ENSG00000023191 |
| GEF25 // SLC26 | Exon Cassette | e31-33 | up | 1,89 | ENSG00000135502 // ENSG00000240771 |
| TBC1D30 | ter. Terminal Exo | e14-16/e15 | up (e14-16) | 1,89 | ENSG00000111490 |
| PLXNC1 | Alter. First Exon | e1-15/e16 | up (e1-15) | 1,89 | ENSG00000136040 |
| DENND5B | Exon Cassette | e4 | up | 1,89 | ENSG00000170456 |
| ZMYM2 | Exon Cassette | e12 | up | 1,89 | ENSG00000121741 |
| 2AP // IGHD // IG | Alter. First Exon | 7,e103/e18- | up (e17,e103) | 1,89 | ENSG00000211896 // ENSG00000211898 // ENSG00000213140 |
| CIRBP | Intron Retention | e6 | up | 1,89 | ENSG00000099622 |
| ZNF254 | Alter. First Exon | e1-2,e11/e9 | up (e1-2,e11) | 1,89 | ENSG00000213096 |

| | | | | | |
|---|---|---|---|---|---|
| 1S1 // IL4I1 // NU | Complex | e2,e3 | up | 1,89 | ENSG00000 104951 // ENSG00000 204673 // ENSG00000 213024 |
| IL17RC | Exon Cassette | e12 | up | 1,89 | ENSG00000 163702 |
| RFTN1 | Alter. First Exon | 2,e4-6,e8-9/ | own (e1-2,e4-6,e8-9 | 1,89 | ENSG00000 131378 |
| APBB2 | Intron Retention | e9 | up | 1,89 | ENSG00000 163697 |
| RUSC2 | Alter. Donor Site | e3 | up | 1,89 | ENSG00000 198853 |
| SH3GLB2 | Alter. First Exon | e1-7/e8 | down (e1-7) | 1,89 | ENSG00000 148341 |
| SEC24C | Intron Retention | e23 | up | 1,88 | ENSG00000 176986 |
| H19 | Exon Cassette | e5 | down | 1,88 | ENSG00000 130600 |
| GALK2 | Exon Cassette | e12 | up | 1,88 | ENSG00000 156958 |
| CORO1A | Intron Retention | e5 | up | 1,88 | ENSG00000 102879 |
| PLA2G15 | Alter. First Exon | e1/e3 | down (e1) | 1,88 | ENSG00000 103066 |
| ACADVL | Intron Retention | e10 | up | 1,88 | ENSG00000 072778 |
| ERBB2 | ter. Terminal Exo | 29/e30,e31-3 | down (e30,e31-33) | 1,88 | ENSG00000 141736 |
| RPL38 | ter. Terminal Exo | e3/e4-5 | up (e3) | 1,88 | ENSG00000 172809 |
| TVP23C // TVP23 | Exon Cassette | e6 | up | 1,88 | ENSG00000 175106 // ENSG00000 239704 // ENSG00000 259024 |
| RNF165 | Exon Cassette | e6-7 | up | 1,88 | ENSG00000 141622 |
| TNFRSF11A | Exon Cassette | e8,e9 | up | 1,88 | ENSG00000 141655 |
| TNFRSF11A | Exon Cassette | e8,e9-10 | up | 1,88 | ENSG00000 141655 |
| MAP2K7 | Exon Cassette | e3 | up | 1,88 | ENSG00000 076984 |
| ZNF69 | Alter. First Exon | e1-4,e6/e5 | down (e1-4,e6) | 1,88 | ENSG00000 198429 |
| TYK2 | Complex | e1/e6-7 | up (e6-7) | 1,88 | ENSG00000 105397 |
| CCDC150 | Alter. First Exon | -18,e20-21,e | 1-12,e14-18,e20-21 | 1,88 | ENSG00000 144395 |

| | | | | | |
|---|---|---|---|---|---|
| BIN1 | Exon Cassette | e18 | up | 1,88 | ENSG00000 136717 |
| PCBP3 | ter. Terminal Exo | e11/e12-16 | up (e12-16) | 1,88 | ENSG00000 183570 |
| EWSR1 | Complex | e9,e10-11 | up | 1,88 | ENSG00000 182944 |
| SACM1L | Exon Cassette | e2 | up | 1,88 | ENSG00000 211456 |
| TMEM175 | lter. Acceptor Sit | e7 | down | 1,88 | ENSG00000 127419 |
| FAM114A1 | Complex | e4-6 | up | 1,88 | ENSG00000 197712 |
| REST | ter. Terminal Exo | 8,e10-11/e14 | up (e5-8,e10-11) | 1,88 | ENSG00000 084093 |
| REST | ter. Terminal Exo | ,e10-11/e13 | up (e5-8,e10-11) | 1,88 | ENSG00000 084093 |
| HINT1 | Alter. Donor Site | e2 | up | 1,88 | ENSG00000 169567 |
| HARS | Intron Retention | e7 | up | 1,88 | ENSG00000 170445 |
| PHF10 | ter. Terminal Exo | 2/e13,e14-1 | down (e12) | 1,88 | ENSG00000 130024 |
| CUTA | Intron Retention | e1,e2 | down | 1,88 | ENSG00000 112514 |
| STAU2 | ter. Terminal Exo | e18/e20,e21 | up (e18) | 1,88 | ENSG00000 040341 |
| BGT1 // RALGD | Alter. First Exon | e2-7/e10 | up (e2-7) | 1,88 | ENSG00000 148288 // ENSG00000 160271 |
| RPL10 | Intron Retention | e5,e6 | up | 1,88 | ENSG00000 147403 |
| MID1 | Complex | e1,e3 | up | 1,88 | ENSG00000 101871 |
| NUP62CL | Exon Cassette | e5 | up | 1,88 | ENSG00000 198088 |
| EPB41 | ter. Terminal Exo | 15,e17-18,e | up (e14) | 1,87 | ENSG00000 159023 |
| EPB41 | ter. Terminal Exo | 4/e18,e20-2 | up (e14) | 1,87 | ENSG00000 159023 |
| ITGB3BP | ter. Terminal Exo | e8/e9-11 | up (e8) | 1,87 | ENSG00000 142856 |
| KIAA1549L | Exon Cassette | e11 | up | 1,87 | ENSG00000 110427 |
| SORD | lter. Acceptor Sit | e4 | up | 1,87 | ENSG00000 140263 |
| STX4 | Intron Retention | e6 | up | 1,87 | ENSG00000 103496 |
| SLC12A4 | Exon Cassette | e5 | up | 1,87 | ENSG00000 124067 |
| PDXDC2P | Exon Cassette | e26 | up | 1,87 | ENSG00000 196696 |

| | | | | | |
|---|---|---|---|---|---|
| ERAL1 | Complex | e4,e5 | up | 1,87 | ENSG00000132591 |
| POLI | Intron Retention | e1 | up | 1,87 | ENSG00000101751 |
| BCAM | Complex | e11-15 | down | 1,87 | ENSG00000187244 |
| DNAJB2 | Intron Retention | e6 | up | 1,87 | ENSG00000135924 |
| PIGF | Exon Cassette | e6 | up | 1,87 | ENSG00000151665 |
| NDUFA10 | Intron Retention | e6,e7 | up | 1,87 | ENSG00000130414 |
| RPN2 | Exon Cassette | e17 | down | 1,87 | ENSG00000118705 |
| UQCC | ter. Terminal Exo | e8/e10-11 | up (e8) | 1,87 | ENSG00000101019 |
| // PRR5 // PRR5 | Exon Cassette | e8,e9-10 | down | 1,87 | ENSG00000186654 // ENSG00000241484 // ENSG00000248405 |
| SETD5 | Complex | e7,e8 | up | 1,87 | ENSG00000168137 |
| OXSR1 | Exon Cassette | e4 | up | 1,87 | ENSG00000172939 |
| PIGG | Complex | e1 | down | 1,87 | ENSG00000174227 |
| PPAP2A | Exon Cassette | e2 | down | 1,87 | ENSG00000067113 |
| RAB24 | Intron Retention | e6 | up | 1,87 | ENSG00000169228 |
| SRSF3 | Exon Cassette | e4 | up | 1,87 | ENSG00000112081 |
| IRPS24 // URGC | Alter. First Exon | e3,e5/e4 | up (e4) | 1,87 | ENSG00000062582 // ENSG00000106608 |
| CHCHD3 | Exon Cassette | e9-10 | up | 1,87 | ENSG00000106554 |
| CHMP5 | Exon Cassette | e7 | down | 1,87 | ENSG00000086065 |
| GALT | Intron Retention | e3 | up | 1,87 | ENSG00000213930 |
| ABCA2 | Intron Retention | e40 | up | 1,87 | ENSG00000107331 |
| BRWD3 | Exon Cassette | e15 | up | 1,87 | ENSG00000165288 |
| DHX9 // NPL | Alter. First Exon | e1,e4-6/e9 | down (e1,e4-6) | 1,86 | ENSG00000135829 // ENSG00000135838 |

| | | | | | |
|---|---|---|---|---|---|
| MYSM1 | Intron Retention | e5-6 | up | 1,86 | ENSG00000162601 |
| ZNF692 | Intron Retention | e4 | up | 1,86 | ENSG00000171163 |
| MYO3A | Alter. First Exon | e1-33/e34 | up (e1-33) | 1,86 | ENSG00000095777 |
| TCF7L2 | ualy Exclusive Ex | e20/e21 | down (e20) | 1,86 | ENSG00000148737 |
| TCF7L2 | Exon Cassette | e18,e19 | up | 1,86 | ENSG00000148737 |
| KCNMA1 | Alter. First Exon | e6-15/e16 | down (e6-15) | 1,86 | ENSG00000156113 |
| ACCS // EXT2 | ter. Terminal Exo | 20-21,e23-2 | 9,e20-21,e23-25,e2 | 1,86 | ENSG00000110455 // ENSG00000151348 |
| RAB6A | Exon Cassette | e6-8 | down | 1,86 | ENSG00000175582 |
| CADM1 | Alter. First Exon | e1,e3-8/e10 | up (e1,e3-8) | 1,86 | ENSG00000182985 |
| C12orf76 | Exon Cassette | e8 | up | 1,86 | ENSG00000174456 |
| EVL | ter. Terminal Exo | 4/e15,e16-1 | up (e14) | 1,86 | ENSG00000196405 |
| PML | ter. Terminal Exo | e4/e8 | up (e8) | 1,86 | ENSG00000140464 |
| NDRG4 | Exon Cassette | e7 | up | 1,86 | ENSG00000103034 |
| ADAP2 | Exon Cassette | e4 | up | 1,86 | ENSG00000184060 |
| SPHK1 | Alter. First Exon | e6/e7,e8 | down (e7,e8) | 1,86 | ENSG00000176170 |
| SLC46A1 | Exon Cassette | e3 | up | 1,86 | --- |
| PLEKHM1 | Exon Cassette | e6 | up | 1,86 | ENSG00000225190 |
| SMARCA4 | Complex | e4,e5 | up | 1,86 | ENSG00000127616 |
| SMARCA4 | Exon Cassette | e4 | up | 1,86 | ENSG00000127616 |
| HNRNPUL1 | Alter. Donor Site | e4 | up | 1,86 | ENSG00000105323 |
| TYK2 | Complex | e1-5 | up | 1,86 | ENSG00000105397 |
| ZNF816 // ZNF8 | ter. Terminal Exo | e4-5/e7 | down (e4-5) | 1,86 | ENSG00000180257 // ENSG00000213801 // ENSG00000221874 |

| | | | | | |
|---|---|---|---|---|---|
| ZNF816 // ZNF8 | ter. Terminal Exo | e5/e7 | down (e5) | 1,86 | ENSG00000 180257 // ENSG00000 213801 // ENSG00000 221874 |
| EPAS1 | Alter. First Exon | e1-11/e12 | down (e1-11) | 1,86 | ENSG00000 116016 |
| UBE2G2 | Alter. First Exon | e1,e3-5/e6 | down (e1,e3-5) | 1,86 | ENSG00000 184787 |
| SMTN | Intron Retention | e17,e18 | up | 1,86 | ENSG00000 183963 |
| NOL12 // TRIOBP | Intron Retention | e6 | up | 1,86 | ENSG00000 100106 // ENSG00000 256872 |
| BID | Complex | e6,e7 | up | 1,86 | ENSG00000 015475 |
| OX17 // POPDC | Exon Cassette | e7-9 | down | 1,86 | ENSG00000 121577 // ENSG00000 138495 |
| STAG1 | ter. Terminal Exo | 0/e11,e12-2 | up (e10) | 1,86 | ENSG00000 118007 |
| MTHFD2L | Exon Cassette | e5 | up | 1,86 | ENSG00000 163738 |
| CYP4V2 | Intron Retention | e7-8 | up | 1,86 | ENSG00000 145476 |
| BRIX1 | Intron Retention | e6 | up | 1,86 | ENSG00000 113460 |
| HACE1 | Exon Cassette | e17 | up | 1,86 | ENSG00000 085382 |
| NRG1 | Complex | e6-7/e10 | up (e6-7) | 1,86 | ENSG00000 157168 |
| COQ4 | Intron Retention | e6 | up | 1,86 | ENSG00000 167113 |
| EPB41 | Exon Cassette | e15 | down | 1,85 | ENSG00000 159023 |
| PSEN2 | Exon Cassette | e4 | down | 1,85 | ENSG00000 143801 |
| TGLF9P // PAR | ter. Terminal Exo | 18-25/e32-3 | up (e18-25) | 1,85 | ENSG00000 227345 |
| ATN1 // C12orf57 | Intron Retention | e14 | up | 1,85 | ENSG00000 111678 |
| ARNTL2 | Alter. Donor Site | e1 | down | 1,85 | ENSG00000 029153 |
| LETMD1 | Exon Cassette | e5,e6-7 | up | 1,85 | ENSG00000 050426 |
| RAB5B | Complex | e1 | up | 1,85 | ENSG00000 111540 |
| CRIP2 | Exon Cassette | e5-8 | down | 1,85 | ENSG00000 182809 |

| | | | | | |
|---|---|---|---|---|---|
| CHD2 // MIR3175 | Intron Retention | e17 | up | 1,85 | ENSG00000173575 // ENSG00000264173 |
| CYLD | Exon Cassette | e4 | up | 1,85 | ENSG00000083799 |
| ACD | Intron Retention | e4 | up | 1,85 | ENSG00000102977 |
| LRRC37B | ter. Terminal Exo | e12/e15-18 | down (e15-18) | 1,85 | ENSG00000185158 |
| --- | Exon Cassette | e6,e9-11 | up | 1,85 | --- |
| NSF // NSFP1 | Complex | e3/e4 | up (e3) | 1,85 | ENSG00000073969 // ENSG00000260075 |
| DNAJC10 | Complex | e11-12 | up | 1,85 | ENSG00000077232 |
| FAM136A | Exon Cassette | e2 | up | 1,85 | ENSG00000035141 |
| BTBD3 | Alter. First Exon | e2/e3 | up (e2) | 1,85 | ENSG00000132640 |
| CENPM | Complex | e4/e5 | up (e4) | 1,85 | ENSG00000100162 |
| SACM1L | Complex | e1,e2 | up | 1,85 | ENSG00000211456 |
| AMT | Exon Cassette | e5 | down | 1,85 | ENSG00000145020 |
| OX17 // POPDC | Exon Cassette | e7,e8 | down | 1,85 | ENSG00000121577 // ENSG00000138495 |
| DOPEY1 | Alter. First Exon | e1/e2 | down (e1) | 1,85 | ENSG00000083097 |
| SOBP | ter. Terminal Exo | e6/e7,e8-11 | down (e7,e8-11) | 1,85 | ENSG00000112320 |
| EPB41L2 | Exon Cassette | e17 | down | 1,85 | ENSG00000079819 |
| RXRB | Intron Retention | e7 | up | 1,85 | ENSG00000204231 |
| TRA2A | Intron Retention | e3 | up | 1,85 | ENSG00000164548 |
| IRPS24 // URGC | Alter. First Exon | e3/e5 | up (e5) | 1,85 | ENSG00000062582 // ENSG00000106608 |
| FIGNL1 | Intron Retention | e2 | up | 1,85 | ENSG00000132436 |
| STX1A | Complex | e3-6 | down | 1,85 | ENSG00000106089 |
| DOCK11 | Exon Cassette | e31 | up | 1,85 | ENSG00000147251 |

| | | | | | |
|---|---|---|---|---|---|
| SH3D21 | Alter. First Exon | e1-3/e4 | down (e1-3) | 1,84 | ENSG00000214193 |
| PLA2G4A | Exon Cassette | e6-7 | down | 1,84 | ENSG00000116711 |
| SYT14 | Complex | e1,e5/e2-4 | up (e2-4) | 1,84 | ENSG00000143469 |
| VPS26A | Exon Cassette | e3 | up | 1,84 | ENSG00000122958 |
| RUFY2 | Alter. First Exon | e1/e2 | up (e1) | 1,84 | ENSG00000204130 |
| CTNND1 // TMX2 | Complex | e10 | up | 1,84 | ENSG00000198561 // ENSG00000213593 |
| TMEM126B | Exon Cassette | e3 | up | 1,84 | ENSG00000171204 |
| AHNAK | ter. Terminal Exo | e5/e8-9 | up (e5) | 1,84 | ENSG00000124942 |
| LETMD1 | Exon Cassette | e6-7 | up | 1,84 | ENSG00000050426 |
| CNOT2 | Intron Retention | e11 | up | 1,84 | ENSG00000111596 |
| DAAM1 | Exon Cassette | e16 | down | 1,84 | ENSG00000100592 |
| HNRNPC | Exon Cassette | e2,e3-4 | up | 1,84 | ENSG00000092199 |
| 2AP // IGHD // IG | Alter. First Exon | 4,e101,e137 | wn (e33-34,e101,e1 | 1,84 | ENSG00000211896 // ENSG00000211898 // ENSG00000213140 |
| 2AP // IGHD // IG | Alter. First Exon | 5-38,e40/e1 | down (e35-38,e40) | 1,84 | ENSG00000211896 // ENSG00000211898 // ENSG00000213140 |
| GLOD4 | Complex | e6,e7 | up | 1,84 | ENSG00000167699 |
| C18orf25 | Exon Cassette | e3 | up | 1,84 | ENSG00000152242 |
| ESCO1 | Exon Cassette | e9 | up | 1,84 | ENSG00000141446 |
| PPP5C | Intron Retention | e6 | up | 1,84 | ENSG00000011485 |
| LPIN1 | ter. Terminal Exo | e13/e14-16 | down (e14-16) | 1,84 | ENSG00000134324 |
| UGGT1 | Alter. Donor Site | e1 | down | 1,84 | ENSG00000136731 |
| BIN1 | Exon Cassette | e17-18 | up | 1,84 | ENSG00000136717 |

| | | | | | |
|---|---|---|---|---|---|
| HPS4 | Intron Retention | e11 | up | 1,84 | ENSG00000 100099 |
| PTPRG | Complex | e2/e4 | up (e2) | 1,84 | ENSG00000 144724 |
| DPH3 | Complex | e1,e2 | up | 1,84 | ENSG00000 154813 |
| EXOSC9 | Intron Retention | e10,e11 | up | 1,84 | ENSG00000 123737 |
| OCLN | Complex | e4-6,e8-9 | down | 1,84 | ENSG00000 197822 |
| TMEM120A | Intron Retention | e6 | up | 1,84 | ENSG00000 189077 |
| CSPP1 | Exon Cassette | e7 | up | 1,84 | ENSG00000 104218 |
| IAA1984 // RABL | ter. Terminal Exo | e25/e26 | up (e25) | 1,84 | ENSG00000 196642 // ENSG00000 213213 |
| SH2D3C | Exon Cassette | e4 | up | 1,84 | ENSG00000 095370 |
| MAGED2 | Intron Retention | e11 | up | 1,84 | ENSG00000 102316 |
| CUL4B | Intron Retention | e16 | up | 1,84 | ENSG00000 158290 |
| SAG4 // MAGEA | Exon Cassette | e3 | up | 1,84 | ENSG00000 183305 // ENSG00000 242599 |
| CTNNBIP1 | ter. Terminal Exo | e2,e4-7/e3 | down (e2,e4-7) | 1,83 | ENSG00000 178585 |
| TTC13 | Exon Cassette | e6,e7 | up | 1,83 | ENSG00000 143643 |
| MYO3A | Alter. First Exon | e2-33/e34 | up (e2-33) | 1,83 | ENSG00000 095777 |
| TRUB1 | Complex | e1 | up | 1,83 | ENSG00000 165832 |
| ITGB1 | Exon Cassette | e18 | up | 1,83 | ENSG00000 150093 |
| TIAL1 | ter. Terminal Exo | e12/e13-14 | up (e12) | 1,83 | ENSG00000 151923 |
| DIXDC1 | Alter. First Exon | e8-12/e13 | up (e8-12) | 1,83 | ENSG00000 150764 |
| WNK1 | Exon Cassette | e13 | up | 1,83 | ENSG00000 060237 |
| IFT81 | Exon Cassette | e20,e21 | up | 1,83 | ENSG00000 122970 |
| MTIF3 | Alter. First Exon | e1-3/e4 | down (e1-3) | 1,83 | ENSG00000 122033 |
| MTRF1 | Intron Retention | e3 | up | 1,83 | ENSG00000 120662 |
| JKAMP | Exon Cassette | e2 | down | 1,83 | ENSG00000 050130 |

| | | | | | |
|---|---|---|---|---|---|
| C16orf93 | Intron Retention | e3 | up | 1,83 | ENSG00000196118 |
| BPTF | ter. Terminal Exc | e32/e33 | up (e32) | 1,83 | ENSG00000171634 |
| CBX2 | ter. Terminal Exc | e3/e4-5 | down (e4-5) | 1,83 | ENSG00000173894 |
| HN1 | Exon Cassette | e3 | up | 1,83 | ENSG00000189159 |
| --- | Exon Cassette | e9-11 | up | 1,83 | --- |
| DMKN | Exon Cassette | e9,e10 | down | 1,83 | ENSG00000161249 |
| BCL2L11 | Exon Cassette | e10,e11 | up | 1,83 | ENSG00000153094 |
| PKP4 | ter. Terminal Exc | e17/e18-23 | up (e17) | 1,83 | ENSG00000144283 |
| PIKFYVE | ter. Terminal Exc | 3/e14,e15-4 | up (e13) | 1,83 | ENSG00000115020 |
| CCT4 | Complex | e1,e2-3 | up | 1,83 | ENSG00000115484 |
| LSM14B | lter. Acceptor Sit | e4 | up | 1,83 | ENSG00000149657 |
| CSNK2A1 | Exon Cassette | e2 | down | 1,83 | ENSG00000101266 |
| MKL1 | Exon Cassette | e2 | up | 1,83 | ENSG00000196588 |
| TRPC1 | Exon Cassette | e3 | down | 1,83 | ENSG00000144935 |
| TSC22D2 | Exon Cassette | e3 | up | 1,83 | ENSG00000196428 |
| FIP1L1 | Intron Retention | e1-2 | up | 1,83 | ENSG00000145216 |
| FIP1L1 | Exon Cassette | e2 | up | 1,83 | ENSG00000145216 |
| SCLT1 | Alter. First Exon | 4,e6,e8-10/e | up (e1-4,e6,e8-10) | 1,83 | ENSG00000151466 |
| PDLIM3 | Alter. Donor Site | e5 | down | 1,83 | ENSG00000154553 |
| GPR98 | Alter. First Exon | e67-85/e86 | down (e67-85) | 1,83 | ENSG00000164199 |
| CDKAL1 | ter. Terminal Exc | e7/e8-17 | down (e8-17) | 1,83 | ENSG00000145996 |
| TAZ | Intron Retention | e8 | down | 1,83 | ENSG00000102125 |
| CD99 | Complex | e2-3/e4 | up (e2-3) | 1,83 | ENSG00000002586 |
| MST1P2 | lter. Acceptor Sit | e4 | up | 1,82 | ENSG00000186301 |
| NDUFS2 | Intron Retention | e14 | up | 1,82 | ENSG00000158864 |
| MDM4 | Exon Cassette | e7,e8-12 | down | 1,82 | ENSG00000198625 |

| | | | | | |
|---|---|---|---|---|---|
| TRIM33 | Exon Cassette | e20 | up | 1,82 | ENSG00000197323 |
| SART3 | Intron Retention | e9 | up | 1,82 | ENSG00000075856 |
| APOPT1 // KLC1 | ter. Terminal Exc | 19-23/e24-2 | up (e19-23) | 1,82 | ENSG00000126214 // ENSG00000256053 |
| GCH1 | Complex | e6 | down | 1,82 | ENSG00000131979 |
| 2AP // IGHD // IG | Alter. First Exon | e103/e35-38 | up (e17,e103) | 1,82 | ENSG00000211896 // ENSG00000211898 // ENSG00000213140 |
| CAPN3 // GANC | Alter. First Exon | e33-35,e37 | (e30-31,e33-35,e3 | 1,82 | ENSG00000092529 // ENSG00000214013 |
| NEDD4 | Exon Cassette | e25-26 | down | 1,82 | ENSG00000069869 |
| NDRG4 | Complex | e6,e7 | up | 1,82 | ENSG00000103034 |
| DHRS7B | ter. Terminal Exc | e3/e4-8 | down (e3) | 1,82 | ENSG00000109016 |
| PSMC5 | Intron Retention | e3 | up | 1,82 | ENSG00000087191 |
| FBXO17 // SARS | Intron Retention | e12,e13 | up | 1,82 | ENSG00000104835 // ENSG00000269190 |
| PLEKHH2 | Alter. First Exon | 1-2,e4-9/e1 | down (e1-2,e4-9) | 1,82 | ENSG00000152527 |
| SPAG16 | Complex | e1/e3 | up (e1) | 1,82 | ENSG00000144451 |
| CLK1 // PPIL3 | Alter. First Exon | e5-8,e10-11 | wn (e1-3,e5-8,e10-1 | 1,82 | ENSG00000013441 // ENSG00000240344 |
| NPEPL1 // STX1 | Alter. First Exon | e9,e12/e14 | down (e9,e12) | 1,82 | ENSG00000124222 // ENSG00000215440 |
| MCM3AP-AS1 | ter. Terminal Exc | e5/e6,e7 | up (e5) | 1,82 | ENSG00000215424 |
| RPL15 | Intron Retention | e1-2 | up | 1,82 | ENSG00000174748 |
| PVRL3 | Alter. Donor Site | e1 | down | 1,82 | ENSG00000177707 |
| AMT | Complex | e5/e6,e7 | down (e5) | 1,82 | ENSG00000145020 |

| | | | | | |
|---|---|---|---|---|---|
| PCGF3 | Exon Cassette | e4 | down | 1,82 | ENSG00000 185619 |
| ZNF76 | ter. Terminal Exo | 0/e11,e12-1 | up (e10) | 1,82 | ENSG00000 065029 |
| S5 // EEF1E1 // T | ter. Terminal Exo | /e7,e9,e11- | down (e7,e9,e11-12 | 1,82 | ENSG00000 124802 // ENSG00000 188428 // ENSG00000 239264 |
| SNK2B // LY6G5 | Intron Retention | e4 | up | 1,82 | ENSG00000 204435 // ENSG00000 240053 |
| BAG6 | Alter. First Exon | e1/e2 | up (e1) | 1,82 | ENSG00000 204463 |
| COA1 | ter. Terminal Exo | e9/e11 | up (e9) | 1,82 | ENSG00000 106603 |
| PMS2P4 | lter. Acceptor Sit | e4 | up | 1,82 | ENSG00000 067601 |
| GTF2I | ter. Terminal Exo | 1/e13,e14-3 | down (e13,e14-34) | 1,82 | ENSG00000 077809 |
| ATP6V1B2 | Intron Retention | e9 | up | 1,82 | ENSG00000 147416 |
| AKNA | Alter. First Exon | e1,e5/e3 | up (e1,e5) | 1,82 | ENSG00000 106948 |
| OGT | ter. Terminal Exo | 4-6/e5,e7-2 | up (e4-6) | 1,82 | ENSG00000 147162 |
| FMR1 | Exon Cassette | e13 | up | 1,82 | ENSG00000 102081 |
| MTMR1 | Alter. First Exon | 2-3,e5-11/e1 | down (e2-3,e5-11) | 1,82 | ENSG00000 063601 |
| TAB3 | Intron Retention | e10 | up | 1,82 | ENSG00000 157625 |
| ZC4H2 | Intron Retention | e6 | up | 1,82 | ENSG00000 126970 |
| AMY2B // RNPC3 | Alter. First Exon | e17/e19,e20 | up (e17) | 1,81 | ENSG00000 185946 // ENSG00000 240038 |
| SOAT1 | Exon Cassette | e2 | up | 1,81 | ENSG00000 057252 |
| NFASC | ter. Terminal Exo | -20/e22,e23 | down (e22,e23-29) | 1,81 | ENSG00000 163531 |
| ZNF678 | Exon Cassette | e3 | up | 1,81 | ENSG00000 181450 |
| S100A13 | Alter. First Exon | e1-2/e3 | down (e1-2) | 1,81 | ENSG00000 189171 |
| RTKN2 | Exon Cassette | e10 | up | 1,81 | ENSG00000 182010 |
| C10orf118 | lter. Acceptor Sit | e14 | up | 1,81 | ENSG00000 165813 |

| | | | | | |
|---|---|---|---|---|---|
| AKAP6 | Alter. First Exon | e1-2/e3 | up (e1-2) | 1,81 | ENSG00000151320 |
| KTN1 | Intron Retention | e43,e44-45 | up | 1,81 | ENSG00000126777 |
| ATXN3 | Complex | ,e6,e8-10/e | down (e2,e6,e8-10) | 1,81 | ENSG00000066427 |
| RAD51 | Exon Cassette | e4 | up | 1,81 | ENSG00000051180 |
| CAPN3 // GANC | Alter. First Exon | -31,e33-42/ | own (e30-31,e33-42 | 1,81 | ENSG00000092529 // ENSG00000214013 |
| ZNF720 | Complex | e7 | down | 1,81 | ENSG00000197302 |
| JPH3 | ter. Terminal Exo | e3/e4,e6-9 | up (e3) | 1,81 | ENSG00000154118 |
| KCTD13 | Complex | e4,e5 | up | 1,81 | ENSG00000174943 |
| SLC12A4 | Exon Cassette | e11 | down | 1,81 | ENSG00000124067 |
| --- | Exon Cassette | e6-11 | up | 1,81 | --- |
| ZNF574 | Alter. First Exon | e2/e3 | up (e3) | 1,81 | ENSG00000105732 |
| RPS9 | Alter. Donor Site | e4 | up | 1,81 | ENSG00000170889 |
| ZNF573 | Exon Cassette | e7-9 | up | 1,81 | ENSG00000189144 |
| RTN2 | Exon Cassette | e6 | up | 1,81 | ENSG00000125744 |
| KIAA1841 | Complex | 18-19/e20-2 | up (e18-19) | 1,81 | ENSG00000162929 |
| ATG4B | Complex | e6/e7 | up (e6) | 1,81 | ENSG00000168397 |
| PI4KA | Alter. First Exon | e1-32/e33 | down (e1-32) | 1,81 | ENSG00000241973 |
| CSNK1E | ter. Terminal Exo | 8/e11,e16-2 | up (e8) | 1,81 | ENSG00000213923 |
| CCDC50 | Exon Cassette | e6 | up | 1,81 | ENSG00000152492 |
| EOGT | Alter. First Exon | e1-2/e3 | down (e1-2) | 1,81 | ENSG00000163378 |
| SLC4A4 | Complex | e4-5 | down | 1,81 | ENSG00000080493 |
| THAP9 | Exon Cassette | e7 | up | 1,81 | ENSG00000168152 |
| ZFR | Intron Retention | e17 | up | 1,81 | ENSG00000056097 |
| HLA-F | ter. Terminal Exo | e5-6/e11,e12 | up (e5-6) | 1,81 | ENSG00000204642 |
| ATAT1 | Intron Retention | e10 | up | 1,81 | ENSG00000137343 |

| | | | | | |
|---|---|---|---|---|---|
| CAMK2B | Exon Cassette | e13 | up | 1,81 | ENSG00000058404 |
| PNPLA8 | Complex | e2/e3,e5 | up (e3,e5) | 1,81 | ENSG00000135241 |
| NRG1 | Exon Cassette | e18 | up | 1,81 | ENSG00000157168 |
| TJP2 | Complex | e24-25 | down | 1,81 | ENSG00000119139 |
| ZNF26 | Exon Cassette | e2 | up | 1,81 | ENSG00000198393 |
| LPHN2 | Exon Cassette | e13 | down | 1,8 | ENSG00000117114 |
| CD55 | Exon Cassette | e9,e14 | down | 1,8 | ENSG00000196352 |
| EXOSC10 | Intron Retention | e15 | up | 1,8 | ENSG00000171824 |
| MARCH8 | Complex | e7,e8 | down | 1,8 | ENSG00000165406 |
| FRS2 | Exon Cassette | e7 | up | 1,8 | ENSG00000166225 |
| UBE3B | Exon Cassette | e21 | up | 1,8 | ENSG00000151148 |
| ITPR2 | ter. Terminal Exo | /e27-43,e45 | own (e27-43,e45-61 | 1,8 | ENSG00000123104 |
| HDAC7 | Complex | 11-25,e27-2 | down | 1,8 | ENSG00000061273 |
| JKAMP | Complex | e1,e2-3 | down | 1,8 | ENSG00000050130 |
| JKAMP | Complex | e1-2/e3 | down (e1-2) | 1,8 | ENSG00000050130 |
| SRSF5 | ter. Terminal Exo | e5/e6,e7-9 | up (e6,e7-9) | 1,8 | ENSG00000100650 |
| GCH1 | ter. Terminal Exo | e5/e6,e7 | up (e5) | 1,8 | ENSG00000131979 |
| 2AP // IGHD // IG | Alter. First Exon | 32,e102/e10 | down (e32,e102) | 1,8 | ENSG00000211896 // ENSG00000211898 // ENSG00000213140 |
| NDRG4 | Complex | 9-11,e13-20 | up (e8) | 1,8 | ENSG00000103034 |
| SLC12A4 | Exon Cassette | e6 | down | 1,8 | ENSG00000124067 |
| MYADM | Alter. First Exon | e1/e2 | up (e2) | 1,8 | ENSG00000179820 |
| --- | Exon Cassette | e3-8 | down | 1,8 | --- |
| --- | Exon Cassette | e2,e4-9 | down | 1,8 | --- |
| FHL2 | Complex | e2 | down | 1,8 | ENSG00000115641 |
| TBC1D20 | Intron Retention | e5,e6 | up | 1,8 | ENSG00000125875 |

| | | | | | |
|---|---|---|---|---|---|
| IDH3B | Intron Retention | e6 | up | 1,8 | ENSG00000 101365 |
| RANGAP1 | Alter. First Exon | e1,e6/e2-3 | up (e1,e6) | 1,8 | ENSG00000 100401 |
| ARIH2 | Exon Cassette | e5 | up | 1,8 | ENSG00000 177479 |
| MTHFD2L | ter. Terminal Exo | e16/e18-19 | up (e16) | 1,8 | ENSG00000 163738 |
| FAM13A | Alter. First Exon | e6-9,e11-23 | p (e2-4,e6-9,e11-23 | 1,8 | ENSG00000 138640 |
| TCERG1 | Exon Cassette | e6 | down | 1,8 | ENSG00000 113649 |
| ERAP1 | Alter. First Exon | e1/e2 | down (e2) | 1,8 | ENSG00000 164307 |
| IAA1984 // RABL | ter. Terminal Exo | 5/e26,e27,e | up (e25) | 1,8 | ENSG00000 196642 // ENSG00000 213213 |
| WRAP73 | Intron Retention | e10 | up | 1,79 | ENSG00000 116213 |
| EXOSC10 | Intron Retention | e12 | up | 1,79 | ENSG00000 171824 |
| ENSA | ter. Terminal Exo | e5/e6 | down (e5) | 1,79 | ENSG00000 143420 |
| MRPL55 | Complex | e2-3 | up | 1,79 | ENSG00000 162910 |
| OPN3 | Alter. Donor Site | e2 | up | 1,79 | ENSG00000 054277 |
| HINFP | Intron Retention | e6 | up | 1,79 | ENSG00000 172273 |
| IFITM10 | Alter. Donor Site | e2 | down | 1,79 | ENSG00000 244242 |
| NUMA1 | Alter. First Exon | e1,e3/e7 | up (e1,e3) | 1,79 | ENSG00000 137497 |
| CCDC90B | Exon Cassette | e2-3 | down | 1,79 | ENSG00000 137500 |
| ATXN3 | Complex | 2-5,e8-10/e1 | down (e2-5,e8-10) | 1,79 | ENSG00000 066427 |
| HYPK // SERF2 | lter. Acceptor Sit | e5 | up | 1,79 | ENSG00000 140264 // ENSG00000 242028 |
| C16orf13 | Intron Retention | e2 | up | 1,79 | ENSG00000 130731 |
| CHRNB1 | Intron Retention | e4 | down | 1,79 | ENSG00000 170175 |
| ABR | Complex | 3,e10-19,e21 | down | 1,79 | ENSG00000 159842 |
| ZNF559 // ZNF55 | Exon Cassette | e5 | down | 1,79 | ENSG00000 188321 // ENSG00000 188629 |

| | | | | | |
|---|---|---|---|---|---|
| HKR1 | Exon Cassette | e13-14 | up | 1,79 | ENSG00000181666 |
| RPS9 | Intron Retention | e4 | up | 1,79 | ENSG00000170889 |
| C19orf12 | Exon Cassette | e3-4 | up | 1,79 | ENSG00000131943 |
| ZNF677 | ter. Terminal Exo | e5/e6,e7-8 | down (e5) | 1,79 | ENSG00000197928 |
| CCDC150 | Exon Cassette | e6,e7 | down | 1,79 | ENSG00000144395 |
| RHBDD1 | Exon Cassette | e2,e3 | up | 1,79 | ENSG00000144468 |
| ATF2 | ter. Terminal Exo | 1-13,e15,e1 | up (e9) | 1,79 | ENSG00000115966 |
| FN1 | Complex | e40-42 | down | 1,79 | ENSG00000115414 |
| UQCC | ter. Terminal Exo | e8/e9-11 | up (e8) | 1,79 | ENSG00000101019 |
| RABL2B | Intron Retention | e1,e2-3 | up | 1,79 | ENSG00000079974 |
| DAG1 | Alter. First Exon | 1,e3/e7,e8-1 | down (e1,e3) | 1,79 | ENSG00000173402 |
| TBL1XR1 | Complex | e1,e3/e4 | down (e1,e3) | 1,79 | ENSG00000177565 |
| REST | Complex | e5/e12 | up (e5) | 1,79 | ENSG00000084093 |
| SLC4A4 | Complex | e11-16 | up | 1,79 | ENSG00000080493 |
| TBCK | Exon Cassette | e5,e6-7 | up | 1,79 | ENSG00000145348 |
| BCKDHB | Exon Cassette | e7 | up | 1,79 | ENSG00000083123 |
| ARID1B | Exon Cassette | e12 | up | 1,79 | ENSG00000049618 |
| NUDT1 | Alter. First Exon | e1-4/e3 | up (e3) | 1,79 | ENSG00000106268 |
| NDUFB2 | Exon Cassette | e2 | up | 1,79 | ENSG00000090266 |
| RADIL | Exon Cassette | e7 | down | 1,79 | ENSG00000157927 |
| GTF2IRD2 | ter. Terminal Exo | e4-8,e10-16, | up (e3) | 1,79 | ENSG00000196275 |
| GTF2I | ter. Terminal Exo | e11/e13-35 | down (e13-35) | 1,79 | ENSG00000077809 |
| GTF2I | ter. Terminal Exo | 1/e12,e13-3 | down (e12,e13-34) | 1,79 | ENSG00000077809 |
| IL7 | Complex | e6/e7 | up (e6) | 1,79 | ENSG00000104432 |
| IAA1984 // RABL | ter. Terminal Exo | 25/e26,e27-3 | up (e25) | 1,79 | ENSG00000196642 // ENSG00000213213 |

| | | | | | |
|---|---|---|---|---|---|
| --- | Exon Cassette | e14 | up | 1,79 | --- |
| AGTRAP | Exon Cassette | e3 | down | 1,78 | ENSG00000177674 |
| SERINC2 | Exon Cassette | e8 | down | 1,78 | ENSG00000168528 |
| TTC39A | ter. Terminal Exo | 17-18/e19-2 | down (e17-18) | 1,78 | ENSG00000085831 |
| VPS72 | Complex | e6,e7 | up | 1,78 | ENSG00000163159 |
| BRSK2 | Complex | e23,e24 | up | 1,78 | ENSG00000174672 |
| TMEM135 | Complex | e3/e5 | up (e5) | 1,78 | ENSG00000166575 |
| C11orf54 | Exon Cassette | e2,e3 | up | 1,78 | ENSG00000182919 |
| NUMA1 | Alter. First Exon | e1-3/e7 | up (e1-3) | 1,78 | ENSG00000137497 |
| PUS3 | Exon Cassette | e2 | down | 1,78 | ENSG00000110060 |
| ZMYM2 | Exon Cassette | e4 | up | 1,78 | ENSG00000121741 |
| SUPT20H | Exon Cassette | e28 | down | 1,78 | ENSG00000102710 |
| ATXN3 | Complex | ,e5-6,e8-10 | own (e2-3,e5-6,e8-1 | 1,78 | ENSG00000066427 |
| ATXN3 | Complex | 2-6,e8-11/e1 | down (e2-6,e8-11) | 1,78 | ENSG00000066427 |
| MAPKBP1 | Complex | e29-30/e31 | down (e29-30) | 1,78 | ENSG00000137802 |
| RHOT2 | Intron Retention | e10 | up | 1,78 | ENSG00000140983 |
| RNPS1 | Exon Cassette | e3 | up | 1,78 | ENSG00000205937 |
| DYNC1LI2 | Intron Retention | e7,e8 | up | 1,78 | ENSG00000135720 |
| SLC12A4 | Complex | e22,e23-29 | up | 1,78 | ENSG00000124067 |
| ERBB2 | ter. Terminal Exo | 9/e30,e31,e | down (e30,e31,e33) | 1,78 | ENSG00000141736 |
| NBR2 | Exon Cassette | e4 | down | 1,78 | ENSG00000198496 |
| GTPBP3 | Complex | e4-10 | up | 1,78 | ENSG00000130299 |
| NF564 // ZNF70 | ter. Terminal Exo | e6-8/e9 | down (e6-8) | 1,78 | ENSG00000242852 // ENSG00000249709 |
| ZNF83 | Alter. First Exon | e8/e9 | down (e8) | 1,78 | ENSG00000167766 |
| LPIN1 | ter. Terminal Exo | /e14-19,e21 | own (e14-19,e21-29 | 1,78 | ENSG00000134324 |

| | | | | | |
|---|---|---|---|---|---|
| // LY75 // LY75- | ter. Terminal Exo | /e14-34,e37 | own (e14-34,e37-41 | 1,78 | ENSG00000 054219 // ENSG00000 241399 // ENSG00000 248672 |
| STK4 | ter. Terminal Exo | 5,e6-12,e14 | up (e2) | 1,78 | ENSG00000 101109 |
| SNHG17 | Exon Cassette | e6,e7 | up | 1,78 | ENSG00000 196756 |
| TMEM189-UBE2V | Complex | e14,e15 | up | 1,78 | ENSG00000 124208 // ENSG00000 240849 // ENSG00000 244687 |
| TMEM189-UBE2V | ualy Exclusive Ex | e13/e14 | up (e14) | 1,78 | ENSG00000 124208 // ENSG00000 240849 // ENSG00000 244687 |
| TMEM189-UBE2V | Exon Cassette | e14 | up | 1,78 | ENSG00000 124208 // ENSG00000 240849 // ENSG00000 244687 |
| KCTD17 | Complex | e7-9 | down | 1,78 | ENSG00000 100379 |
| CTNNB1 | Intron Retention | e18-19 | down | 1,78 | ENSG00000 168036 |
| GFM1 | Exon Cassette | e6 | up | 1,78 | ENSG00000 168827 |
| FAM13A | Alter. First Exon | e7-8,e15-23 | p (e2-4,e7-8,e15-23 | 1,78 | ENSG00000 138640 |
| --- | ter. Terminal Exo | e3/e5 | up (e3) | 1,78 | --- |
| NR2F1-AS1 | Exon Cassette | e5,e6-7,e10 | up | 1,78 | ENSG00000 237187 |
| GUSBP9 | Exon Cassette | e25,e26-27 | up | 1,78 | ENSG00000 215630 |
| RXRB | Intron Retention | e6 | up | 1,78 | ENSG00000 204231 |
| WDR46 | Complex | e1,e2-3 | down | 1,78 | ENSG00000 227057 |
| HLA-DQB1 | Alter. First Exon | e1/e2 | down (e1) | 1,78 | ENSG00000 179344 |
| ZNF138 | Exon Cassette | e2 | up | 1,78 | ENSG00000 197008 |
| FBXL6 | Intron Retention | e4 | up | 1,78 | ENSG00000 182325 |

| | | | | | |
|---|---|---|---|---|---|
| MSL3 | Alter. First Exon | e1/e2 | down (e1) | 1,78 | ENSG00000 005302 |
| HAUS7 // TREX2 | ter. Terminal Exc | 9-12,e14-15 | up (e8) | 1,78 | ENSG00000 183479 |
| FPGT-TNNI3K // | ter. Terminal Exc | e22/e23-27 | up (e22) | 1,77 | ENSG00000 116783 // ENSG00000 254685 // ENSG00000 259030 |
| SOAT1 | Exon Cassette | e2,e4 | up | 1,77 | ENSG00000 057252 |
| NFASC | ter. Terminal Exc | 21,e22-25,e2 | (e21,e22-25,e27-29 | 1,77 | ENSG00000 163531 |
| CDH23 | ter. Terminal Exc | 14-28,e30-4 | 13,e14-28,e30-48,e | 1,77 | ENSG00000 107736 |
| RRM1 | Alter. Donor Site | e10 | up | 1,77 | ENSG00000 167325 |
| SDHD | Exon Cassette | e3,e5 | up | 1,77 | ENSG00000 204370 |
| SUV420H1 | Exon Cassette | e6 | up | 1,77 | ENSG00000 110066 |
| CCDC90B | Complex | e1/e2-3 | up (e1) | 1,77 | ENSG00000 137500 |
| // CRYAB // FDX | Alter. First Exon | e5/e6 | up (e6) | 1,77 | ENSG00000 086848 // ENSG00000 255561 |
| DIP2B | Alter. First Exon | e1-16/e17 | down (e1-16) | 1,77 | ENSG00000 066084 |
| TRIM13 | Intron Retention | e1 | up | 1,77 | ENSG00000 204977 |
| ATXN3 | Complex | ,e5-6,e8-11 | own (e2-3,e5-6,e8-1 | 1,77 | ENSG00000 066427 |
| ATXN3 | Complex | e2-10/e12 | down (e2-10) | 1,77 | ENSG00000 066427 |
| 2AP // IGHD // IG | Alter. First Exon | 7,e103/e32-1 | up (e17,e103) | 1,77 | ENSG00000 211896 // ENSG00000 211898 // ENSG00000 213140 |
| CLK3 | Intron Retention | e4,e5 | up | 1,77 | ENSG00000 179335 |
| NGRN // TTLL13 | Complex | e4-7 | down | 1,77 | ENSG00000 182768 // ENSG00000 213471 |
| PITPNC1 | Exon Cassette | e9 | up | 1,77 | ENSG00000 154217 |
| BPTF | Exon Cassette | e7 | down | 1,77 | ENSG00000 171634 |

| | | | | | |
|---|---|---|---|---|---|
| SDF2 | Complex | e1-2 | up | 1,77 | ENSG00000132581 |
| C18orf54 | Alter. Donor Site | e3 | down | 1,77 | ENSG00000166845 |
| ZNF532 | Alter. First Exon | e6-10,e12/e | up (e1,e6-10,e12) | 1,77 | ENSG00000074657 |
| MAU2 | Intron Retention | e11,e12 | up | 1,77 | ENSG00000129933 |
| FCGRT | ter. Terminal Exo | e6/e7-8 | down (e6) | 1,77 | ENSG00000104870 |
| PRMT1 | Alter. First Exon | e1-2/e4 | up (e1-2) | 1,77 | ENSG00000126457 |
| 1S1 // IL4I1 // NU | Alter. First Exon | e2/e3 | down (e2) | 1,77 | ENSG00000104951 // ENSG00000204673 // ENSG00000213024 |
| ZNF816 // ZNF8 | ter. Terminal Exo | e5/e7,e8 | down (e5) | 1,77 | ENSG00000180257 // ENSG00000213801 // ENSG00000221874 |
| LPIN1 | Exon Cassette | e11 | up | 1,77 | ENSG00000134324 |
| ATF2 | ter. Terminal Exo | 0-13,e15,e1 | up (e9) | 1,77 | ENSG00000115966 |
| ORMDL1 | Intron Retention | e2,e3 | up | 1,77 | ENSG00000128699 |
| PGAP1 | Exon Cassette | e3 | up | 1,77 | ENSG00000197121 |
| UBE2C | Alter. First Exon | e1/e2 | down (e1) | 1,77 | ENSG00000175063 |
| HPS4 | Intron Retention | e10,e11 | up | 1,77 | ENSG00000100099 |
| AP1AR | Exon Cassette | e6 | up | 1,77 | ENSG00000138660 |
| --- | Complex | e2,e4 | down | 1,77 | --- |
| SDHAP3 | ter. Terminal Exo | e3/e4,e6-8 | down (e4,e6-8) | 1,77 | ENSG00000185986 |
| HYMAI // PLAGL1 | Alter. Donor Site | e6 | up | 1,77 | ENSG00000118495 |
| SKIV2L | Intron Retention | e7 | up | 1,77 | ENSG00000204351 |
| VPS52 | ter. Terminal Exo | e13/e14-20 | up (e13) | 1,77 | ENSG00000223501 |
| STAG3L4 | Exon Cassette | e3 | up | 1,77 | ENSG00000106610 |
| CCDC146 | Alter. First Exon | e1-6/e7 | down (e1-6) | 1,77 | ENSG00000135205 |

| | | | | | |
|---|---|---|---|---|---|
| ASNS | Exon Cassette | e3 | down | 1,77 | ENSG00000070669 |
| PTPRN2 | Complex | e4/e5 | down (e4) | 1,77 | ENSG00000155093 |
| PTPRN2 | Complex | e1,e4/e5 | down (e1,e4) | 1,77 | ENSG00000155093 |
| IAA1984 // RABL | ter. Terminal Exo | e24/e26 | down (e26) | 1,77 | ENSG00000196642 // ENSG00000213213 |
| SMC1A | Alter. First Exon | e1/e2 | down (e1) | 1,77 | ENSG00000072501 |
| UBE4B | Exon Cassette | e7,e8 | up | 1,76 | ENSG00000130939 |
| EPB41 | ter. Terminal Exo | /e17-18,e20 | up (e14) | 1,76 | ENSG00000159023 |
| PLEKHA1 | Intron Retention | e14,e15 | up | 1,76 | ENSG00000107679 |
| SEPHS1 | ter. Terminal Exo | e6/e7-10 | up (e6) | 1,76 | ENSG00000086475 |
| LDLRAD3 | Exon Cassette | e5 | up | 1,76 | ENSG00000179241 |
| MYL6 | ter. Terminal Exo | e8/e9 | up (e8) | 1,76 | ENSG00000092841 |
| SAV1 | Exon Cassette | e2 | up | 1,76 | ENSG00000151748 |
| GCH1 | ter. Terminal Exo | e5/e6-8 | up (e5) | 1,76 | ENSG00000131979 |
| ATXN3 | Complex | 2-3,e5-11/e1 | down (e2-3,e5-11) | 1,76 | ENSG00000066427 |
| ATXN3 | Complex | e5-6,e8,e10 | wn (e2-3,e5-6,e8,e1 | 1,76 | ENSG00000066427 |
| ATXN3 | Exon Cassette | e3,e5-6,e8,e | down | 1,76 | ENSG00000066427 |
| ATXN3 | Exon Cassette | e2,e4-5 | down | 1,76 | ENSG00000066427 |
| PPCDC | Alter. First Exon | e1-2/e3 | down (e1-2) | 1,76 | ENSG00000138621 |
| DMXL2 | Exon Cassette | e34 | up | 1,76 | ENSG00000104093 |
| NFAT5 | Exon Cassette | e5 | up | 1,76 | ENSG00000102908 |
| SLC12A4 | Complex | e22-29/e29 | up (e22-29) | 1,76 | ENSG00000124067 |
| RFWD3 | Exon Cassette | e2 | up | 1,76 | ENSG00000168411 |
| RABEP1 | Exon Cassette | e17 | up | 1,76 | ENSG00000029725 |
| EPN2 | Exon Cassette | e2 | up | 1,76 | ENSG00000072134 |
| AKAP1 | Alter. First Exon | e2/e5 | down (e2) | 1,76 | ENSG00000121057 |

| | | | | | |
|---|---|---|---|---|---|
| CDRT1 // TRIM16 | ter. Terminal Exc | e11,e13-21 | down (e10) | 1,76 | ENSG00000 221926 // ENSG00000 241322 |
| FAM222B | Complex | e1,e4/e2 | up (e2) | 1,76 | ENSG00000 173065 |
| FAM222B | Exon Cassette | e2 | up | 1,76 | ENSG00000 173065 |
| EXOC7 | Exon Cassette | e7,e8 | down | 1,76 | ENSG00000 182473 |
| AKT2 | Exon Cassette | e3 | up | 1,76 | ENSG00000 105221 |
| CASP8 | Complex | e4-5,e7 | down | 1,76 | ENSG00000 064012 |
| SDC1 | Complex | e4-7 | up | 1,76 | ENSG00000 115884 |
| SMEK2 | Exon Cassette | e10 | up | 1,76 | ENSG00000 138041 |
| ABCB6 // ATG9A | ter. Terminal Exc | 16,e17-30,e | vn (e16,e17-30,e32- | 1,76 | ENSG00000 115657 // ENSG00000 198925 |
| EPB41L1 | Complex | e2,e10/e9 | up (e9) | 1,76 | ENSG00000 088367 |
| SLC4A11 | lter. Acceptor Sit | e15 | up | 1,76 | ENSG00000 088836 |
| MPST | Intron Retention | e3 | up | 1,76 | ENSG00000 128309 |
| FAM211B | Alter. Donor Site | e1 | up | 1,76 | ENSG00000 178026 |
| CRBN | Intron Retention | e6,e7 | up | 1,76 | ENSG00000 113851 |
| SLC4A7 | Exon Cassette | e14 | up | 1,76 | ENSG00000 033867 |
| FAM193A | Exon Cassette | e8 | up | 1,76 | ENSG00000 125386 |
| EXOSC9 | Intron Retention | e10,e11-12 | up | 1,76 | ENSG00000 123737 |
| TJAP1 | Intron Retention | e12 | up | 1,76 | ENSG00000 137221 |
| NCOA7 | Alter. First Exon | e5-14/e15 | down (e5-14) | 1,76 | ENSG00000 111912 |
| MYB | Exon Cassette | e10,e12 | up | 1,76 | ENSG00000 118513 |
| ELN | Complex | 18-21,e24,e2 | up | 1,76 | ENSG00000 049540 |
| GTF2I | ter. Terminal Exc | e11/e12-35 | down (e12-35) | 1,76 | ENSG00000 077809 |
| VPS13D | Complex | e40/e41 | down (e40) | 1,75 | ENSG00000 048707 |
| HP1BP3 | lter. Acceptor Sit | e13 | up | 1,75 | ENSG00000 127483 |

| | | | | | |
|---|---|---|---|---|---|
| FAF1 | Exon Cassette | e20 | up | 1,75 | ENSG00000 185104 |
| SEC24C | Intron Retention | e19 | up | 1,75 | ENSG00000 176986 |
| EPC1 | Complex | e2-3 | down | 1,75 | ENSG00000 120616 |
| RHGAP19 // SLI | ter. Terminal Exo | 20/e21,e22-2 | up (e21,e22-28) | 1,75 | ENSG00000 187122 // ENSG00000 213390 |
| DIXDC1 | Alter. First Exon | e10-12/e13 | up (e10-12) | 1,75 | ENSG00000 150764 |
| PARPBP | Exon Cassette | e14 | up | 1,75 | ENSG00000 185480 |
| STAT2 | Complex | e6-12 | down | 1,75 | ENSG00000 170581 |
| GNS | Exon Cassette | e2 | down | 1,75 | ENSG00000 135677 |
| EXOSC8 | Intron Retention | e4 | up | 1,75 | ENSG00000 120699 |
| ATXN3 | Complex | e2-5/e12 | down (e2-5) | 1,75 | ENSG00000 066427 |
| ATXN3 | Exon Cassette | e2-5 | down | 1,75 | ENSG00000 066427 |
| MAPKBP1 | Complex | e29-30 | down | 1,75 | ENSG00000 137802 |
| TAOK2 | ter. Terminal Exo | e17/e18-20 | up (e17) | 1,75 | ENSG00000 149930 |
| C17orf76-AS1 | Complex | e3,e4 | up | 1,75 | ENSG00000 175061 |
| ZNF565 | Alter. First Exon | e1,e4/e3 | up (e1,e4) | 1,75 | ENSG00000 196357 |
| ZNF565 | Exon Cassette | e4 | up | 1,75 | ENSG00000 196357 |
| BOLA3-AS1 | Intron Retention | e1,e2 | down | 1,75 | ENSG00000 225439 |
| CIAO1 | Intron Retention | e6 | up | 1,75 | ENSG00000 144021 |
| MAP4K4 | Exon Cassette | e18 | down | 1,75 | ENSG00000 071054 |
| --- | Exon Cassette | e4-9 | down | 1,75 | --- |
| --- | Exon Cassette | e2-9 | down | 1,75 | --- |
| PUM2 | Exon Cassette | e14 | up | 1,75 | ENSG00000 055917 |
| GTF3C2 | Exon Cassette | e3 | up | 1,75 | ENSG00000 115207 |
| ATF2 | ter. Terminal Exo | e11-13,e15 | up (e9) | 1,75 | ENSG00000 115966 |
| PAX3 | ter. Terminal Exo | e6,e7-8,e10 | up (e6,e7-8,e10-11) | 1,75 | ENSG00000 135903 |
| NAPB | Exon Cassette | e4 | down | 1,75 | ENSG00000 125814 |

| | | | | | |
|---|---|---|---|---|---|
| E1 // NFS1 // RB | Exon Cassette | e22,e23 | up | 1,75 | ENSG00000 214078 // ENSG00000 244005 // ENSG00000 244462 |
| RPL15 | Complex | e1 | down | 1,75 | ENSG00000 174748 |
| RPL15 | Complex | e1/e2 | down (e1) | 1,75 | ENSG00000 174748 |
| PHF15 | Complex | e12 | up | 1,75 | ENSG00000 043143 |
| PHF15 | Complex | e12,e13 | up | 1,75 | ENSG00000 043143 |
| FAM135A | Alter. First Exon | -10,e12-15/e | up (e4-10,e12-15) | 1,75 | ENSG00000 082269 |
| CDK19 | Complex | e3-4 | down | 1,75 | ENSG00000 155111 |
| NUDT1 | Alter. First Exon | e1,e4/e2-3 | up (e2-3) | 1,75 | ENSG00000 106268 |
| FOXP2 | Complex | e7/e8-9 | down (e8-9) | 1,75 | ENSG00000 128573 |
| FOXP2 | Exon Cassette | e8,e9 | down | 1,75 | ENSG00000 128573 |
| FOXP2 | Exon Cassette | e19-20 | down | 1,75 | ENSG00000 128573 |
| ERCC6L2 | Exon Cassette | e5-6 | up | 1,75 | ENSG00000 182150 |
| LPAR1 | Alter. First Exon | e3/e6-8 | up (e3) | 1,75 | ENSG00000 198121 |
| AKNA | Alter. First Exon | e3/e5 | down (e3) | 1,75 | ENSG00000 106948 |
| VPS13D | Exon Cassette | e40 | down | 1,74 | ENSG00000 048707 |
| EPS15 | Alter. First Exon | -8,e10-13/e | up (e1-8,e10-13) | 1,74 | ENSG00000 085832 |
| MRPL55 | Complex | e1,e3 | up | 1,74 | ENSG00000 162910 |
| TGLF9P // PAR | ter. Terminal Ex | -25/e30,e31 | up (e18-25) | 1,74 | ENSG00000 227345 |
| SORBS1 | Exon Cassette | e30,e31 | up | 1,74 | ENSG00000 095637 |
| DGKZ | Alter. First Exon | e1-29/e30 | up (e1-29) | 1,74 | ENSG00000 149091 |
| DGKZ | Alter. First Exon | -9,e11-29/e | up (e2-9,e11-29) | 1,74 | ENSG00000 149091 |
| TTC9C | Exon Cassette | e2 | up | 1,74 | ENSG00000 162222 |
| FOXRED1 | Alter. Donor Site | e1 | up | 1,74 | ENSG00000 110074 |

| | | | | | |
|---|---|---|---|---|---|
| IGEF25 // SLC26 | Complex | e31,e32-33 | up | 1,74 | ENSG00000 135502 // ENSG00000 240771 |
| P2RX7 | ualy Exclusive Ex | e5/e6 | down (e5) | 1,74 | ENSG00000 089041 |
| TNFRSF1A | Alter. First Exon | 3,e5-6,e8-1 | p (e1,e3,e5-6,e8-11 | 1,74 | ENSG00000 067182 |
| TNFRSF1A | Alter. First Exon | 3,e5-6,e8-11 | up (e1-3,e5-6,e8-11 | 1,74 | ENSG00000 067182 |
| // TRAV12-1 // T | ter. Terminal Exo | 44,e152-15 | n (e35,e144,e152-1 | 1,74 | ENSG00000 211785 // ENSG00000 211800 // ENSG00000 229164 |
| ATXN3 | Exon Cassette | e2-3,e5 | down | 1,74 | ENSG00000 066427 |
| SLC7A5P1 | Alter. First Exon | e1/e2 | up (e1) | 1,74 | ENSG00000 260727 |
| ARHGAP44 | Exon Cassette | e18-19 | down | 1,74 | ENSG00000 006740 |
| C17orf76-AS1 | Complex | e3 | up | 1,74 | ENSG00000 175061 |
| WSB1 | Complex | e6-7 | up | 1,74 | ENSG00000 109046 |
| ACSF2 | Alter. First Exon | e1,e3-11/e12 | down (e1,e3-11) | 1,74 | ENSG00000 167107 |
| BRSK1 | Alter. First Exon | e3-11/e12 | down (e3-11) | 1,74 | ENSG00000 160469 |
| ZNF573 | Exon Cassette | e6-7 | up | 1,74 | ENSG00000 189144 |
| 1S1 // IL4I1 // NU | Alter. First Exon | e1/e3,e4 | up (e3,e4) | 1,74 | ENSG00000 104951 // ENSG00000 204673 // ENSG00000 213024 |
| CEP68 | Complex | e1-2 | down | 1,74 | ENSG00000 011523 |
| NYAP2 | Exon Cassette | e4-5 | up | 1,74 | ENSG00000 144460 |
| EIF2B4 | Intron Retention | e9 | up | 1,74 | ENSG00000 115211 |
| PRV1 // PCBP1-A | ter. Terminal Exo | 1/e12,e14-1 | up (e11) | 1,74 | ENSG00000 179818 // ENSG00000 244617 |
| EPB41L1 | Complex | e2-10/e4,e9 | up (e4,e9) | 1,74 | ENSG00000 088367 |
| NELFCD | Alter. Donor Site | e7 | up | 1,74 | ENSG00000 101158 |

| | | | | | |
|---|---|---|---|---|---|
| BID | Complex | e2/e4-6 | up (e4-6) | 1,74 | ENSG00000 015475 |
| PDE12 | Intron Retention | e2 | up | 1,74 | ENSG00000 174840 |
| TSC22D2 | Exon Cassette | e3,e4 | up | 1,74 | ENSG00000 196428 |
| EOGT | Complex | e16-20 | up | 1,74 | ENSG00000 163378 |
| :OX17 // POPDC | Exon Cassette | e7,e8-10 | down | 1,74 | ENSG00000 121577 // ENSG00000 138495 |
| FNIP2 | Intron Retention | e7 | up | 1,74 | ENSG00000 052795 |
| GNPDA1 | Complex | e1,e2 | up | 1,74 | ENSG00000 113552 |
| ZNF76 | ter. Terminal Exo | e10/e11-15 | up (e10) | 1,74 | ENSG00000 065029 |
| ZNF76 | ter. Terminal Exo | e11,e12,e1 | up (e10) | 1,74 | ENSG00000 065029 |
| OPRM1 | Exon Cassette | e8-9,e12 | down | 1,74 | ENSG00000 112038 |
| IICAL1 // ZBTB2 | ter. Terminal Exo | e2/e4,e5-8 | up (e2) | 1,74 | ENSG00000 112365 // ENSG00000 135596 |
| ATAT1 | ter. Terminal Exo | 1/e12,e13- | down (e11) | 1,74 | ENSG00000 137343 |
| IKBKB | Exon Cassette | e16 | down | 1,74 | ENSG00000 104365 |
| CREB3 | Intron Retention | e3 | up | 1,74 | ENSG00000 107175 |
| SMC2 | Exon Cassette | e25 | up | 1,74 | ENSG00000 136824 |
| --- | Exon Cassette | e12 | up | 1,74 | --- |
| ARMCX2 | Alter. Donor Site | e4 | up | 1,74 | ENSG00000 184867 |
| ADC | Alter. First Exon | e1-2/e5 | down (e1-2) | 1,73 | ENSG00000 142920 |
| AMY2B // RNPC3 | Alter. First Exon | 17-18/e19,e2 | up (e17-18) | 1,73 | ENSG00000 185946 // ENSG00000 240038 |
| DDX20 | Intron Retention | e9 | up | 1,73 | ENSG00000 064703 |
| CD46 | Exon Cassette | e13 | up | 1,73 | ENSG00000 117335 |
| MARK1 | Exon Cassette | e17 | up | 1,73 | ENSG00000 116141 |
| MRPL55 | Exon Cassette | e2-3 | up | 1,73 | ENSG00000 162910 |

| | | | | | |
|---|---|---|---|---|---|
| NID1 | Exon Cassette | e10-12 | up | 1,73 | ENSG00000116962 |
| WAC | Intron Retention | e14 | up | 1,73 | ENSG00000095787 |
| NRP1 | Exon Cassette | e4 | up | 1,73 | ENSG00000099250 |
| HMBS | Intron Retention | e11 | up | 1,73 | ENSG00000256269 |
| H19 | Complex | e5 | down | 1,73 | ENSG00000130600 |
| DKK3 | Complex | e5,e7-10 | up | 1,73 | ENSG00000050165 |
| ABCC8 | Exon Cassette | e2 | up | 1,73 | ENSG00000006071 |
| SLC37A4 | Complex | e3 | up | 1,73 | --- |
| WNK1 | Exon Cassette | e29,e30 | down | 1,73 | ENSG00000060237 |
| FAM66C | Exon Cassette | e4 | up | 1,73 | ENSG00000226711 |
| TNFRSF1A | Alter. First Exon | ,e4-6,e8-11/ | up (e1,e4-6,e8-11) | 1,73 | ENSG00000067182 |
| CALCOCO1 | Intron Retention | e14 | up | 1,73 | ENSG00000012822 |
| ANKRD10 | Exon Cassette | e4 | up | 1,73 | ENSG00000088448 |
| IRF9 // RNF31 | ter. Terminal Exc | e28/e30-31 | up (e30-31) | 1,73 | ENSG00000092098 // ENSG00000213928 |
| TIMM9 | Exon Cassette | e2-3 | up | 1,73 | ENSG00000100575 |
| ATXN3 | Exon Cassette | e2,e5 | down | 1,73 | ENSG00000066427 |
| PRC1 | Exon Cassette | e14 | up | 1,73 | ENSG00000198901 |
| NDRG4 | Complex | /e9-11,e13- | up (e8) | 1,73 | ENSG00000103034 |
| ACSF2 | Alter. First Exon | e1-11/e12 | down (e1-11) | 1,73 | ENSG00000167107 |
| NARF | Exon Cassette | e11 | up | 1,73 | ENSG00000141562 |
| SRSF1 | Intron Retention | e3 | up | 1,73 | ENSG00000136450 |
| EXOC7 | ter. Terminal Exc | 5-7,e9-13,e1 | up (e4) | 1,73 | ENSG00000182473 |
| EXOC7 | Intron Retention | e11 | up | 1,73 | ENSG00000182473 |

| | | | | | |
|---|---|---|---|---|---|
| HERP // MED26 | Exon Cassette | e35-37 | up | 1,73 | ENSG00000085872 // ENSG00000105085 // ENSG00000127526 // ENSG00000269058 |
| BCL2L11 | Complex | 2/e3,e11,e1 | up (e3,e11,e13) | 1,73 | ENSG00000153094 |
| BCL2L11 | Exon Cassette | e3,e11 | up | 1,73 | ENSG00000153094 |
| SPAG16 | Complex | e2,e3 | up | 1,73 | ENSG00000144451 |
| STK11IP | Intron Retention | e3 | up | 1,73 | ENSG00000144589 |
| PCGF1 | Alter. First Exon | e1/e2 | down (e1) | 1,73 | ENSG00000115289 |
| SNRNP200 | Exon Cassette | e44 | up | 1,73 | ENSG00000144028 |
| BID | Exon Cassette | e6 | up | 1,73 | ENSG00000015475 |
| RPL15 | Alter. Donor Site | e1 | down | 1,73 | ENSG00000174748 |
| SAP30L | Exon Cassette | e3 | up | 1,73 | ENSG00000164576 |
| FBN2 | ter. Terminal Exo | 4/e15,e17-7 | up (e15,e17-73) | 1,73 | ENSG00000138829 |
| C7orf55-LUC7L2 | Exon Cassette | e7 | up | 1,73 | ENSG00000146963 // ENSG00000164898 // ENSG00000269955 |
| NSMF | Exon Cassette | e5 | up | 1,73 | ENSG00000165802 |
| ZNF711 | Intron Retention | e11 | up | 1,73 | ENSG00000147180 |
| MAP7D2 | Exon Cassette | e6 | down | 1,73 | ENSG00000184368 |
| HS6ST2 | Alter. First Exon | e1-2/e4 | up (e1-2) | 1,73 | ENSG00000171004 |
| FAM3A | Exon Cassette | e3 | down | 1,73 | ENSG00000071889 |
| PTPRF | Exon Cassette | e25 | down | 1,72 | ENSG00000142949 |
| CHTOP | ter. Terminal Exo | e3/e6 | up (e3) | 1,72 | ENSG00000160679 |
| HP1BP3 | Complex | e3/e4 | up (e3) | 1,72 | ENSG00000127483 |
| MSANTD2 | Exon Cassette | e2 | down | 1,72 | ENSG00000120458 |

| | | | | | |
|---|---|---|---|---|---|
| VEZT | Exon Cassette | e7 | up | 1,72 | ENSG00000 028203 |
| HDAC7 | Complex | e11-28 | down | 1,72 | ENSG00000 061273 |
| 2AP // IGHD // IG | ter. Terminal Exo | 03,e130-13 | (e75,e103,e130-13 | 1,72 | ENSG00000 211896 // ENSG00000 211898 // ENSG00000 213140 |
| 2AP // IGHD // IG | ter. Terminal Exo | /e103,e130 | up (e103,e130-133) | 1,72 | ENSG00000 211896 // ENSG00000 211898 // ENSG00000 213140 |
| HERC2 | Alter. First Exon | 2,e4,e6-49/ | down (e1-2,e4,e6-49 | 1,72 | ENSG00000 128731 |
| PKM | Complex | e2,e5-6 | up | 1,72 | ENSG00000 067225 |
| JPH3 | ter. Terminal Exo | e3/e4,e6-8 | up (e3) | 1,72 | ENSG00000 154118 |
| CYBA | Complex | e1/e2,e3-6 | down (e1) | 1,72 | ENSG00000 051523 |
| CYBA | Complex | e1/e2-6 | down (e1) | 1,72 | ENSG00000 051523 |
| PHF12 | Complex | e3-4 | up | 1,72 | ENSG00000 109118 |
| ANKRD12 | ter. Terminal Exo | e10/e12-16 | up (e12-16) | 1,72 | ENSG00000 101745 |
| NEDD4L | Complex | e1,e11/e7 | down (e7) | 1,72 | ENSG00000 049759 |
| SS18 | Exon Cassette | e3-4 | up | 1,72 | ENSG00000 141380 |
| MFSD12 | Complex | e1-10 | up | 1,72 | ENSG00000 161091 |
| TTPAL | Alter. First Exon | e1-2/e3 | down (e1-2) | 1,72 | ENSG00000 124120 |
| DBNDD2 // SYS1 | Complex | e7 | down | 1,72 | ENSG00000 204070 // ENSG00000 244274 |
| --- | Alter. First Exon | e1/e2 | down (e1) | 1,72 | --- |
| TCEA2 | Complex | e2-4 | up | 1,72 | ENSG00000 171703 |
| SNX5 | Alter. First Exon | e2/e3 | down (e2) | 1,72 | ENSG00000 089006 |
| HAND2-AS1 | Exon Cassette | e4 | up | 1,72 | ENSG00000 237125 |
| PPM1K | Exon Cassette | e4 | up | 1,72 | ENSG00000 163644 |

| | | | | | |
|---|---|---|---|---|---|
| DDX41 | Intron Retention | e4 | up | 1,72 | ENSG00000183258 |
| MYB | Complex | e10-12 | up | 1,72 | ENSG00000118513 |
| PNISR | Intron Retention | e11 | up | 1,72 | ENSG00000132424 |
| CCHCR1 | ter. Terminal Exo | e16/e17-19 | up (e16) | 1,72 | ENSG00000204536 |
| FTSJ2 | Complex | e3,e4 | down | 1,72 | ENSG00000122687 |
| GTF2IRD2 | ter. Terminal Exo | e4,e5-8,e10 | up (e3) | 1,72 | ENSG00000196275 |
| ABCB1 | Alter. First Exon | e1-21/e22 | up (e1-21) | 1,72 | ENSG00000085563 |
| ASAH1 | Alter. First Exon | e1/e2 | down (e1) | 1,72 | ENSG00000104763 |
| ZNF706 | Exon Cassette | e3,e4 | up | 1,72 | ENSG00000120963 |
| TMEM234 | Exon Cassette | e4 | down | 1,71 | ENSG00000160055 |
| PLEKHA1 | Exon Cassette | e15 | up | 1,71 | ENSG00000107679 |
| BBIP1 | Exon Cassette | e4-5 | up | 1,71 | ENSG00000214413 |
| NSMCE4A | Intron Retention | e1,e2 | up | 1,71 | ENSG00000107672 |
| OAT | lter. Acceptor Sit | e5 | up | 1,71 | ENSG00000065154 |
| DDB2 | Exon Cassette | e6 | down | 1,71 | ENSG00000134574 |
| CTNND1 // TMX2 | lter. Acceptor Sit | e3 | down | 1,71 | ENSG00000198561 // ENSG00000213593 |
| ACAD8 | Intron Retention | e5 | up | 1,71 | ENSG00000151498 |
| RNH1 | Complex | e3,e4 | up | 1,71 | ENSG00000023191 |
| SLC37A4 | Exon Cassette | e2 | down | 1,71 | --- |
| DCP1B | Exon Cassette | e5 | up | 1,71 | ENSG00000151065 |
| CALCOCO1 | Intron Retention | e14,e15 | up | 1,71 | ENSG00000012822 |
| SUCLA2 | Alter. First Exon | e1-2/e3 | up (e1-2) | 1,71 | ENSG00000136143 |
| KTN1 | Exon Cassette | e44 | up | 1,71 | ENSG00000126777 |
| ATXN3 | Complex | 3-6,e8-10/e1 | down (e3-6,e8-10) | 1,71 | ENSG00000066427 |
| CCNDBP1 | Intron Retention | e2,e3 | up | 1,71 | ENSG00000166946 |

| | | | | | |
|---|---|---|---|---|---|
| EDC4 | Intron Retention | e8 | up | 1,71 | ENSG00000038358 |
| PDXDC2P | Exon Cassette | e24 | down | 1,71 | ENSG00000196696 |
| C17orf76-AS1 | Complex | e2,e3-4 | up | 1,71 | ENSG00000175061 |
| C17orf76-AS1 | Exon Cassette | e3 | up | 1,71 | ENSG00000175061 |
| ZNF254 | Alter. First Exon | 2,e4,e11-12 | up (e1-2,e4,e11-12) | 1,71 | ENSG00000213096 |
| ZNF507 | ter. Terminal Exo | e4/e5-7 | up (e4) | 1,71 | ENSG00000168813 |
| ZNF551 | Exon Cassette | e2 | up | 1,71 | ENSG00000204519 |
| CEP68 | Exon Cassette | e2 | down | 1,71 | ENSG00000011523 |
| CCDC150 | Alter. First Exon | e19-25/e26 | down (e19-25) | 1,71 | ENSG00000144395 |
| PREPL | Complex | e1,e2 | up | 1,71 | ENSG00000138078 |
| CCT4 | Exon Cassette | e2 | up | 1,71 | ENSG00000115484 |
| PRMT2 | Exon Cassette | e2 | up | 1,71 | ENSG00000160310 |
| ZBTB21 | Exon Cassette | e3 | up | 1,71 | ENSG00000173276 |
| CMTM8 | Exon Cassette | e2 | up | 1,71 | ENSG00000170293 |
| FIP1L1 | lter. Acceptor Sit | e17 | up | 1,71 | ENSG00000145216 |
| REST | Exon Cassette | e5,e8 | up | 1,71 | ENSG00000084093 |
| ME1 | Exon Cassette | e2 | down | 1,71 | ENSG00000065833 |
| CDK19 | Exon Cassette | e3 | down | 1,71 | ENSG00000155111 |
| 5-SAPCD1 // SAI | lter. Acceptor Sit | e27 | down | 1,71 | ENSG00000228727 // ENSG00000255152 |
| SDK1 | Alter. First Exon | e1-33/e34 | up (e1-33) | 1,71 | ENSG00000146555 |
| INSIG1 | Exon Cassette | e6 | up | 1,71 | ENSG00000186480 |
| PRKAR1B | Alter. First Exon | ,e6,e8-11/e | down (e4,e6,e8-11) | 1,71 | ENSG00000188191 |
| YTHDF2 | Intron Retention | e1 | up | 1,7 | ENSG00000198492 |
| DMAP1 | lter. Acceptor Sit | e7 | down | 1,7 | ENSG00000178028 |
| PIAS3 | Intron Retention | e8 | up | 1,7 | ENSG00000131788 |

| | | | | | |
|---|---|---|---|---|---|
| EXOSC10 | Intron Retention | e16 | up | 1,7 | ENSG00000171824 |
| S3MT // C10orf3 | Alter. First Exon | e7/e8 | up (e7) | 1,7 | ENSG00000166275 // ENSG00000214435 |
| ZNF248 | Intron Retention | e7 | down | 1,7 | ENSG00000198105 |
| PPP3CB | Exon Cassette | e2 | up | 1,7 | ENSG00000107758 |
| DNAJC24 | ter. Terminal Ex | e3/e4,e5 | up (e4,e5) | 1,7 | ENSG00000170946 |
| MUS81 | Intron Retention | e12 | up | 1,7 | ENSG00000172732 |
| TBCEL | Exon Cassette | e2,e3 | up | 1,7 | ENSG00000154114 |
| TBRG1 | Exon Cassette | e4 | up | 1,7 | ENSG00000154144 |
| NHG1 // SNORD | Intron Retention | e7 | up | 1,7 | ENSG00000207487 // ENSG00000255717 |
| NUMA1 | Alter. First Exon | e2-3/e7 | up (e2-3) | 1,7 | ENSG00000137497 |
| SLC6A15 | ter. Terminal Ex | e5/e6,e7-12 | down (e5) | 1,7 | ENSG00000072041 |
| ADSSL1 | Alter. First Exon | e2-3/e4 | up (e2-3) | 1,7 | ENSG00000185100 |
| ATXN3 | Complex | e5-6,e8-11/ | down (e3,e5-6,e8-11 | 1,7 | ENSG00000066427 |
| ATXN3 | Complex | 4,e6,e8-10/ | down (e2-4,e6,e8-10 | 1,7 | ENSG00000066427 |
| TCF12 | Alter. First Exon | e2-4/e17-20 | up (e2-4) | 1,7 | ENSG00000140262 |
| ZSCAN32 | Exon Cassette | e2 | down | 1,7 | ENSG00000140987 |
| DNAH3 | ter. Terminal Ex | 5/e46,e47- | down (e46,e47-66) | 1,7 | ENSG00000158486 |
| SGSM2 | Intron Retention | e5 | up | 1,7 | ENSG00000141258 |
| SPHK1 | Alter. First Exon | e4-8/e6 | down (e4-8) | 1,7 | ENSG00000176170 |
| MIEN1 | Intron Retention | e3 | up | 1,7 | ENSG00000141741 |
| EXOC7 | ter. Terminal Ex | 4/e5-7,e9-2 | up (e4) | 1,7 | ENSG00000182473 |
| EXOC7 | ter. Terminal Ex | 5-7,e9-13,e1 | up (e4) | 1,7 | ENSG00000182473 |
| ZNF254 | Alter. First Exon | 1-2,e11-12/e | up (e1-2,e11-12) | 1,7 | ENSG00000213096 |
| TTC31 | Intron Retention | e11 | up | 1,7 | ENSG00000115282 |

| | | | | | |
|---|---|---|---|---|---|
| SDC1 | Complex | e4-5,e7 | up | 1,7 | ENSG00000115884 |
| ERCC3 | Intron Retention | e4 | up | 1,7 | ENSG00000163161 |
| CACNB4 | Exon Cassette | e12 | down | 1,7 | ENSG00000182389 |
| ICA1L | Intron Retention | e16 | down | 1,7 | ENSG00000163596 |
| PLCB1 | Alter. First Exon | e1/e2,e4-5 | down (e2,e4-5) | 1,7 | ENSG00000182621 |
| BTG3 | Exon Cassette | e4 | down | 1,7 | ENSG00000154640 |
| ZBTB21 | Exon Cassette | e3,e4 | up | 1,7 | ENSG00000173276 |
| FLNB | Alter. First Exon | e3-26/e28 | down (e3-26) | 1,7 | ENSG00000136068 |
| PLS1 | Alter. First Exon | e1/e2,e5 | down (e2,e5) | 1,7 | ENSG00000120756 |
| WHSC1 | Exon Cassette | e5 | down | 1,7 | ENSG00000109685 |
| UGDH | Exon Cassette | e2 | down | 1,7 | ENSG00000109814 |
| SLC1A3 | Complex | e2,e4-5 | up | 1,7 | ENSG00000079215 |
| NR2F1-AS1 | Exon Cassette | e5-7 | up | 1,7 | ENSG00000237187 |
| FAM13B | Exon Cassette | e14 | up | 1,7 | ENSG00000031003 |
| TFAP2A-AS1 | Intron Retention | e3 | down | 1,7 | ENSG00000229950 |
| MYLIP | Exon Cassette | e2 | up | 1,7 | ENSG00000007944 |
| TRAF3IP2-AS1 | Exon Cassette | e6,e7 | down | 1,7 | ENSG00000231889 |
| WASF1 | Exon Cassette | e4 | up | 1,7 | ENSG00000112290 |
| GTF2H4 // VARS2 | Intron Retention | e25 | up | 1,7 | ENSG00000137411 // ENSG00000213780 |
| PPP1R11 | Exon Cassette | e2 | up | 1,7 | ENSG00000204619 |
| TRIM27 | Intron Retention | e7 | up | 1,7 | ENSG00000204713 |
| GTPBP10 | Exon Cassette | e4 | up | 1,7 | ENSG00000105793 |
| PRKAR1B | Alter. First Exon | 1-6,e8-11/e1 | down (e1-6,e8-11) | 1,7 | ENSG00000188191 |
| ETV1 | Alter. First Exon | e4,e6-8/e10 | up (e4,e6-8) | 1,7 | ENSG00000006468 |
| COA1 | ter. Terminal Exo | e9/e10 | up (e9) | 1,7 | ENSG00000106603 |

| | | | | | |
|---|---|---|---|---|---|
| MAGED1 | Intron Retention | e11 | up | 1,7 | ENSG00000179222 |
| NF182 // ZNF63 | Exon Cassette | e16 | up | 1,7 | ENSG00000147118 // ENSG00000221994 |
| FAM104B | Alter. Donor Site | e1 | up | 1,7 | ENSG00000182518 |
| DDX20 | Intron Retention | e8 | up | 1,69 | ENSG00000064703 |
| TTLL7 | Exon Cassette | e25-26 | up | 1,69 | ENSG00000137941 |
| MRPL55 | Complex | e2-4 | up | 1,69 | ENSG00000162910 |
| PPP3CB | Exon Cassette | e16 | down | 1,69 | ENSG00000107758 |
| KCNMA1 | Alter. First Exon | e8,e10-15/e | down (e1,e8,e10-15 | 1,69 | ENSG00000156113 |
| TSPAN4 | Alter. First Exon | e1/e3-4 | down (e1) | 1,69 | ENSG00000214063 |
| PPHLN1 | Exon Cassette | e5 | up | 1,69 | ENSG00000134283 |
| 3HDM2 // STAC | Exon Cassette | e33 | down | 1,69 | ENSG00000179912 // ENSG00000185482 |
| RBM26 | Intron Retention | e21 | up | 1,69 | ENSG00000139746 |
| PRPF39 | Exon Cassette | e4 | up | 1,69 | ENSG00000185246 |
| APOPT1 // KLC1 | Alter. Donor Site | e18 | up | 1,69 | ENSG00000126214 // ENSG00000256053 |
| ACTN1 | Alter. First Exon | 1,e4-13/e1 | down (e1,e4-13) | 1,69 | ENSG00000072110 |
| ACTN1 | Alter. First Exon | e2-13/e14 | down (e2-13) | 1,69 | ENSG00000072110 |
| ACTN1 | Alter. First Exon | e3-13/e14 | down (e3-13) | 1,69 | ENSG00000072110 |
| ATXN3 | Exon Cassette | e2-3 | down | 1,69 | ENSG00000066427 |
| 2AP // IGHD // IG | Alter. First Exon | e24-25/e71 | up (e24-25) | 1,69 | ENSG00000211896 // ENSG00000211898 // ENSG00000213140 |

| | | | | | |
|---|---|---|---|---|---|
| 2AP // IGHD // IG | Alter. First Exon | 17,e101/e7 | up (e17,e101) | 1,69 | ENSG00000211896 // ENSG00000211898 // ENSG00000213140 |
| 2AP // IGHD // IG | Alter. First Exon | 4-25/e71,e1 | up (e24-25) | 1,69 | ENSG00000211896 // ENSG00000211898 // ENSG00000213140 |
| 2AP // IGHD // IG | Alter. First Exon | 17/e71,e10 | up (e17) | 1,69 | ENSG00000211896 // ENSG00000211898 // ENSG00000213140 |
| 2AP // IGHD // IG | Alter. First Exon | e17/e71 | up (e17) | 1,69 | ENSG00000211896 // ENSG00000211898 // ENSG00000213140 |
| CTDSPL2 | Complex | e1 | up | 1,69 | ENSG00000137770 |
| USP3 | Alter. First Exon | e1-2/e3 | up (e1-2) | 1,69 | ENSG00000140455 |
| COMMD4 | Intron Retention | e2 | up | 1,69 | ENSG00000140365 |
| ZFAND6 | Alter. First Exon | e1,e7/e4 | up (e1,e7) | 1,69 | ENSG00000086666 |
| --- | Intron Retention | e2 | up | 1,69 | --- |
| WSB1 | Alter. First Exon | e1,e4-5/e7 | down (e1,e4-5) | 1,69 | ENSG00000109046 |
| SPHK1 | Alter. First Exon | e1-3,e7-8/e6 | down (e1-3,e7-8) | 1,69 | ENSG00000176170 |
| RAD51D // RFFL | Exon Cassette | e6,e7 | down | 1,69 | ENSG00000092871 // ENSG00000185379 |
| ZNF507 | ter. Terminal Exo | e4/e5,e6-7 | up (e4) | 1,69 | ENSG00000168813 |
| DMKN | Complex | e23,e24 | down | 1,69 | ENSG00000161249 |
| ZNF772 | Exon Cassette | e3-4 | up | 1,69 | ENSG00000197128 |
| AUP1 | Intron Retention | e4 | up | 1,69 | ENSG00000115307 |
| PLCB1 | Alter. First Exon | 2,e4-5/e6,e | down (e2,e4-5) | 1,69 | ENSG00000182621 |

| | | | | | |
|---|---|---|---|---|---|
| BID | Alter. First Exon | e1/e2,e4-7 | up (e2,e4-7) | 1,69 | ENSG00000 015475 |
| FLNB | Alter. First Exon | 1-2,e4-26/e2 | down (e1-2,e4-26) | 1,69 | ENSG00000 136068 |
| CYP2U1 | ter. Terminal Exo | e2/e3-7 | down (e3-7) | 1,69 | ENSG00000 155016 |
| LINC01001 | Exon Cassette | e13 | up | 1,69 | ENSG00000 230724 |
| DBN1 | Intron Retention | e6 | up | 1,69 | ENSG00000 113758 |
| PPARD | ter. Terminal Exo | e8/e9-10 | up (e9-10) | 1,69 | ENSG00000 112033 |
| ICAL1 // ZBTB2 | ter. Terminal Exo | e2/e3,e4-8 | up (e2) | 1,69 | ENSG00000 112365 // ENSG00000 135596 |
| CEP85L | Alter. First Exon | e1-2/e4 | down (e4) | 1,69 | ENSG00000 111860 |
| CEP85L | Alter. First Exon | e3/e4 | down (e4) | 1,69 | ENSG00000 111860 |
| HYMAI // PLAGL1 | Complex | e7/e8-10 | up (e8-10) | 1,69 | ENSG00000 118495 |
| HLA-E | Complex | e2-3 | down | 1,69 | ENSG00000 204592 |
| ZMIZ2 | Complex | -8,e10-12,e | down | 1,69 | ENSG00000 122515 |
| CCDC146 | Complex | e3,e4 | up | 1,69 | ENSG00000 135205 |
| ADAM22 | ter. Terminal Exo | e29/e31-32 | up (e29) | 1,69 | ENSG00000 008277 |
| NAA38 | Intron Retention | e1 | up | 1,69 | ENSG00000 128534 |
| --- | Alter. First Exon | 22-23,e36/e3 | up (e22-23,e36) | 1,69 | --- |
| MAGI2 | Alter. First Exon | 1,e3,e6-7/e8 | up (e1,e3,e6-7) | 1,69 | ENSG00000 187391 |
| ESYT2 | Exon Cassette | e16 | down | 1,69 | ENSG00000 117868 |
| OMMD5 // ZNF25 | Complex | e6 | down | 1,69 | ENSG00000 170619 // ENSG00000 196150 |
| ERCC6L2 | Exon Cassette | e6 | up | 1,69 | ENSG00000 182150 |
| GEMIN8 | Alter. First Exon | e1/e2 | down (e1) | 1,69 | ENSG00000 046647 |
| AIFM1 | Exon Cassette | e2 | up | 1,69 | ENSG00000 156709 |
| PRPF38B | ter. Terminal Exo | e5-6/e8-9 | up (e5-6) | 1,68 | ENSG00000 134186 |
| DIP2C | ter. Terminal Exo | /e31-35,e37 | up (e30) | 1,68 | ENSG00000 151240 |

| | | | | | |
|---|---|---|---|---|---|
| ZNF37BP | Intron Retention | e2 | up | 1,68 | ENSG00000234420 |
| DIXDC1 | Alter. First Exon | e3-7/e8 | down (e3-7) | 1,68 | ENSG00000150764 |
| STT3A | Exon Cassette | e2-3 | up | 1,68 | ENSG00000134910 |
| PIDD | Intron Retention | e3,e4 | down | 1,68 | ENSG00000177595 |
| NUMA1 | Complex | e26,e27-32 | down | 1,68 | ENSG00000137497 |
| NUMA1 | Exon Cassette | e26 | up | 1,68 | ENSG00000137497 |
| ETS1 | Complex | e9-10 | up | 1,68 | ENSG00000134954 |
| SLC38A1 | Alter. First Exon | e1/e4 | down (e1) | 1,68 | ENSG00000111371 |
| --- | Complex | e1,e2 | up | 1,68 | --- |
| ATXN2 | Complex | e1-2 | up | 1,68 | ENSG00000204842 |
| IRF9 // RNF31 | ter. Terminal Exo | e28/e29-31 | up (e29-31) | 1,68 | ENSG00000092098 // ENSG00000213928 |
| ATXN3 | Complex | 2-3,e5-10/e1 | down (e2-3,e5-10) | 1,68 | ENSG00000066427 |
| SGSM2 | Exon Cassette | e12 | up | 1,68 | ENSG00000141258 |
| SPAG9 | Exon Cassette | e7 | up | 1,68 | ENSG00000008294 |
| FSD1 | Complex | e2 | down | 1,68 | ENSG00000105255 |
| R3HDM1 | Exon Cassette | e16 | down | 1,68 | ENSG00000048991 |
| CCDC150 | Alter. First Exon | 8-12,e14-17 | e1-2,e5,e8-12,e14-1 | 1,68 | ENSG00000144395 |
| ATG4B | Exon Cassette | e6 | up | 1,68 | ENSG00000168397 |
| ABCA12 | Exon Cassette | e24 | up | 1,68 | ENSG00000144452 |
| FARSB | Exon Cassette | e2-3 | up | 1,68 | ENSG00000116120 |
| RABL2B | Intron Retention | e1,e2 | up | 1,68 | ENSG00000079974 |
| DAG1 | Alter. First Exon | e1/e7,e8-11 | down (e1) | 1,68 | ENSG00000173402 |
| RBM5 // RBM6 | Complex | e3-6,e9-17 | down | 1,68 | ENSG00000003756 // ENSG00000004534 |
| CAMKV | Complex | e6-8 | up | 1,68 | ENSG00000164076 |

| | | | | | |
|---|---|---|---|---|---|
| HAND2-AS1 | lter. Acceptor Sit | e3 | down | 1,68 | ENSG00000237125 |
| CDKL3 // PPP2CA | ter. Terminal Exo | e11,e12-13 | up (e11,e12-13,e17 | 1,68 | ENSG00000006837 // ENSG00000113575 |
| VEGFA | Alter. First Exon | e1-3/e4 | down (e1-3) | 1,68 | ENSG00000112715 |
| FAM135A | Alter. First Exon | e4,e6-15/e16 | up (e4,e6-15) | 1,68 | ENSG00000082269 |
| FAM135A | Alter. First Exon | e4-15/e16 | up (e4-15) | 1,68 | ENSG00000082269 |
| MCM9 | ter. Terminal Exo | e7/e9,e10-14 | up (e7) | 1,68 | ENSG00000111877 |
| HDDC2 | Exon Cassette | e3 | up | 1,68 | ENSG00000111906 |
| MAP7 | Alter. First Exon | e2/e4 | down (e2) | 1,68 | ENSG00000135525 |
| EPM2A | Alter. First Exon | e2/e3 | up (e2) | 1,68 | ENSG00000112425 |
| AUTS2 | Exon Cassette | e11 | up | 1,68 | ENSG00000158321 |
| STEAP2 | Alter. First Exon | e2-3/e5 | down (e2-3) | 1,68 | ENSG00000157214 |
| HUS1 | Intron Retention | e8 | down | 1,68 | ENSG00000136273 |
| CSPP1 | Exon Cassette | e6,e7 | up | 1,68 | ENSG00000104218 |
| RPS20 | Alter. First Exon | e1-4/e5 | up (e1-4) | 1,68 | ENSG00000008988 |
| CYHR1 | Alter. First Exon | e1/e2 | down (e2) | 1,68 | ENSG00000187954 |
| IAA1984 // RABL | ter. Terminal Exo | 24/e26-27,e3 | down (e26-27,e33) | 1,68 | ENSG00000196642 // ENSG00000213213 |
| FNBP1 | Alter. First Exon | ,e14-18,e20 | (e1-10,e12,e14-18 | 1,68 | ENSG00000187239 |
| RMCX5-GPRASP | lter. Acceptor Sit | e11 | up | 1,68 | ENSG00000125962 // ENSG00000158301 |
| AGO3 | ter. Terminal Exo | 7/e10,e11-2 | up (e7) | 1,67 | ENSG00000126070 |
| PRKACB | Alter. First Exon | e1,e7/e8 | up (e1,e7) | 1,67 | ENSG00000142875 |
| PRPF38B | ter. Terminal Exo | e5-6/e8,e9 | up (e5-6) | 1,67 | ENSG00000134186 |
| PIAS3 | Intron Retention | e7,e8 | up | 1,67 | ENSG00000131788 |
| LEPRE1 | Intron Retention | e8 | down | 1,67 | ENSG00000117385 |

| | | | | | |
|---|---|---|---|---|---|
| MKNK1 // MOB3C | ter. Terminal Exo | 5/e7,e9,e11 | down (e3-5) | 1,67 | ENSG00000 079277 // ENSG00000 142961 |
| APH1A | Intron Retention | e2 | up | 1,67 | ENSG00000 117362 |
| ARNT | Exon Cassette | e2 | up | 1,67 | ENSG00000 143437 |
| JTB | Intron Retention | e2 | up | 1,67 | ENSG00000 143543 |
| ARHGEF11 | Intron Retention | e38 | up | 1,67 | ENSG00000 132694 |
| SEC24C | Exon Cassette | e8 | down | 1,67 | ENSG00000 176986 |
| WDR11 | Intron Retention | e28 | up | 1,67 | ENSG00000 120008 |
| CAMK2G | Exon Cassette | e16 | down | 1,67 | ENSG00000 148660 |
| PTDSS2 | Alter. First Exon | e1/e3 | down (e1) | 1,67 | ENSG00000 174915 |
| GAS6 | Alter. First Exon | e1-2,e4/e5 | up (e1-2,e4) | 1,67 | ENSG00000 183087 |
| ATXN3 | Exon Cassette | e6,e8-10 | down | 1,67 | ENSG00000 066427 |
| 2AP // IGHD // IG | Alter. First Exon | 8,e40,e103/ | wn (e35-38,e40,e10 | 1,67 | ENSG00000 211896 // ENSG00000 211898 // ENSG00000 213140 |
| PKM | Complex | e2,e5 | up | 1,67 | ENSG00000 067225 |
| TTC23 | Exon Cassette | e4,e5 | down | 1,67 | ENSG00000 103852 |
| TAOK2 | ter. Terminal Exo | 16-17/e18-2 | up (e16-17) | 1,67 | ENSG00000 149930 |
| DPH1 // OVCA2 | Intron Retention | e13 | up | 1,67 | ENSG00000 108963 // ENSG00000 262664 |
| MED24 | Complex | e1,e2 | up | 1,67 | ENSG00000 008838 |
| NEDD4L | Exon Cassette | e21 | up | 1,67 | ENSG00000 049759 |
| PIN1 | Exon Cassette | e3 | up | 1,67 | ENSG00000 127445 |
| BRSK1 | Alter. First Exon | e1-11/e12 | down (e1-11) | 1,67 | ENSG00000 160469 |
| TMEM205 | Complex | e2,e3 | down | 1,67 | ENSG00000 105518 |
| MERTK | ter. Terminal Exo | e19-20/e21 | down (e19-20) | 1,67 | ENSG00000 153208 |

| | | | | | |
|---|---|---|---|---|---|
| // LY75 // LY75- | ter. Terminal Exo | /e14-33,e37 | own (e14-33,e37-41 | 1,67 | ENSG00000054219 // ENSG00000241399 // ENSG00000248672 |
| PATZ1 | ter. Terminal Exo | e4/e5,e7 | up (e4) | 1,67 | ENSG00000100105 |
| NKTR | Exon Cassette | e8 | up | 1,67 | ENSG00000114857 |
| PTPRG | Alter. First Exon | e1/e2 | down (e1) | 1,67 | ENSG00000144724 |
| RNF7 | Complex | e1,e2 | up | 1,67 | ENSG00000114125 |
| RNF7 | Exon Cassette | e2 | up | 1,67 | ENSG00000114125 |
| EIF2B5 | Intron Retention | e3 | up | 1,67 | ENSG00000145191 |
| SEPSECS | Exon Cassette | e3 | down | 1,67 | ENSG00000109618 |
| SREK1 | Intron Retention | e10 | up | 1,67 | ENSG00000153914 |
| OCLN | Complex | e4/e5-6,e8-9 | up (e4) | 1,67 | ENSG00000197822 |
| DDX46 | Intron Retention | e20 | up | 1,67 | ENSG00000145833 |
| CEP120 | Complex | e15-17 | up | 1,67 | ENSG00000168944 |
| ARAP3 | Exon Cassette | e32 | down | 1,67 | ENSG00000120318 |
| FAXDC2 | Exon Cassette | e2 | up | 1,67 | ENSG00000170271 |
| ZNF76 | ter. Terminal Exo | e10/e11,e15 | up (e10) | 1,67 | ENSG00000065029 |
| FAM135A | Alter. First Exon | e3-4,e6-15/e | up (e1,e3-4,e6-15) | 1,67 | ENSG00000082269 |
| KCNQ5 | Exon Cassette | e10,e11-12 | up | 1,67 | ENSG00000185760 |
| 5-SAPCD1 // SAF | Exon Cassette | e9 | down | 1,67 | ENSG00000228727 // ENSG00000255152 |
| RBAKDN // RBAK | ter. Terminal Exo | e15/e17,e18 | down (e15) | 1,67 | ENSG00000146587 // ENSG00000196204 |
| FAM221A | Exon Cassette | e7 | up | 1,67 | ENSG00000188732 |
| ZMIZ2 | Complex | e3-12,e14-19 | down | 1,67 | ENSG00000122515 |

| | | | | | |
|---|---|---|---|---|---|
| IAA1984 // RABL | ter. Terminal Exo | e24/e26-33 | down (e26-33) | 1,67 | ENSG00000 196642 // ENSG00000 213213 |
| IAA1984 // RABL | ter. Terminal Exo | 24/e26,e27-3 | down (e26,e27-33) | 1,67 | ENSG00000 196642 // ENSG00000 213213 |
| POLE3 | Intron Retention | e3 | up | 1,67 | ENSG00000 148229 |
| SCML1 | Alter. First Exon | e1/e2,e3-4 | down (e1) | 1,67 | ENSG00000 047634 |
| WDTC1 | Intron Retention | e14 | up | 1,66 | ENSG00000 142784 |
| ZNF691 | Intron Retention | e2-3 | up | 1,66 | ENSG00000 164011 |
| AKR1A1 | Intron Retention | e5 | up | 1,66 | ENSG00000 117448 |
| SETDB1 | Intron Retention | e8 | up | 1,66 | ENSG00000 143379 |
| EPS15 | ter. Terminal Exo | e10-13,e15- | up (e9) | 1,66 | ENSG00000 085832 |
| TRIM33 | ter. Terminal Exo | e17,e18-19 | up (e16) | 1,66 | ENSG00000 197323 |
| MRPL55 | Exon Cassette | e3 | up | 1,66 | ENSG00000 162910 |
| RGS7 | Complex | e6-9/e7-8 | up (e7-8) | 1,66 | ENSG00000 182901 |
| POLR1D | Complex | e4,e5 | up | 1,66 | ENSG00000 186184 |
| COL4A1 | Complex | e24,e26-33 | down | 1,66 | ENSG00000 187498 |
| FUT8 | Complex | e1,e5/e3 | up (e3) | 1,66 | ENSG00000 033170 |
| ATXN3 | Complex | 3,e6,e8-10/ | down (e2-3,e6,e8-10 | 1,66 | ENSG00000 066427 |
| 2AP // IGHD // IG | Alter. First Exon | 6-27,e101/e | up (e26-27,e101) | 1,66 | ENSG00000 211896 // ENSG00000 211898 // ENSG00000 213140 |
| 2AP // IGHD // IG | Alter. First Exon | 6-27/e71,e1 | up (e26-27) | 1,66 | ENSG00000 211896 // ENSG00000 211898 // ENSG00000 213140 |
| WDR61 | Intron Retention | e2 | up | 1,66 | ENSG00000 140395 |
| C17orf76-AS1 | Complex | e2,e3 | up | 1,66 | ENSG00000 175061 |

| | | | | | |
|---|---|---|---|---|---|
| C17orf76-AS1 | Complex | e1/e2,e3 | up (e2,e3) | 1,66 | ENSG00000 175061 |
| CDK5RAP3 | Alter. First Exon | e2-6/e7 | down (e2-6) | 1,66 | ENSG00000 108465 |
| PRKAR1A | Alter. First Exon | e1/e3 | up (e3) | 1,66 | ENSG00000 108946 |
| SLC25A19 | Alter. First Exon | e1-3/e4 | down (e1-3) | 1,66 | ENSG00000 125454 |
| GPN1 // ZNF512 | Alter. First Exon | ,e8-15,e19/ | own (e1-6,e8-15,e1 | 1,66 | ENSG00000 198522 // ENSG00000 243943 |
| PP1CB // SPDY | Complex | e2 | up | 1,66 | ENSG00000 163806 // ENSG00000 213639 |
| PAPOLG | Intron Retention | e20 | up | 1,66 | ENSG00000 115421 |
| NAGK | Intron Retention | e5 | up | 1,66 | ENSG00000 124357 |
| FMNL2 | Exon Cassette | e27 | up | 1,66 | ENSG00000 157827 |
| FASTKD2 | Intron Retention | e11 | up | 1,66 | ENSG00000 118246 |
| SMC6 | Exon Cassette | e6 | up | 1,66 | ENSG00000 163029 |
| IDH3B | Intron Retention | e7 | up | 1,66 | ENSG00000 101365 |
| STAU1 | Exon Cassette | e4-5 | down | 1,66 | ENSG00000 124214 |
| TMEM189-UBE2 | Exon Cassette | e11,e15 | up | 1,66 | ENSG00000 124208 // ENSG00000 240849 // ENSG00000 244687 |
| PSMG1 | Exon Cassette | e2 | down | 1,66 | ENSG00000 183527 |
| BID | Complex | e2,e7/e4-6 | up (e4-6) | 1,66 | ENSG00000 015475 |
| RABL2B | Alter. Donor Site | e2 | up | 1,66 | ENSG00000 079974 |
| PXK | Exon Cassette | e2-3,e5 | up | 1,66 | ENSG00000 168297 |
| PXK | Exon Cassette | e3,e5 | up | 1,66 | ENSG00000 168297 |
| POLR1C | Intron Retention | e4 | up | 1,66 | ENSG00000 171453 |
| SOBP | ter. Terminal Ex | e6/e7-10 | down (e7-10) | 1,66 | ENSG00000 112320 |
| IQCE | Exon Cassette | e2-3 | down | 1,66 | ENSG00000 106012 |

| | | | | | |
|---|---|---|---|---|---|
| UPP1 | Exon Cassette | e5-7 | up | 1,66 | ENSG00000183696 |
| STEAP2 | Alter. First Exon | e2/e5 | down (e2) | 1,66 | ENSG00000157214 |
| ZNHIT1 | Exon Cassette | e2 | up | 1,66 | ENSG00000106400 |
| ARF5 | Intron Retention | e3 | up | 1,66 | ENSG00000004059 |
| TBL2 | lter. Acceptor Sit | e3 | down | 1,66 | ENSG00000106638 |
| OTUD6B | Exon Cassette | e4 | up | 1,66 | ENSG00000155100 |
| PTPDC1 | ter. Terminal Exo | e8/e9-11 | up (e8) | 1,66 | ENSG00000158079 |
| STAG2 | Complex | e1/e4-15 | up (e4-15) | 1,66 | ENSG00000101972 |
| MAP7D2 | Complex | e7,e8-9 | up | 1,66 | ENSG00000184368 |
| LCN6 // NPPA-AS | ter. Terminal Exo | e4/e5-23 | up (e4) | 1,65 | ENSG00000011021 // ENSG00000242349 |
| LCN6 // NPPA-AS | ter. Terminal Exo | e4/e5,e6-25 | up (e4) | 1,65 | ENSG00000011021 // ENSG00000242349 |
| GNPAT | Exon Cassette | e3 | up | 1,65 | ENSG00000116906 |
| TTLL7 | Exon Cassette | e25 | up | 1,65 | ENSG00000137941 |
| SUFU | ter. Terminal Exo | e12/e13,e14 | down (e13,e14) | 1,65 | ENSG00000107882 |
| ARFIP2 | ter. Terminal Exo | e3/e4-8 | up (e3) | 1,65 | ENSG00000132254 |
| ARFIP2 | ter. Terminal Exo | e3/e5,e6-8 | up (e3) | 1,65 | ENSG00000132254 |
| MYL6 | Complex | e8/e9 | up (e8) | 1,65 | ENSG00000092841 |
| ZNF10 // ZNF268 | Complex | e1,e2 | up | 1,65 | ENSG00000090612 // ENSG00000256223 |
| RPLP0 | Intron Retention | e2,e3 | up | 1,65 | ENSG00000089157 |
| SCFD1 | Exon Cassette | e8 | up | 1,65 | ENSG00000092108 |
| FUT8 | Complex | e1-5/e3-4 | up (e3-4) | 1,65 | ENSG00000033170 |
| ATXN3 | Complex | 3,e6,e8-11/ | down (e2-3,e6,e8-11 | 1,65 | ENSG00000066427 |
| FANCI | Complex | e1,e2-3 | up | 1,65 | ENSG00000140525 |

| | | | | | |
|---|---|---|---|---|---|
| FANCI | Intron Retention | e18 | up | 1,65 | ENSG00000140525 |
| PPIP5K1 | Complex | e24-26 | up | 1,65 | ENSG00000168781 |
| HAGH | Exon Cassette | e2 | down | 1,65 | ENSG00000063854 |
| CNTROB | Intron Retention | e17 | up | 1,65 | ENSG00000170037 |
| ABHD1 | Exon Cassette | e4 | down | 1,65 | ENSG00000143994 |
| PREB | Intron Retention | e2 | up | 1,65 | ENSG00000138073 |
| MYNN | Exon Cassette | e2 | up | 1,65 | ENSG00000085274 |
| FGD5-AS1 | ter. Terminal Exo | e4/e5,e6-9 | down (e4) | 1,65 | ENSG00000225733 |
| C31A // THAP9-A | Exon Cassette | e30 | up | 1,65 | ENSG00000138674 // ENSG00000251022 |
| SREK1 | Intron Retention | e9,e10 | up | 1,65 | ENSG00000153914 |
| OCLN | Exon Cassette | e4 | up | 1,65 | ENSG00000197822 |
| RASA1 | Complex | e1,e3 | up | 1,65 | ENSG00000145715 |
| SOBP | ter. Terminal Exo | e6/e7,e8 | down (e7,e8) | 1,65 | ENSG00000112320 |
| DDR1 | lter. Acceptor Sit | e16 | up | 1,65 | ENSG00000204580 |
| SNK2B // LY6G5 | Exon Cassette | e6 | down | 1,65 | ENSG00000204435 // ENSG00000240053 |
| DLD | lter. Acceptor Sit | e6 | up | 1,65 | ENSG00000091140 |
| POLM | Intron Retention | e8-9 | up | 1,65 | ENSG00000122678 |
| PON2 | Complex | e2/e3 | up (e2) | 1,65 | ENSG00000105854 |
| COPS5 | Intron Retention | e1,e2 | up | 1,65 | ENSG00000121022 |
| --- | Exon Cassette | e2 | up | 1,65 | --- |
| NTRK2 | ter. Terminal Exo | e19/e20-22 | up (e20-22) | 1,65 | ENSG00000148053 |
| C9orf89 | Alter. First Exon | e1-2/e3 | down (e1-2) | 1,65 | ENSG00000165233 |
| --- | Exon Cassette | e7-8 | up | 1,65 | --- |
| NAA10 | Intron Retention | e5 | up | 1,65 | ENSG00000102030 |

| | | | | | |
|---|---|---|---|---|---|
| LCN6 // NPPA-AS | ter. Terminal Exo | e4/e5,e6-22 | up (e4) | 1,64 | ENSG00000 011021 // ENSG00000 242349 |
| RERE | Exon Cassette | e2 | up | 1,64 | ENSG00000 142599 |
| EPS15 | ter. Terminal Exo | 11-13,e15-1 | up (e9) | 1,64 | ENSG00000 085832 |
| CDCP2 // CYB5R | Complex | e4,e5 | down | 1,64 | ENSG00000 157211 // ENSG00000 215883 |
| DBT | lter. Acceptor Sit | e9 | up | 1,64 | ENSG00000 137992 |
| MTMR11 | Exon Cassette | e8-13 | up | 1,64 | ENSG00000 014914 |
| MRPL55 | Complex | e1-2 | up | 1,64 | ENSG00000 162910 |
| DDB2 | Complex | e6 | down | 1,64 | ENSG00000 134574 |
| C11orf80 // RCE1 | Alter. First Exon | e1,e4-15/e16 | down (e1,e4-15) | 1,64 | ENSG00000 173653 // ENSG00000 173715 |
| ANKRD13D | Intron Retention | e8 | up | 1,64 | ENSG00000 172932 |
| APBB1 | Intron Retention | e16 | up | 1,64 | ENSG00000 166313 |
| NUMA1 | Complex | 25-26,e32-3 | up | 1,64 | ENSG00000 137497 |
| WASF3 | ualy Exclusive Ex | e7/e8 | down (e7) | 1,64 | ENSG00000 132970 |
| FLT1 | Alter. First Exon | e24/e25 | down (e24) | 1,64 | ENSG00000 102755 |
| BRF1 | Complex | e11,e12-13 | down | 1,64 | ENSG00000 185024 |
| FBF1 // MRPL38 | Complex | e31,e32 | up | 1,64 | ENSG00000 188878 // ENSG00000 204316 |
| ACOX1 | Complex | e2-3,e5 | up | 1,64 | ENSG00000 161533 |
| BSG | Alter. First Exon | e2,e4/e3-5 | down (e2,e4) | 1,64 | ENSG00000 172270 |
| SNAPC2 | Alter. First Exon | e1/e2 | up (e2) | 1,64 | ENSG00000 104976 |
| ZNF525 // ZNF765 | ualy Exclusive Ex | e2/e3 | down (e2) | 1,64 | ENSG00000 196417 // ENSG00000 203326 |

| | | | | | |
|---|---|---|---|---|---|
| M228A // FAM22 | Exon Cassette | e13 | up | 1,64 | ENSG00000186453 // ENSG00000219626 |
| SLC30A6 | Exon Cassette | e10 | down | 1,64 | ENSG00000152683 |
| AAK1 | ter. Terminal Ex | e14-18/e22 | down (e14-18) | 1,64 | ENSG00000115977 |
| CSNK2A1 | Exon Cassette | e2,e3 | down | 1,64 | ENSG00000101266 |
| NAPB | Alter. First Exon | e1-2/e4 | up (e1-2) | 1,64 | ENSG00000125814 |
| DSN1 | Intron Retention | e2 | up | 1,64 | ENSG00000149636 |
| ) // CRYZL1 // D | lter. Acceptor Sit | e18 | up | 1,64 | ENSG00000159147 // ENSG00000205758 // ENSG00000241837 |
| CRELD2 | Intron Retention | e9 | down | 1,64 | ENSG00000184164 |
| PRAME | Complex | e2-3,e6 | up | 1,64 | ENSG00000185686 |
| ASTE1 | Exon Cassette | e5 | up | 1,64 | ENSG00000034533 |
| ANK2 | Complex | e33-42 | up | 1,64 | ENSG00000145362 |
| RXRB | ter. Terminal Ex | e7/e8-11 | up (e7) | 1,64 | ENSG00000204231 |
| IQCE | Exon Cassette | e3 | down | 1,64 | ENSG00000106012 |
| IRPS24 // URGC | Alter. First Exon | e1/e3 | up (e1) | 1,64 | ENSG00000062582 // ENSG00000106608 |
| SEMA3C | Alter. First Exon | e1/e2 | down (e2) | 1,64 | ENSG00000075223 |
| EPHB4 | Alter. Donor Site | e8 | up | 1,64 | ENSG00000196411 |
| C7orf49 | Alter. First Exon | e1,e4/e5 | down (e1,e4) | 1,64 | ENSG00000122783 |
| CERCAM | Complex | e3-5 | up | 1,64 | ENSG00000167123 |
| SCML1 | Alter. First Exon | e1/e2-3 | down (e1) | 1,64 | ENSG00000047634 |
| MID1 | Exon Cassette | e15 | down | 1,64 | ENSG00000101871 |
| LCN6 // NPPA-A | ter. Terminal Ex | e4/e5,e6-23 | up (e4) | 1,63 | ENSG00000011021 // ENSG00000242349 |

| | | | | | |
|---|---|---|---|---|---|
| PRPF38B | ter. Terminal Exo | e6/e8-9 | up (e6) | 1,63 | ENSG00000134186 |
| PRPF38B | ter. Terminal Exo | e6/e8,e9 | up (e6) | 1,63 | ENSG00000134186 |
| SLC35E2B | Intron Retention | e6 | up | 1,63 | ENSG00000189339 |
| TPM3 | Alter. First Exon | e5,e7-9/e10 | down (e5,e7-9) | 1,63 | ENSG00000143549 |
| GPR161 | Alter. First Exon | e1-2,e6/e7 | up (e1-2,e6) | 1,63 | ENSG00000143147 |
| FGFR2 | Exon Cassette | e16,e17 | up | 1,63 | ENSG00000066468 |
| CADM1 | Exon Cassette | e9 | up | 1,63 | ENSG00000182985 |
| SPATS2 | Exon Cassette | e2 | down | 1,63 | ENSG00000123352 |
| NABP2 | Alter. First Exon | e1/e2 | up (e1) | 1,63 | ENSG00000139579 |
| PARPBP | Exon Cassette | e5,e7,e9-14 | up | 1,63 | ENSG00000185480 |
| RBM25 | Intron Retention | e17 | up | 1,63 | ENSG00000119707 |
| PAPOLA | Intron Retention | e8 | up | 1,63 | ENSG00000090060 |
| ATXN3 | Exon Cassette | e6-10 | down | 1,63 | ENSG00000066427 |
| 2AP // IGHD // IG | Alter. First Exon | 1/e72-73,e1 | down (e71) | 1,63 | ENSG00000211896 // ENSG00000211898 // ENSG00000213140 |
| 2AP // IGHD // IG | Alter. First Exon | e71/e72-73 | down (e71) | 1,63 | ENSG00000211896 // ENSG00000211898 // ENSG00000213140 |
| 2AP // IGHD // IG | Alter. First Exon | 1,e101/e72- | down (e71,e101) | 1,63 | ENSG00000211896 // ENSG00000211898 // ENSG00000213140 |
| 2AP // IGHD // IG | Alter. First Exon | /e72,e73,e1 | down (e71) | 1,63 | ENSG00000211896 // ENSG00000211898 // ENSG00000213140 |

| | | | | | |
|---|---|---|---|---|---|
| 2AP // IGHD // IG | Alter. First Exon | 1/e72-73,e1 | down (e71) | 1,63 | ENSG00000 211896 // ENSG00000 211898 // ENSG00000 213140 |
| 2AP // IGHD // IG | Alter. First Exon | 1,e100/e72- | down (e71,e100) | 1,63 | ENSG00000 211896 // ENSG00000 211898 // ENSG00000 213140 |
| TTC23 | Exon Cassette | e4 | down | 1,63 | ENSG00000 103852 |
| ACD | Intron Retention | e6 | up | 1,63 | ENSG00000 102977 |
| KSR1 | Exon Cassette | e7 | up | 1,63 | ENSG00000 141068 |
| NKLE1 // BABAM | Intron Retention | e7 | up | 1,63 | ENSG00000 105393 // ENSG00000 160117 |
| ZNF480 | Exon Cassette | e4 | up | 1,63 | ENSG00000 198464 |
| MAP4K3 | Exon Cassette | e15,e16 | up | 1,63 | ENSG00000 011566 |
| CCT4 | Complex | e1,e2 | up | 1,63 | ENSG00000 115484 |
| FN1 | Exon Cassette | e40-42 | down | 1,63 | ENSG00000 115414 |
| UQCC | ter. Terminal Exo | e8/e9-10 | up (e8) | 1,63 | ENSG00000 101019 |
| TIMP3 | Complex | e1,e2-5 | up | 1,63 | ENSG00000 100234 |
| TEX264 | Exon Cassette | e3,e5 | up | 1,63 | ENSG00000 164081 |
| THAP9 | Exon Cassette | e4 | up | 1,63 | ENSG00000 168152 |
| USP53 | Exon Cassette | e4 | down | 1,63 | ENSG00000 145390 |
| APBB2 | Alter. First Exon | -8,e10-13/e | up (e2-8,e10-13) | 1,63 | ENSG00000 163697 |
| NSD1 | Intron Retention | e4 | down | 1,63 | ENSG00000 165671 |
| BBS9 | Exon Cassette | e17-18 | down | 1,63 | ENSG00000 122507 |
| SLC35D2 | Exon Cassette | e9-11 | up | 1,63 | ENSG00000 130958 |
| AKNA | Alter. First Exon | e1,e5/e3-4 | up (e1,e5) | 1,63 | ENSG00000 106948 |
| NSMF | Intron Retention | e8 | up | 1,63 | ENSG00000 165802 |

| | | | | | |
|---|---|---|---|---|---|
| OGT | lter. Acceptor Sit | e2 | up | 1,63 | ENSG00000147162 |
| DFFB | Complex | e2/e3 | up (e3) | 1,62 | ENSG00000169598 |
| DFFB | Complex | e3/e4 | up (e3) | 1,62 | ENSG00000169598 |
| DFFB | Exon Cassette | e3 | up | 1,62 | ENSG00000169598 |
| AGO3 | ter. Terminal Exc | e7-8/e10-23 | up (e7-8) | 1,62 | ENSG00000126070 |
| CASP9 | lter. Acceptor Sit | e5 | up | 1,62 | ENSG00000132906 |
| CDC42SE1 | Intron Retention | e3 | up | 1,62 | ENSG00000197622 |
| TAF5L | ter. Terminal Exc | e4/e5 | up (e4) | 1,62 | ENSG00000135801 |
| SHOC2 | Exon Cassette | e3 | down | 1,62 | ENSG00000108061 |
| VTI1A | Exon Cassette | e2 | up | 1,62 | ENSG00000151532 |
| SORCS1 | Complex | e2/e3 | down (e2) | 1,62 | ENSG00000108018 |
| TBRG1 | Exon Cassette | e5 | up | 1,62 | ENSG00000154144 |
| SPATS2 | Exon Cassette | e3 | up | 1,62 | ENSG00000123352 |
| LETMD1 | Exon Cassette | e4 | up | 1,62 | ENSG00000050426 |
| CPNE8 | Alter. First Exon | e1/e2 | down (e2) | 1,62 | ENSG00000139117 |
| COMMD6 | Exon Cassette | e3 | down | 1,62 | ENSG00000188243 |
| PRC1 | Complex | e13-14 | up | 1,62 | ENSG00000198901 |
| FAM65A | Complex | e15,e16-24 | up | 1,62 | ENSG00000039523 |
| C17orf76-AS1 | Complex | e1,e2-3 | up | 1,62 | ENSG00000175061 |
| MBTD1 | Intron Retention | e17 | up | 1,62 | ENSG00000011258 |
| EXOC7 | ter. Terminal Exc | /e5-13,e15- | up (e4) | 1,62 | ENSG00000182473 |
| -RAB4B // RAB4 | ter. Terminal Exc | e6/e8-14 | down (e8-14) | 1,62 | ENSG00000167578 // ENSG00000171570 // ENSG00000261857 // ENSG00000268975 // ENSG00000269858 |

| | | | | | |
|---|---|---|---|---|---|
| SCLY // UBE2F | Exon Cassette | e19 | up | 1,62 | ENSG00000 132330 // ENSG00000 184182 |
| Sep-02 | Exon Cassette | e5 | up | 1,62 | ENSG00000 168385 |
| C2orf43 | Exon Cassette | e2 | down | 1,62 | ENSG00000 118961 |
| CACNB4 | ter. Terminal Exo | 2/e13-15,e1 | up (e13-15,e17) | 1,62 | ENSG00000 182389 |
| ARFGAP1 | Intron Retention | e13,e14-15 | up | 1,62 | ENSG00000 101199 |
| SNX5 | Alter. First Exon | e1-2/e3 | down (e1-2) | 1,62 | ENSG00000 089006 |
| NAPB | Alter. First Exon | e1-3/e4 | up (e1-3) | 1,62 | ENSG00000 125814 |
| UQCC | Complex | e4,e5,e8-9 | down | 1,62 | ENSG00000 101019 |
| E1 // NFS1 // RB | Complex | e21,e22-23 | up | 1,62 | ENSG00000 214078 // ENSG00000 244005 // ENSG00000 244462 |
| SNHG17 | ter. Terminal Exo | 2/e3,e5,e8-9 | up (e2) | 1,62 | ENSG00000 196756 |
| THOC5 | Exon Cassette | e14 | up | 1,62 | ENSG00000 100296 |
| IL17RC | Complex | e18 | down | 1,62 | ENSG00000 163702 |
| NR2C2 | Exon Cassette | e4 | up | 1,62 | ENSG00000 177463 |
| PXK | Exon Cassette | e5 | up | 1,62 | ENSG00000 168297 |
| NIT2 | Complex | e5 | up | 1,62 | ENSG00000 114021 |
| QRICH1 | Exon Cassette | e3 | up | 1,62 | ENSG00000 198218 |
| FOXP1 | Intron Retention | e27,e28 | up | 1,62 | ENSG00000 114861 |
| ABI3BP | ter. Terminal Exo | 14,e15-31,e | vn (e14,e15-31,e36- | 1,62 | ENSG00000 154175 |
| FRYL | ter. Terminal Exo | e6,e7-57,e59 | up (e5) | 1,62 | ENSG00000 075539 |
| OCLN | Exon Cassette | e4-5 | up | 1,62 | ENSG00000 197822 |
| TMEM161B | Exon Cassette | e2,e3 | up | 1,62 | ENSG00000 164180 |
| NUDT12 | Intron Retention | e2 | up | 1,62 | ENSG00000 112874 |
| YIPF3 | Intron Retention | e6 | up | 1,62 | ENSG00000 137207 |

| | | | | | |
|---|---|---|---|---|---|
| SFT2D1 | Intron Retention | e7 | down | 1,62 | ENSG00000198818 |
| WIPI2 | Exon Cassette | e2 | up | 1,62 | ENSG00000157954 |
| WBSCR22 | Intron Retention | e7 | up | 1,62 | ENSG00000071462 |
| SH3GLB2 | Alter. First Exon | e1-5,e7/e8 | down (e1-5,e7) | 1,62 | ENSG00000148341 |
| MAP7D2 | Complex | e9,e10-13 | down | 1,62 | ENSG00000184368 |
| AIFM1 | Exon Cassette | e2,e4-10 | up | 1,62 | ENSG00000156709 |
| MECP2 | Complex | e1,e3/e4 | up (e1,e3) | 1,62 | ENSG00000169057 |
| HDAC1 | Exon Cassette | e2 | up | 1,61 | ENSG00000116478 |
| RAP1A | Exon Cassette | e4 | up | 1,61 | ENSG00000116473 |
| GUK1 | Exon Cassette | e3 | down | 1,61 | ENSG00000143774 |
| C1orf63 | lter. Acceptor Sit | e5 | up | 1,61 | ENSG00000117616 |
| BAI2 | Alter. First Exon | e1/e2-4 | up (e2-4) | 1,61 | ENSG00000121753 |
| OMMD3 // COMM | Intron Retention | e4 | up | 1,61 | ENSG00000148444 // ENSG00000168283 // ENSG00000269897 |
| P4HA1 | Exon Cassette | e2 | down | 1,61 | ENSG00000122884 |
| TSPAN4 | Exon Cassette | e4 | up | 1,61 | ENSG00000214063 |
| ILK | Intron Retention | e6 | up | 1,61 | ENSG00000166333 |
| PLEKHA5 | Alter. First Exon | e1/e6,e7 | down (e1) | 1,61 | ENSG00000052126 |
| MDM2 | Complex | 8,e9,e11,e1 | down | 1,61 | ENSG00000135679 |
| MDM2 | Complex | e5,e7,e14 | down | 1,61 | ENSG00000135679 |
| CDK17 | Intron Retention | e16 | down | 1,61 | ENSG00000059758 |
| CAMKK2 | Exon Cassette | e15 | down | 1,61 | ENSG00000110931 |
| CDC16 | Intron Retention | e6 | up | 1,61 | ENSG00000130177 |
| FUT8 | Complex | e1,e4/e3 | up (e3) | 1,61 | ENSG00000033170 |
| FUT8 | Exon Cassette | e3 | up | 1,61 | ENSG00000033170 |

| | | | | | |
|---|---|---|---|---|---|
| ATXN3 | Exon Cassette | e3-5 | down | 1,61 | ENSG00000066427 |
| MOK | Intron Retention | e11-12 | up | 1,61 | ENSG00000080823 |
| C15orf41 | Intron Retention | e14 | down | 1,61 | ENSG00000186073 |
| F6 // HEXA // PA | Exon Cassette | e48 | up | 1,61 | ENSG00000137817 // ENSG00000140488 // ENSG00000213614 |
| TRAPPC2L | Complex | e3/e4,e5 | up (e4,e5) | 1,61 | ENSG00000167515 |
| ZNF397 | ter. Terminal Exc | e3/e5,e6,e8 | down (e3) | 1,61 | ENSG00000186812 |
| ZNF302 | Complex | e6 | up | 1,61 | ENSG00000089335 |
| CACNB4 | ter. Terminal Exc | e12/e13-17 | up (e13-17) | 1,61 | ENSG00000182389 |
| USP25 | Exon Cassette | e19,e20 | up | 1,61 | ENSG00000155313 |
| HSF2BP | Exon Cassette | e3 | up | 1,61 | ENSG00000160207 |
| BID | Alter. First Exon | e1/e2,e4-6 | up (e2,e4-6) | 1,61 | ENSG00000015475 |
| IP6K2 | Intron Retention | e9-10 | up | 1,61 | ENSG00000068745 |
| HEG1 | Exon Cassette | e6 | down | 1,61 | ENSG00000173706 |
| TET2 | Exon Cassette | e5 | up | 1,61 | ENSG00000168769 |
| APBB2 | Alter. First Exon | 8,e10-13/e1 | up (e8,e10-13) | 1,61 | ENSG00000163697 |
| BTF3 | Complex | e1,e2 | down | 1,61 | ENSG00000145741 |
| ARHGEF28 | ter. Terminal Exc | e15/e16-36 | up (e16-36) | 1,61 | ENSG00000214944 |
| GNPDA1 | Intron Retention | e1,e2 | up | 1,61 | ENSG00000113552 |
| ZNF204P | Alter. First Exon | e1-3/e4 | down (e1-3) | 1,61 | ENSG00000204789 |
| IYMAI // PLAGL1 | Complex | e7/e8,e9-10 | up (e8,e9-10) | 1,61 | ENSG00000118495 |
| GNL1 | Complex | e1,e2-8 | down | 1,61 | ENSG00000204590 |
| CUTA | Alter. First Exon | e1/e2 | up (e1) | 1,61 | ENSG00000112514 |
| CCM2 | Exon Cassette | e4 | down | 1,61 | ENSG00000136280 |
| COA1 | Exon Cassette | e3 | up | 1,61 | ENSG00000106603 |

| | | | | | |
|---|---|---|---|---|---|
| ABCB1 | Complex | e2 | down | 1,61 | ENSG00000 085563 |
| PMS2P1 | Exon Cassette | e5-6 | down | 1,61 | ENSG00000 078319 |
| POMT1 | Alter. First Exon | e1,e4/e9 | down (e1,e4) | 1,61 | ENSG00000 130714 |
| C9orf41 | Exon Cassette | e4 | up | 1,61 | ENSG00000 156017 |
| IKBKAP | Alter. First Exon | e1-8/e9 | down (e1-8) | 1,61 | ENSG00000 070061 |
| MAOB | Exon Cassette | e15 | down | 1,61 | ENSG00000 069535 |
| SAG4 // MAGEA2 | Exon Cassette | e2,e3 | up | 1,61 | ENSG00000 183305 // ENSG00000 242599 |
| CMC4 // MTCP1 | Complex | e2,e3-5 | up | 1,61 | ENSG00000 182712 // ENSG00000 214827 |
| UHMK1 | Intron Retention | e3 | up | 1,6 | ENSG00000 152332 |
| SLC35E2B | ter. Terminal Exo | e6/e7,e8-10 | up (e6) | 1,6 | ENSG00000 189339 |
| AKR7A2 | ter. Terminal Exo | e3/e6-7 | up (e3) | 1,6 | ENSG00000 053371 |
| MTMR11 | Exon Cassette | e3 | down | 1,6 | ENSG00000 014914 |
| TPM3 | Alter. First Exon | e5,e8-9/e10 | down (e5,e8-9) | 1,6 | ENSG00000 143549 |
| TPM3 | Alter. First Exon | e5-9/e10 | down (e5-9) | 1,6 | ENSG00000 143549 |
| NVL | Exon Cassette | e6-7 | up | 1,6 | ENSG00000 143748 |
| NVL | Exon Cassette | e6 | up | 1,6 | ENSG00000 143748 |
| TCF7L2 | Exon Cassette | e4,e5 | up | 1,6 | ENSG00000 148737 |
| CAMK2G | Exon Cassette | e20,e22 | up | 1,6 | ENSG00000 148660 |
| POLL | ter. Terminal Exo | e7/e8-10 | up (e7) | 1,6 | ENSG00000 166169 |
| MRPL48 | Exon Cassette | e2-3 | up | 1,6 | ENSG00000 175581 |
| RNH1 | Alter. First Exon | e1/e2-4 | down (e1) | 1,6 | ENSG00000 023191 |
| ZNF195 | Exon Cassette | e6,e7 | up | 1,6 | ENSG00000 005801 |
| DYRK4 | ter. Terminal Exo | e9/e10-16 | up (e10-16) | 1,6 | ENSG00000 010219 |
| MCRS1 | Intron Retention | e3,e4 | up | 1,6 | ENSG00000 187778 |

| | | | | | |
|---|---|---|---|---|---|
| CSAD | Exon Cassette | e13 | up | 1,6 | ENSG00000 139631 |
| CBX5 | Alter. First Exon | e1/e2 | up (e1) | 1,6 | ENSG00000 094916 |
| GALNT16 | Complex | e1-10 | up | 1,6 | ENSG00000 100626 |
| SIVA1 | Intron Retention | e3 | up | 1,6 | ENSG00000 184990 |
| MTA1 | Exon Cassette | e5 | down | 1,6 | ENSG00000 182979 |
| HECTD1 | Alter. First Exon | 1-3,e6-29/e3 | down (e1-3,e6-29) | 1,6 | ENSG00000 092148 |
| HECTD1 | Alter. First Exon | e6-20,e22-2 | wn (e2-3,e6-20,e22- | 1,6 | ENSG00000 092148 |
| ATXN3 | Exon Cassette | e2-3,e8-10 | down | 1,6 | ENSG00000 066427 |
| SETD6 | Intron Retention | e6,e7 | up | 1,6 | ENSG00000 103037 |
| AP1G1 | Exon Cassette | e3-4 | up | 1,6 | ENSG00000 166747 |
| ANKRD11 | Exon Cassette | e6 | down | 1,6 | ENSG00000 167522 |
| MED24 | Alter. First Exon | e1/e2 | down (e1) | 1,6 | ENSG00000 008838 |
| CARD8 | Intron Retention | e6,e7-8 | up | 1,6 | ENSG00000 105483 |
| ZNF615 | Exon Cassette | e6 | up | 1,6 | ENSG00000 197619 |
| ELMOD3 | Intron Retention | e2,e3-4 | up | 1,6 | ENSG00000 115459 |
| OBSL1 | ter. Terminal Exo | e10,e12,e14 | up (e9) | 1,6 | ENSG00000 124006 |
| XYLB | ter. Terminal Exo | e18/e19 | up (e18) | 1,6 | ENSG00000 093217 |
| PHC3 | ter. Terminal Exo | e4/e5 | up (e4) | 1,6 | ENSG00000 173889 |
| REST | ualy Exclusive Ex | e5/e7 | up (e5) | 1,6 | ENSG00000 084093 |
| REST | ualy Exclusive Ex | e5/e6 | up (e5) | 1,6 | ENSG00000 084093 |
| REST | Exon Cassette | e5 | up | 1,6 | ENSG00000 084093 |
| APBB2 | Alter. First Exon | e11-13/e14 | up (e11-13) | 1,6 | ENSG00000 163697 |
| WDFY3 | Exon Cassette | e46 | up | 1,6 | ENSG00000 163625 |
| S5 // EEF1E1 // T | Exon Cassette | e15 | down | 1,6 | ENSG00000 124802 // ENSG00000 188428 // ENSG00000 239264 |

| | | | | | |
|---|---|---|---|---|---|
| IICAL1 // ZBTB2 | Intron Retention | e16 | up | 1,6 | ENSG00000 112365 // ENSG00000 135596 |
| ELN | Complex | 18-21,e24-2 | up | 1,6 | ENSG00000 049540 |
| COA1 | Alter. First Exon | 2,e4-6,e8-10 | up (e1-2,e4-6,e8-10 | 1,6 | ENSG00000 106603 |
| AP3M2 | Exon Cassette | e2 | up | 1,6 | ENSG00000 070718 |
| SH3D21 | Complex | e12,e14 | up | 1,59 | ENSG00000 214193 |
| PRPF38B | Exon Cassette | e7 | up | 1,59 | ENSG00000 134186 |
| MOV10 | Intron Retention | e13 | down | 1,59 | ENSG00000 155363 |
| SLC50A1 | Exon Cassette | e3 | up | 1,59 | ENSG00000 169241 |
| PBX1 | Exon Cassette | e8 | down | 1,59 | ENSG00000 185630 |
| DCAF6 | Exon Cassette | e12,e14 | up | 1,59 | ENSG00000 143164 |
| VASH2 | ter. Terminal Exo | e7-8/e9-11 | up (e9-11) | 1,59 | ENSG00000 143494 |
| VASH2 | Complex | e2,e3 | up | 1,59 | ENSG00000 143494 |
| ACOT7 | Complex | e5-6 | down | 1,59 | ENSG00000 097021 |
| ENAH | Exon Cassette | e12 | down | 1,59 | ENSG00000 154380 |
| MLLT10 | Exon Cassette | e19-20 | up | 1,59 | ENSG00000 078403 |
| ZSWIM8 | ter. Terminal Exo | e1-2/e3-27 | up (e1-2) | 1,59 | ENSG00000 214655 |
| TYSND1 | Alter. First Exon | e2/e3 | down (e2) | 1,59 | ENSG00000 156521 |
| ATAD1 | Exon Cassette | e9 | up | 1,59 | ENSG00000 138138 |
| SORBS1 | Exon Cassette | e36 | up | 1,59 | ENSG00000 095637 |
| SFXN4 | Intron Retention | e1 | down | 1,59 | ENSG00000 183605 |
| TBCEL | Complex | e1,e2-3 | up | 1,59 | ENSG00000 154114 |
| PIDD | Intron Retention | e15 | down | 1,59 | ENSG00000 177595 |
| PICALM | Exon Cassette | e14 | down | 1,59 | ENSG00000 073921 |
| DHRS12 | Exon Cassette | e3 | up | 1,59 | ENSG00000 102796 |
| SRSF5 | Intron Retention | e6 | up | 1,59 | ENSG00000 100650 |

| | | | | | |
|---|---|---|---|---|---|
| ATXN3 | Complex | 2-3,e6-10/e1 | down (e2-3,e6-10) | 1,59 | ENSG00000066427 |
| ATXN3 | Exon Cassette | e3,e5 | down | 1,59 | ENSG00000066427 |
| 2AP // IGHD // IG | Alter. First Exon | 4-65/e71,e1 | up (e64-65) | 1,59 | ENSG00000211896 // ENSG00000211898 // ENSG00000213140 |
| 2AP // IGHD // IG | Alter. First Exon | 4-65,e100/e | up (e64-65,e100) | 1,59 | ENSG00000211896 // ENSG00000211898 // ENSG00000213140 |
| 2AP // IGHD // IG | Alter. First Exon | e64-65/e71 | up (e64-65) | 1,59 | ENSG00000211896 // ENSG00000211898 // ENSG00000213140 |
| SRP14-AS1 | ter. Terminal Exo | e3-4/e5,e6 | up (e3-4) | 1,59 | ENSG00000248508 |
| CA12 | Exon Cassette | e3 | up | 1,59 | ENSG00000074410 |
| :15orf38 // C15or | Exon Cassette | e11 | up | 1,59 | ENSG00000157823 // ENSG00000242498 // ENSG00000250021 |
| RRN3P1 | Alter. First Exon | e3-8/e10 | up (e3-8) | 1,59 | ENSG00000248124 |
| SLC12A4 | Alter. First Exon | e2/e3 | down (e2) | 1,59 | ENSG00000124067 |
| ALDOC | Alter. Donor Site | e2 | up | 1,59 | ENSG00000109107 |
| ZSCAN30 | Alter. Donor Site | e1 | up | 1,59 | ENSG00000186814 |
| ZNF283 | ter. Terminal Exo | e7/e8 | up (e7) | 1,59 | ENSG00000167637 |
| BRSK1 | ter. Terminal Exo | e13/e14-22 | down (e14-22) | 1,59 | ENSG00000160469 |
| MFF | Exon Cassette | e8 | up | 1,59 | ENSG00000168958 |
| Sep-02 | Alter. First Exon | e1-4,e7/e9 | up (e1-4,e7) | 1,59 | ENSG00000168385 |
| U2AF1 | Alter. First Exon | e1/e2 | down (e1) | 1,59 | ENSG00000160201 |
| TPTEP1 | Exon Cassette | e11 | down | 1,59 | ENSG00000100181 |

| | | | | | |
|---|---|---|---|---|---|
| ITPR1 | Exon Cassette | e12 | down | 1,59 | ENSG00000 150995 |
| DAG1 | Alter. First Exon | 1,e6/e7,e8-1 | down (e1,e6) | 1,59 | ENSG00000 173402 |
| DAG1 | Complex | e1,e3 | down | 1,59 | ENSG00000 173402 |
| PBRM1 | Exon Cassette | e29 | up | 1,59 | ENSG00000 163939 |
| APBB2 | Alter. First Exon | e2-13/e14 | up (e2-13) | 1,59 | ENSG00000 163697 |
| ARHGEF28 | ter. Terminal Exo | 5/e16,e17-3 | up (e16,e17-38) | 1,59 | ENSG00000 214944 |
| ARHGAP26 | Complex | e21/e23,e24 | up (e23,e24) | 1,59 | ENSG00000 145819 |
| ARHGAP26 | Exon Cassette | e23 | up | 1,59 | ENSG00000 145819 |
| CDKL3 // PPP2CA | ter. Terminal Exo | e11,e12-14 | up (e11,e12-14,e17 | 1,59 | ENSG00000 006837 // ENSG00000 113575 |
| FAM135A | Alter. First Exon | -10,e12-15/e | up (e1-10,e12-15) | 1,59 | ENSG00000 082269 |
| SLC16A10 | ter. Terminal Exo | e3/e4,e5-6 | down (e3) | 1,59 | ENSG00000 112394 |
| QKI | Intron Retention | e6,e7 | up | 1,59 | ENSG00000 112531 |
| HLA-DOB // TAP2 | ter. Terminal Exo | e17/e18-20 | up (e18-20) | 1,59 | ENSG00000 204267 // ENSG00000 241106 |
| CALU | Exon Cassette | e5 | up | 1,59 | ENSG00000 128595 |
| ATP6V0E2 | Exon Cassette | e3 | up | 1,59 | ENSG00000 171130 |
| FASTK | Alter. First Exon | e1/e2 | down (e1) | 1,59 | ENSG00000 164896 |
| CDC26 // FKBP15 | Alter. First Exon | e6/e20-34 | up (e20-34) | 1,59 | ENSG00000 119321 // ENSG00000 176386 |
| AKNA | Alter. First Exon | e3-4/e5 | down (e3-4) | 1,59 | ENSG00000 106948 |
| RAPGEF1 | Exon Cassette | e14,e15 | up | 1,59 | ENSG00000 107263 |
| DDX3X | ter. Terminal Exo | /e6-12,e14- | up (e4) | 1,59 | ENSG00000 215301 |
| DDX3X | ter. Terminal Exo | e4/e6,e7-13 | up (e4) | 1,59 | ENSG00000 215301 |
| ARHGEF9 | Complex | e2,e4 | down | 1,59 | ENSG00000 131089 |
| BCAP31 | Complex | e2-3 | down | 1,59 | ENSG00000 185825 |

| | | | | | |
|---|---|---|---|---|---|
| LCN6 // NPPA-AS | Exon Cassette | e12 | down | 1,58 | ENSG00000 011021 // ENSG00000 242349 |
| ZCCHC17 | Exon Cassette | e3 | up | 1,58 | ENSG00000 121766 |
| HDAC1 | ter. Terminal Exc | e6/e7-14 | up (e6) | 1,58 | ENSG00000 116478 |
| RAD54L | Complex | e1,e2-4 | up | 1,58 | ENSG00000 085999 |
| TRIM46 | Complex | e3,e7 | down | 1,58 | ENSG00000 163462 |
| ZNF678 | Exon Cassette | e4 | down | 1,58 | ENSG00000 181450 |
| SLC35E2B | ter. Terminal Exc | e6/e7,e8-9 | up (e6) | 1,58 | ENSG00000 189339 |
| FHL3 | Intron Retention | e4 | up | 1,58 | ENSG00000 183386 |
| --- | Exon Cassette | e27-42 | up | 1,58 | --- |
| B4GALT3 | Intron Retention | e2 | up | 1,58 | ENSG00000 158850 |
| SMNDC1 | Intron Retention | e1 | up | 1,58 | ENSG00000 119953 |
| SLC39A13 | Alter. First Exon | e5/e6-7 | down (e6-7) | 1,58 | ENSG00000 165915 |
| IRAK4 | Exon Cassette | e2-3,e5 | up | 1,58 | ENSG00000 198001 |
| SLC4A8 | ter. Terminal Exc | e10-23,e25- | own (e10-23,e25-3( | 1,58 | ENSG00000 050438 |
| HELB | Exon Cassette | e5 | up | 1,58 | ENSG00000 127311 |
| SYT1 | Complex | e5-8 | up | 1,58 | ENSG00000 067715 |
| COG3 | ter. Terminal Exc | 2/e13,e14-2 | down (e13,e14-23) | 1,58 | ENSG00000 136152 |
| ATXN3 | Exon Cassette | 2,e3-5,e8-1( | down | 1,58 | ENSG00000 066427 |
| CIB1 | Complex | e2 | down | 1,58 | ENSG00000 185043 |
| FAM65A | Complex | e15-24 | up | 1,58 | ENSG00000 039523 |
| FAM96B | Intron Retention | e1,e2 | up | 1,58 | ENSG00000 166595 |
| ERBB2 | Alter. First Exon | e7-8/e9 | down (e7-8) | 1,58 | ENSG00000 141736 |
| CNOT3 | Intron Retention | e18 | up | 1,58 | ENSG00000 088038 |
| PAPOLG | Exon Cassette | e22 | up | 1,58 | ENSG00000 115421 |
| BCL2L11 | Exon Cassette | e11 | up | 1,58 | ENSG00000 153094 |

| | | | | | |
|---|---|---|---|---|---|
| FMNL2 | Complex | e1,e2-16 | down | 1,58 | ENSG00000157827 |
| DTNB | Exon Cassette | e12 | up | 1,58 | ENSG00000138101 |
| GPR75 // GPR7 | Alter. Donor Site | e3 | up | 1,58 | ENSG00000115239 // ENSG00000119737 |
| FAM161A | Exon Cassette | e5,e6 | up | 1,58 | ENSG00000170264 |
| CDC142 // MRPL | Alter. First Exon | e1-8/e10 | up (e1-8) | 1,58 | ENSG00000135637 // ENSG00000204822 |
| BAZ2B | Complex | e16,e17 | up | 1,58 | ENSG00000123636 |
| APP | Exon Cassette | e11 | up | 1,58 | ENSG00000142192 |
| U2AF1 | Intron Retention | e2 | up | 1,58 | ENSG00000160201 |
| MED15 | Complex | e11-12 | up | 1,58 | ENSG00000099917 |
| TIMP3 | Complex | e1-5 | up | 1,58 | ENSG00000100234 |
| PATZ1 | Exon Cassette | e6 | up | 1,58 | ENSG00000100105 |
| SACM1L | Alter. First Exon | e1,e3-14/e15 | down (e1,e3-14) | 1,58 | ENSG00000211456 |
| SACM1L | Alter. First Exon | 1-4,e6-14/e1 | down (e1-4,e6-14) | 1,58 | ENSG00000211456 |
| KCTD6 | Alter. First Exon | e1/e2 | up (e2) | 1,58 | ENSG00000168301 |
| CEP63 | Exon Cassette | e14,e15 | up | 1,58 | ENSG00000182923 |
| NT5DC2 | Alter. Donor Site | e1 | up | 1,58 | ENSG00000168268 |
| ARHGEF28 | ter. Terminal Exo | 5/e16-36,e3 | up (e16-36,e38) | 1,58 | ENSG00000214944 |
| PHF15 | Alter. First Exon | e1/e3 | up (e1) | 1,58 | ENSG00000043143 |
| ARHGAP26 | Alter. Donor Site | e23 | up | 1,58 | ENSG00000145819 |
| FBN2 | ter. Terminal Exo | 4/e15,e17-4 | up (e15,e17-42) | 1,58 | ENSG00000138829 |
| GMDS-AS1 | Complex | e4-7 | down | 1,58 | ENSG00000250903 |
| ANKS1A | Exon Cassette | e28 | up | 1,58 | ENSG00000064999 |
| TRMT11 | ter. Terminal Exo | e11/e12-16 | down (e12-16) | 1,58 | ENSG00000066651 |
| PEX6 | ter. Terminal Exo | 8/e9,e10-18 | down (e9,e10-18) | 1,58 | ENSG00000124587 |

| | | | | | |
|---|---|---|---|---|---|
| SYNE1 | Alter. First Exon | 19-88,e90-1 | e8-15,e19-88,e90-1 | 1,58 | ENSG00000 131018 |
| SYNE1 | Alter. First Exon | e77-88,e90-1 | e19-75,e77-88,e90- | 1,58 | ENSG00000 131018 |
| RGL2 | Exon Cassette | e2 | up | 1,58 | ENSG00000 237441 |
| SNK2B // LY6G5 | Complex | e2/e9-10 | up (e9-10) | 1,58 | ENSG00000 204435 // ENSG00000 240053 |
| GNL1 | Complex | e1-7/e8 | down (e1-7) | 1,58 | ENSG00000 204590 |
| SDK1 | Alter. First Exon | e34-40/e41 | down (e34-40) | 1,58 | ENSG00000 146555 |
| DMTF1 | lter. Acceptor Sit | e14 | up | 1,58 | ENSG00000 135164 |
| ETV1 | Exon Cassette | e11 | down | 1,58 | ENSG00000 006468 |
| FAM126A | Exon Cassette | e11 | down | 1,58 | ENSG00000 122591 |
| ZNF746 | Complex | e1-4 | up | 1,58 | ENSG00000 181220 |
| FAM49B | Complex | e5-6/e8 | up (e5-6) | 1,58 | ENSG00000 153310 |
| NUDT2 | Exon Cassette | e2 | up | 1,58 | ENSG00000 164978 |
| CBWD1 | ter. Terminal Exo | e5,e7-11,e14 | up (e4) | 1,58 | ENSG00000 172785 |
| DDX3X | ter. Terminal Exo | /e5-12,e14- | up (e4) | 1,58 | ENSG00000 215301 |
| DDX3X | ter. Terminal Exo | e5,e6-12,e14 | up (e4) | 1,58 | ENSG00000 215301 |
| DDX3X | ter. Terminal Exo | 6-12,e14-15 | up (e4) | 1,58 | ENSG00000 215301 |
| HDAC6 | Intron Retention | e15 | up | 1,58 | ENSG00000 094631 |
| ALG13 | Alter. First Exon | e1/e2 | down (e1) | 1,58 | ENSG00000 101901 |
| AMPD2 | Exon Cassette | e5 | down | 1,57 | ENSG00000 116337 |
| HSPB11 | Intron Retention | e5 | up | 1,57 | ENSG00000 081870 |
| FUBP1 | Exon Cassette | e3 | up | 1,57 | ENSG00000 162613 |
| STIM1 | Alter. Donor Site | e12 | down | 1,57 | ENSG00000 167323 |
| TMEM135 | Complex | e1-4,e7 | down | 1,57 | ENSG00000 166575 |
| DYRK4 | ter. Terminal Exo | 9/e10,e11-1 | up (e10,e11-16) | 1,57 | ENSG00000 010219 |
| ATXN3 | Exon Cassette | e2-6,e8-10 | down | 1,57 | ENSG00000 066427 |

| | | | | | |
|---|---|---|---|---|---|
| ATXN3 | Exon Cassette | 2-3,e5-6,e8-1 | down | 1,57 | ENSG00000 066427 |
| ATXN3 | Exon Cassette | e3,e4-5 | down | 1,57 | ENSG00000 066427 |
| TUBGCP4 | Exon Cassette | e14 | up | 1,57 | ENSG00000 137822 |
| BBS2 | Intron Retention | e9 | up | 1,57 | ENSG00000 125124 |
| .SCR3 // TMEM2 | Alter. First Exon | e1-3,e6-7/e5 | up (e1-3,e6-7) | 1,57 | ENSG00000 187838 // ENSG00000 205544 |
| ZNF18 | Alter. First Exon | e1/e2-4 | up (e2-4) | 1,57 | ENSG00000 154957 |
| EXOC7 | Exon Cassette | e7-8 | down | 1,57 | ENSG00000 182473 |
| CSNK1D | Complex | e9,e10 | up | 1,57 | ENSG00000 141551 |
| ATRAID | Complex | e1/e2 | down (e1) | 1,57 | ENSG00000 138085 |
| SNX17 | Exon Cassette | e4 | up | 1,57 | ENSG00000 115234 |
| BCS1L | Complex | e2 | up | 1,57 | ENSG00000 074582 |
| GIGYF2 | ter. Terminal Exo | 9,e10-13,e1 | up (e8) | 1,57 | ENSG00000 204120 |
| GIGYF2 | ter. Terminal Exo | 9,e10-13,e1 | up (e8) | 1,57 | ENSG00000 204120 |
| PREB | Intron Retention | e3 | up | 1,57 | ENSG00000 138073 |
| MAP4K3 | Exon Cassette | e15 | up | 1,57 | ENSG00000 011566 |
| OBSL1 | ter. Terminal Exo | 9/e10,e11-1 | up (e9) | 1,57 | ENSG00000 124006 |
| SEC23B | Complex | e1,e2 | up | 1,57 | ENSG00000 101310 |
| GNAS | Alter. First Exon | e2/e8 | up (e2) | 1,57 | ENSG00000 087460 |
| SNHG17 | Alter. First Exon | e1-3/e4 | down (e1-3) | 1,57 | ENSG00000 196756 |
| DIDO1 | ter. Terminal Exo | 3/e9,e10-17, | up (e7-8) | 1,57 | ENSG00000 101191 |
| // PRR5 // PRR5 | Exon Cassette | e14 | up | 1,57 | ENSG00000 186654 // ENSG00000 241484 // ENSG00000 248405 |
| CLTCL1 | Alter. Donor Site | e24 | down | 1,57 | ENSG00000 070371 |
| IFT122 | Intron Retention | e28 | up | 1,57 | ENSG00000 163913 |

| | | | | | |
|---|---|---|---|---|---|
| CAMKV | Complex | e7-8 | up | 1,57 | ENSG00000 164076 |
| ARHGAP10 | Alter. First Exon | e6-18/e19 | up (e6-18) | 1,57 | ENSG00000 071205 |
| PHYKPL | Complex | e7-8 | down | 1,57 | ENSG00000 175309 |
| IYMAI // PLAGL1 | Exon Cassette | e8-9 | up | 1,57 | ENSG00000 118495 |
| GNL1 | Complex | e1-8 | down | 1,57 | ENSG00000 204590 |
| Sep-07 | Complex | e1,e3 | down | 1,57 | ENSG00000 122545 |
| LANCL2 | Complex | e3-4 | down | 1,57 | ENSG00000 132434 |
| ELN | Complex | e18,e19-26 | up | 1,57 | ENSG00000 049540 |
| KMT2C | Intron Retention | e59 | up | 1,57 | ENSG00000 055609 |
| --- | Alter. First Exon | e1-3/e4 | down (e1-3) | 1,57 | --- |
| RFX3 | Exon Cassette | e8 | down | 1,57 | ENSG00000 080298 |
| MPDZ | Exon Cassette | e27-28 | up | 1,57 | ENSG00000 107186 |
| BGT1 // RALGD | Alter. First Exon | e9/e10 | up (e9) | 1,57 | ENSG00000 148288 // ENSG00000 160271 |
| SEC16A | Exon Cassette | e32-33 | up | 1,57 | ENSG00000 148396 |
| ZMAT1 | Exon Cassette | e6,e7-9 | up | 1,57 | ENSG00000 166432 |
| BCAP31 | Alter. First Exon | e2/e3 | down (e3) | 1,57 | ENSG00000 185825 |
| --- | Intron Retention | e6 | up | 1,57 | --- |
| STRIP1 | Intron Retention | e18 | up | 1,56 | ENSG00000 143093 |
| DCAF6 | Exon Cassette | e14 | up | 1,56 | ENSG00000 143164 |
| MDM4 | Exon Cassette | e8-12 | down | 1,56 | ENSG00000 198625 |
| RERE | Alter. First Exon | e1-9/e10 | down (e1-9) | 1,56 | ENSG00000 142599 |
| RERE | Alter. First Exon | e3-9/e10 | down (e3-9) | 1,56 | ENSG00000 142599 |
| TTC39A | ter. Terminal Exo | e17-18/e19 | down (e17-18) | 1,56 | ENSG00000 085831 |
| ZRANB2 | Intron Retention | e9,e10 | up | 1,56 | ENSG00000 132485 |
| HSPA14 | Exon Cassette | e4 | down | 1,56 | ENSG00000 187522 |
| ZNF248 | ter. Terminal Exo | e7/e9,e10 | down (e7) | 1,56 | ENSG00000 198105 |

| | | | | | |
|---|---|---|---|---|---|
| ATE1 | Exon Cassette | e3-4 | down | 1,56 | ENSG00000107669 |
| C11orf30 | Exon Cassette | e18 | up | 1,56 | ENSG00000158636 |
| RNH1 | lter. Acceptor Sit | e4 | up | 1,56 | ENSG00000023191 |
| TMEM218 | lter. Acceptor Sit | e5 | up | 1,56 | ENSG00000150433 |
| WNK1 | Exon Cassette | e9 | up | 1,56 | ENSG00000060237 |
| PPHLN1 | Alter. First Exon | e1,e3-8/e9 | up (e1,e3-8) | 1,56 | ENSG00000134283 |
| TSC22D1 | Alter. First Exon | e1/e3 | down (e1) | 1,56 | ENSG00000102804 |
| SUCLA2 | Alter. First Exon | e1-2,e5/e3 | up (e1-2,e5) | 1,56 | ENSG00000136143 |
| SCFD1 | Exon Cassette | e4-5 | down | 1,56 | ENSG00000092108 |
| JKAMP | Complex | e2/e3 | down (e2) | 1,56 | ENSG00000050130 |
| ATXN3 | Exon Cassette | e2-6,e8-11 | down | 1,56 | ENSG00000066427 |
| ATXN3 | Exon Cassette | 2,e4-6,e8-1( | down | 1,56 | ENSG00000066427 |
| ATXN3 | Exon Cassette | ,e3,e5-6,e8- | down | 1,56 | ENSG00000066427 |
| CCNDBP1 | ter. Terminal Exc | e9/e10,e11 | up (e10,e11) | 1,56 | ENSG00000166946 |
| SLTM | Intron Retention | e7 | up | 1,56 | ENSG00000137776 |
| NAGPA | Exon Cassette | e8 | up | 1,56 | ENSG00000103174 |
| TMEM170A | Alter. First Exon | e1/e2 | up (e1) | 1,56 | ENSG00000166822 |
| PLEKHM1P | Alter. Donor Site | e17 | down | 1,56 | ENSG00000214176 |
| CSNK1D | Alter. First Exon | e1,e3/e4 | down (e1,e3) | 1,56 | ENSG00000141551 |
| LTBP4 | Alter. First Exon | e2-4/e5 | down (e2-4) | 1,56 | ENSG00000090006 |
| --- | Intron Retention | e4 | up | 1,56 | --- |
| ATP5SL | Exon Cassette | e10,e11 | down | 1,56 | ENSG00000105341 |
| 1S1 // IL4I1 // NU | Complex | e2-3 | up | 1,56 | ENSG00000104951 // ENSG00000204673 // ENSG00000213024 |
| MPHOSPH10 | ter. Terminal Exc | e5/e6-11 | up (e5) | 1,56 | ENSG00000124383 |

| | | | | | |
|---|---|---|---|---|---|
| CARF | Exon Cassette | e4 | up | 1,56 | ENSG00000138380 |
| SDC1 | Complex | e4-6/e7 | up (e4-6) | 1,56 | ENSG00000115884 |
| SEC23B | Complex | e1-2 | up | 1,56 | ENSG00000101310 |
| DDX27 | Intron Retention | e11-12 | up | 1,56 | ENSG00000124228 |
| --- | Alter. First Exon | e2/e3 | up (e2) | 1,56 | --- |
| NAPB | Exon Cassette | e3,e4 | down | 1,56 | ENSG00000125814 |
| DIDO1 | ter. Terminal Exc | e17-18/e19 | up (e17-18) | 1,56 | ENSG00000101191 |
| DIDO1 | ter. Terminal Exc | 8/e9-17,e1 | up (e8) | 1,56 | ENSG00000101191 |
| ARFRP1 | Alter. First Exon | e1/e2 | down (e1) | 1,56 | ENSG00000101246 |
| SLC2A11 | ter. Terminal Exc | e11/e12-15 | up (e12-15) | 1,56 | ENSG00000133460 |
| TUG1 | Exon Cassette | e2 | up | 1,56 | ENSG00000253352 |
| SMTN | Complex | e19-21 | down | 1,56 | ENSG00000183963 |
| SGSM3 | Alter. First Exon | e1-17/e18 | up (e1-17) | 1,56 | ENSG00000100359 |
| TXNRD2 | Alter. First Exon | e2/e3 | down (e3) | 1,56 | ENSG00000184470 |
| GSTT1 | Exon Cassette | e3,e4 | down | 1,56 | ENSG00000184674 |
| DAG1 | Alter. First Exon | ,e4-6/e7,e8- | down (e1,e4-6) | 1,56 | ENSG00000173402 |
| SLC4A4 | Alter. First Exon | e2-3/e4 | down (e2-3) | 1,56 | ENSG00000080493 |
| ATP8A1 | ualy Exclusive E | e7/e8 | up (e7) | 1,56 | ENSG00000124406 |
| SLC10A7 | Complex | e4/e7 | down (e7) | 1,56 | ENSG00000120519 |
| WASF1 | Exon Cassette | e3,e4 | up | 1,56 | ENSG00000112290 |
| ZNRD1 | Intron Retention | e1 | up | 1,56 | ENSG00000066379 |
| GSTK1 | Intron Retention | e4 | up | 1,56 | ENSG00000197448 |
| RNF32 | Exon Cassette | e6 | up | 1,56 | ENSG00000105982 |
| GUSB | Intron Retention | e5 | down | 1,56 | ENSG00000169919 |
| C7orf49 | Alter. First Exon | e1-2,e4/e5 | down (e1-2,e4) | 1,56 | ENSG00000122783 |
| FUT10 | ter. Terminal Exc | e5/e7,e8 | up (e5) | 1,56 | ENSG00000172728 |

| | | | | | |
|---|---|---|---|---|---|
| DAB2IP | lter. Acceptor Sit | e16 | down | 1,56 | ENSG00000136848 |
| CBWD1 | ter. Terminal Exo | e5,e7-11,e1 | up (e4) | 1,56 | ENSG00000172785 |
| STS | Alter. First Exon | e2/e3 | down (e2) | 1,56 | ENSG00000101846 |
| STAG2 | Complex | e1/e5-15 | up (e5-15) | 1,56 | ENSG00000101972 |
| MAP7D2 | Complex | e7,e9 | up | 1,56 | ENSG00000184368 |
| MAP7D2 | Exon Cassette | e7 | up | 1,56 | ENSG00000184368 |
| MORF4L2 | Exon Cassette | e4 | up | 1,56 | ENSG00000123562 |
| LRP8 | Exon Cassette | e19 | up | 1,55 | ENSG00000157193 |
| ZNF692 | Complex | e4 | up | 1,55 | ENSG00000171163 |
| WAC | Complex | e5/e6 | up (e5) | 1,55 | ENSG00000095787 |
| WAC | Exon Cassette | e5 | up | 1,55 | ENSG00000095787 |
| C11orf63 | ter. Terminal Exo | e4/e5-10 | down (e5-10) | 1,55 | ENSG00000109944 |
| SLC25A3 | Intron Retention | e2,e3 | up | 1,55 | ENSG00000075415 |
| RSRC2 | Exon Cassette | e5 | up | 1,55 | ENSG00000111011 |
| CDK2AP1 | Alter. First Exon | e1/e4 | up (e1) | 1,55 | ENSG00000111328 |
| CDK2AP1 | Alter. First Exon | e1/e3 | up (e1) | 1,55 | ENSG00000111328 |
| DPP8 | Exon Cassette | e19-20 | up | 1,55 | ENSG00000074603 |
| MGRN1 | Exon Cassette | e11,e12 | up | 1,55 | ENSG00000102858 |
| TNRC6A | Alter. First Exon | e14/e15 | up (e14) | 1,55 | ENSG00000090905 |
| COQ9 | Intron Retention | e4 | up | 1,55 | ENSG00000088682 |
| MTSS1L | Complex | e9,e10-14 | up | 1,55 | ENSG00000132613 |
| SPHK1 | Alter. First Exon | e5-8/e6 | down (e5-8) | 1,55 | ENSG00000176170 |
| INO80C | Exon Cassette | e2,e3 | down | 1,55 | ENSG00000153391 |
| NFIC | Exon Cassette | e10,e11 | up | 1,55 | ENSG00000141905 |
| PRMT1 | Exon Cassette | e2,e3 | up | 1,55 | ENSG00000126457 |
| MED25 | Complex | e8/e9 | down (e9) | 1,55 | ENSG00000104973 |

| | | | | | |
|---|---|---|---|---|---|
| ZNF772 | Exon Cassette | e2,e3-4 | up | 1,55 | ENSG00000 197128 |
| TTC31 | Intron Retention | e5 | up | 1,55 | ENSG00000 115282 |
| DNAJB2 | Intron Retention | e3 | up | 1,55 | ENSG00000 135924 |
| GIGYF2 | ter. Terminal Exo | 0-13,e17-33 | up (e8) | 1,55 | ENSG00000 204120 |
| ALS2 | ter. Terminal Exo | e23/e25-35 | up (e23) | 1,55 | ENSG00000 003393 |
| ALS2 | ter. Terminal Exo | 23/e24,e25-3 | up (e23) | 1,55 | ENSG00000 003393 |
| E1 // NFS1 // RB | ter. Terminal Exo | 21-23/e24-3 | up (e21-23) | 1,55 | ENSG00000 214078 // ENSG00000 244005 // ENSG00000 244462 |
| USP16 | Intron Retention | e13,e14 | up | 1,55 | ENSG00000 156256 |
| SCAF4 | ter. Terminal Exo | e19/e20 | up (e19) | 1,55 | ENSG00000 156304 |
| RPP14 | Alter. First Exon | e1,e3/e2 | up (e1,e3) | 1,55 | ENSG00000 163684 |
| GLB1 // TMPPE | Exon Cassette | e5 | down | 1,55 | ENSG00000 170266 // ENSG00000 188167 |
| IP6K2 | ter. Terminal Exo | e11/e12-15 | up (e11) | 1,55 | ENSG00000 068745 |
| INTU | Exon Cassette | e7,e8 | up | 1,55 | ENSG00000 164066 |
| TMEM128 | Intron Retention | e3 | up | 1,55 | ENSG00000 132406 |
| C31A // THAP9-A | Exon Cassette | e19-20 | down | 1,55 | ENSG00000 138674 // ENSG00000 251022 |
| TSPAN17 | Intron Retention | e4 | up | 1,55 | ENSG00000 048140 |
| orf165 // SLC35 | Alter. First Exon | e1-11/e12 | down (e1-11) | 1,55 | ENSG00000 164414 // ENSG00000 213204 |
| PEX6 | Complex | e10/e11-12 | down (e11-12) | 1,55 | ENSG00000 124587 |
| RARS2 | Alter. Donor Site | e1 | down | 1,55 | ENSG00000 146282 |
| CRCP | Exon Cassette | e2,e4-5 | down | 1,55 | ENSG00000 241258 |
| STAU2 | ter. Terminal Exo | 8/e19,e20-2 | up (e18) | 1,55 | ENSG00000 040341 |

| | | | | | |
|---|---|---|---|---|---|
| FUBP3 | Exon Cassette | e3 | up | 1,55 | ENSG00000107164 |
| POMT1 | Exon Cassette | e4-5 | down | 1,55 | ENSG00000130714 |
| CBWD1 | Exon Cassette | e2 | up | 1,55 | ENSG00000172785 |
| MPDZ | Exon Cassette | e27 | up | 1,55 | ENSG00000107186 |
| XIST | lter. Acceptor Sit | e9 | down | 1,55 | ENSG00000229807 |
| AKR1A1 | Exon Cassette | e2-3 | up | 1,54 | ENSG00000117448 |
| AHCYL1 | lter. Acceptor Sit | e18 | up | 1,54 | ENSG00000168710 |
| MOV10 | Complex | e3,e4 | up | 1,54 | ENSG00000155363 |
| RGL1 | Exon Cassette | e5-8 | down | 1,54 | ENSG00000143344 |
| POMGNT1 | Intron Retention | e13 | up | 1,54 | ENSG00000085998 |
| ZZZ3 | Intron Retention | e10 | up | 1,54 | ENSG00000036549 |
| CCT3 | Exon Cassette | e2,e4 | up | 1,54 | ENSG00000163468 |
| CDH23 | ter. Terminal Ex | 3/e34,e35-3 | down (e34,e35-38) | 1,54 | ENSG00000107736 |
| ABI1 | Exon Cassette | e11 | up | 1,54 | ENSG00000136754 |
| TYSND1 | Alter. First Exon | e1-2/e3 | down (e1-2) | 1,54 | ENSG00000156521 |
| LDLRAD3 | Exon Cassette | e2 | down | 1,54 | ENSG00000179241 |
| VEGFB | lter. Acceptor Sit | e6 | up | 1,54 | ENSG00000173511 |
| RNH1 | Exon Cassette | e3 | up | 1,54 | ENSG00000023191 |
| NAP1L4 | Exon Cassette | e16 | up | 1,54 | ENSG00000205531 |
| PPHLN1 | Alter. First Exon | e1,e4-8/e9 | up (e1,e4-8) | 1,54 | ENSG00000134283 |
| AM101A // ZNF6 | Exon Cassette | e4-5 | up | 1,54 | ENSG00000178882 // ENSG00000179195 |
| RNF41 | Exon Cassette | e5 | up | 1,54 | ENSG00000181852 |
| SETDB2 | Exon Cassette | e6 | down | 1,54 | ENSG00000136169 |
| JKAMP | Complex | e1,e2 | down | 1,54 | ENSG00000050130 |
| TINF2 | Intron Retention | e4 | up | 1,54 | ENSG00000092330 |

| | | | | | |
|---|---|---|---|---|---|
| NEK9 | Intron Retention | e12 | up | 1,54 | ENSG00000 119638 |
| ATXN3 | Exon Cassette | e3-6,e8-10 | down | 1,54 | ENSG00000 066427 |
| ATXN3 | Exon Cassette | 3,e5-6,e8-1( | down | 1,54 | ENSG00000 066427 |
| ATXN3 | Exon Cassette | ,e3,e5-6,e8- | down | 1,54 | ENSG00000 066427 |
| ATXN3 | Exon Cassette | e2,e3,e5-11 | down | 1,54 | ENSG00000 066427 |
| ATXN3 | Exon Cassette | 3,e4-6,e8-1( | down | 1,54 | ENSG00000 066427 |
| NFE2L1 | Exon Cassette | e6 | up | 1,54 | ENSG00000 082641 |
| ZNF606 | Exon Cassette | e5 | up | 1,54 | ENSG00000 166704 |
| CENPO | Complex | e1-2 | up | 1,54 | ENSG00000 138092 |
| SPTBN1 | Alter. First Exon | e2-3/e4 | down (e2-3) | 1,54 | ENSG00000 115306 |
| NAGK | Intron Retention | e3 | up | 1,54 | ENSG00000 124357 |
| PUS10 | Exon Cassette | e8 | up | 1,54 | ENSG00000 162927 |
| ANKRD36 | Exon Cassette | e4,e5-6 | up | 1,54 | ENSG00000 135976 |
| PCIF1 | Intron Retention | e16 | up | 1,54 | ENSG00000 100982 |
| LIMK2 | Alter. First Exon | e1-2/e3 | up (e1-2) | 1,54 | ENSG00000 182541 |
| SMC4 | Intron Retention | e1,e2 | up | 1,54 | ENSG00000 113810 |
| EOMES | lter. Acceptor Sit | e7 | up | 1,54 | ENSG00000 163508 |
| SLC26A6 | Intron Retention | e17,e18 | down | 1,54 | ENSG00000 225697 |
| THAP6 | ter. Terminal Exc | e4/e5 | up (e4) | 1,54 | ENSG00000 174796 |
| ARHGAP10 | Alter. First Exon | e1-18/e19 | up (e1-18) | 1,54 | ENSG00000 071205 |
| C4orf21 | ter. Terminal Exc | e21/e22-31 | up (e22-31) | 1,54 | ENSG00000 138658 |
| FCHO2 | Alter. First Exon | e8-10,e12-1 | b (e1-6,e8-10,e12-1 | 1,54 | ENSG00000 157107 |
| GPR98 | Alter. First Exon | e24-26,e28- | n (e1-22,e24-26,e28 | 1,54 | ENSG00000 164199 |
| SAP30L | Complex | e2,e4 | down | 1,54 | ENSG00000 164576 |
| BRD9 | Complex | e22-23/e24 | up (e22-23) | 1,54 | ENSG00000 028310 |
| ATF6B | Complex | e36-46 | down | 1,54 | ENSG00000 213676 |

| | | | | | |
|---|---|---|---|---|---|
| CUTA | Intron Retention | e1-2 | down | 1,54 | ENSG00000112514 |
| C7orf49 | Alter. First Exon | e2,e4/e5 | down (e2,e4) | 1,54 | ENSG00000122783 |
| NEK6 | Alter. First Exon | e2/e4 | down (e2) | 1,54 | ENSG00000119408 |
| CBWD1 | ter. Terminal Exo | 5,e7-11,e14 | up (e4) | 1,54 | ENSG00000172785 |
| ZCCHC6 | Intron Retention | e13 | up | 1,54 | ENSG00000083223 |
| DDX31 | Alter. First Exon | e1-2/e3 | up (e1-2) | 1,54 | ENSG00000125485 |
| PQBP1 | Intron Retention | e7 | up | 1,54 | ENSG00000102103 |
| MAGED1 | Intron Retention | e9 | up | 1,54 | ENSG00000179222 |
| FGD1 | Intron Retention | e15 | up | 1,54 | ENSG00000102302 |
| AIFM1 | Exon Cassette | e2,e4-11 | up | 1,54 | ENSG00000156709 |
| SRRM1 | Exon Cassette | e15 | down | 1,53 | ENSG00000133226 |
| KIAA1324 | Complex | e4-6 | down | 1,53 | ENSG00000116299 |
| CDK18 | Intron Retention | e13 | down | 1,53 | ENSG00000117266 |
| RCC2 | Alter. First Exon | e1/e2 | down (e1) | 1,53 | ENSG00000179051 |
| ERCC6-PGBD3 | Intron Retention | e8,e9 | up | 1,53 | ENSG00000225830 // ENSG00000243251 // ENSG00000258838 |
| SORCS1 | ter. Terminal Exo | e25/e26 | up (e25) | 1,53 | ENSG00000108018 |
| CD151 | Alter. First Exon | e1,e4/e3 | down (e1,e4) | 1,53 | ENSG00000177697 |
| HIPK3 | Exon Cassette | e15 | up | 1,53 | ENSG00000110422 |
| DGKZ | Alter. First Exon | -9,e11-29/e | up (e3-9,e11-29) | 1,53 | ENSG00000149091 |
| DKK3 | Complex | e5-10 | up | 1,53 | ENSG00000050165 |
| LPXN | Exon Cassette | e4 | up | 1,53 | ENSG00000110031 |
| HDAC7 | Intron Retention | e24 | up | 1,53 | ENSG00000061273 |
| RIMBP2 | Exon Cassette | e5-6 | up | 1,53 | ENSG00000060709 |
| IFT88 | Complex | e2,e3 | up | 1,53 | ENSG00000032742 |

| | | | | | |
|---|---|---|---|---|---|
| SDR39U1 | Intron Retention | e3 | up | 1,53 | ENSG00000100445 |
| GCH1 | Complex | e6-7 | down | 1,53 | ENSG00000131979 |
| SMEK1 | ter. Terminal Exo | 9,e10-11,e1 | up (e8) | 1,53 | ENSG00000100796 |
| ATXN3 | Complex | 2-6,e8-10/e1 | down (e2-6,e8-10) | 1,53 | ENSG00000066427 |
| ATXN3 | Exon Cassette | 2-3,e6,e8-1( | down | 1,53 | ENSG00000066427 |
| ATXN3 | Exon Cassette | 3,e5-6,e8-1 | down | 1,53 | ENSG00000066427 |
| COG1 | Exon Cassette | e15 | up | 1,53 | ENSG00000166685 |
| CANT1 | Intron Retention | e3 | down | 1,53 | ENSG00000171302 |
| TMEM241 | Exon Cassette | e9 | up | 1,53 | ENSG00000134490 |
| / RPL17 // RPL1 | Complex | e1,e2 | up | 1,53 | ENSG00000177576 // ENSG00000215472 // ENSG00000265681 |
| MBD1 | Exon Cassette | e13 | up | 1,53 | ENSG00000141644 |
| PNPLA6 | Alter. First Exon | e2,e4/e5 | down (e2,e4) | 1,53 | ENSG00000032444 |
| ZNF136 | Exon Cassette | e2 | down | 1,53 | ENSG00000196646 |
| LTBP4 | Exon Cassette | e25 | up | 1,53 | ENSG00000090006 |
| BCL2L11 | Complex | 2/e9,e11,e1 | up (e9,e11,e13) | 1,53 | ENSG00000153094 |
| SDC1 | Alter. First Exon | e1-2/e3 | up (e1-2) | 1,53 | ENSG00000115884 |
| EPB41L1 | Alter. First Exon | e3-6/e8 | up (e3-6) | 1,53 | ENSG00000088367 |
| ARFGEF2 | Exon Cassette | e24 | up | 1,53 | ENSG00000124198 |
| E1 // NFS1 // RB | ter. Terminal Exo | ,e24-38/e22 | down (e21,e24-38) | 1,53 | ENSG00000214078 // ENSG00000244005 // ENSG00000244462 |
| KCTD17 | Complex | e8-9 | down | 1,53 | ENSG00000100379 |
| PXK | Exon Cassette | e2 | up | 1,53 | ENSG00000168297 |
| C3orf52 | Exon Cassette | e4,e7 | up | 1,53 | ENSG00000114529 |

| | | | | | |
|---|---|---|---|---|---|
| MAP3K13 | ter. Terminal Exc | e12/e13-19 | down (e13-19) | 1,53 | ENSG00000073803 |
| FOXP1 | Complex | e13/e14,e17 | down (e14,e17) | 1,53 | ENSG00000114861 |
| FOXP1 | Exon Cassette | e28 | up | 1,53 | ENSG00000114861 |
| TBL1XR1 | Alter. First Exon | e1/e3 | up (e1) | 1,53 | ENSG00000177565 |
| APBB2 | Alter. First Exon | e1-13/e14 | up (e1-13) | 1,53 | ENSG00000163697 |
| LIN54 | Alter. Donor Site | e3 | down | 1,53 | ENSG00000189308 |
| C31A // THAP9-A | ter. Terminal Exc | ,e28-29,e31 | (e3,e8-26,e28-29,e3 | 1,53 | ENSG00000138674 // ENSG00000251022 |
| FCHO2 | Alter. First Exon | -10,e12-17/e | up (e1-10,e12-17) | 1,53 | ENSG00000157107 |
| ARHGAP26 | Complex | e23,e24-25 | up | 1,53 | ENSG00000145819 |
| TDP2 | Alter. Donor Site | e2 | up | 1,53 | ENSG00000111802 |
| PNISR | Intron Retention | e5 | up | 1,53 | ENSG00000132424 |
| AHI1 | Exon Cassette | e2,e3 | up | 1,53 | ENSG00000135541 |
| ATAT1 | ter. Terminal Exc | 10-11/e12-1 | down (e10-11) | 1,53 | ENSG00000137343 |
| HCG18 | Exon Cassette | e2 | up | 1,53 | ENSG00000231074 |
| PSMG3-AS1 | Exon Cassette | e4 | up | 1,53 | ENSG00000230487 |
| SDK1 | Alter. First Exon | e27-40/e41 | down (e27-40) | 1,53 | ENSG00000146555 |
| ELN | Exon Cassette | e22,e23 | up | 1,53 | ENSG00000049540 |
| ADAM22 | ter. Terminal Exc | 29/e30,e31-3 | up (e29) | 1,53 | ENSG00000008277 |
| EZH2 | Intron Retention | e13 | up | 1,53 | ENSG00000106462 |
| FAM49B | Complex | e1,e4/e2 | down (e2) | 1,53 | ENSG00000153310 |
| MSL3 | Alter. First Exon | e1/e3 | down (e1) | 1,53 | ENSG00000005302 |
| ACADM | Exon Cassette | e2 | up | 1,52 | ENSG00000117054 |
| RABGGTB | Complex | e4,e5-6 | up | 1,52 | ENSG00000137955 |
| INTS3 | Intron Retention | e5 | up | 1,52 | ENSG00000143624 |
| MRPL55 | Complex | e1,e2 | up | 1,52 | ENSG00000162910 |

| | | | | | |
|---|---|---|---|---|---|
| SHOC2 | Exon Cassette | e2-3 | down | 1,52 | ENSG00000108061 |
| LRRC20 | Alter. First Exon | e2/e3 | up (e2) | 1,52 | ENSG00000172731 |
| AMBRA1 | Exon Cassette | e10,e11 | up | 1,52 | ENSG00000110497 |
| MRE11A | ter. Terminal Exo | e10-17,e19- | up (e9) | 1,52 | ENSG00000020922 |
| EXPH5 | Exon Cassette | e7 | down | 1,52 | ENSG00000110723 |
| DPAGT1 | Intron Retention | e10 | up | 1,52 | ENSG00000172269 |
| SETD8 | Alter. First Exon | e1/e2,e3 | up (e1) | 1,52 | ENSG00000183955 |
| SETD8 | Alter. First Exon | e1/e4 | up (e1) | 1,52 | ENSG00000183955 |
| ATXN2 | Exon Cassette | e26 | up | 1,52 | ENSG00000204842 |
| EIF2B1 | ter. Terminal Exo | e5/e6-9 | up (e5) | 1,52 | ENSG00000111361 |
| IFT88 | Exon Cassette | e2-3 | up | 1,52 | ENSG00000032742 |
| ATXN3 | Exon Cassette | e3,e4-10 | down | 1,52 | ENSG00000066427 |
| ATXN3 | Exon Cassette | 2,e3,e6,e8-1 | down | 1,52 | ENSG00000066427 |
| PML | ter. Terminal Exo | e4/e5,e6-8 | up (e5,e6-8) | 1,52 | ENSG00000140464 |
| PML | Exon Cassette | e5 | up | 1,52 | ENSG00000140464 |
| ADPGK | Exon Cassette | e2-3 | up | 1,52 | ENSG00000159322 |
| TM2D3 | ter. Terminal Exo | e4/e5,e7 | up (e4) | 1,52 | ENSG00000184277 |
| TELO2 | Complex | e13-15/e16 | up (e13-15) | 1,52 | ENSG00000100726 |
| CUN1D3 // ERI2 | lter. Acceptor Sit | e15 | down | 1,52 | ENSG00000188215 // ENSG00000196678 |
| PIPB3 // SLC7A5 | Alter. First Exon | e11/e12 | up (e11) | 1,52 | ENSG00000169246 // ENSG00000258186 |
| DOK4 | Exon Cassette | e4 | down | 1,52 | ENSG00000125170 |
| VAC14 | ter. Terminal Exo | /e11-15,e17 | own (e11-15,e17-21 | 1,52 | ENSG00000103043 |
| C17orf76-AS1 | Exon Cassette | e8-9 | up | 1,52 | ENSG00000175061 |
| MPRIP | Exon Cassette | e25 | up | 1,52 | ENSG00000133030 |

| | | | | | |
|---|---|---|---|---|---|
| WSB1 | Alter. First Exon | e1-5/e7 | down (e1-5) | 1,52 | ENSG00000 109046 |
| SPATA20 | Intron Retention | e15 | up | 1,52 | ENSG00000 006282 |
| CAMTA2 | Exon Cassette | e3 | up | 1,52 | ENSG00000 108509 |
| TBC1D16 | Alter. First Exon | e1-5,e7/e6 | down (e1-5,e7) | 1,52 | ENSG00000 167291 |
| ANKRD12 | Exon Cassette | e5 | down | 1,52 | ENSG00000 101745 |
| ZNF519 | Exon Cassette | e6,e7 | up | 1,52 | ENSG00000 175322 |
| MBD1 | Complex | e13/e14 | up (e13) | 1,52 | ENSG00000 141644 |
| ZNF317 | Intron Retention | e3 | up | 1,52 | ENSG00000 130803 |
| DNM2 // QTRT1 | ualy Exclusive Ex | e22/e23 | up (e22) | 1,52 | ENSG00000 079805 // ENSG00000 213339 |
| GTPBP3 | lter. Acceptor Sit | e4 | down | 1,52 | ENSG00000 130299 |
| PM3P9 // ZNF76 | Alter. First Exon | e1,e4-6/e7 | up (e1,e4-6) | 1,52 | ENSG00000 160336 // ENSG00000 241015 |
| BCL2L11 | Exon Cassette | e9,e11 | up | 1,52 | ENSG00000 153094 |
| THUMPD2 | Exon Cassette | e8 | up | 1,52 | ENSG00000 138050 |
| ANKRD36B | Exon Cassette | e4-5 | up | 1,52 | ENSG00000 196912 |
| FHL2 | Complex | e2-4 | down | 1,52 | ENSG00000 115641 |
| STK4 | ter. Terminal Exo | e11,e12,e1 | up (e10) | 1,52 | ENSG00000 101109 |
| GNAS | Complex | e9/e10,e13 | down (e10,e13) | 1,52 | ENSG00000 087460 |
| E1 // NFS1 // RB | ter. Terminal Exo | 22-23/e24-3 | up (e22-23) | 1,52 | ENSG00000 214078 // ENSG00000 244005 // ENSG00000 244462 |
| ZBTB21 | Exon Cassette | e2,e3 | up | 1,52 | ENSG00000 173276 |
| ITFP1 // SEC14L | Complex | e11-14 | up | 1,52 | ENSG00000 100003 // ENSG00000 242114 |
| MCM5 | Intron Retention | e15 | up | 1,52 | ENSG00000 100297 |

| | | | | | |
|---|---|---|---|---|---|
| GCR2 // DGCR1 | Complex | e1/e2 | up (e2) | 1,52 | ENSG00000 070413 |
| TRMT2A | Intron Retention | e3 | up | 1,52 | ENSG00000 099899 |
| TADA3 | Complex | e1/e2-4 | down (e2-4) | 1,52 | ENSG00000 171148 |
| CCNL1 | Intron Retention | e6-8 | up | 1,52 | ENSG00000 163660 |
| NCEH1 | Exon Cassette | e2 | up | 1,52 | ENSG00000 144959 |
| REST | Complex | e5,e8,e11-12 | up | 1,52 | ENSG00000 084093 |
| UGT8 | Alter. First Exon | e1/e2 | down (e1) | 1,52 | ENSG00000 174607 |
| SREK1 | Intron Retention | e11,e12 | up | 1,52 | ENSG00000 153914 |
| SMAD5 | Exon Cassette | e3 | down | 1,52 | ENSG00000 113658 |
| SPOCK1 | Complex | e2,e4/e3 | down (e3) | 1,52 | ENSG00000 152377 |
| MAP3K7 | Exon Cassette | e12 | up | 1,52 | ENSG00000 135341 |
| ATAT1 | Complex | e1 | up | 1,52 | ENSG00000 137343 |
| SNK2B // LY6G5 | Complex | e3-4/e9-10 | down (e3-4) | 1,52 | ENSG00000 204435 // ENSG00000 240053 |
| MRPL32 | Alter. Donor Site | e2 | up | 1,52 | ENSG00000 106591 |
| NUB1 | ter. Terminal Exo | e9/e12-18 | down (e9) | 1,52 | ENSG00000 013374 |
| IRPS24 // URGC | Intron Retention | e5 | up | 1,52 | ENSG00000 062582 // ENSG00000 106608 |
| C7orf49 | Alter. First Exon | e3-4/e5 | down (e3-4) | 1,52 | ENSG00000 122783 |
| FASTK | Intron Retention | e2 | up | 1,52 | ENSG00000 164896 |
| POMT1 | Exon Cassette | e2-3 | up | 1,52 | ENSG00000 130714 |
| AKNA | Alter. First Exon | e1/e3-4 | down (e3-4) | 1,52 | ENSG00000 106948 |
| NSMF | Alter. First Exon | e1-4/e6 | down (e1-4) | 1,52 | ENSG00000 165802 |
| MAP7D2 | Complex | e6,e7-9 | up | 1,52 | ENSG00000 184368 |
| AIFM1 | Alter. First Exon | 1-2,e4-10/e1 | up (e1-2,e4-10) | 1,52 | ENSG00000 156709 |
| RGL1 | Exon Cassette | e6-8 | down | 1,51 | ENSG00000 143344 |

| | | | | | |
|---|---|---|---|---|---|
| GUK1 | Intron Retention | e5 | up | 1,51 | ENSG00000143774 |
| COG2 | Intron Retention | e10 | up | 1,51 | ENSG00000135775 |
| 1A // CDK11B // | Intron Retention | e38 | up | 1,51 | ENSG00000008128 // ENSG00000078369 // ENSG00000248333 |
| TPM3 | Exon Cassette | e14 | up | 1,51 | ENSG00000143549 |
| WDR26 | Exon Cassette | e10 | up | 1,51 | ENSG00000162923 |
| ANKRD26 | Exon Cassette | e7 | up | 1,51 | ENSG00000107890 |
| CD151 | Complex | e1-2 | up | 1,51 | ENSG00000177697 |
| ARHGEF12 | Exon Cassette | e6 | up | 1,51 | ENSG00000196914 |
| ACAD8 | Complex | e10,e11 | up | 1,51 | ENSG00000151498 |
| MRE11A | ter. Terminal Exo | e9/e10-22 | up (e9) | 1,51 | ENSG00000020922 |
| LRRC23 | Complex | e1,e2 | up | 1,51 | ENSG00000010626 |
| UBE3B | Complex | e30 | up | 1,51 | ENSG00000151148 |
| SMEK1 | Exon Cassette | e12 | up | 1,51 | ENSG00000100796 |
| ATXN3 | Complex | 5-6,e8-10/e1 | down (e5-6,e8-10) | 1,51 | ENSG00000066427 |
| ATXN3 | Complex | e6-10/e12 | down (e6-10) | 1,51 | ENSG00000066427 |
| ATXN3 | Exon Cassette | e2,e3,e6-10 | down | 1,51 | ENSG00000066427 |
| MOK | Exon Cassette | e4 | up | 1,51 | ENSG00000080823 |
| PML | ter. Terminal Exo | e4/e5-8 | up (e5-8) | 1,51 | ENSG00000140464 |
| PML | Complex | 2,e3-7,e9-1 | up | 1,51 | ENSG00000140464 |
| WHAMMP3 | Exon Cassette | e9 | down | 1,51 | ENSG00000187667 |
| SLC25A11 | Intron Retention | e3 | up | 1,51 | ENSG00000108528 |
| MYL12B | Alter. First Exon | e1/e2 | down (e1) | 1,51 | ENSG00000118680 |
| ATP9B | Exon Cassette | e11 | up | 1,51 | ENSG00000166377 |
| NFIC | Exon Cassette | e11 | up | 1,51 | ENSG00000141905 |

| | | | | | |
|---|---|---|---|---|---|
| NF587 // ZNF587 | Complex | e3 | down | 1,51 | ENSG00000083828 // ENSG00000152443 // ENSG00000198466 // ENSG00000269343 |
| GIGYF2 | ter. Terminal Exo | e10-13,e17- | up (e8) | 1,51 | ENSG00000204120 |
| KANSL3 | Alter. First Exon | e1/e5 | down (e1) | 1,51 | ENSG00000114982 |
| OBSL1 | ter. Terminal Exo | 9/e10,e11-2 | up (e9) | 1,51 | ENSG00000124006 |
| CBFA2T2 | Exon Cassette | e6-7 | down | 1,51 | ENSG00000078699 |
| ELMO2 | Intron Retention | e23 | up | 1,51 | ENSG00000062598 |
| DIDO1 | ter. Terminal Exo | 7-8/e9,e10-1 | up (e7-8) | 1,51 | ENSG00000101191 |
| NF2 | Exon Cassette | e17 | down | 1,51 | ENSG00000186575 |
| SGSM3 | Alter. First Exon | 1-2,e4-17/e1 | up (e1-2,e4-17) | 1,51 | ENSG00000100359 |
| CHEK2 | Exon Cassette | e7,e8 | down | 1,51 | ENSG00000183765 |
| AGTR1 | Exon Cassette | e2 | down | 1,51 | ENSG00000144891 |
| RFTN1 | Alter. First Exon | 3-6,e8-9/e1 | down (e3-6,e8-9) | 1,51 | ENSG00000131378 |
| CLCN3 | ter. Terminal Exo | /e5,e6-15,e | lown (e5,e6-15,e17 | 1,51 | ENSG00000109572 |
| CUL7 | Alter. First Exon | e1-10/e11 | down (e1-10) | 1,51 | ENSG00000044090 |
| ZNRD1-AS1 | Alter. First Exon | e1/e3 | down (e1) | 1,51 | ENSG00000204623 |
| CUTA | Complex | e1,e2 | down | 1,51 | ENSG00000112514 |
| GTF2IRD1 | Intron Retention | e27 | up | 1,51 | ENSG00000006704 |
| RHBDD2 | Exon Cassette | e2 | down | 1,51 | ENSG00000005486 |
| NSUN5 | Intron Retention | e9,e10 | up | 1,51 | ENSG00000130305 |
| C8orf59 | Exon Cassette | e2 | up | 1,51 | ENSG00000176731 |
| ZNF706 | Alter. First Exon | e2/e4 | down (e2) | 1,51 | ENSG00000120963 |
| SEC16A | Exon Cassette | e33 | up | 1,51 | ENSG00000148396 |
| SYP | Intron Retention | e2 | up | 1,51 | ENSG00000102003 |

| | | | | | |
|---|---|---|---|---|---|
| MORF4L2 | Exon Cassette | e4-5 | up | 1,51 | ENSG00000123562 |
| CHTOP | ter. Terminal Exo | e3/e4-6 | up (e3) | 1,5 | ENSG00000160679 |
| YY1AP1 | Exon Cassette | e4-5 | up | 1,5 | ENSG00000163374 |
| HINFP | Intron Retention | e4,e5 | up | 1,5 | ENSG00000172273 |
| GANAB | Exon Cassette | e6 | up | 1,5 | ENSG00000089597 |
| XRRA1 | Exon Cassette | e3 | down | 1,5 | ENSG00000166435 |
| IRAK4 | Exon Cassette | e2,e3-5 | up | 1,5 | ENSG00000198001 |
| SRGAP1 | ter. Terminal Exo | 0/e11,e13-2 | up (e10) | 1,5 | ENSG00000196935 |
| WARS | Alter. First Exon | e1/e2,e3-4 | down (e1) | 1,5 | ENSG00000140105 |
| OCA2 | Exon Cassette | e10 | down | 1,5 | ENSG00000104044 |
| EHD4 | Complex | e1-4 | up | 1,5 | ENSG00000103966 |
| C17orf76-AS1 | Exon Cassette | e7-9 | up | 1,5 | ENSG00000175061 |
| PRKAR1A | Alter. Donor Site | e3 | up | 1,5 | ENSG00000108946 |
| CSNK1D | Complex | e1-3 | up | 1,5 | ENSG00000141551 |
| TGIF1 | Alter. First Exon | e4-8/e7 | down (e7) | 1,5 | ENSG00000177426 |
| TGIF1 | Alter. First Exon | e7/e10 | down (e7) | 1,5 | ENSG00000177426 |
| CLASRP | Alter. First Exon | e2-11/e12 | up (e2-11) | 1,5 | ENSG00000104859 |
| CLASRP | Alter. First Exon | 1-3,e5-11/e1 | up (e1-3,e5-11) | 1,5 | ENSG00000104859 |
| SYT3 | Alter. First Exon | e1-2/e3 | up (e1-2) | 1,5 | ENSG00000213023 |
| ELMOD3 | Exon Cassette | e3 | up | 1,5 | ENSG00000115459 |
| OBSL1 | ter. Terminal Exo | 0,e12-15,e1 | up (e9) | 1,5 | ENSG00000124006 |
| E1 // NFS1 // RB | Complex | e21,e22 | up | 1,5 | ENSG00000214078 // ENSG00000244005 // ENSG00000244462 |
| DAG1 | Complex | e1 | down | 1,5 | ENSG00000173402 |

| | | | | | |
|---|---|---|---|---|---|
| C31A // THAP9-A | ter. Terminal Exo | ,e28-29,e31 | (e3,e8-26,e28-29,e3 | 1,5 | ENSG00000138674 // ENSG00000251022 |
| ZSCAN16 | ter. Terminal Exo | e3/e4 | up (e3) | 1,5 | ENSG00000196812 |
| RNF146 | Complex | e1 | up | 1,5 | ENSG00000118518 |
| HYMAI // PLAGL1 | Exon Cassette | e8 | down | 1,5 | ENSG00000118495 |
| MPP6 | Exon Cassette | e2 | down | 1,5 | ENSG00000105926 |
| PRKAG2 | Complex | e7/e8 | up (e7) | 1,5 | ENSG00000106617 |
| VPS13B | Exon Cassette | e31 | down | 1,5 | ENSG00000132549 |
| COQ4 | Alter. First Exon | e2-4/e6 | down (e2-4) | 1,5 | ENSG00000167113 |
| --- | Intron Retention | e6,e7 | up | 1,5 | --- |
| MORF4L2 | Exon Cassette | e4,e5 | up | 1,5 | ENSG00000123562 |
| BCAP31 | Complex | e2,e3 | down | 1,5 | ENSG00000185825 |
