## Supplementary material for "Regulation of neuronal mRNA splicing and Tau isoform ratio by ATXN3 through deubiquitylation of splicing factors": Table S3

| Pathway description (KEGG) | Number of genes in the pathway | Number of altered genes | p value |
|---|---|---|---|
| Nucleotide excision repair | 44 | 21 | 1.24E-04 |
| Adherens junction | 77 | 30 | 1.40E-04 |
| Spliceosome | 126 | 42 | 1.93E-04 |
| Ubiquitin mediated proteolysis | 137 | 43 | 3.16E-04 |
| Endometrial cancer | 52 | 21 | 5.37E-04 |
| Neurotrophin signaling pathway | 124 | 38 | 1.23E-03 |
| Axon guidance | 129 | 39 | 1.39E-03 |
| Pathogenic Escherichia coli infection | 57 | 22 | 2.03E-03 |
| Endocytosis | 184 | 52 | 3.36E-03 |
| Prostate cancer | 89 | 30 | 4.20E-03 |
| RNA degradation | 57 | 20 | 5.01E-03 |
| Ribosome | 87 | 28 | 6.12E-03 |
| Pathways in cancer | 328 | 79 | 7.35E-03 |
| Focal adhesion | 201 | 52 | 1.25E-02 |
| MAPK signaling pathway | 267 | 64 | 1.79E-02 |
| Apoptosis | 87 | 25 | 2.26E-02 |
| VEGF signaling pathway | 75 | 22 | 2.70E-02 |
| Colorectal cancer | 84 | 24 | 2.75E-02 |
| Pancreatic cancer | 72 | 21 | 3.30E-02 |
| Type I diabetes mellitus | 42 | 17 | 3.45E-02 |
| Fc gamma R-mediated phagocytosis | 95 | 27 | 3.53E-02 |
| Selenoamino acid metabolism | 26 | 10 | 3.71E-02 |
| Alzheimer's disease | 163 | 40 | 4.49E-02 |
