## Supplementary material for "Regulation of neuronal mRNA splicing and Tau isoform ratio by ATXN3 through deubiquitylation of splicing factors": Table S4

| | Up | Down | Up | Down |
|---|---|---|---|---|
| Alter. First exon | 380 | 362 | 51% | **49%** |
| Alter. Terminal exon | 165 | 329 | 33% | **67%** |
| Exon cassete | 438 | 810 | 35% | **65%** |
| Mutually exclusive exons | 11 | 26 | 30% | **70%** |
| Alter. Acceptor splice site | 20 | 53 | 27% | **73%** |
| Alter donor splice site | 22 | 50 | 31% | **70%** |
| Intron retention | 42 | 396 | 10% | **90%** |
| Complex | 230 | 350 | 40% | **60%** |
